# Food supplementation with molybdenum complexes improves honey bee health

**DOI:** 10.1101/2025.02.25.640117

**Authors:** Arcadie Fuior, Loic Colin, Valentina Cebotari, Amélie Noel, Isabelle Ribaud, Isabelle Gerard, Olga Garbuz, Mathieu Fregnaux, Xavier López, Virginie Larcher, Michael A. Shestopalov, Anastasiya O. Solovieva, Tatiana N. Pozmogova, Olesea Gliga, Nadejda Railean, Précillia Cochard, Benjamin Poirot, Leonidas Charistos, Fani Hatjina, Andrea Somogyi, Kadda Medjoubi, Sébastien Gaumer, Aurelian Gulea, Ion Toderas, Jean-Christophe Sandoz, Sébastien Floquet

**Affiliations:** Institut Lavoisier de Versailles, Université Paris-Saclay, UVSQ, CNRS, UMR 8180, 78000 Versailles, France; State University of Moldova, MD-2009 Chisinau, Republic of Moldova; Evolution Genomes Behaviour & Ecology, University Paris-Saclay, CNRS, IRD, 91198 Gif Sur Yvette, France; Institute of Zoology, MD-2028 Chisinau, Republic of Moldova; Université Paris Saclay, CNRS, IN2P3, IJCLab, 91403 Orsay, France; Universitat Rovira i Virgili, Departament de Química Física i Inorgànica, Marcel·lí Domingo 1, 43007 Tarragona, Spain; Nikolaev Institute of Inorganic Chemistry SB RAS, 630090 Novosibirsk, Russia; Research Institute of Clinical and Experimental Lymphology – Branch of the ICG SB RAS, 630090 Novosibirsk, Russia; Novosibirsk State University, 630090 Novosibirsk, Russia; Apinov S.A.S, 17140 Lagord (La Rochelle), France; Department of Apiculture, Institute of Animal Science, ELGO ‘DIMITRA’, Nea Moudania, 63200 Greece; Nanoscopium beamline, Synchrotron SOLEIL, L’Orme des Merisiers, Saint-Aubin, 91192 Gif-sur-Yvette, France; Genetics and Cell Biology Laboratory, Université Paris-Saclay, UVSQ, UR 4589, 78180 Montigny-le-Bretonneux, France

**Keywords:** Honeybees, Nutrition, Molybdenum complexes, Trace element

## Abstract

In this study, we evaluated the impact of supplementing honey bee feed with molybdenum-based compounds. Among a dozen of dinuclear Mo(V) complexes, we first selected the most stable and non-toxic complexes, which were tested as food supplements in an extensive eight-year campaign involving more than 700 beehives across various environmental conditions in Moldova, France, Greece and USA. This unprecedented field campaign revealed that a few milligrams of compounds **Na-Mo**_**2**_**O**_**4**_**-EDTA** or **Li-Mo**_**2**_**O**_**4**_**-EDTA** provided in spring or autumn, increased queen fecundity, hygienic behavior, and honey production, while infestation rates of worker bees and brood by the mite *Varroa destructor*, and mortality rates in winter were dramatically reduced. Hive monitoring showed that the Mo-containing syrup can be consumed over 1.5 months and is well assimilated by larvae and workers within the hive. In particular, Mo levels increased significantly in the head of the bees. X-ray fluorescence measurements demonstrated that **Na-Mo**_**2**_**O**_**4**_**-EDTA** increases Mo levels in brain, neurolemma and hypopharyngeal glands. The metabolism of Mo complexes was addressed using X-Ray photoelectron spectroscopy (XPS) on bee faeces, which revealed that the complexes are oxidized into Mo(VI) species and thus suggesting that Mo complexes may function as antioxidant agents in bees. These findings offer promising solutions for the beekeeping industry, which is struggling with weakening honey bee colonies.

**Significance Statement:** Honey bees play a crucial role as pollinators. Unfortunately, colony losses have increased significantly over the past decades through the action of multiple environmental stressors. Currently, solutions are sought to strengthen bees’ health and resilience. This study shows that feeding bees with small amounts of trace elements like molybdenum can improve honey bee condition and increase the production of hive products. Through an unprecedented series of field and lab tests it notably demonstrates that molybdenum complexes are i) non-toxic, ii) consumed by bees over several generations, iii) assimilated particularly in the brain and hypopharyngeal glands, iv) acting as antioxidant. These results open new avenues for using trace elements to improve pollinator health.

## 1. Introduction

About 80% of all cultivated plant species and 40% of our diet directly depend on pollination [1]. The economic impact of insect pollinators, especially bees, has thus been estimated to 150 billion euros in 2005 [2]. While the honey bee *Apis mellifera* plays a major role in insect-mediated pollination, it is gravely endangered by colony losses [3], which have systematically increased in the last decades through a multifactorial process involving biological agents (*Varroa destructor, Vairimorpha*, etc.), the use of pesticides, and a lack of resources among others [4]. In this context, the development of new solutions to protect bees from these external aggressions and strengthen their health and resilience is therefore of great interest.

One way to improve honey bees’ health and resilience to stress is to optimize their nutrition. This is a highly complex issue as it requires a perfect balance between macronutrients (carbohydrates, proteins, and lipids) and micronutrients (vitamins, minerals). Floral sources are honey bees’ major source of macronutrients. Pollen supplies bees with lipids and proteins, while nectar provides them with carbohydrates. Micronutrients, and especially metals, are also important for animals’ health as they play an essential role in numerous physiological processes. Surprisingly, micronutrients have been poorly studied in honey bees [5]. Honey bees likely obtain micronutrients, vitamins and minerals from two main sources: pollen and “dirty” or turbid water. Yet, very little is known about the metals that are needed for honey bees’ health [5]. Several studies have been focused on the physiological role and/or toxicity of elements like Fe, Mn, Cu and Zn, as well as the deleterious effects of pollution by heavy elements such as As, Cd, and Pb [6-10]. However, trace elements like molybdenum have been generally overlooked, leaving us with limited knowledge of their potential role or of the consequence of a deficiency in these elements.

Molybdenum (Mo) is a trace element of essential importance for life in almost all living organisms from bacteria to humans [11]. Mo is present in about fifty enzymes, namely molybdoenzymes, which are essential constituents of the global carbon, sulfur and nitrogen metabolism in plants and animals. In these enzymes, Mo atoms play a key role in the catalysis of diverse redox reactions thanks to their oxidation state which can vary from Mo(+VI) to Mo(+IV) [11, 12]. Molybdoenzymes are generally classified into different families. In eukaryotes, these enzymes mainly belong to the Sulfite Oxidase (SO) and Xanthine Oxidase (XO) families [12]. The role of SO is important for sulfites detoxification. XO molybdoenzymes are very useful for the degradation of xenobiotics and possess a broad substrate spectrum. As we demonstrate here, Mo is also present in honey bees, with an average content of about 0.4 ppm (see Part I, SI) but to our knowledge its role in this insect is utterly unknown. To fill this gap, we evaluated the impact of supplementing the bee feed with Mo-based compounds in an extensive eight-year campaign in Moldova, France, Greece and USA. For this, we chose simple coordination complexes formed by association of a Mo-based cluster with organic ligands, as this should facilitate their uptake by bees and rapidly involve redox processes due to the reduced oxidation state of the Mo core.

## 2. Results and discussion

### 2.1 Selection of Mo-complexes for the beekeeping tests

Numerous dinuclear Mo complexes combining the [Mo_2_O_4_]^2+^ and [Mo_2_O_2_S_2_]^2+^ clusters with organic ligands, such as EDTA or Cysteine (L-Cys), have emerged in the literature as biomimetic models of molybdoenzymes since the 1970’s [13]. Quite recently, it was demonstrated that these complexes have little or no toxicity [14,15] and some of them can be used to detoxify cyanides [16], are capable of penetrating animal cells [17] and can stimulate biomass production or act as excellent antioxidants [15]. In particular, Floquet et al. studied a dozen complexes of general formula (cation)_x_[Mo_2_O_2_E_2_(L)_y_] (x = 0, 2, 4; y = 1 or 2, E = O or S), hereafter referred to as **Cation-Core-Ligand**, where the ligands L are commercial ligands and the counter-cations are alkali or organic cations [15]. The aim of the first step of this study was to select the most promising complexes among them in terms of chemical stability (Part II, SI) and lack of toxicity (Part III, SI) when used in honey bees’ food. This process led to the selection of 3 complexes: [Mo_2_O_4_(EDTA)]^2-^, [Mo_2_O_2_S_2_(EDTA)]^2-^, and [Mo_2_O_2_S_2_(L-Cys)_2_]^2-^ (**Figure 1**). [Mo_2_O_4_(EDTA)]^2-^ will be used as PPh4+, Na^+^ and Li^+^ salts (**PPh4-Mo2O4-EDTA, Na-Mo2O4-EDTA** and **Li-Mo2O4-EDTA**), the sulphurated analogue [Mo2O2S2(EDTA)]^2-^ will be used as PPh^4+^ salt (**PPh**_**4**_**-Mo**_**2**_**O**_**2**_**S**_**2**_**-EDTA**) and [Mo_2_O_2_S_2_(LCys)_2_]^2-^ will be used as K^+^ salt (**K-Mo**_**2**_**O**_**2**_**S**_**2**_**-LCys**).

**Figure 1.**
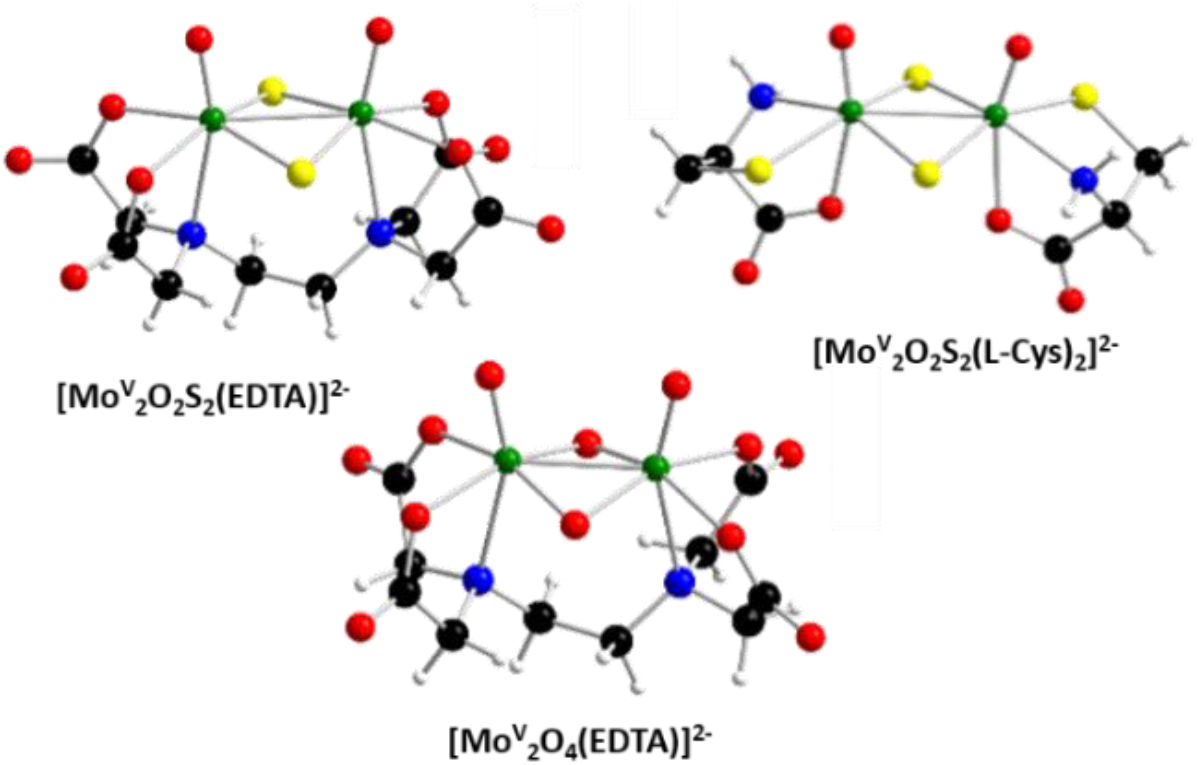
Structures of the three complexes tested in beehives. Color code: Mo in green, S in yellow, O in red, N in blue, C in black, H in white

### 2.2 Tests in beehives

#### Choice of the most efficient complex in beehives

The first two field test campaigns were performed in Moldova (forest environment 20 km outside the region of Chisinau) on 48 honey beehives of the *carpatica* ecotype of *Apis mellifera carnica*. The colonies were fed at the beginning of spring every second day during a period of two weeks, with sugar syrup containing **PPh**_**4**_**-Mo**_**2**_**O**_**4**_**-EDTA, PPh**_**4**_**-Mo**_**2**_**O**_**2**_**S**_**2**_**-EDTA** or **K-Mo**_**2**_**O**_**2**_**S**_**2**_**-L-Cys** complexes for a total dose of 2 mg of complex per hive and compared to a control group fed only with syrup. More details and all raw data are available in Part IV of the SI. Among the three complexes the best and most significant results were obtained with **PPh**_**4**_**-Mo**_**2**_**O**_**4**_**-EDTA** (Table SIV.1, SI), so that sulphurated complexes were later discarded. Feeding with **PPh**_**4**_**-Mo**_**2**_**O**_**4**_**-EDTA** induced a significant increase in queen fecundity (+10.5%, Mann-Whitney U test, p = 1.5 10^−7^), hygienic behavior (+4.3 %, U-test, p = 8.8 10^−8^), bee bread production (fermented stored pollen; +21.8%, U-test, p = 8.8 10^−5^), honey production (+19.6%, U-test, p = 6.7 10^−6^) and wax production by +39% (U-test, p = 6.7 10^−6^) [18].

#### Choice of the most efficient salts of complex [Mo_2_O_4_(EDTA)]^2-^

Two more field campaigns were performed in Moldova to evaluate the influence of the counter cations PPh^4+^, Na^+^ and Li^+^ associated to the complex [Mo_2_O_4_(EDTA)]^2-^, applying 2 mg of each compound per beehive. In addition to the previously monitored parameters, infestation by the mite *Varroa destructor*, a major pest for the honey bee, was also evaluated. All data are presented in Part IV of the SI (Table SIV.2), while the most significant results are depicted in Figure 2. Queen fecundity increased by +13.3 % and +11.7 % for **Na-Mo**_**2**_**O**_**4**_**-EDTA** and **Li-Mo**_**2**_**O**_**4**_**-EDTA** but not significantly compared with the control (Kruskal Wallis test, p = 0.574). Conversely, the three complexes induced a decrease in the infestation of bee workers by the mite *Varroa destructor* (Kruskal Wallis test, p < 0.001). The lithium salt **Li-Mo**_**2**_**O**_**4**_**-EDTA** had the strongest effect (−42.9% on worker bees, multi comparison Dunn’s test, p < 0.001), while **PPh**_**4**_**-Mo**_**2**_**O**_**4**_**-EDTA** and **Na-Mo**_**2**_**O**_**4**_**-EDTA** salts produced less pronounced and non-significant decreases (respectively -21.3% / p = 0.36 and -13.5 % / p = 1). Finally, honey production increased by +13.0% for PPh_4_-Mo_2_O_4_-EDTA (p = 1), by +42.9 % for **Li-Mo**_**2**_**O**_**4**_**-EDTA** (Kruskal Wallis test withal data, p = 0.41; U-test compared to control, p = 0.03), and up to +49.6 % for **Na-Mo**_**2**_**O**_**4**_**-EDTA** (Kruskal Wallis test with all data, p = 0.13; U-test compared to control, p = 0.05), which translated in an average honey production of 31.1 ± 3.0 kg per beehive for the later *vs* 20.8 ± 3.5 kg for the control group. Mo feeding did not affect honey quality. In particular, no trace of Mo was found in the produced honey and all measured physical parameters fully comply with European Union regulations (Table SIV.3, Part IV, SI). In summary, these test campaigns confirmed the two alkali salts **Li-Mo**_**2**_**O**_**4**_**-EDTA**, and **Na-Mo**_**2**_**O**_**4**_**-EDTA**, as the two most promising complexes. The most significant results of this campaign are depicted in Figure 2A.

**Figure 2.**
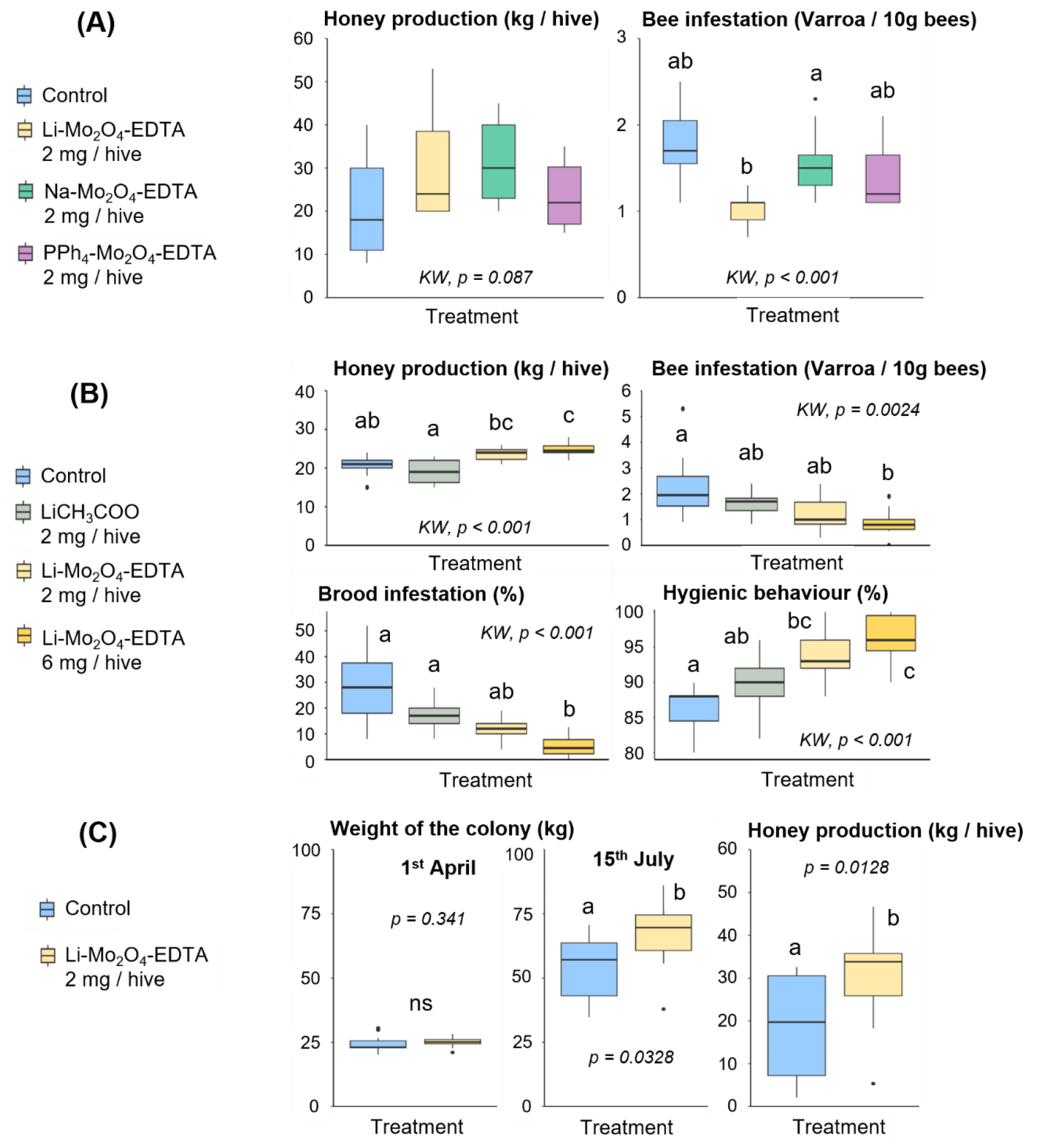
Representative results obtained in the course of 3 test campaigns on honey bee colonies. (A) Effect of Na-Mo_2_O_4_-EDTA (2 mg/hive), Li-Mo_2_O_4_-EDTA (2 mg/hive) and PPh_4_-Mo_2_O_4_-EDTA (2 mg/hive) on honey production and worker bees infestation by Varroa from the field test in Moldova (2018) (B) Effect of lithium acetate or Li-Mo_2_O_4_-EDTA at 2 and 6 mg/hive on worker bees and brood infestation by Varroa, hygienic behaviour and honey production from the field test in Moldova (2019); (C) Effect of Li-Mo_2_O_4_-EDTA at 2 mg/hive on colony weight and honey production from the field test in France (2019). For each campaign, n=10 hives per treatment. Letters indicate significant differences between treatments (p<0.05) based on the Dunn’s post hoc test with Bonferroni correction (A and B) or Mann-Whitney U test (C). Boxplots show median (horizontal crossbar) and interquartile ranges. KW : Kruskal-Wallis.

#### Effects against Varroa infestation

Lithium cations are known to have a deleterious effect on *Varroa* [19]. A further campaign in Moldova (2019) focused on the impact of the **Li-Mo**_**2**_**O**_**4**_**-EDTA** complex on *Varroa* infestation of worker bees and larvae. The complex was used at two dosages (2 and 6 mg per beehive), in comparison with a control group, and a group treated with lithium acetate as reference (LiOAc, 2 mg/beehive). The results are gathered in Table SIV.4 (Part IV, SI) and the most significant results are depicted in Figure 2B.

The treatment had a significant effect on the worker bee infestation rate by Varroa (Kruskal Wallis test, p = 0.0024). With an overall dose of 2 mg of **Li-Mo**_**2**_**O**_**4**_**-EDTA** per beehive, infestation of worker bees by *Varroa* fell by -47% compared to the control group (multi comparison Dunn’s test, p = 0.080), similarly to the previous campaign (−42.9%). This effect increased to -61.3% with 6 mg per beehive (Dunn’s test, p = 0.002). Despite a Lithium concentration 3.7 times higher, the effect of lithium acetate was weaker and non-significant (−30.4%, p = 1). It could suggest that the Mo-complex in **Li-Mo**_**2**_**O**_**4**_**-EDTA** has a protective action against *Varroa*, in synergy with Li^+^ cations. Interestingly, the hygienic behavior of hives fed with 2 and 6 mg of **Li-Mo**_**2**_**O**_**4**_**-EDTA** concomitantly increased by +9.3% (p = 0.007) and +12 % (p = <0.001), respectively. It is known that honey bee colonies with higher hygienic behavior tend to keep varroa infestation at lower levels. [20]

We also observed a significant effect of the treatment on the brood infestation rate by Varroa (Kruskal Wallis test, p < 0.001). Interestingly, *Varroa* infestation was very high in control hives, *i*.*e*. 28.2%. After feeding with the reference compound LiOAc (2 mg/hive), this rate is lower by -35.1% for this group (Dunn’s test, p = 1). To date no substance appears capable of protecting bee brood against *Varroa*. Again, the **Li-Mo**_**2**_**O**_**4**_**-EDTA** complex produced stronger effects, reducing brood infestation by -57.4% compared to the control group with 2 mg (Dunn’s test, t, p = 0.067), and up to -81.6% with 6 mg per beehive (Dunn’s test, p = <0.001). Beyond this result, it also suggests that the complex is consumed not only by adult workers, but is also transferred to the brood.

#### Test under operating conditions

The feeding protocol used in previous campaigns is not applicable in professional beekeeping conditions. A test campaign was thus carried out in France, using *Apis mellifera “Buckfast”* bees, which are widely used by professional beekeepers. The test was performed on 22 beehives in Gif-sur-Yvette (France), split into two groups: 11 control and 11 test colonies. For the test colonies, 2 mg of **Li-Mo**_**2**_**O**_**4**_**-EDTA** complex was introduced in April in only one syrup feeding (0.5 L of syrup at 4 mg/L). We monitored colony weight as a function of time and the quantity of honey produced on 15^th^ July (Table SIV.5, Part IV, SI). The results depicted in Figure 2C show that a single dose of **Li-Mo**_**2**_**O**_**4**_**-EDTA** complex introduced at the beginning of spring resulted in a +23.6% increase in colony weight on July 15 compared with the reference batch (U-test, p = 0.033), and in a +58.7 % increase in honey production (30.3 kg on average for the test batch versus 19.1 kg for the control beehives (U-test, p = 0.014). This result is comparable to that obtained in the second campaign in Moldova with 2 mg of the **Li-Mo**_**2**_**O**_**4**_**-EDTA** given every second day over a period of two weeks (+47%, U-test compared to control group, p = 0.03). This suggests that the syrup is consumed over several weeks. More importantly, this experiment validated the efficacy of our complexes when applied in a single feeding, in accordance with professional beekeepers’ practices and confirmed the efficacy of **Li-Mo**_**2**_**O**_**4**_**-EDTA** in a second geographical location and on another honey bee subspecies.

#### Impact on winter colony mortality

During the autumn-winter period, colony losses can be very high, which is a major issue for professional beekeepers. Two test campaigns were carried out during the winters of 2019 and 2020 on apiaries in the San Francisco area (USA), where reported colony losses are particularly important [21]. In a first campaign, 151 beehives were distributed across 6 different apiaries. In each apiary, the colonies were randomly divided into a control group (76 hives) and a group receiving the **Li-Mo**_**2**_**O**_**4**_**-EDTA** complex (75 hives). At the end of October 2019, the colonies were fed with 1 US Gallon (3.78 liters) of sugar syrup. For the test group, 4 mg of **Li-Mo**_**2**_**O**_**4**_**-EDTA** per beehive were added. The surviving colonies were counted on 7^th^ January 2020. In the control group, 47 colonies were lost out of 76. *i*.*e*. 61.8% winter mortality, while after feeding with 4 mg **Li-Mo**_**2**_**O**_**4**_**-EDTA** only 26 colonies were lost, *i*.*e*. 34.7% winter mortality. This -43.8% mortality drop was significant (chi^2^ test, p = 0.0008).

A second campaign (winter 2020) involved 220 beehives, distributed across 11 different apiaries. The beehives were divided into 4 groups of 55 colonies each fed with 1 US gallon of syrup in September and in October: beehives fed with sugar syrup only (control), beehives receiving 8 mg **Li-Mo**_**2**_**O**_**4**_**-EDTA** in September and sugar syrup in October, beehives fed with syrup in September and 8 mg **Li-Mo**_**2**_**O**_**4**_**-EDTA** in October, and beehives fed with 4 mg **Li-Mo**_**2**_**O**_**4**_**-EDTA** both in September and in October. Thus, the three test colonies received 8 mg of the complex and only differed in the schedule of **Li-Mo**_**2**_**O**_**4**_**-EDTA** treatment. Lost colonies were counted on December 31^st^, 2020. The period of the feeding strongly and significantly impacted the results. The mortality rate in the control group was 15/55, *i*.*e*. 27.3% loss and feeding with the complex in October only reduced mortality by -33% compared with the control group (10/55, *i*.*e*. 18.2% loss), in agreement with the -44% reduction obtained the previous year, although it was not significant (Chi^2^ test, p = 0.25). However, feeding in September or in September and October greatly reduced colony mortality: indeed, no colonies were lost when feeding in September only (Chi^2^ test, p = 3.1 10^−5^), while 3 colonies out of 55 (5.5% loss, Chi^2^ test, p = 0.002) were lost with feeding in September and in October, representing an 80% reduction in winter mortality compared to the control group.

#### Is there a maximum dose?

In previous campaigns, colonies were given amounts of Mo-complexes ranging from 2 to 30 mg over several weeks or months. The question obviously arose as to the maximum quantities that could be used without causing harm. In a test campaign, carried out from December 2020 to April 2021 in Nea Moudania region, Greece, we focused on the **Li-Mo**_**2**_**O**_**4**_**-EDTA** complex (See section IV.4 of the part IV, SI, for more details). Thirty beehives were divided into 3 batches: a control batch, a batch receiving complex in the form of candy in December (40 mg the **Li-Mo**_**2**_**O**_**4**_**-EDTA** per beehive, “MoLi-B” batch) and a batch receiving 80 mg of **Li-Mo**_**2**_**O**_**4**_**-EDTA** in the form of candy in December and then in syrup in March (“MoLi-A” batch). The populations of the colonies were followed throughout this period (Part IV, SI). During the first period (December to March), we found that feeding the bee colonies with high **Li-Mo**_**2**_**O**_**4**_**-EDTA** quantities (colonies from both MoLi-A and MoLi-B groups) had deleterious effects on the colonies. On average, about 185 dead bees were found in front of the colonies in these groups, compared to about 130 for the control group (p = 0.0001). This effect appeared more pronounced during the second period (March to April) in the “MoLi-A” group (160 dead bees/hive *vs* 97 dead bees/hive for the control group, p = 0.0001), while the increased mortality stopped for the MoLi-B group receiving syrup only (57 dead bees/hive for the second period). This result contrasts with the chronic and acute toxicity studies (see part III, SI) in laboratory conditions. This difference may relate to the presence of a wider range of stressors in natural beehives compared to laboratory conditions. Because of this negative effect, we decided to focus on **Na-Mo**_**2**_**O**_**4**_**-EDTA** for further studies for which such deleterious effects were not observed (see below).

### 2.3 Tracking molybdenum in beehives

Feeding beehives with **Na-Mo**_**2**_**O**_**4**_**-EDTA** showed positive effects on the colonies over several months, although feeding was applied over a short period of time. To understand this long-term effect, we tracked Mo in honey bee colonies for 2 months after the feeding (May-June 2022). Five groups of 8 beehives were each fed with 4 L of syrup (2 times 2L with 1 week between the two feeding) containing 0, 1, 2, 4 or 20 mg/L of **Na-Mo**_**2**_**O**_**4**_**-EDTA**, which corresponds to a global feeding with 0, 4, 8, 16 and 80 mg of complex per beehive, respectively. The Mo-contents were measured in worker bees, in larvae, and in the food stored in the brood frames (see Figure 3A). Mortality was also monitored for all batches over 2 months thanks to Gary traps positioned in front of the hives. No significant difference in mortality was measured between **Na-Mo**_**2**_**O**_**4**_**-EDTA** treated and control hives, even at the highest dose (80 mg, p = 0.57, see Figure SIII.24, Part III, SI).

**Figure 3.**
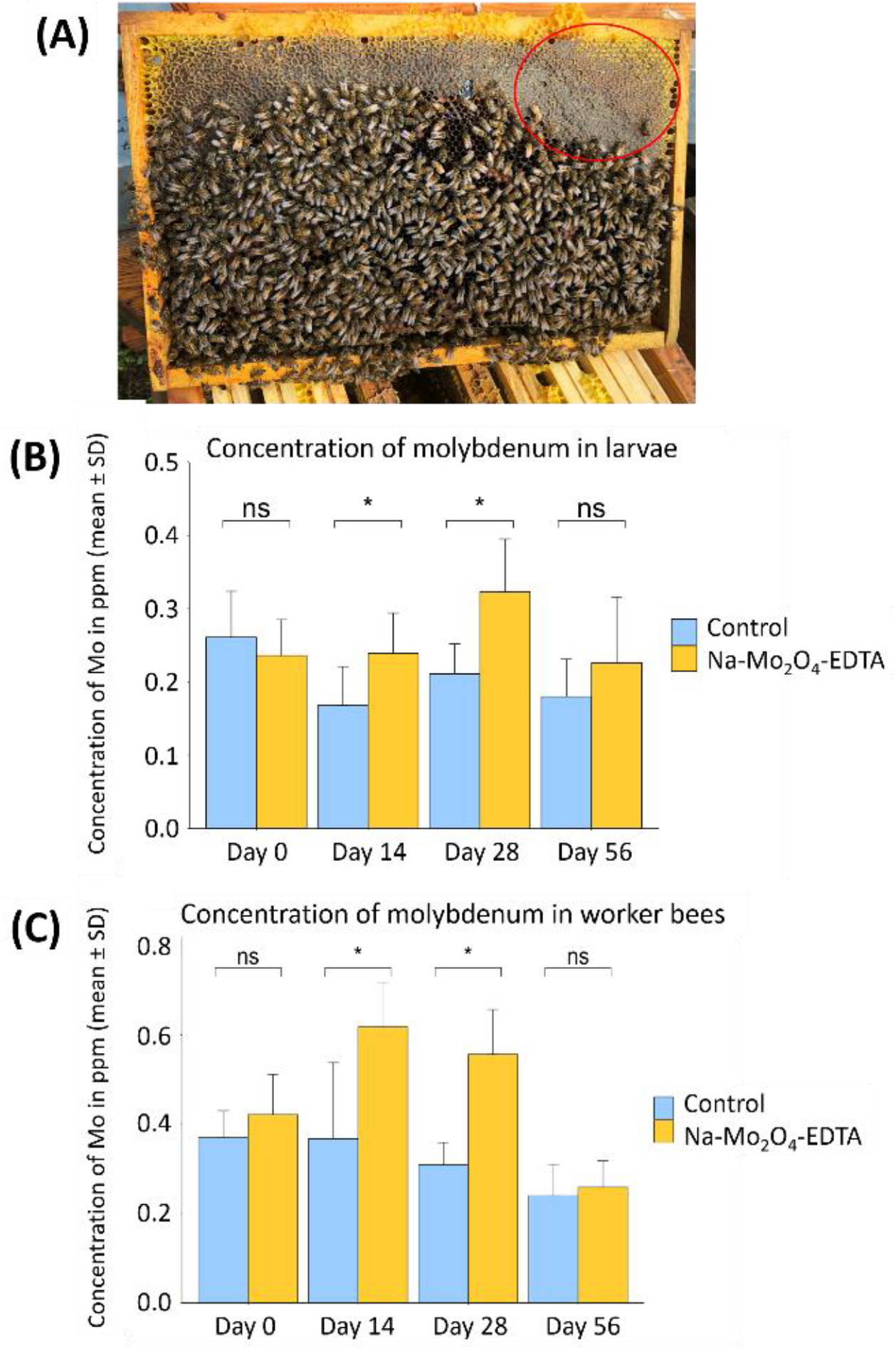
(A) A typical frame found in the experimental beehives. The red circle indicates the zone where food is stored and where the samples are taken; (B) Mo content in larvae in μg/g sampled at D0, D14, D28, D56 for the control group and the group treated with **Na-Mo**_**2**_**O**_**4**_**-EDTA** at 8 mg/beehive; (C) Mo content in worker bees in ppm sampled at D0, D14, D28, D56 for the control group and the group treated with **Na-Mo**_**2**_**O**_**4**_**-EDTA** at 8 mg/beehive. Non-parametric U-tests were performed. Differences at D0 and D56 are not significant (ns), while Mo-contents significantly differ between groups for larvae and worker bees at D14 and D28 (* : p < 0.05).

At Day 0 (hence D0), the level of Mo in the colonies’ food stores was below 0.1 ppm in all groups (Table SV.1, Part V, SI). At D28 and D42 the Mo-content closely matched the levels of Mo in the syrups given to the five test batches, evidencing that the syrup collected by the bees was stored around the brood. At D56, the Mo level was again below 0.1 ppm in almost all batches. These samples corresponded this time to honey produced and stored by the bees and not to any stored syrup, as confirmed by ^1^H NMR studies evidencing the absence of disaccharides. This result demonstrates that after feeding the colonies, the syrup is stored within the beehives and consumed over several weeks by the colonies, over 1 month and a half for young colonies in our case. This spread-out consumption enables several generations of bees to be fed with **Na-Mo**_**2**_**O**_**4**_**-EDTA**. Figures 3B and 3C show the variations in Mo-contents in bee larvae and in worker bees between D0 and D56 for control and 8 mg/hive batches. In bee larvae, at D0, Mo levels were similar in the control and 8 mg/beehive groups: 0.24 *vs* 0.26 ppm. After two weeks (D14) a difference of +42% in Mo level was measured in the 8 mg/hive group (U-test, p = 0.02). This difference increased again at D28, reaching +53% Mo (p = 0.004). The syrup stored within the beehives is thus consumed and assimilated by bee larvae, which provokes a significant increase, especially at D28. At D56, the difference between the two groups decreased and was not significant (p = 0.36).

In workers, Mo levels were also similar at D0 in the control and 8 mg/beehive groups: around 0.4 ppm. At D14, the Mo level increased by +68% in the 8 mg/hive group (0.62 ppm, U-test, p = 0.01), reaching +80% at D28 (p = 0.001). The workers consumed the syrup and assimilated Mo. At D56, Mo levels were again similar in both groups (p = 0.46).

These experiments demonstrate that i) the syrup containing **Na-Mo**_**2**_**O**_**4**_**-EDTA** is consumed by several generations of bees, ii) **Na-Mo**_**2**_**O**_**4**_**-EDTA** is assimilated by both larvae and worker bees within beehives and iii) feeding hives with 80 mg or less of **Na-Mo**_**2**_**O**_**4**_**-EDTA** does not increase mortality.

### 2.4 Effect of Chronic feeding of bees in laboratory conditions

A possible explanation for an increased honey production in beehives fed with Mo-complexes would be a higher longevity of the bees. Laboratory experiments were thus conducted to evaluate the effects of chronic feeding with Mo complexes. Three different concentrations were used in addition to a control group: 2, 20 and 400 mg/L for both **Na-Mo**_**2**_**O**_**4**_**-EDTA** and **Li-Mo**_**2**_**O**_**4**_**-EDTA** complexes. For each group, 3 cages of 50 workers collected at emergence were fed *ad libitum* with water and with Mo-enriched sugar solution, and survival was monitored throughout (Figure 4A). The survival curves are depicted in Figure 4B and 4C (see also section III.4, Part III, SI). For both complexes, three distinct periods can be observed: The first phase, before 30 days, with very little mortality (<10%). In the second phase, between 30 and 60 days, mortality increased. Then, after 60-70 days, when populations have been strongly reduced, the mortality rate decreased again.

**Figure 4.**
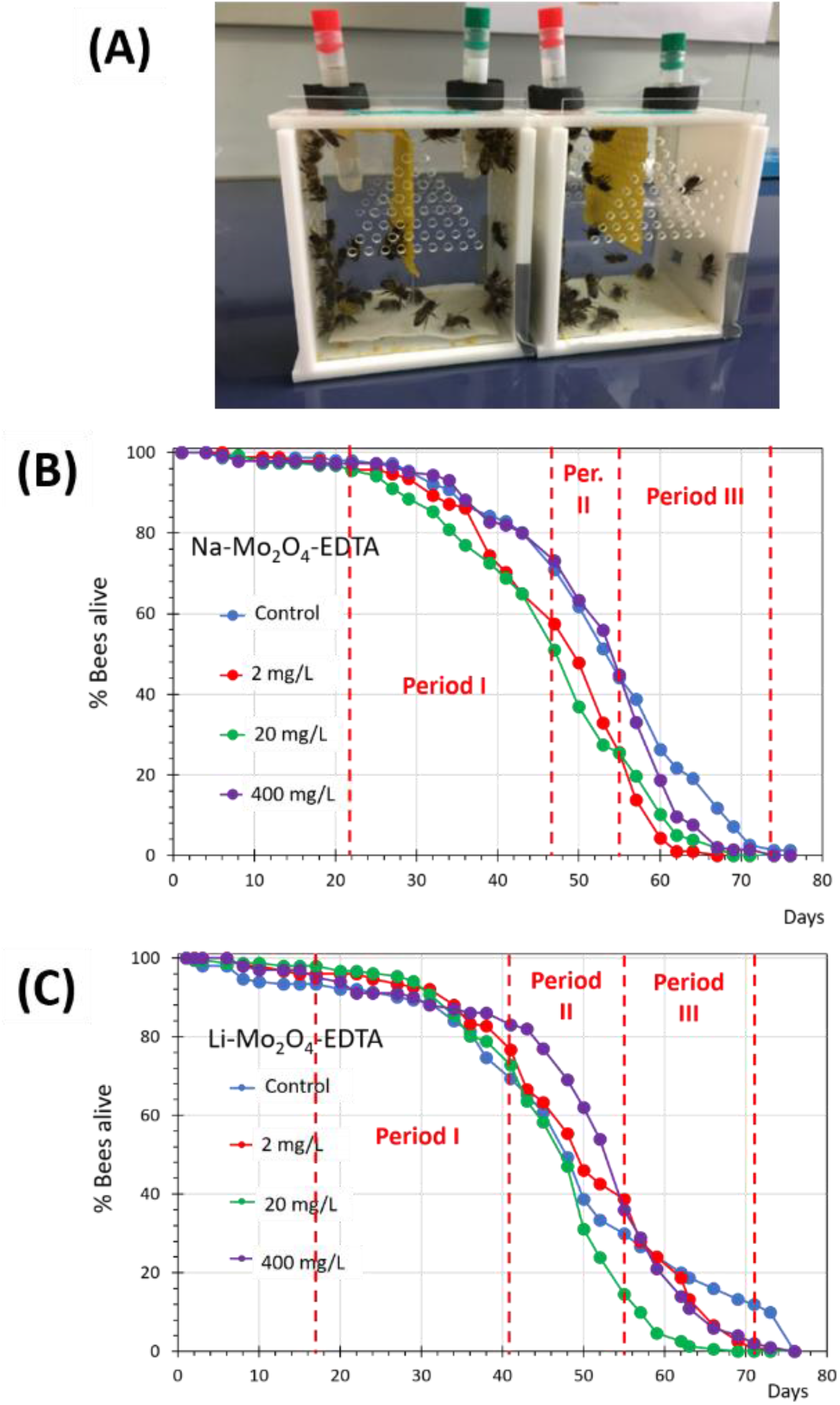
Population cages used in this experiment (A); Mortality curves obtained with complex **Na-Mo**_**2**_**O**_**4**_**-EDTA** (B) and **Li-Mo**_**2**_**O**_**4**_**-EDTA** (C) highlighting the 3 periods defined to follow Mo and Na contents in bees; Controls are indicated in blue, 2 mg/L solution in red, 20 mg/L in green, 400 mg/L in purple.

We found that generally, the average curves obtained for control batches and for bees fed with 2, 20 and 400 mg/L solutions of both complexes are similar. The statistical analysis confirms that no difference in mortality appeared between the control group and the groups fed with both **Na-Mo**_**2**_**O**_**4**_**-EDTA** and **Li-Mo**_**2**_**O**_**4**_**-EDTA** complexes during the first 10 days (Table SIII.10, Part III, SI) and even after for all concentrations of **Li-Mo**_**2**_**O**_**4**_**-EDTA** (Table SIII.11, Part III, SI). In contrast, the cox model indicates slight negative impact of **Na-Mo**_**2**_**O**_**4**_**-EDTA** at 2 and 20 mg/L but positive impact at 400 mg/L. Nevertheless, no notable positive effect of feeding with Mo-complexes was observed on bees’ longevity in laboratory conditions, nor any notable excess of mortality related to chronic feeding with the two complexes, even at a very high concentration.

### 2.5 Assimilation of Mo by the honey bees in laboratory

Dead bees collected in the previous experiments were used to evaluate the assimilation of Mo complexes within the body of the bees, as a function of **Na-Mo**_**2**_**O**_**4**_**-EDTA** or **Li-Mo**_**2**_**O**_**4**_**-EDTA** concentration. The dead bees were divided into 3 batches (Periods 1, 2, 3, see Figures 4b and 4c) depending on feeding duration, and Mo-content was then determined in heads, thoraxes and abdomens by ICP-MS (see Tables SVI.2 and SVI.3, Part VI, SI). First, we found low Mo levels in the control bees, with values respectively around 0.2 ppm for **Na-Mo**_**2**_**O**_**4**_**-EDTA** experiments and 0.5-0.7 ppm for **Li-Mo**_**2**_**O**_**4**_**-EDTA** experiments in the head and in the abdomen, and lower levels in the thorax (0.12 and 0.25 ppm respectively). An increase in Mo level was found in head, thorax and abdomen when the concentration of **Na-Mo**_**2**_**O**_**4**_**-EDTA** or **Li-Mo**_**2**_**O**_**4**_**-EDTA** increased in the feeding syrup. For each concentration Mo level reached a different plateau which did not depend on the duration of feeding. More precisely, Mo level in the head increased from ∼0.19 ppm in the control group, to 0.58, 1.83 and 27 ppm, respectively for the groups fed with syrups at 2, 20 and 400 mg/L of **Na-Mo**_**2**_**O**_**4**_**-EDTA**, *i*.*e*. up to a 140-fold increase. At the same time, Mo-level variations were higher in the thorax, from 0.12 ppm to 0.73, 2.09 and 58 ppm with the same syrups. Similar observations were made with **Li-Mo**_**2**_**O**_**4**_**-EDTA**, with levels reaching 34.8 ppm in the head and 75.2 ppm in the thorax. With both complexes, as bees defecate very little while in cages, strong accumulation was observed in the bees’ abdomen, in particular in the faeces (up to 3600 ppm). In the case of **Li-Mo**_**2**_**O**_**4**_**-EDTA**, an accumulation of Lithium was also observed, in particular in the head in which we estimate that 5.6 Li^+^ are assimilated for each Mo atom (see Figure SVI.5, Part VI, SI).

### 2.6 X-Ray fluorescence studies

X-ray fluorescence spectra were recorded on the Nanoscopium beamline of Synchrotron Soleil (Gif-sur-Yvette, France) on 20-50 μm thick slices of head, thorax and abdomen and on hypopharyngeal glands extracted from the heads of worker bees to locate Mo more precisely within the bees. The study was conducted on bees fed during 14 days after their emergence with a sugar syrup containing **Na-Mo**_**2**_**O**_**4**_**-EDTA** at 400 mg/L or unspiked syrup for the control group. A first series of spectra on abdomen slices revealed that this technique can be employed and that the faeces present in the rectum of bees fed with **Na-Mo**_**2**_**O**_**4**_**-EDTA** are highly loaded with Mo as expected from part 2.5, while Mo is negligible in the rectum of control bees (see Figures SVII.6 and SVII7, Part VII, SI). This part of the bees was not analyzed further. Even if it is really difficult to confidently and precisely quantify the levels of Mo for different reasons (amount of Mo in the nanogram range for each slice, limited access to the facility, different recording conditions between samples to find the good ones, limited number of samples…) such analysis applied to slices and glands of bees not supplemented with **Na-Mo**_**2**_**O**_**4**_**-EDTA** qualitatively revealed that Mo-content i) is negligible in the brain (see Figure 5), ii) is low in the neurolemma, hypopharyngeal glands and muscles (see Figures 6A and 7A, and Figures SVII.9, SVII.13 and SVII.16, Part VII, SI) as evidenced by a shoulder on the X-ray fluorescence spectra at the energy expected for the element Mo (16.9 to 17.9 eV) and iii) is moderate but at least two times higher in the cuticle parts, which appears as the richest part of the control bees, as shown for instance in the figure SVII.18 (SI).

**Figure 5.**
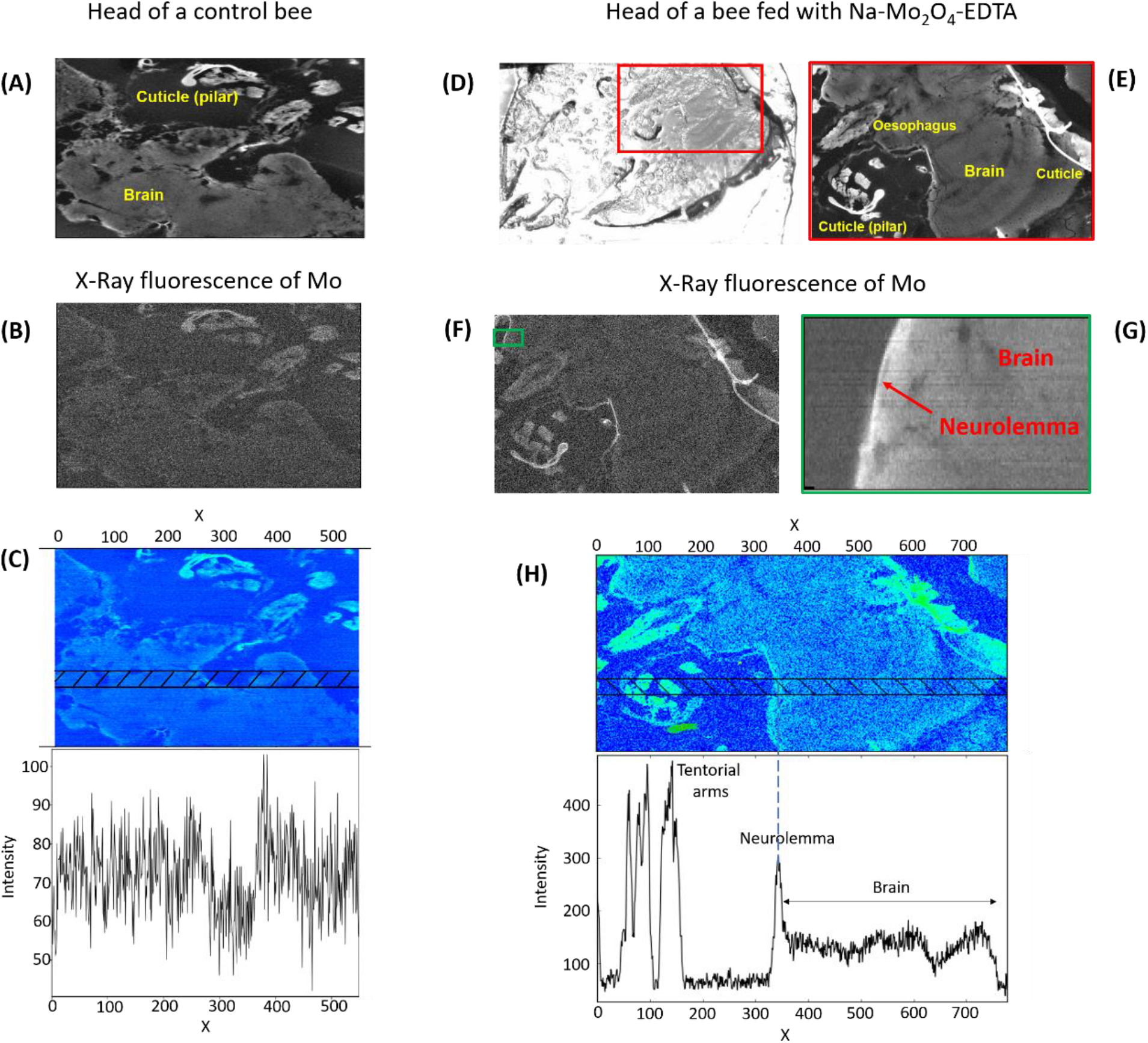
Comparisons of heads of a worker bee control (left part) and a bee fed with Na-Mo_2_O_4_-EDTA (right part). Control bee: Picture of 20 μm thick slice of the head obtained by X-ray fluorescence of all elements (A) or just by X-ray fluorescence of Mo of a portion of brain of a control bee (B); (C) Variation of intensity of the X-Ray fluorescence of Mo in the hatched area (in the energy range 16.9 to 17.9 eV) along X in a 20 μm thickness slice of the head of a worker bee control. Bee fed with Na-Mo_2_O_4_-EDTA: Picture of a 20 μm thick slice of the head of a worker bee obtained by microscope (D), by X-ray fluorescence of all elements in the red rectangle (E, size 2000 μm x 1000 μm, pixel size 2×2 μm^2^, acquisition time/pixel 100 ms), and by X-ray fluorescence of Mo in the whole zone (F) or focused on the neurolemma (G, green rectangle of size 181 × 64 μm^2^ ; pixel size 1×1 μm^2^, acquisition time/pixel 100 ms); (H) Variation of intensity of the X-Ray fluorescence of Mo in the hatched area (in the energy range 16.9 to 17.9 eV) along X in a 20 μm thickness slice of the head of a worker bee, evidencing the presence of Mo in tentorial arms (cuticle), neurolemma and brain.

**Figure 6.**
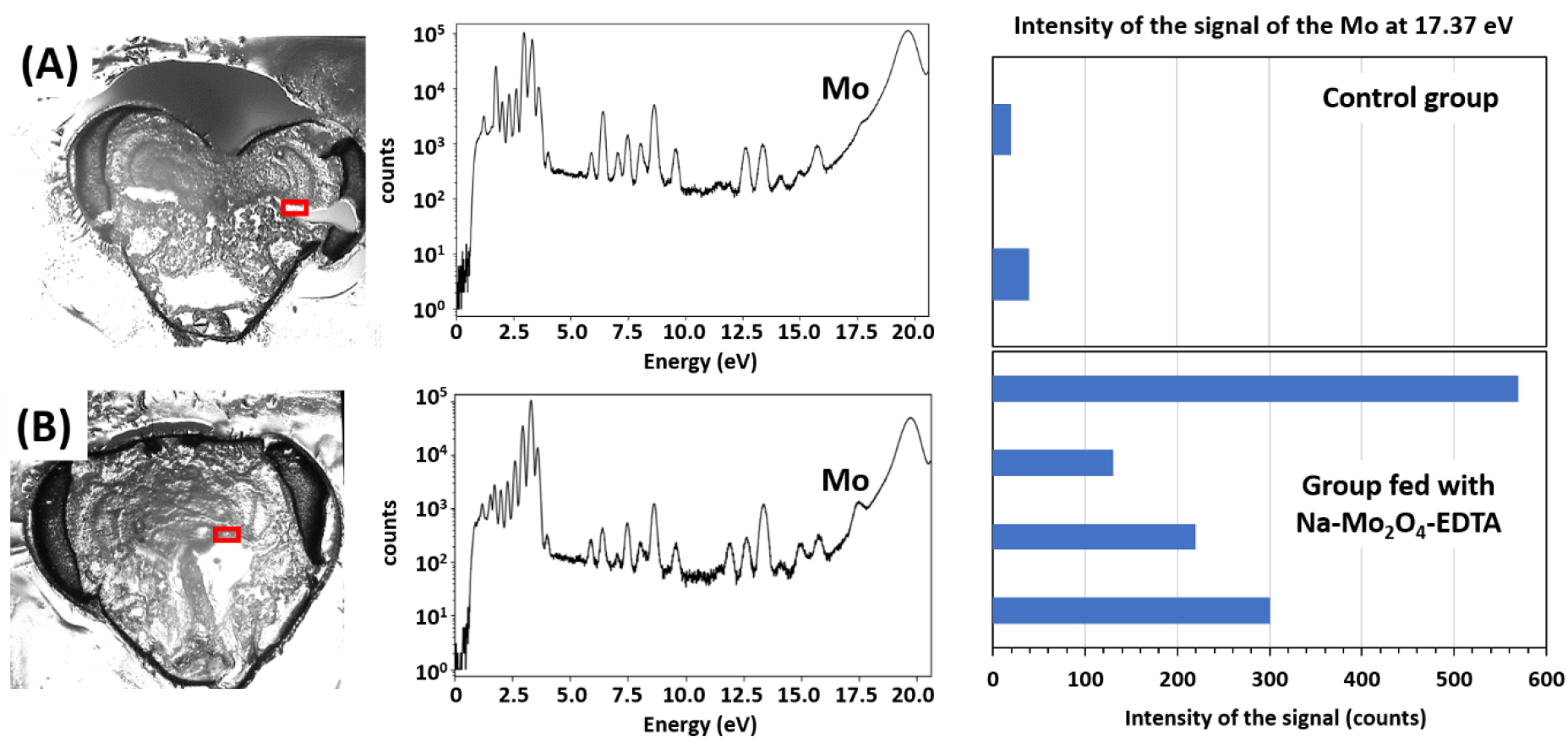
Pictures of 40 μm thick slices of the head of a worker bee from the control group (A) and bees fed with Mo (B) with the corresponding fluorescence spectra of all elements in the red rectangle zones corresponding to the neurolemma (A: Pixels 0.5×0.5 μm^2^, size 100×60 μm^2^, acquisition time/pixel 300 ms; B: Pixels 0.5×0.5 μm^2^, size 136×33 μm^2^, acquisition time/pixel 300 ms). The energy of the spectrum corresponding to Mo is indicated.

Figures 5D-5H show the distribution of Mo on a 20 μm thick slice of the head of a bee fed with the complex **Na-Mo**_**2**_**O**_**4**_**-EDTA**. This slice contains elements of the brain, neurolemma, cuticle and esophagus. After feeding with **Na-Mo**_**2**_**O**_**4**_**-EDTA** at 400 mg/L, the intensity of the fluorescence of Mo in the cuticle (external cuticle and tentorial arms) and in the muscles were similar as in the control group (see also analysis of thorax, Figures SVII.18-SVII.20, Part VII, SI) and it was not possible to clearly evidence any Mo increase by feeding. More interestingly, in bees fed with **Na-Mo**_**2**_**O**_**4**_**-EDTA** containing syrup, the fluorescence of Mo is clearly seen in the brain and found especially in the neurolemma around the brain (Figure 5F and 5G) in contrast with control bees (Figure 5B). The Mo level appears slightly enhanced within the brain after feeding from an intensity of ca 70-80 counts in control bees (Figure 5C), comparable to the baseline, to ca 130-140 counts in fed bees (Figure 5H).

Concomitantly, the increase of Mo in the neurolemma is much more pronounced. From several X-ray spectra recorded in the same experimental conditions for several individuals from both groups (Figure 6 and Figures SVII13 and SVII14, Part VII, SI), Mo-content in bees fed with **Na-Mo**_**2**_**O**_**4**_**-EDTA** appeared indeed around 10-fold higher than in the control group, showing an accumulation of Mo in the neurolemma.

#### Hypopharyngeal glands

The hypopharyngeal glands found in the head of honey bee workers are crucial for bees’ nutrition and health [23]. Glands were taken from control bees or bees fed with the complex **Na-Mo**_**2**_**O**_**4**_**-EDTA** at 400 mg/L for 14 days and analyzed by X-Ray fluorescence (Figure 7 and Figures SVII-16-SVII-17, Part VII, SI).

**Figure 7.**
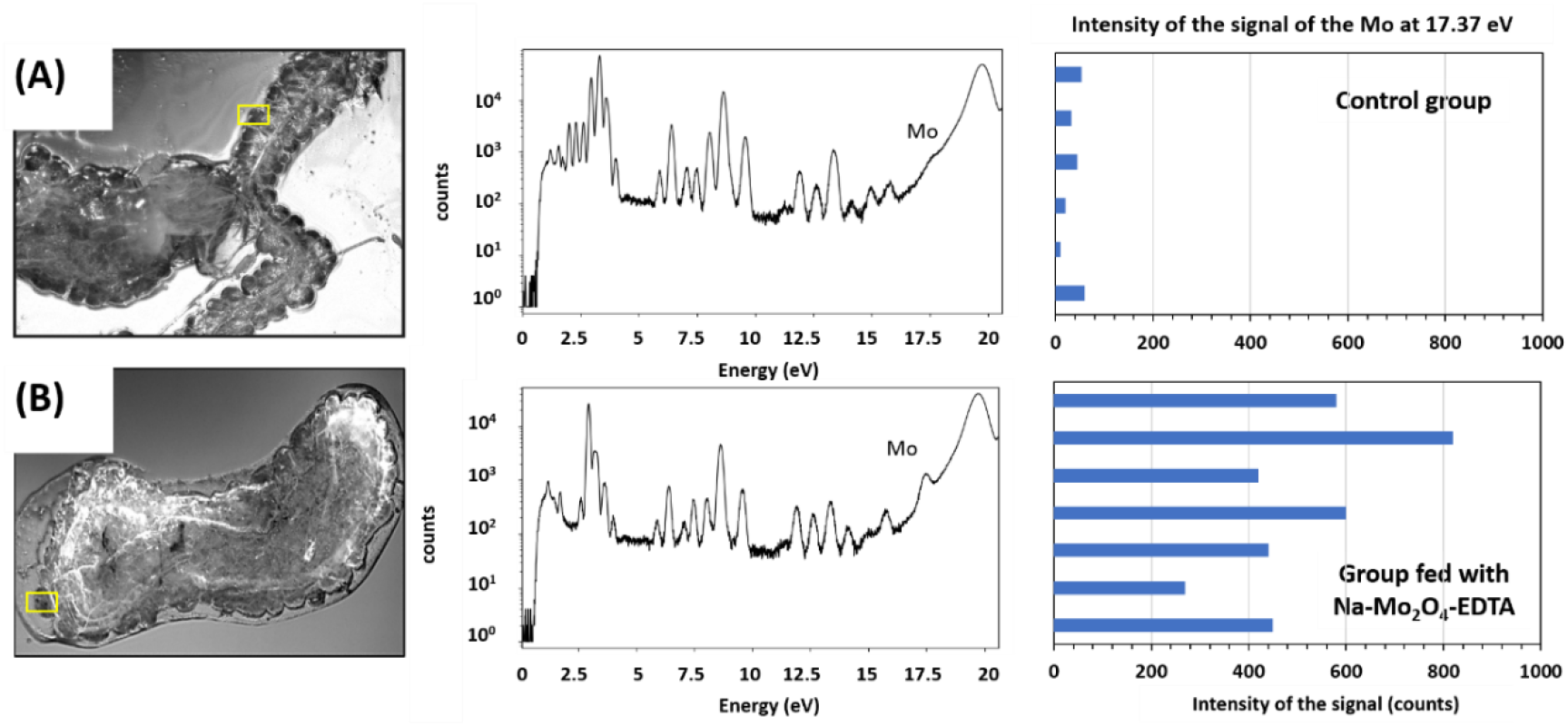
Hypopharyngeal glands extracted from workers of the control group (A) and bees fed with Mo (B) with their corresponding total X-ray fluorescence spectra. The yellow rectangles indicate the part of the gland on which the fluorescence spectrum was measured (A: Pixels 0.5×0.5 μm^2^, size 108×49 μm^2^, acquisition time/pixel 200 ms; B: Pixels 0.5×0.5 μm^2^, size 70×80 μm^2^, acquisition time/pixel 200 ms).

In control bees (Figure 7A), the fluorescence spectrum reveals a low, natural, presence of Mo in these glands. Feeding with a sugar syrup containing the **Na-Mo**_**2**_**O**_**4**_**-EDTA** complex induced a remarkable fluorescence peak, as shown in Figure 7B. The results obtained for 7 *acini* of hypopharyngeal glands from 2 different individuals are similar (see Part VII, SI). A 14x increase in Mo content was estimated in bees’ hypopharyngeal glands after feeding bees with **Na-Mo**_**2**_**O**_**4**_**-EDTA** at 400 mg/L. These results evidence that the hypopharyngeal glands are a primary target for the Mo complex, which increases the Mo level naturally present in these glands and due to the importance of these glands could impact the health of the bees.

### 2.7 Metabolism and antioxidant properties of Mo-complexes within the bee organism

**Na-Mo**_**2**_**O**_**4**_**-EDTA** is assimilated by bees. X-ray photoelectron spectroscopy (XPS) experiments were performed on the faeces of individuals fed with **Na-Mo**_**2**_**O**_**4**_**-EDTA** at 400 mg/L to address the question of its metabolism within the bees. The Mo 3d spectral region of **Na-Mo**_**2**_**O**_**4**_**-EDTA** and of bee faeces, as well as their associated reconstruction, are presented in Figure 8.

**Figure 8.**
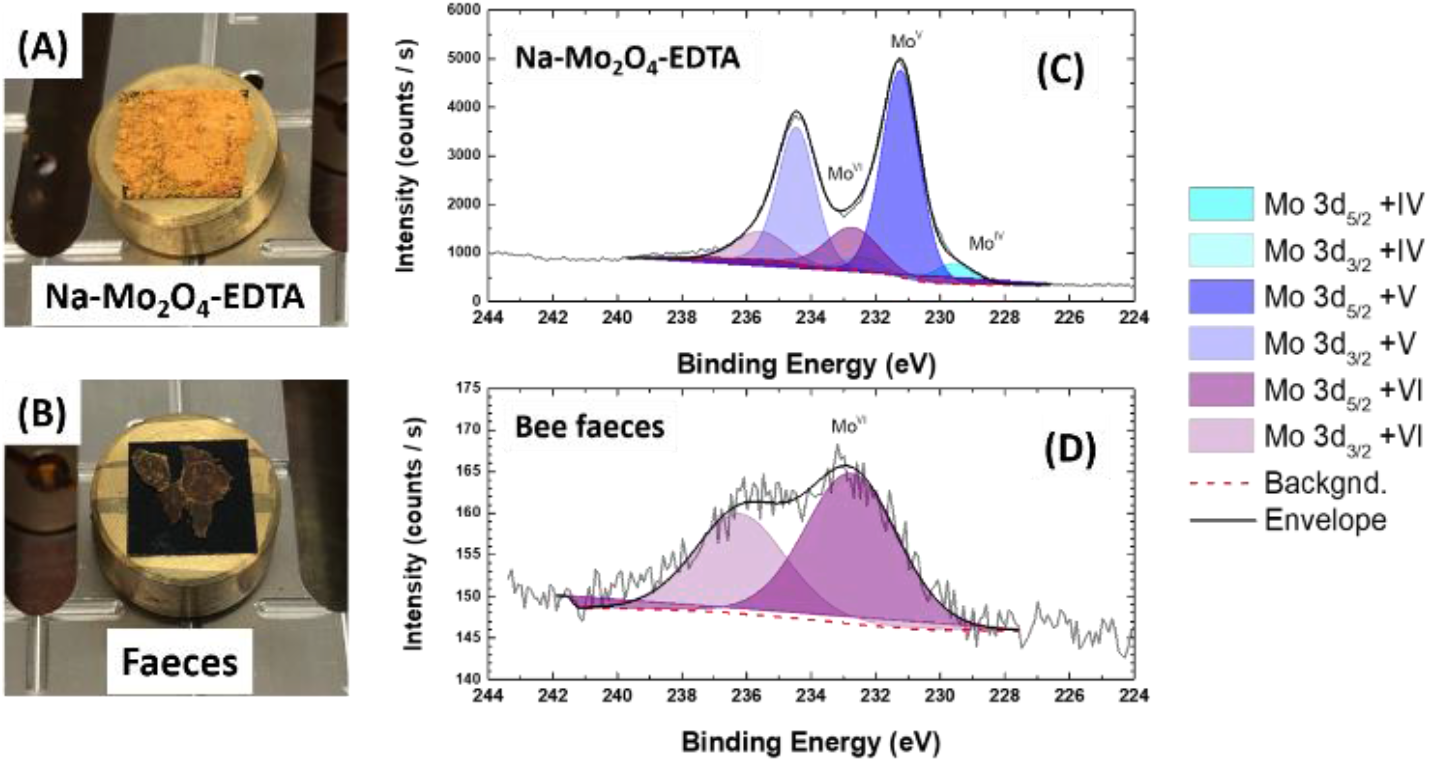
Samples of **Na-Mo**_**2**_**O**_**4**_**-EDTA** and bee faeces deposited on carbon tape (A and B, respectively). The diameter of the sample holder is 1 cm; XPS high-resolution spectra (Mo3d region) of **Na-Mo**_**2**_**O**_**4**_**-EDTA** (C) and bee faeces (D). The different contributions and the background used for the reconstruction are plotted as well as the final envelope.

The **Na-Mo**_**2**_**O**_**4**_**-EDTA** spectrum can be resolved in three doublets, with the most intense corresponding, as expected, to the Mo^+V^ oxidation state (Mo 3d_5/2_ = 231.2 eV, full width at half maximum FWHM = 1.4 eV) [24]. For the bee faeces, the Mo 3d core level spectrum displays only one broad doublet (Mo 3d_5/2_ at BE 232.7 eV, FWHM = 3.5 eV). The energy position is fully consistent with a Mo^+VI^ phase[24]. The presence of Mo in +V or +IV oxidation states was not identified. This experiment thus unambiguously evidences that the complex **Na-Mo**_**2**_**O**_**4**_**-EDTA** has been oxidized inside the bees, probably into molybdate anion MoO_4_^2-^. It constitutes the first element suggesting that the **Na-Mo**_**2**_**O**_**4**_**-EDTA** complex may act as an antioxidant by oxidation of the Mo(V) atoms into the Mo(VI) oxidation state.

Such antioxidant activity (AOA) in both bees and larvae can hold particular significance for insects with high metabolic rates, inherently generating substantial volumes of free radicals [25]. Therefore, implementing proactive measures to mitigate oxidative stress effects can boost the resilience of honey bees substantially [26].

The AOA of the complexes **Na-Mo**_**2**_**O**_**4**_**-EDTA** and **Li-Mo**_**2**_**O**_**4**_**-EDTA** were evaluated on *Apis mellifera carnica*. Experimental beehives were fed at the beginning of spring for 14 days with 50% sugar syrup enriched or not with **Na-Mo**_**2**_**O**_**4**_**-EDTA** and **Li-Mo**_**2**_**O**_**4**_**-EDTA** at 0.2 mg/L. Sodium molybdate Na_2_MoO_4_.2H_2_O was also used, for comparison, with a Mo content equivalent to that of the **Na-Mo**_**2**_**O**_**4**_**-EDTA** and **Li-Mo**_**2**_**O**_**4**_**-EDTA** compounds. The AOA was evaluated by ABTS method [27] on hemolymph from worker bees and larvae, wax, honey, propolis, royal jelly and bee bread. The results are expressed as IC_50_ values, the half-maximal Inhibitory Concentration, defined as the concentration that causes the loss of 50% of the activity of ABTS radicals. The lower the IC_50_ values, the higher the antioxidative activity. Some selected results are depicted in Figure 9 while the other results are given in the supporting information (Figures SVIII-6 to SVIII-12, Part VIII, SI). As shown in Figure 9, the IC_50_ values obtained for the hemolymph of worker bees are low, indicating a naturally good antioxidant activity (Figure 9A). The AOA of hemolymph in worker bees increased for both **Na-Mo**_**2**_**O**_**4**_**-EDTA** and **Li-Mo**_**2**_**O**_**4**_**-EDTA** complexes compared to the control group, suggesting that these complexes act on the organisms of bees as antioxidants or induce AOA. The AOA of both complexes appears similar, slightly better with **Na-Mo**_**2**_**O**_**4**_**-EDTA**. Interestingly, even though sodium molybdate Na_2_MoO_4_ cannot chemically act as a direct antioxidant, the AOA of the hemolymph of bees treated with this compound increased, suggesting a more complex mechanism. In the case of larval hemolymph (Figure 9B), the findings are similar for the control group and sodium molybdate, showing no increase in AOA. Both the complexes **Na-Mo**_**2**_**O**_**4**_**-EDTA** and **Li-Mo**_**2**_**O**_**4**_**-EDTA** showed an important increase in AOA in larvae with ABTS method.

**Figure 9.**
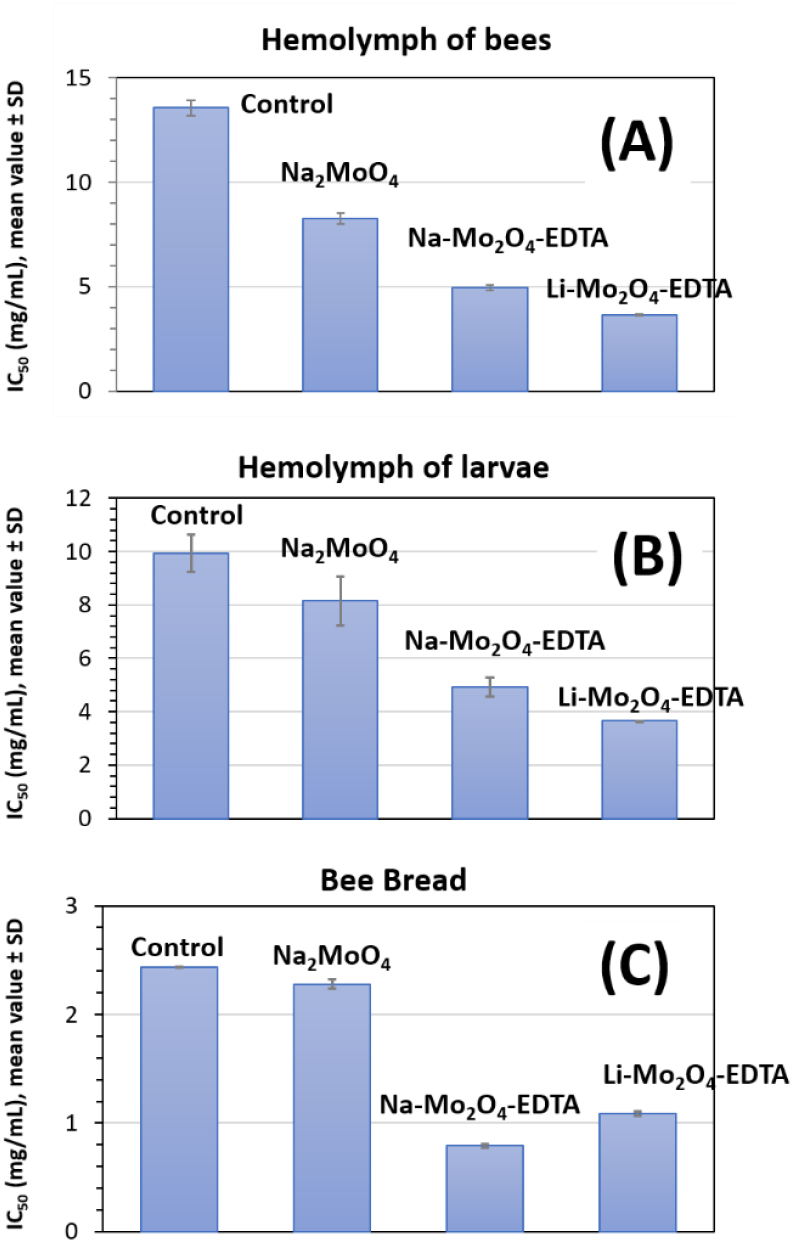
Antioxidant properties of hemolymph of worker bees (A) and larvae (B) and beebread (C) obtained from beehives of the control group or the groups fed with Na_2_MoO_4_.2H_2_O, **Na-Mo**_**2**_**O**_**4**_**-EDTA** and **Li-Mo**_**2**_**O**_**4**_**-EDTA**. Each IC_50_ value corresponds to an average value between three replicates.

Regarding hive products (see Part VIII, SI), the AOA of honey samples was similar for control, sodium molybdate and **Na-Mo**_**2**_**O**_**4**_**-EDTA** groups but strongly enhanced with **Li-Mo**_**2**_**O**_**4**_**-EDTA**. This result is surprising and could be either due to traces of the feeding syrup in the honey for this group or to an action of this complex on the enzymes produced in the bees’ crop to produce honey. The AOA of royal jelly and bee wax taken from the 4 groups were similar in all cases. Finally, both complexes significantly amplified the antioxidant activity of bee bread (Figure 9C), showcasing remarkable efficacy in enhancing its overall antioxidant potential [28] for the benefits of all the colony.

## 3. Conclusion

Based on a large set of field test campaigns and lab experiments, this study establishes the importance of the molybdenum trace element for honey bee health. The field tests were performed on more than 700 beehives, on several honey bee sub-species, and in three different countries in Europe and the United States. To our knowledge, the scale of these field studies is unprecedented in the literature.

The results showed that feeding with only a few milligrams of complexes **Li-Mo**_**2**_**O**_**4**_**-EDTA** or **Na-Mo**_**2**_**O**_**4**_**-EDTA** during spring or autumn improves key parameters of colony performance: it enhances in particular colony development, and honey and wax productions. Furthermore, it also reduces infestation by the mite *Varroa destructor* and decreases winter colony losses. Importantly, feeding with Mo-complexes had no effect on the quality of the produced honey.

In-depth studies in the field, in the laboratory and at the synchrotron facility have demonstrated that these complexes have a prolonged action in the beehives, as they are consumed over a long period of time and are assimilated by the bees (larvae and workers). We observed that Mo is naturally present in bees’ cuticle, muscles, hypopharyngeal glands and in the neurolemma. Interestingly, Mo levels increased in the neurolemma and in the hypopharyngeal glands after supplementation with Mo-complexes. Lastly, our experiments suggest that the complexes **Li-Mo**_**2**_**O**_**4**_**-EDTA** and **Na-Mo**_**2**_**O**_**4**_**-EDTA** act as antioxidant agents in bees.

These results are promising for the beekeeping industry, which is currently suffering from numerous challenges weakening honey bee colonies. The Mo-complexes were demonstrated to have positive effects for beekeeping, improving winter survival, hygienic behavior and colony productivity. The presence of Mo in multiple organs and the antioxidant activity of the complexes could explain the important effects found on colony health. Further studies are currently in progress to assess the impact of Mo-complexes on the resilience of bees towards pesticides and the expression of Mo-enzymes. More generally, this study paves the way for further research into the possible positive impacts that feeding with other micronutrient-based complexes may provide on honey bee health.

## Materials and Methods

### Materials, chemicals and methods

All chemicals were purchased from Sigma Aldrich or Acros Chemical Company Ltd. and used without further purification. The syntheses of complexes are given in Part II of the Supporting Information. All the methods used in this study, with the raw data are detailed in Supporting Information.

### Supporting Information

To facilitate the reading, the supporting information is divided into 8 independent parts built as reports to cover each topic. Each part contains, materials and methods, results and discussion and raw data. Part I contains data about Mo level measured in hundreds honeybee samples by ICP-MS or ICP-OES to establish a reference level; Part II contains all details on the synthesis of complexes, their characterization and the stability studies by DFT calculation, _1_H NMR and UV-visible spectroscopies; Part III contains the toxicity studies of Mo-complexes on mice, *Daphnia Magna*, and bees both in laboratory and in hives; Part IV gathers all the field tests in beehives and analyses on honey with the raw data obtained for each test campaign; Part V is focused on the cycle of Mo in beehives after feeding by monitoring Mo content by ICP-MS in larvae, worker bees, honey/syrup, wax, and bee bread; Part VI contains the data about the assimilation of Mo by honey bees in laboratory conditions in head, thorax and abdomen. The Mo, Na/Li contents are measured by ICP-MS; Part VII gathered all data of X-Ray fluorescence acquired at synchrotron SOLEIL; Part VIII focuses on the metabolism and the antioxidant properties of Mo-complexes consumed by honey bees thank to a XPS study of bees faeces and spectroscopic determination of anti-oxidant activity of hemolymph of bees and larvae as well as hive products by ABTS and DPPH methods.

## Supporting information

Table of contents of Supporting Information

Molybdenum contents in honeybees

Synthesis and stability studies of Molybdenum complexes for this study

Toxicity studies (Mice, Daphnia, Bees)

Tests in beehives

Tracking Mo in hives

Assimilation of Mo-complexes by honeybees

X-ray fluorescence studies at synchrotron SOLEIL

XPS stuides of bee faeces and Anti-Oxydant properties

## Acknowledgments

University of Versailles, the “Institut Universitaire de France, IUF” and the CNRS are gratefully acknowledged for financial support. AF gratefully acknowledge Campus France for Excellence Eiffel grant as well as State University of Moldova for funding his PhD thesis. This work was funded by the Charmmmat labex (project “COMPA”), the IDEX Paris-Saclay (project “COMPA2”), the “lune de miel” foundation, the UVSQ foundation (project “COMPA3, and the SATT Paris-Saclay (projects “COMPA4” and “APIMONA”). This project has received financial support from CNRS through the MITI interdisciplinary programs. The financial support from National Agency for Research and Development (ANCD) of the Republic of Moldova (Project No 20.80009.5007.10) is also acknowledged. Dr Clémence Riva is gratefully acknowledged for her help for the statistical analyses.

