## Supplementary material for "Food supplementation with molybdenum complexes improves honey bee health": Table of contents of Supporting Information

k) Hellenic Agriculture Org. "DIMITRA", Institute of Animal Science, Department of Apiculture, 63200 Nea Moudania, Greece,

### *Supporting Information*

### Table of contents

### Table of contents


#### Part I. Molybdenum in honey bees (10 pages)

#### Part II. Molybdenum-based complexes (14 pages)

II.1 Synthesis and characterization

II.2. Stability studies

i. DFT Studies

ii. $^{1}$H NMR and electronic spectroscopy studies

#### Part III. Toxicity studies (36 pages)

III.1- Toxicity on mice

III.2- Toxicity on *Daphnia Magna*

III.3- Acute toxicity on bees

III.4- Chronic toxicity on bees: mortality studies

III.5. Tolerance studies in beehives

#### Part IV. Tests in Beehives and analyses of honey (66 pages)

IV.1- Tests in the apiary of Institute of Zoology, Moldova: 2013-2019

IV.2- Tests in France in operating conditions: 2019.

IV.3- Tests in California, USA: 2019-2020

IV.4- Tests in Greece: 2020-2021

IV.5- General conclusions

V.6- Raw data

#### Part V. Tracking Mo in hives after the feeding (17 pages)

V.1 Objectives of the study

V.2 Materials and methods

V.3 Results

V.3.1 Molybdenum-content in honey

V.3.2 Molybdenum-content in bee larvae

V.3.3 Molybdenum-content in worker bees

V.3.4 Molybdenum-content in wax

V.4 Conclusions

#### Part VI. Assimilation of Mo by bees : ICP-MS/ICP-OES studies (8 pages)

VI.1 Experimental procedures

VI.2 Results and discussion

VI.3 Conclusion

**Part VII. Synchrotron radiation X-ray fluorescence microscopy (19 pages)**

VII.1 Experimental procedures

1°) Animal preparation

2°) Cross sections

3°) X-Ray Fluorescence spectra.

VII.2 Bee Cross-section results

VII.3 Results of synchrotron measurements

VII.3.1 X-Ray Fluorescence spectrum of resin on $Si_3N_4$ membrane

VII.3.2 X-Ray fluorescence spectra of a honey bee abdomen

VII.3.3 X-Ray fluorescence spectra of a honey bee head

1°) Localization of the Mo in a large area of the head

2°) Focus on the neurolemma

3°) Focus on hypopharyngeal glands

VII.3.4 X-Ray fluorescence spectra of cuticle / thorax

VII.4 Conclusions of X-Ray fluorescence studies

**Part VIII. Antioxidant properties of Na-$Mo_2O_4$-EDTA and Li- $Mo_2O_4$-EDTA complexes (23 pages)**

VIII-1 : XPS studies on bees' faeces

VIII-1-1 Experimental section

VIII-1-2 XPS analysis

VIII-1-3 Conclusion

VIII-2 : Antioxidant properties measured in honey bees and products of the hive.

VIII.2.1 Experimental protocols.

1°) Apiary and conditions of feeding.

2°) Sampling

3°) Anti-oxidant activity (AOA) determinations

VIII.2.2 Results.

VIII.2.3 Discussion / conclusions
