## Supplementary material for "Food supplementation with molybdenum complexes improves honey bee health": Molybdenum contents in honeybees

### *Supporting Information*

#### Part I

#### Molybdenum in honey bees

### Molybdenum in honey bees

Samples of honey bee workers were collected in different areas and environments to assess natural levels of molybdenum in France (see map, Figure SI-1), in Nea Moudania, Thessaloniki and Polygyros region of Greece (respectively urban/, industrial and natural environments) and in Moldova (forest environments in the region of Chisinau).

The bees were frozen after collection, then lyophilized, ground, and mineralized before analysis of their molybdenum content by ICP-MS. Analyses were performed by LEAV (Laboratoire de l’Environnement et d’Analyses de Vendée), La Roche-sur-Yon, France.

The results found for Mo contents in ppm in dehydrated bees are gathered in Table SI-1. The uncertainty is estimated to ± 0.02 ppm (µg/g).

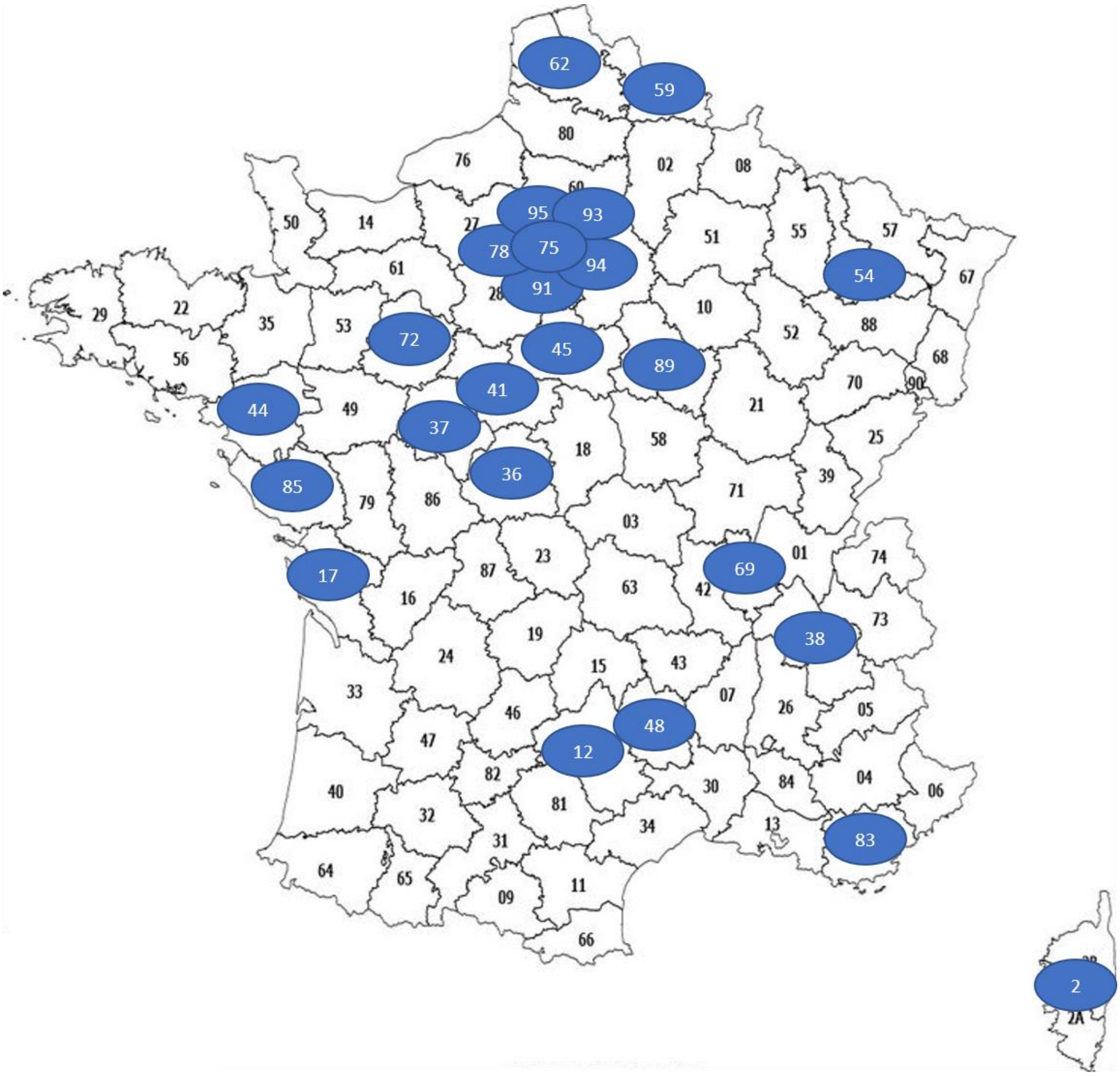


*Figure SI-1. Map of the areas in France, where the bees were collected. Numbers correspond to the administrative numbers given to each department in France. The samples collected cover different type of environments and climates.*

*Table SI-1. Mo content in ppm (µg/g of dried bee) as a function of sampling areas, month and year of sampling and environment. NAT, AGRI, URB, INDUS are given for Natural, Agricultural, Urban or Industrial environments, respectively.*

| N° | Sampling Area | Month | Year | Environment[a] | Mo content (ppm) |
|---|---|---|---|---|---|
| 1 | France (12) | 4 | 2021 | AGRI | 0.31 |
| 2 | France (12) | 7 | 2021 | AGRI | 0.34 |
| 3 | France (12) | 9 | 2021 | AGRI | 0.36 |
| 4 | France (17) | 4 | 2013 | AGRI | 0.39 |
| 5 | France (17) | 4 | 2014 | AGRI | 1.19 |
| 6 | France (17) | 7 | 2014 | AGRI | 0.25 |
| 7 | France (17) | 9 | 2014 | AGRI | 0.22 |
| 8 | France (17) | 5 | 2015 | AGRI | 0.25 |
| 9 | France (17) | 7 | 2015 | AGRI | 0.34 |
| 10 | France (17) | 9 | 2015 | AGRI | 0.29 |
| 11 | France (17) | 5 | 2022 | AGRI | 0.37 |
| 12 | France (17) | 5 | 2022 | AGRI | 0.36 |
| 13 | France (17) | 5 | 2022 | AGRI | 0.49 |
| 14 | France (17) | 5 | 2022 | AGRI | 0.30 |
| 15 | France (17) | 5 | 2022 | AGRI | 0.41 |
| 16 | France (17) | 5 | 2022 | AGRI | 0.30 |
| 17 | France (17) | 5 | 2022 | AGRI | 0.35 |
| 18 | France (17) | 5 | 2022 | AGRI | 0.38 |
| 19 | France (17) | 5 | 2022 | AGRI | 0.27 |
| 20 | France (17) | 5 | 2022 | AGRI | 0.28 |
| 21 | France (17) | 5 | 2022 | AGRI | 0.42 |
| 22 | France (17) | 5 | 2022 | AGRI | 0.75 |
| 23 | France (17) | 5 | 2022 | AGRI | 0.26 |
| 24 | France (17) | 5 | 2022 | AGRI | 0.30 |
| 25 | France (17) | 5 | 2022 | AGRI | 0.42 |
| 26 | France (17) | 5 | 2022 | AGRI | 0.24 |
| 27 | France (17) | 6 | 2022 | AGRI | 0.33 |
| 28 | France (17) | 6 | 2022 | AGRI | 0.29 |
| 29 | France (17) | 6 | 2022 | AGRI | 0.39 |
| 30 | France (17) | 6 | 2022 | AGRI | 0.34 |
| 31 | France (17) | 6 | 2022 | AGRI | 0.23 |
| 32 | France (17) | 6 | 2022 | AGRI | 0.28 |
| 33 | France (17) | 6 | 2022 | AGRI | 0.34 |
| 34 | France (17) | 6 | 2022 | AGRI | 0.27 |
| 35 | France (17) | 7 | 2022 | AGRI | 0.22 |
| 36 | France (17) | 7 | 2022 | AGRI | 0.26 |
| 37 | France (17) | 7 | 2022 | AGRI | 0.36 |
| 38 | France (17) | 7 | 2022 | AGRI | 0.30 |
| 39 | France (17) | 7 | 2022 | AGRI | 0.23 |
| 40 | France (17) | 7 | 2022 | AGRI | 0.21 |

| | | | | | |
|---|---|---|---|---|---|
| 41 | France (17) | 7 | 2022 | AGRI | 0.16 |
| 42 | France (17) | 7 | 2022 | AGRI | 0.18 |
| 43 | France (2) | 4 | 2013 | AGRI | 1.35 |
| 44 | France (2) | 4 | 2013 | AGRI | 0.68 |
| 45 | France (2) | 6 | 2013 | AGRI | 0.43 |
| 46 | France (2) | 6 | 2013 | AGRI | 0.46 |
| 47 | France (36) | 4 | 2019 | AGRI | 0.09 |
| 48 | France (36) | 7 | 2019 | AGRI | 0.19 |
| 49 | France (36) | 9 | 2019 | AGRI | 0.21 |
| 50 | France (36) | 4 | 2020 | AGRI | 0.33 |
| 51 | France (36) | 7 | 2020 | AGRI | 0.35 |
| 52 | France (36) | 9 | 2020 | AGRI | 0.36 |
| 53 | France (36) | 4 | 2021 | AGRI | 0.4 |
| 54 | France (36) | 6 | 2021 | AGRI | 0.59 |
| 55 | France (36) | 9 | 2021 | AGRI | 0.48 |
| 56 | France (37) | 7 | 2014 | NAT | 0.31 |
| 57 | France (37) | 9 | 2014 | NAT | 0.47 |
| 58 | France (37) | 4 | 2015 | NAT | 0.21 |
| 59 | France (37) | 7 | 2015 | NAT | 0.16 |
| 60 | France (37) | 9 | 2015 | NAT | 0.20 |
| 61 | France (37) | 3 | 2016 | NAT | 0.17 |
| 62 | France (37) | 7 | 2016 | NAT | 0.23 |
| 63 | France (37) | 9 | 2016 | NAT | 0.29 |
| 64 | France (37) | 3 | 2017 | NAT | 0.41 |
| 65 | France (37) | 7 | 2017 | NAT | 0.21 |
| 66 | France (37) | 9 | 2017 | NAT | 0.21 |
| 67 | France (37) | 4 | 2018 | NAT | 0.33 |
| 68 | France (37) | 7 | 2018 | NAT | 0.298 |
| 69 | France (37) | 9 | 2018 | NAT | 0.234 |
| 70 | France (37) | 4 | 2019 | NAT | 0.08 |
| 71 | France (37) | 7 | 2019 | NAT | 0.21 |
| 72 | France (37) | 9 | 2019 | NAT | 0.59 |
| 73 | France (37) | 4 | 2020 | NAT | 0.025 |
| 74 | France (37) | 7 | 2020 | NAT | 0.15 |
| 75 | France (37) | 9 | 2020 | NAT | 0.36 |
| 76 | France (37) | 4 | 2021 | NAT | 0.2 |
| 77 | France (37) | 6 | 2021 | NAT | 0.35 |
| 78 | France (37) | 9 | 2021 | NAT | 0.38 |
| 79 | France (38) | 8 | 2016 | URB | 0.82 |
| 80 | France (38) | 8 | 2016 | AGRI | 0.44 |
| 81 | France (38) | 9 | 2016 | URB | 0.435 |
| 82 | France (38) | 9 | 2016 | AGRI | 0.34 |
| 83 | France (41) | 6 | 2013 | AGRI | 0.45 |
| 84 | France (41) | 4 | 2014 | AGRI | 1.31 |
| 85 | France (41) | 7 | 2014 | AGRI | 0.39 |
| 86 | France (41) | 9 | 2014 | AGRI | 0.37 |

| | | | | | |
|---|---|---|---|---|---|
| 87 | France (41) | 4 | 2015 | AGRI | 0.23 |
| 88 | France (41) | 7 | 2015 | AGRI | 0.19 |
| 89 | France (41) | 9 | 2015 | AGRI | 0.24 |
| 90 | France (41) | 3 | 2016 | AGRI | 0.28 |
| 91 | France (41) | 7 | 2016 | AGRI | 0.37 |
| 92 | France (41) | 9 | 2016 | AGRI | 0.33 |
| 93 | France (41) | 3 | 2017 | AGRI | 0.38 |
| 94 | France (41) | 7 | 2017 | AGRI | 0.28 |
| 95 | France (41) | 9 | 2017 | AGRI | 0.54 |
| 96 | France (41) | 4 | 2018 | AGRI | 0.29 |
| 97 | France (41) | 7 | 2018 | AGRI | 0.45 |
| 98 | France (41) | 9 | 2018 | AGRI | 0.40 |
| 99 | France (41) | 4 | 2019 | AGRI | 0.08 |
| 100 | France (41) | 7 | 2019 | AGRI | 0.23 |
| 101 | France (41) | 9 | 2019 | AGRI | 0.24 |
| 102 | France (41) | 4 | 2020 | AGRI | 0.025 |
| 103 | France (41) | 7 | 2020 | AGRI | 0.35 |
| 104 | France (41) | 9 | 2020 | AGRI | 0.34 |
| 105 | France (41) | 4 | 2021 | AGRI | 0.20 |
| 106 | France (41) | 6 | 2021 | AGRI | 0.31 |
| 107 | France (41) | 9 | 2021 | AGRI | 0.33 |
| 108 | France (44) | 6 | 2014 | AGRI | 0.35 |
| 109 | France (44) | 7 | 2014 | AGRI | 0.41 |
| 110 | France (44) | 10 | 2014 | AGRI | 0.22 |
| 111 | France (44) | 4 | 2015 | AGRI | 0.3 |
| 112 | France (44) | 7 | 2015 | AGRI | 0.34 |
| 113 | France (44) | 9 | 2015 | AGRI | 0.34 |
| 114 | France (44) | 4 | 2016 | AGRI | 0.35 |
| 115 | France (44) | 7 | 2016 | AGRI | 0.25 |
| 116 | France (44) | 9 | 2016 | AGRI | 0.395 |
| 117 | France (45) | 4 | 2013 | AGRI | 0.34 |
| 118 | France (45) | 6 | 2013 | AGRI | 0.28 |
| 119 | France (45) | 4 | 2014 | AGRI | 1.19 |
| 120 | France (45) | 7 | 2014 | AGRI | 0.48 |
| 121 | France (45) | 9 | 2014 | AGRI | 0.33 |
| 122 | France (45) | 5 | 2015 | AGRI | 0.18 |
| 123 | France (45) | 7 | 2015 | AGRI | 0.26 |
| 124 | France (45) | 9 | 2015 | AGRI | 0.41 |
| 125 | France (45) | 3 | 2016 | AGRI | 0.36 |
| 126 | France (45) | 7 | 2016 | AGRI | 0.48 |
| 127 | France (45) | 9 | 2016 | AGRI | 0.535 |
| 128 | France (45) | 3 | 2017 | AGRI | 0.44 |
| 129 | France (45) | 7 | 2017 | AGRI | 0.41 |
| 130 | France (45) | 9 | 2017 | AGRI | 0.47 |
| 131 | France (45) | 4 | 2018 | AGRI | 0.58 |
| 132 | France (45) | 7 | 2018 | AGRI | 0.62 |

| | | | | | |
|---|---|---|---|---|---|
| 133 | France (45) | 9 | 2018 | AGRI | 0.395 |
| 134 | France (45) | 4 | 2019 | AGRI | 0.19 |
| 135 | France (45) | 7 | 2019 | AGRI | 0.42 |
| 136 | France (45) | 9 | 2019 | AGRI | 0.44 |
| 137 | France (45) | 4 | 2020 | AGRI | 0.025 |
| 138 | France (45) | 7 | 2020 | AGRI | 0.28 |
| 139 | France (45) | 9 | 2020 | AGRI | 0.49 |
| 140 | France (45) | 4 | 2021 | AGRI | 0.35 |
| 141 | France (45) | 6 | 2021 | AGRI | 0.69 |
| 142 | France (45) | 9 | 2021 | AGRI | 0.36 |
| 143 | France (48) | 4 | 2021 | NAT | 0.45 |
| 144 | France (48) | 7 | 2021 | NAT | 0.38 |
| 145 | France (48) | 9 | 2021 | NAT | 0.35 |
| 146 | France (54) | 6 | 2019 | AGRI | 0.35 |
| 147 | France (54) | 6 | 2019 | AGRI | 0.29 |
| 148 | France (54) | 7 | 2019 | AGRI | 0.35 |
| 149 | France (54) | 7 | 2019 | AGRI | 0.21 |
| 150 | France (54) | 9 | 2019 | AGRI | 0.22 |
| 151 | France (54) | 9 | 2019 | AGRI | 0.24 |
| 152 | France (54) | 6 | 2020 | AGRI | 0.30 |
| 153 | France (54) | 6 | 2020 | AGRI | 0.28 |
| 154 | France (54) | 8 | 2020 | AGRI | 0.31 |
| 155 | France (54) | 8 | 2020 | AGRI | 0.24 |
| 156 | France (54) | 9 | 2020 | AGRI | 0.28 |
| 157 | France (54) | 9 | 2020 | AGRI | 0.29 |
| 158 | France (59) | 9 | 2020 | AGRI | 0.52 |
| 159 | France (59) | 9 | 2020 | AGRI | 0.57 |
| 160 | France (59) | 9 | 2020 | AGRI | 0.63 |
| 161 | France (59) | 6 | 2021 | AGRI | 0.43 |
| 162 | France (59) | 6 | 2021 | AGRI | 0.42 |
| 163 | France (59) | 6 | 2021 | AGRI | 0.41 |
| 164 | France (59) | 6 | 2021 | AGRI | 0.44 |
| 165 | France (59) | 8 | 2021 | AGRI | 0.39 |
| 166 | France (59) | 8 | 2021 | AGRI | 0.52 |
| 167 | France (59) | 8 | 2021 | AGRI | 0.47 |
| 168 | France (59) | 8 | 2021 | AGRI | 0.46 |
| 169 | France (62) | 5 | 2016 | AGRI | 0.64 |
| 170 | France (62) | 7 | 2016 | AGRI | 0.57 |
| 171 | France (62) | 9 | 2016 | AGRI | 0.37 |
| 172 | France (62) | 5 | 2017 | AGRI | 0.71 |
| 173 | France (62) | 7 | 2017 | AGRI | 1.48 |
| 174 | France (62) | 9 | 2017 | AGRI | 0.78 |
| 175 | France (69) | 9 | 2021 | URB | 0.42 |
| 176 | France (72) | 7 | 2017 | NAT | 0.41 |
| 177 | France (72) | 7 | 2017 | NAT | 0.24 |
| 178 | France (72) | 9 | 2017 | NAT | 0.43 |

| | | | | | |
|---|---|---|---|---|---|
| 179 | France (72) | 9 | 2017 | NAT | 0.27 |
| 180 | France (72) | 5 | 2018 | NAT | 0.43 |
| 181 | France (72) | 5 | 2018 | NAT | 0.395 |
| 182 | France (75) | 9 | 2015 | URB | 0.68 |
| 183 | France (75) | 4 | 2019 | URB | 0.27 |
| 184 | France (75) | 4 | 2019 | URB | 0.30 |
| 185 | France (75) | 7 | 2019 | NAT | 2.31 |
| 186 | France (75) | 7 | 2019 | NAT | 0.69 |
| 187 | France (75) | 9 | 2019 | AGRI | 0.49 |
| 188 | France (75) | 9 | 2019 | AGRI | 0.51 |
| 189 | France (75) | 4 | 2020 | URB | 0.36 |
| 190 | France (75) | 7 | 2020 | NAT | 0.64 |
| 191 | France (75) | 9 | 2020 | AGRI | 0.77 |
| 192 | France (78) | 4 | 2013 | URB | 0.35 |
| 193 | France (78) | 6 | 2013 | URB | 0.47 |
| 194 | France (78) | 4 | 2014 | URB | 1.12 |
| 195 | France (78) | 7 | 2014 | URB | 0.46 |
| 196 | France (78) | 9 | 2014 | URB | 0.39 |
| 197 | France (78) | 4 | 2015 | URB | 0.26 |
| 198 | France (78) | 7 | 2015 | URB | 0.49 |
| 199 | France (78) | 9 | 2015 | URB | 0.40 |
| 200 | France (83) | 4 | 2013 | NAT | 0.45 |
| 201 | France (83) | 4 | 2013 | NAT | 0.42 |
| 202 | France (83) | 4 | 2013 | NAT | 0.35 |
| 203 | France (83) | 4 | 2014 | NAT | 0.53 |
| 204 | France (83) | 7 | 2014 | NAT | 0.36 |
| 205 | France (83) | 9 | 2014 | NAT | 0.35 |
| 206 | France (85) | 3 | 2022 | NAT | 0.23 |
| 207 | France (85) | 4 | 2022 | NAT | 0.11 |
| 208 | France (85) | 5 | 2022 | NAT | 0.10 |
| 209 | France (85) | 5 | 2022 | NAT | 0.09 |
| 210 | France (85) | 6 | 2022 | NAT | 0.07 |
| 211 | France (85) | 6 | 2022 | NAT | 0.23 |
| 212 | France (85) | 6 | 2022 | NAT | 0.14 |
| 213 | France (85) | 6 | 2022 | AGRI | 0.23 |
| 214 | France (85) | 6 | 2022 | AGRI | 0.14 |
| 215 | France (85) | 7 | 2022 | NAT | 0.31 |
| 216 | France (85) | 7 | 2022 | NAT | 0.21 |
| 217 | France (85) | 7 | 2022 | AGRI | 0.31 |
| 218 | France (85) | 7 | 2022 | AGRI | 0.25 |
| 219 | France (85) | 8 | 2022 | NAT | 0.25 |
| 220 | France (85) | 8 | 2022 | AGRI | 0.25 |
| 221 | France (85) | 9 | 2022 | NAT | 0.27 |
| 222 | France (85) | 9 | 2022 | AGRI | 0.27 |
| 223 | France (89) | 4 | 2020 | URB | 0.37 |
| 224 | France (89) | 4 | 2020 | AGRI | 0.41 |

| | | | | | |
|---|---|---|---|---|---|
| 225 | France (89) | 4 | 2021 | URB | 0.39 |
| 226 | France (89) | 4 | 2021 | AGRI | 0.26 |
| 227 | France (89) | 6 | 2021 | URB | 0.72 |
| 228 | France (89) | 6 | 2021 | AGRI | 0.46 |
| 229 | France (89) | 9 | 2021 | URB | 0.59 |
| 230 | France (89) | 9 | 2021 | AGRI | 0.48 |
| 231 | France (91) | 6 | 2013 | AGRI | 0.23 |
| 232 | France (91) | 5 | 2015 | AGRI | 0.2 |
| 233 | France (91) | 7 | 2015 | AGRI | 0.19 |
| 234 | France (91) | 9 | 2015 | AGRI | 0.27 |
| 235 | France (91) | 4 | 2019 | AGRI | 0.13 |
| 236 | France (91) | 7 | 2019 | AGRI | 0.35 |
| 237 | France (91) | 9 | 2019 | AGRI | 0.42 |
| 238 | France (91) | 4 | 2020 | AGRI | 0.025 |
| 239 | France (91) | 7 | 2020 | AGRI | 0.32 |
| 240 | France (91) | 9 | 2020 | AGRI | 0.31 |
| 241 | France (91) | 4 | 2021 | AGRI | 0.21 |
| 242 | France (91) | 6 | 2021 | AGRI | 0.29 |
| 243 | France (91) | 9 | 2021 | AGRI | 0.38 |
| 244 | France (91) | 3 | 2022 | AGRI | 0.82 |
| 245 | France (91) | 4 | 2022 | AGRI | 0.49 |
| 246 | France (91) | 5 | 2022 | AGRI | 0.45 |
| 247 | France (91) | 5 | 2022 | AGRI | 0.46 |
| 248 | France (91) | 6 | 2022 | AGRI | 0.50 |
| 249 | France (91) | 6 | 2022 | AGRI | 0.39 |
| 250 | France (91) | 7 | 2022 | AGRI | 0.35 |
| 251 | France (91) | 7 | 2022 | AGRI | 0.41 |
| 252 | France (91) | 8 | 2022 | AGRI | 0.40 |
| 253 | France (91) | 8 | 2022 | AGRI | 0.42 |
| 254 | France (91) | 9 | 2022 | AGRI | 0.44 |
| 255 | France (91) | 9 | 2022 | AGRI | 0.49 |
| 256 | France (91) | 10 | 2022 | AGRI | 0.48 |
| 257 | France (93) | 4 | 2019 | URB | 0.28 |
| 258 | France (93) | 7 | 2019 | NAT | 0.55 |
| 259 | France (93) | 9 | 2019 | AGRI | 0.25 |
| 260 | France (94) | 4 | 2019 | URB | 0.40 |
| 261 | France (94) | 7 | 2019 | NAT | 0.67 |
| 262 | France (94) | 9 | 2019 | AGRI | 0.66 |
| 263 | France (94) | 4 | 2020 | URB | 0.46 |
| 264 | France (94) | 7 | 2020 | NAT | 0.54 |
| 265 | France (94) | 9 | 2020 | AGRI | 0.68 |
| 266 | France (95) | 4 | 2015 | URB | 0.35 |
| 267 | France (95) | 4 | 2015 | NAT | 0.37 |
| 268 | France (95) | 7 | 2015 | URB | 0.46 |
| 269 | France (95) | 7 | 2015 | NAT | 0.27 |
| 270 | France (95) | 9 | 2015 | URB | 0.51 |

| | | | | | |
|---|---|---|---|---|---|
| 271 | France (95) | 9 | 2015 | NAT | 0.26 |
| 272 | France (95) | 3 | 2016 | URB | 0.35 |
| 273 | France (95) | 7 | 2016 | URB | 0.40 |
| 274 | France (95) | 9 | 2016 | URB | 0.385 |
| 275 | Greece | 4 | 2022 | URB | 0.48 |
| 276 | Greece | 4 | 2022 | URB | 0.32 |
| 277 | Greece | 4 | 2022 | INDUS | 0.34 |
| 278 | Greece | 4 | 2022 | INDUS | 0.52 |
| 279 | Greece | 4 | 2022 | NAT | 0.21 |
| 280 | Greece | 4 | 2022 | NAT | 0.22 |
| 281 | Greece | 5 | 2022 | URB | 0.61 |
| 282 | Greece | 5 | 2022 | URB | 0.56 |
| 283 | Greece | 5 | 2022 | INDUS | 0.53 |
| 284 | Greece | 5 | 2022 | INDUS | 0.46 |
| 285 | Greece | 5 | 2022 | NAT | 0.32 |
| 286 | Greece | 5 | 2022 | NAT | 0.26 |
| 287 | Greece | 6 | 2022 | INDUS | 0.23 |
| 288 | Greece | 6 | 2022 | INDUS | 0.38 |
| 289 | Greece | 6 | 2022 | URB | 0.53 |
| 290 | Greece | 6 | 2022 | URB | 0.34 |
| 291 | Greece | 6 | 2022 | NAT | 0.23 |
| 292 | Greece | 6 | 2022 | NAT | 0.38 |
| 293 | Greece | 7 | 2022 | INDUS | 0.28 |
| 294 | Greece | 7 | 2022 | INDUS | 0.29 |
| 295 | Greece | 7 | 2022 | URB | 0.50 |
| 296 | Greece | 7 | 2022 | URB | 0.47 |
| 297 | Greece | 7 | 2022 | NAT | 0.28 |
| 298 | Greece | 7 | 2022 | NAT | 0.29 |
| 299 | Greece | 8 | 2022 | INDUS | 0.32 |
| 300 | Greece | 8 | 2022 | INDUS | 0.32 |
| 301 | Greece | 8 | 2022 | URB | 0.38 |
| 302 | Greece | 8 | 2022 | URB | 0.35 |
| 303 | Greece | 8 | 2022 | NAT | 0.32 |
| 304 | Greece | 8 | 2022 | NAT | 0.32 |
| 305 | Greece | 9 | 2022 | INDUS | 0.43 |
| 306 | Greece | 9 | 2022 | INDUS | 0.25 |
| 307 | Greece | 9 | 2022 | URB | 0.53 |
| 308 | Greece | 9 | 2022 | URB | 0.55 |
| 309 | Greece | 9 | 2022 | NAT | 0.43 |
| 310 | Greece | 9 | 2022 | NAT | 0.25 |
| 311 | Moldova | 3 | 2022 | NAT | 0.50 |
| 312 | Moldova | 4 | 2022 | NAT | 0.67 |
| 313 | Moldova | 5 | 2022 | NAT | 1.20 |
| 314 | Moldova | 6 | 2022 | NAT | 0.93 |
| 315 | Moldova | 7 | 2022 | NAT | 0.53 |
| 316 | Moldova | 8 | 2022 | NAT | 0.30 |

| | | | | | |
|---|---|---|---|---|---|
| 317 | Moldova | 9 | 2022 | NAT | 0.26 |
| 318 | Moldova | 10 | 2022 | NAT | 0.30 |

The analysis of the 318 samples shown in the table SI-1 gives an overall **average value of <u>0.39 ppm for Mo in honey bees</u>.** The distribution of Mo-content within these 318 bee samples are represented in the Figure SI.2. 75% of the bees analyzed display a Mo content in the 0.2-0.5 ppm range.

As shown in Table SI.2, this average content varies according to the environment but the standard deviation is rather high, especially in natural environment. It remains around 0.39 ppm in agricultural environment, slightly decreases to 0.36 ppm in natural environment, but rises slightly to 0.44 ppm in urban or industrial environments, suggesting an anthropogenic contribution in this case.

**Table SI.2.** Average values found for Mo content in bees as a function of their environment.

| | NAT | AGRI | URB | INDUS |
|---|---|---|---|---|
| Nb of samples | 78 | 188 | 40 | 12 |
| Mean Mo (ppm) | 0.364 | 0.386 | 0.463 | 0.363 |
| Standard Deviation SD | 0.291 | 0.207 | 0.162 | 0.102 |

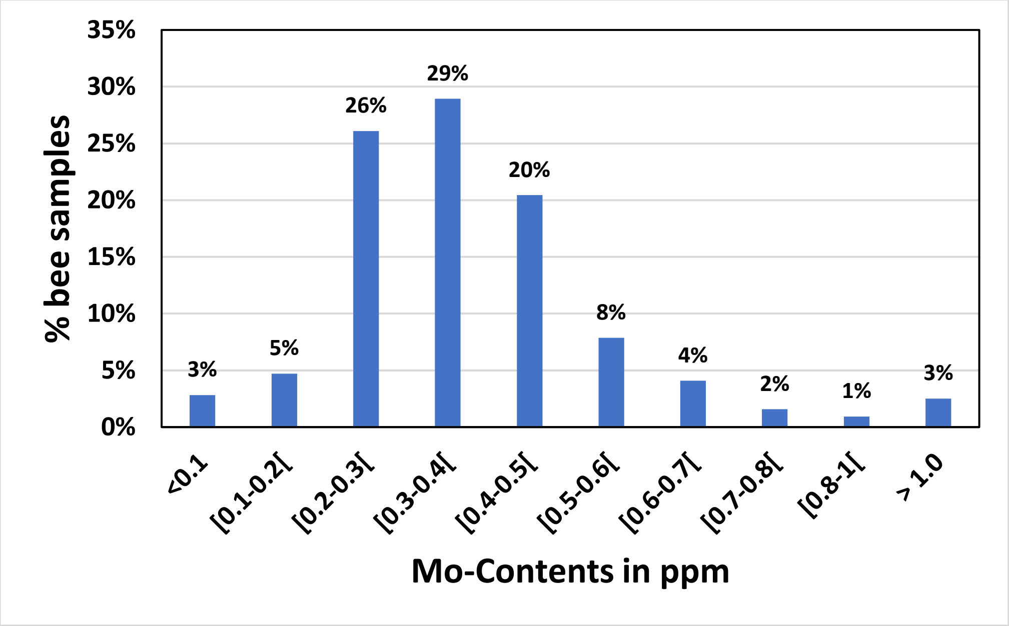


**Figure SI-2.** Representation of the distribution of Mo-content in the 318 bee samples analyzed. The main part, 75%, are found between 0.2 and 0.5 ppm.
