## Supplementary material for "Food supplementation with molybdenum complexes improves honey bee health": Synthesis and stability studies of Molybdenum complexes for this study

### *Supporting Information*

#### Part II. Molybdenum-based complexes.

II.1 Synthesis and characterization

II.2. Stability studies

i. DFT Studies
ii. NMR and electronic spectroscopy studies

### Part II. Molybdenum-based complexes.

#### II.1 Synthesis and characterization

For this study, we focused on complexes of the type $[Mo_2O_2E_2(L)_x]^{n-}$ (E = O or S, x = 1 or 2, n = 0, 2 or 4) in which L is a bidentate, tridentate or hexadentate ligand linked to a central cluster $[Mo_2O_4]^{2+}$ or $[Mo_2O_2S_2]^{2+}$, both well known in the literature. [1] These clusters comprise two Mo(+V) atoms, each possessing a single electron and which is shared in a Mo-Mo bond, rendering these clusters diamagnetic. On each Mo atom, an oxo ligand is present in a Mo=O bond. The coordination sphere of Mo(+V) atoms is completed by L ligands and possibly by water to give distorted octahedral environments. There have been a large number of complexes of this type in the literature since the 1960s and they are generally presented as biomimetic of molybdenum-containing enzymes [2].

Within the framework of this study, we initially considered 12 different compounds depicted in Figure SII.1, which can be isolated either in neutral form with the L-histidine ligand, or in the form of various salts for the others: $Na^+$, $Li^+$, $PPh_4^+$. The syntheses were carried out according to the protocols described by Fuior *et al*. [1] and characterized by routine methods (FT-IR, NMR).

For clarity, the complexes will be abbreviated as « **cation-core-Ligand** ». For example, $Na_2[Mo_2O_4(EDTA)]$ or $K_2[Mo_2O_2S_2(L\text{-}cys)_2]$ will be abbreviated **Na-$Mo_2O_4$-EDTA** and **K-$Mo_2O_2S_2$-LCys**, respectively. In this study, we started with 9 complexes, declined as different salts, mainly alkali, for producing 12 compounds listed in Table SII.1 and we applied a drastic selection to keep the best candidates for the safeguard of bees based on their solubility, toxicity and chemical stability. Note that all complexes are more or less hydrated but for clarity the water molecules are not given in the Table.

***Table SII.1*** *Table of Mo-based complexes initially considered in our study. $TBA^+$ and $PPh_4^+$ correspond respectively to tetrabutylammonium and tetraphenyl phosphonium cations.*

| **Complex** | Salt | Abbreviation |
|---|---|---|
| **$[Mo_2O_2S_2(C_2O_4)_2(H_2O)]^{2-}$** | $Cs^+$, $Na^+$ | **$Cs_{1.5}Na_{0.5}$-$Mo_2O_2S_2$-Ox** |
| **$[Mo_2O_4(C_2O_4)_2(H_2O)]^{2-}$** | $TBA^+$, $Na^+$ | **$TBA_{0.5}Na_{1.5}$-$Mo_2O_4$-Ox** |
| **$[Mo_2O_4(HNTA)_2]^{2-}$** | $K^+$ | **K-$Mo_2O_4$-HNTA** |
| **$[Mo_2O_2S_2(HNTA)_2]^{2-}$** | $K^+$ | **K-$Mo_2O_2S_2$-HNTA** |
| **$[Mo_2O_4(L\text{-}his)_2]$** | - | **$Mo_2O_4$-L-his** |
| **$[Mo_2O_4(L\text{-}Cys)_2]^{2-}$** | $Na^+$ | **Na-$Mo_2O_4$-L-cys** |
| **$[Mo_2O_2S_2(L\text{-}Cys)_2]^{2-}$** | $K^+$ | **K-$Mo_2O_2S_2$-L-cys** |
| **$[Mo_2O_4(EDTA)]^{2-}$** | $PPh_4^+$ | **$PPh_4$-$Mo_2O_4$-EDTA** |
| | $Na^+$ | **Na-$Mo_2O_4$-EDTA** |
| | $Li^+$ | **Li-$Mo_2O_4$-EDTA** |
| **$[Mo_2O_2S_2(EDTA)]^{2-}$** | $PPh_4^+$ | **$PPh_4$-$Mo_2O_2S_2$-EDTA** |
| | $K^+$ | **K-$Mo_2O_2S_2$-EDTA** |

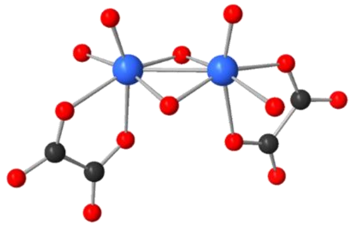

$[Mo^V_2O_4(ox)_2(H_2O)_2]^{2-} = [Mo^V_2O_2S_2(C_2O_4)_2(H_2O)_2]^{2-}$

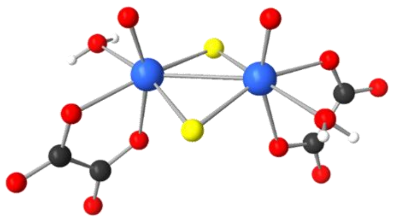

$[Mo^V_2O_2S_2(ox)_2(H_2O)_2]^{2-} = [Mo^V_2O_2S_2(C_2O_4)_2(H_2O)_2]^{2-}$

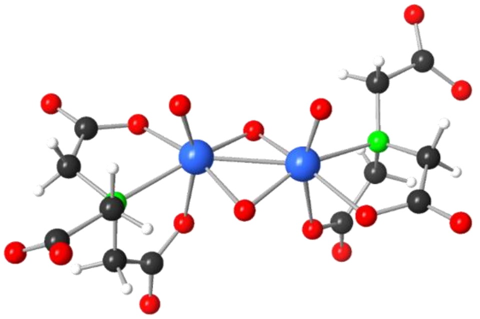

$[Mo^V_2O_4(NTA)_2]^{4-} = [Mo^V_2O_4(NC_6H_6O_6)_2]^{4-}$

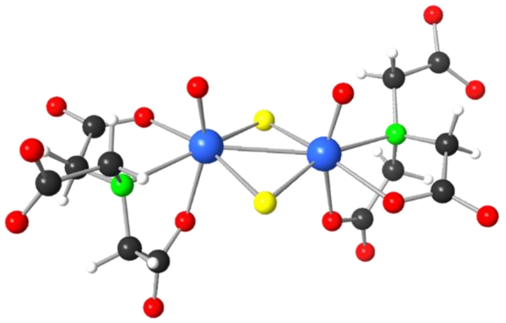

$[Mo^V_2O_2S_2(HNTA)_2]^{2-} = [Mo^V_2O_2S_2(NC_6H_7O_6)_2]^{2-}$

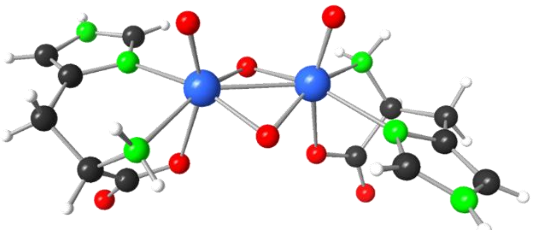

$[Mo^V_2O_4(L\text{-histidine})_2] = [Mo^V_2O_4(C_6H_5N_3O_2)_2]$

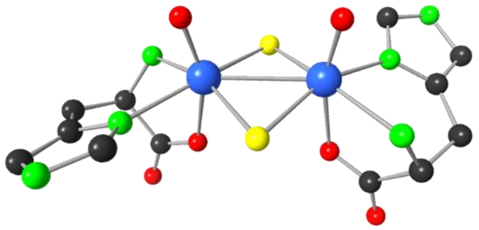

$[Mo^V_2O_2S_2(L\text{-histidine})_2] = [Mo^V_2O_2S_2(C_6H_5N_3O_2)_2]^{2-}$

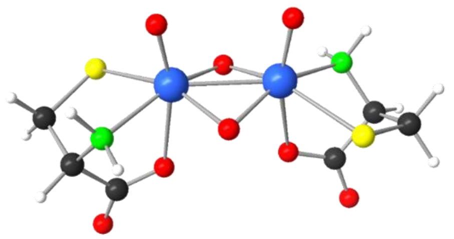

$[Mo^V_2O_4(L\text{-Cysteine})_2]^{2-} = [Mo^V_2O_4(C_3H_5NSO_2)_2]^{2-}$

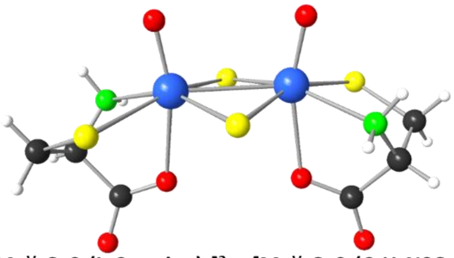

$[Mo^V_2O_2S_2(L\text{-Cysteine})_2]^{2-} = [Mo^V_2O_2S_2(C_3H_5NSO_2)_2]^{2-}$

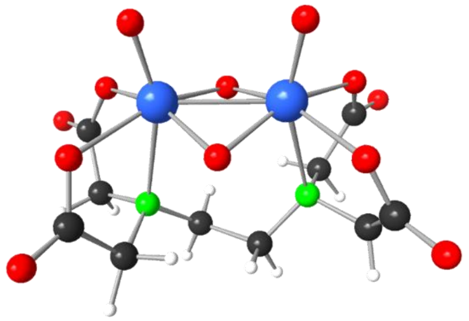

$[Mo^V_2O_4(EDTA)]^{2-} = [Mo^V_2O_4(C_{10}H_{12}O_8N_2)]^{2-}$

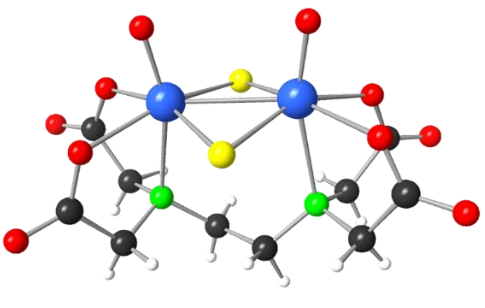

$[Mo^V_2O_2S_2(EDTA)]^{2-} = [Mo^V_2O_2S_2(C_{10}H_{12}O_8N_2)]^{2-}$

***Figure SII.1*** *Structures of $[Mo_2O_2S_2]^{2+}$ and $[Mo_2O_4]^{2+}$ complexes with $H_2ox$, $H_3NTA$, $H_4EDTA$, $H_2L$-cys and HL-his ligands. Color code: Mo(blue), O(red), S(yellow), N(green), C(black), H(white)*

#### II.1 Stability studies

Testing a large panel of complexes in honey bee hives is not realistic given the need to test each on a significant number of bee colonies. Given the low concentrations used in the syrups given to the bees (2 to 8 $mg.L^{-1}$, which corresponds to 3 to $12.10^{-6}$ $mol.L^{-1}$), it appeared necessary to focus on the most chemically stable complexes in solution. To access these chemical stabilities and select the best candidates, a theoretical study by the density functional theory (DFT) was first carried out then completed by experimental studies in water or in sugar syrup by NMR for concentrations up to $5.10^{-5}$ M and by UV-Vis spectroscopy for concentrations up to $2.10^{-7}$ M.

##### i) DFT studies

##### Computational details.

The calculations were based on the density functional theory (DFT) as implemented in the ADF 2016 program [3,4]. The molecular geometries and electronic energies were determined with the hybrid B3LYP functional [5] along with Grimme's dispersion corrections [6] and high numerical accuracy. Since we explore the behaviour of molecules in aqueous solution, we introduced the *conductor-like screening model* (COSMO), a computationally inexpensive recourse to mimic the influence of the liquid solution (the water solvent molecules and the necessary counterions compensating the net charge in case of an ionic solute) upon the solute molecules.[7,8] COSMO treats the liquid phase in an approximate way as if it was a polarizable dielectric continuum material that reacts to the presence of the solute, and vice versa. Hence, water molecules are not explicitly present in the calculations as defined entities and, thus, formation/disruption of the water network cannot be examined.

To describe the electron distribution of the molecules studied, we used atomic basis sets composed of Slater-type functions of triple-$\zeta$ + double polarization (TZ2P) quality for the valence electrons of all atoms. For atoms of the 2nd row of the periodic table and beyond, we applied the frozen core approximation, in which the internal electron shells —non-valence or *core* shells, 1s for C, N and O; 2p for S; 3d for Mo— are kept frozen during the electron density optimization. This approach is applied routinely saving much computational time with no significant loss of accuracy.

To obtain the electronic energies of the molecules analyzed, we first carried out geometry optimization calculations to generate the most stable (lowest energy) atomic arrangement of of the molecules. Such an arrangement corresponds to a minimum in the potential energy hypersurface, which is that of a stable chemical entity.

#### Results

DFT calculations were performed to estimate the stabilities of the complexes by computing their energies and those of the fragments (two or three, depending on the considered starting complex) into which they might break down by a proposed decomposition process. Therefore, optimized molecular geometries and energies were obtained for complexes presented in Figure SII.1 and their corresponding fragments.

The complexes and the resulting fragments are differently charged species: anions, cations or neutral molecules, as shown in Table SII.2. In all cases, the formal charge of Mo is +5 and the $Mo_2O_2E_2$ (E = O or S) moiety carries a charge of +2. Present calculations simulating an aqueous solution largely stabilize the ionic systems in comparison with gas-phase conditions. The fragments into which each complex decomposes in our proposal are listed in Table SII.2 below.

Table SII.2 also contains the molecular energies obtained for complexes and their fragments, as well as the energy increments upon fragmentation (Decomposition Energy: DE, DE = sum of energies of fragments – energy of the complex), which are proposed as a quantification of the relative stability of the complexes.

***Table SII.2*** *Computed molecular energies of the initial complexes and the fragments proposed upon decomposition, and decomposition energies (DE). Values in eV.*

| Complex | Fragment 1 | Fragment 2 | Fragment 3 | DE |
|---|---|---|---|---|
| **$[Mo_2O_4(EDTA)]^{2-}$** | $[Mo_2O_4]^{2+}$ | $(EDTA)^{4-}$ | | |
| Energy : -316.181 | Energy : -49.718 | Energy : -256.59 | | 9.873 |
| **$[Mo_2O_2S_2(EDTA)]^{2-}$** | $[Mo_2O_2S_2]^{2+}$ | $(EDTA)^{4-}$ | | |
| -308.375 | -42.740 | -256.590 | | 9.045 |
| **$[Mo_2O_4(L\text{-}Cys)_2]^{2-}$** | $[Mo_2O_4(L\text{-}Cys)]$ | $(L\text{-}Cys)^{2-}$ | | |
| -241.438 | -146.502 | -91.060 | | 3.876 |
| **$[Mo_2O_2S_2(L\text{-}Cys)_2]^{2-}$** | $[Mo_2O_2S_2(L\text{-}Cys)]$ | $(L\text{-}Cys)^{2-}$ | | |
| -234.150 | -139.165 | -91.034 | | 3.951 |
| **$[Mo_2O_4(NTA)_2]^{4-}$** | $[Mo_2O_4(NTA)]^{-}$ | $(NTA)^{3-}$ | | |
| -379.768 | -215.549 | -160.350 | | 3.869 |
| **$[Mo_2O_2S_2(NTA)_2]^{4-}$** | $[Mo_2O_2S_2(NTA)]^{-}$ | $(NTA)^{3-}$ | | |
| -372.270 | -208.166 | -160.350 | | 3.755 |
| **$[Mo_2O_4(C_2O_4)_2(H_2O)_2]^{2-}$** | $[Mo_2O_4(C_2O_4)(H_2O)]$ | $(C_2O_4)^{2-}$ | $H_2O$ | |
| -219.983 | -135.475 | -63.525 | -17.375 | 3.607 |
| **$[Mo_2O_2S_2(C_2O_4)_2(H_2O)_2]^{2-}$** | $[Mo_2O_2S_2(C_2O_4)(H_2O)]$ | $(C_2O_4)^{2-}$ | $H_2O$ | |
| -212.309 | -127.985 | -63.525 | -17.375 | 3.424 |

We observe that all the DEs are very positive between 3.4 and 9.9 eV (ca. 80 – 230 kcal.mol$^{-1}$) indicating, in principle, that the complexes in solution are stable to highly stable. Since, in general, we are dealing with ions breaking down into other ions, the balance between the charge of the initial and final species is relevant for the DE.

When the decomposition implies the formation of some highly charged fragment (even more than the initial complex), DEs tend to reach more positive values, implying a less favoured breakdown process. This is the case of **$[Mo_2O_4(EDTA)]^{2-}$** and **$[Mo_2O_2S_2(EDTA)]^{2-}$**, with the largest DEs (>9 eV). It is worth mentioning the case of species **$[Mo_2O_4(C_2O_4)_2(H_2O)_2]^{2-}$** and **$[Mo_2O_2S_2(C_2O_4)_2(H_2O)_2]^{2-}$**, with the lowest DEs within the series. Even if the complexes are not highly charged, they decompose into two neutral and one equally charged ion, which appears to be not very energetically penalized compared to others. In addition, these are the smallest complexes, with 24 atoms each, thus their negative charges are more concentrated and thus less stabilized than compounds **$[Mo_2O_4(EDTA)]^{2-}$** and **$[Mo_2O_2S_2(EDTA)]^{2-}$**. As a consistent point, we see that the two compounds with EDTA, by far the ones with the largest DEs, have, as one of the final fragments, the $[Mo_2O_2E_2]^{2+}$ unit, which comparatively penalizes more the breakdown process within this series. Also, all the $Mo_2O_4$-based systems tend to produce more positive or nearly equal DEs, implying that the oxo derivatives are thermodynamically more stable than the thio (E = S) ones. The HOMO-LUMO gaps, which may be viewed as an indicator of molecular stability, tend to be larger in $Mo_2O_4$ derivatives compared with their thio homologues (Table SII.3).

***Table SII.3.*** *Frontier orbital energies and HOMO-LUMO gaps of complexes. Values in eV.*

| Complex | HOMO energy | LUMO energy | HOMO-LUMO gap | DE |
|---|---|---|---|---|
| **$[Mo_2O_4(EDTA)]^{2-}$** | -5.93 | -1.78 | 4.16 | 9.87 |
| **$[Mo_2O_2S_2(EDTA)]^{2-}$** | -6.14 | -2.12 | 4.02 | 9.04 |
| **$[Mo_2O_4(L\text{-}Cys)_2]^{2-}$** | -5.72 | -1.74 | 3.98 | 3.88 |
| **$[Mo_2O_2S_2(L\text{-}Cys)_2]^{2-}$** | -5.70 | -1.96 | 3.74 | 3.95 |
| **$[Mo_2O_4(NTA)_2]^{4-}$** | -5.93 | -1.94 | 3.99 | 3.87 |
| **$[Mo_2O_2S_2(NTA)_2]^{4-}$** | -6.11 | -2.13 | 3.98 | 3.75 |
| **$[Mo_2O_4(C_2O_4)_2(H_2O)_2]^{2-}$** | -6.17 | -2.21 | 3.96 | 3.61 |
| **$[Mo_2O_2S_2(C_2O_4)_2(H_2O)_2]^{2-}$** | -6.31 | -2.43 | 3.88 | 3.42 |

We see no significant correlation between HOMO-LUMO gaps and DEs for the whole series. However, the DEs do show some correlation with the LUMO energies in the sense that higher LUMOs are usually linked with more positive DE values, that is, more stable complexes.

From Table SII.2 showing Des, we logically see that oxalato complexes are the least stable ones, followed by **$[Mo_2O_2E_2(HNTA)_2]^{2-}$**, and then **$[Mo_2O_2E_2(L\text{-}Cys)_2]^{2-}$** in an increasing trend of stability. EDTA complexes are by far more stable with DEs more than two times higher. Figure SII.2 graphically summarizes this trend.

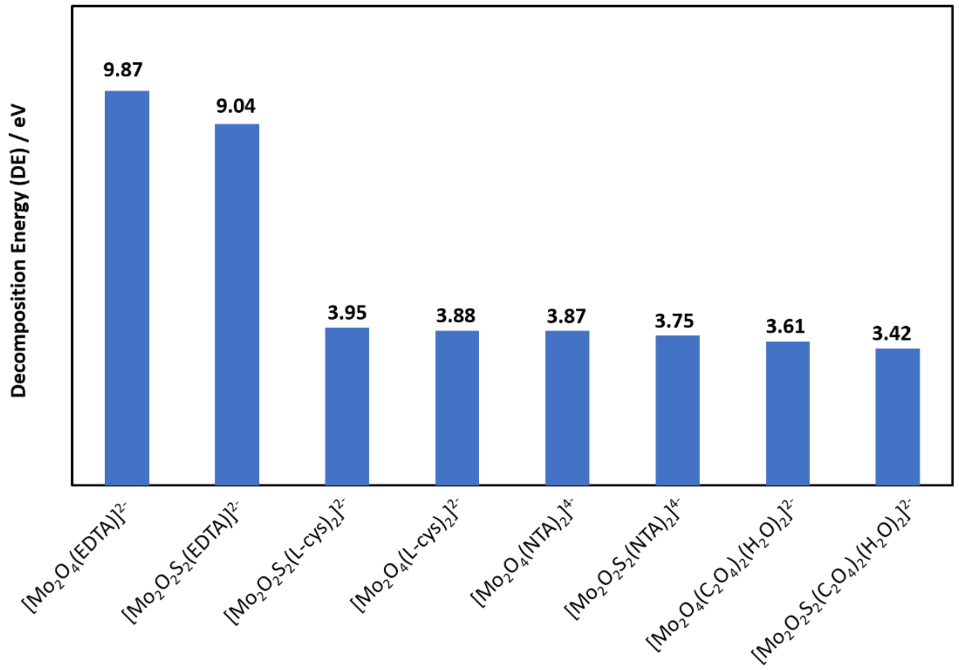


**Figure SII.2** Complexes stabilities ranked in terms of decomposition energies (in eV).

In conclusion, DFT calculations evidence that complex **$[Mo_2O_4(EDTA)]^{2-}$**is the most stable, closely followed by **$[Mo_2O_2S_2(EDTA)]^{2-}$** and further by complex **$[Mo_2O_2S_2(L\text{-}Cys)_2]^{2-}$**, which appears only sigthly more stable than all the other complexes.

#### ii) NMR and electronic spectroscopy studies

#### Materials and methods

**NMR Studies.** Liquid state $^1H$ NMR spectra were recorded in $D_2O$ relative to TMS as external standard (δ = 0 ppm). The experiments were conducted at room temperature on a Bruker Avance 400 spectrometer operating at a resonance frequency of 400 MHz on solutions containing complexes as alkali salts in the concentration range $5.10^{-3}$ to $5.10^{-5}$ mol.L$^{-1}$.

**UV-vis spectra** were recorded on a Perkin-Elmer UV-vis-NIR Lambda-750 spectrometer using calibrated 1 mm, 2 mm, 10 mm, 100 mm Quartz cells. Studies were performed either in deionized water, in phosphate buffer aqueous medium (1,3609g $KH_2PO_4$ + 0.3128g NaOH/200 mL $H_2O$, pH 8.0) or in sugar syrup medium (50% w/w sucrose in water).

#### Results

##### *NMR studies*

Since the **$[Mo_2O_4(L\text{-}his)_2]$** complex is not soluble in aqueous medium, its study in aqueous solution could not be performed. For complexes obtained with oxalato ligands, there is no easy NMR probe ($^1H$) for NMR studies in solution, so that their stability study by NMR was also not possible. Therefore, the stability studies by NMR were focused on alkaki salts of complexes **$[Mo_2O_4(HNTA)_2]^{2-}$**, **$[Mo_2O_2S_2(HNTA)_2]^{2-}$**, **$[Mo_2O_4(L\text{-}Cys)_2]^{2-}$**, **$[Mo_2O_2S_2(L\text{-}Cys)_2]^{2-}$**, **$[Mo_2O_4(EDTA)]^{2-}$**, and **$[Mo_2O_2S_2(EDTA)]^{2-}$**.

The spectra recorded at $5.10^{-3}$M for complexes **$[Mo_2O_4(L\text{-}Cys)_2]^{2-}$**, **$[Mo_2O_4(HNTA)_2]^{2-}$** and **$[Mo_2O_2S_2(HNTA)_2]^{2-}$** display a significant NMR peak assigned to uncoordinated ligands. It evidences that even at this relatively high concentration these three complexes are already partially decomposed (Figures SII.3 and SII.4) and according to Ostwald law, the decoordination will increase at lower concentrations.

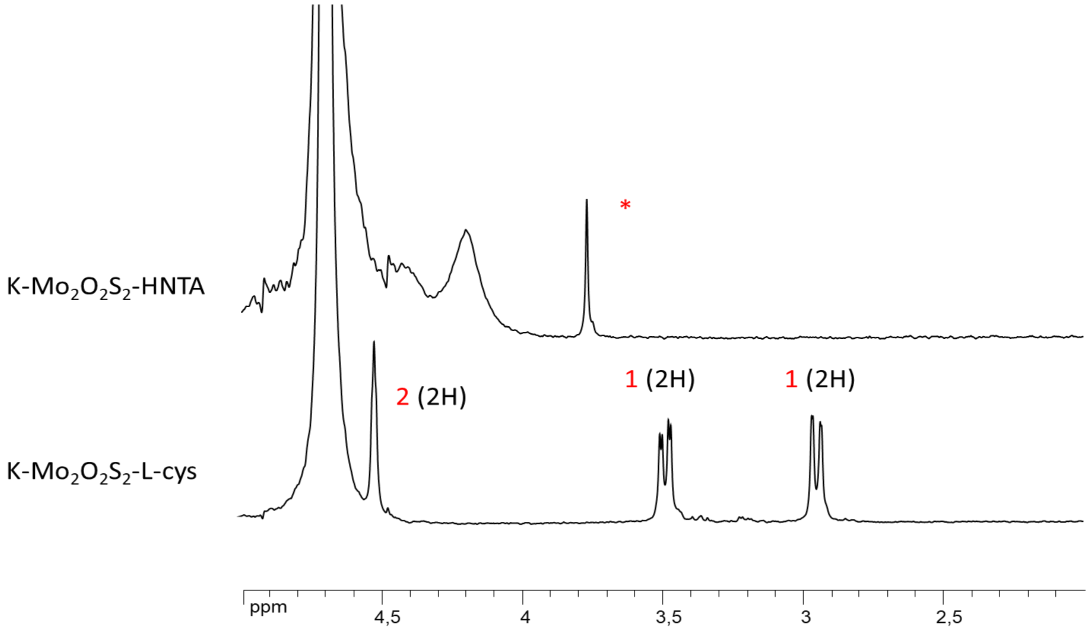


***Figure SII.3*** *$^1$H-NMR spectra of $Mo_2O_2S_2$-based complexes in $D_2O$ at $5.10^{-3}$ M. * indicates the presence of some uncoordinated ligand.*

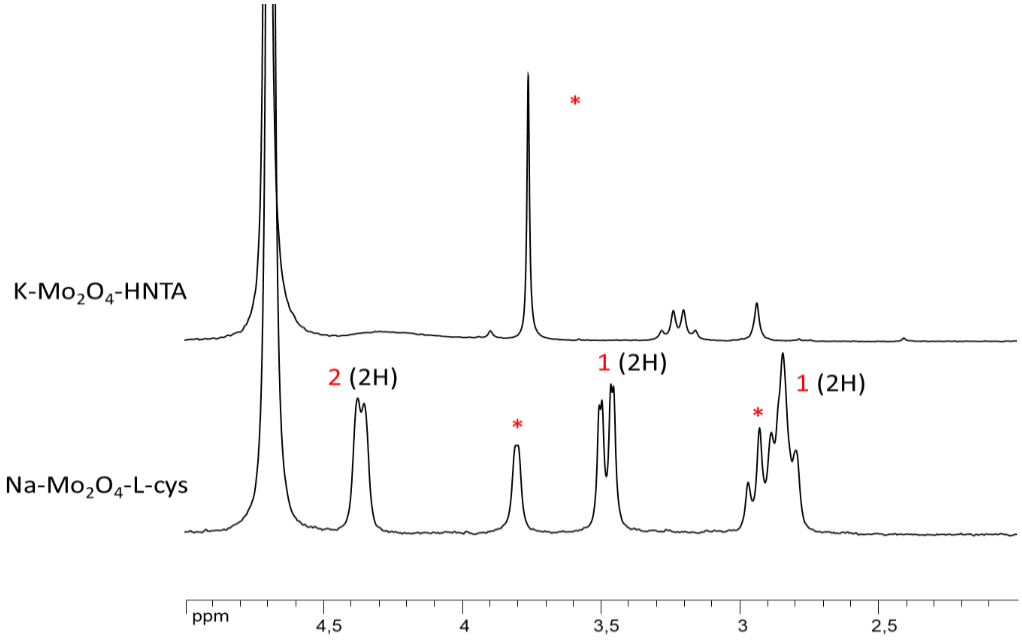


***Figure SII.4*** *$^1$H-NMR spectra of $Mo_2O_4$-based complexes in $D_2O$ at $5.10^{-3}$ M. * indicates the presence of some uncoordinated ligand.*

In contrast, the $^1H$ NMR spectra of the complexes **$[Mo_2O_2S_2(L\text{-}Cys)_2]^{2-}$**, **$[Mo_2O_4(EDTA)]^{2-}$, and $[Mo_2O_2S_2(EDTA)]^{2-}$** display only the signals of complexes at $5.10^{-3}$ M (Figures SII.5-SII.7), according to a higher chemical stability in solution of these complexes, in agreement with DFT calculations. For these three complexes, we thus followed the evolution of $^1H$-NMR spectra of the complexes in aqueous solutions while progressively diluting them more until $5.10^{-5}$ M, which corresponds to the limit of detection by NMR for these complexes (Figures SII.5-SII.7).

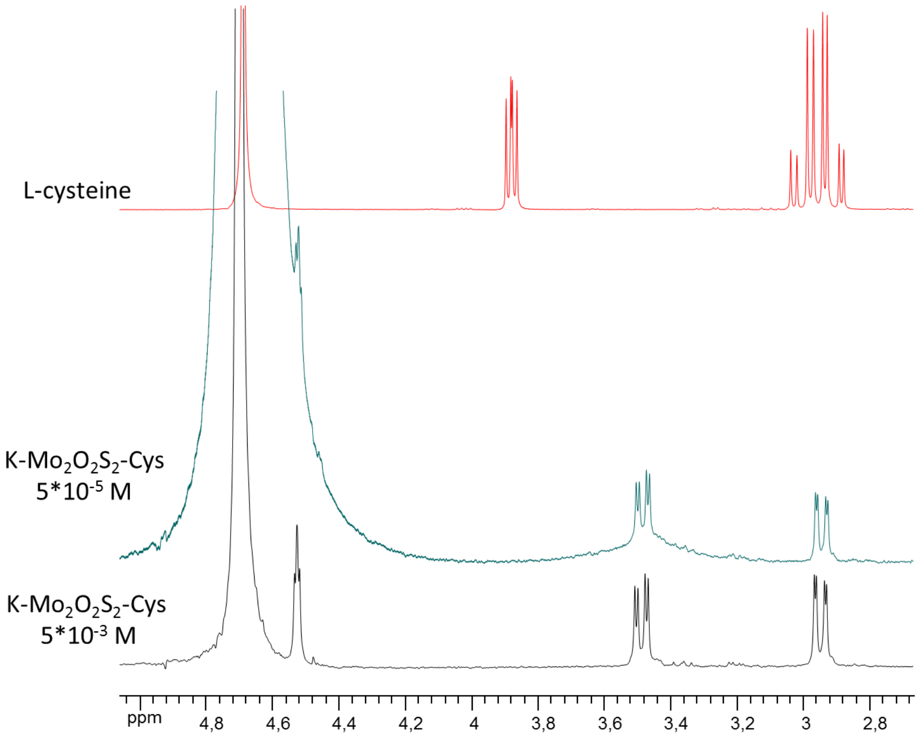


***Figure SII.5.*** *$^1H$-NMR spectra of K-$Mo_2O_2S_2$-Cys at $5*10^{-3}$ and $5*10^{-5}$ M in comparison with free ligand in $D_2O$*

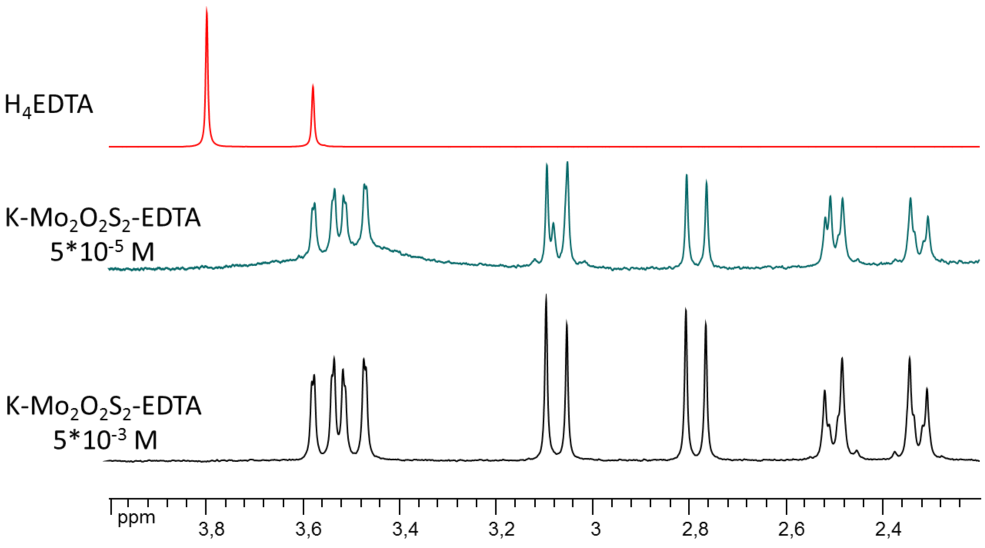


***Figure SII.6.*** *$^1H$-NMR spectra of K-$Mo_2O_2S_2$-EDTA at $5*10^{-3}$ and $5*10^{-5}$ M in comparison with free ligand in $D_2O$*

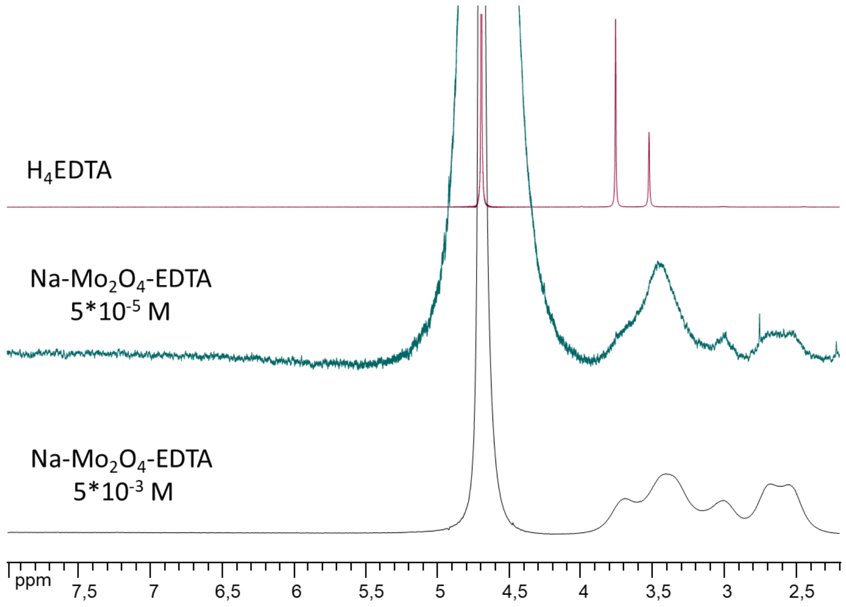


***Figure SII.7.*** *[1]H-NMR spectra of Na-$Mo_2O_4$-EDTA at $5*10^{-3}$ and $5*10^{-5}$ M in comparison with free ligand in $D_2O$*

Figures SII.5-SII.7 show that compounds **K-$Mo_2O_2S_2$-Cys, K-$Mo_2O_2S_2$-EDTA** as well as **Na-$Mo_2O_4$-EDTA** show no sign of instability in the $5.10^{-3}$ – $5.10^{-5}$ M range since the spectra remain similar without appearance of any free ligand. Their integrity is maintained. The three complexes **$[Mo_2O_2S_2(L\text{-}Cys)_2]^{2-}$**, **$[Mo_2O_4(EDTA)]^{2-}$**, and **$[Mo_2O_2S_2(EDTA)]^{2-}$** thus appear to be the most stable in water in the $5.10^{-3}$ – $5.10^{-5}$ M, according to DFT calculations.

***Electronic spectroscopy studies***

The complexes formed with clusters $[Mo_2O_4]^{2+}$ and $[Mo_2O_2S_2]^{2+}$ are orange colored due to Ligand to Metal Charge Transfer (LMCT) bands. Even if the absorption coefficients are not very high, by playing on the length of the cell up to 10 cm, we can measure electronic spectra up to $2.10^{-7}$ $mol.L^{-1}$, which is of the same order as the concentration used for tests in honey bee hives.

We focused our attention on complexes **K-$Mo_2O_2S_2$-EDTA**, **Na-$Mo_2O_4$-EDTA** and **K-$Mo_2O_2S_2$-Cys** in water, then in phosphate buffer medium and finally in a sugar syrup. For each, the data were plotted as a $\varepsilon = f(\lambda)$ function in the 200-800 nm range. The spectra recorded in water are depicted in Figures SII.8-SII.10.

Looking at the absorption spectra of **K-$Mo_2O_2S_2$-Cys** in water (Figure SII.8), we notice a strong change in ε value at $5.10^{-6}$ M and higher, which thus corresponds to the limit od stability for this complex. The other two complexes with EDTA ligand are far more stable (Figures SII.9 and SII.10).

Figure SII.10 shows changes at $2.10^{-7}$ M in the middle-UV region of wavelength (200 – 300 nm) for **Na-$Mo_2O_4$-EDTA**, which is slightly more stable than the sulfido-bridged analogue **K-$Mo_2O_2S_2$-EDTA** (Figure SII.9). Even though the absorption spectra show changes in ε values.

the characteristic $\lambda_{max}$ values do not shift while going extremely low in concentration and they agree with published data: **Na-Mo$_2$O$_4$-EDTA** ($\lambda_{max}$ = 298 nm. 392 nm), **K-Mo$_2$O$_2$S$_2$-EDTA** ($\lambda_{max}$ = 278nm. 308(sh) nm. 347(sh) nm. 468(sh) nm) [2]. This means that the Mo$_2$O$_2$E$_2$-EDTA compounds do not undergo any structural modifications at 2.10$^{-7}$ M.

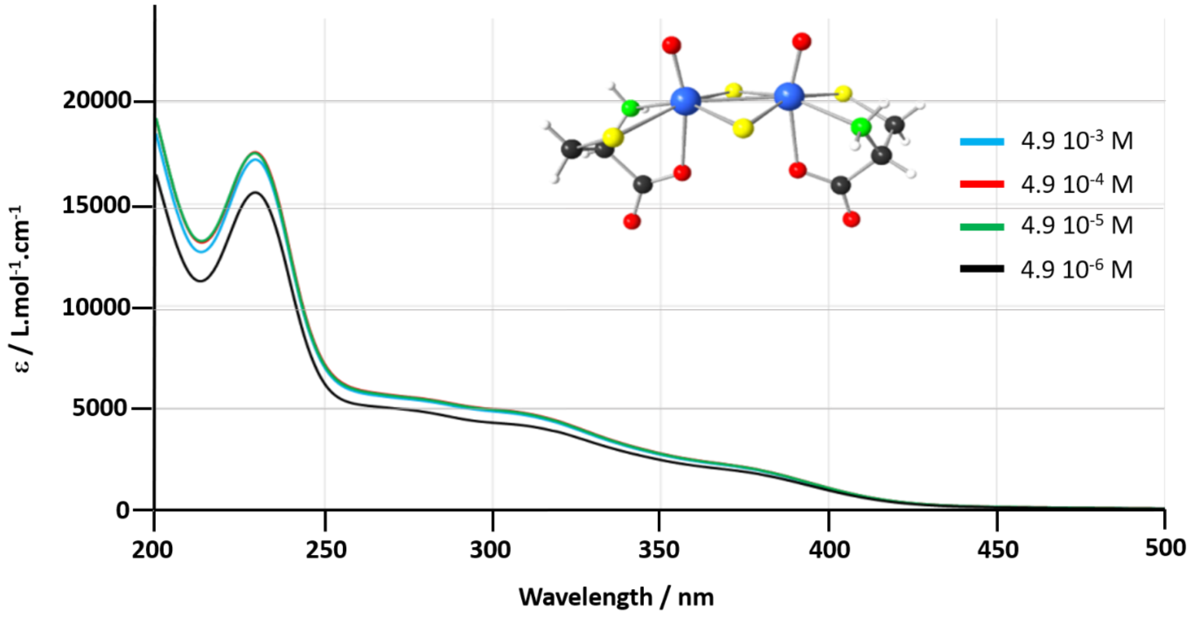


***Figure SII.8.*** *Absorption spectra of K-Mo$_2$O$_2$S$_2$-Cys in diluted water solutions*

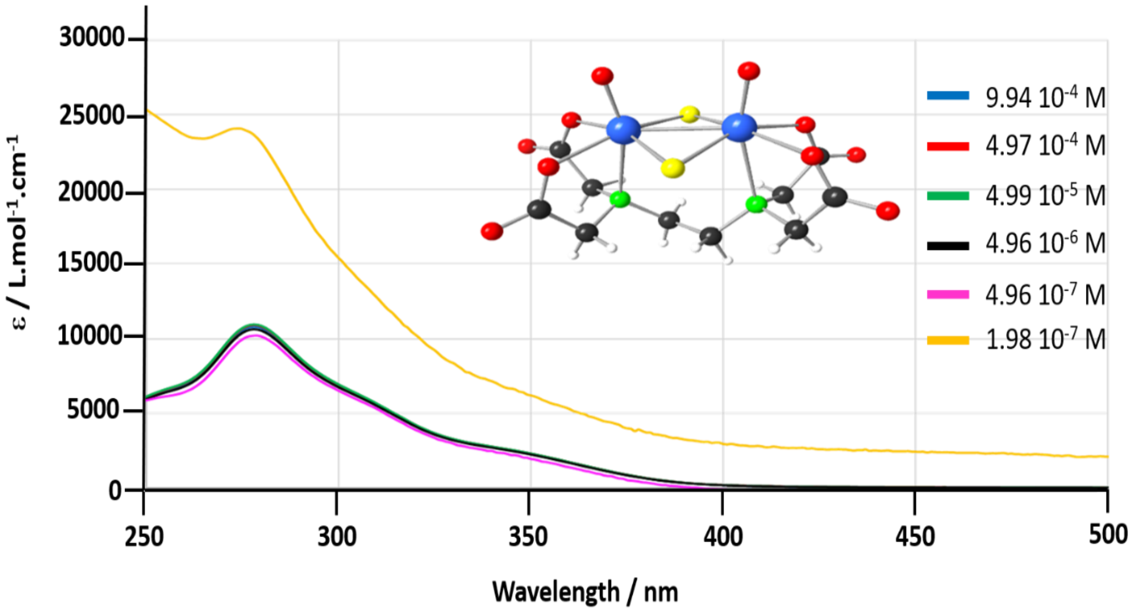


***Figure SII.9.*** *UV-vis absorption spectra of K-Mo$_2$O$_2$S$_2$-EDTA in diluted water solutions*

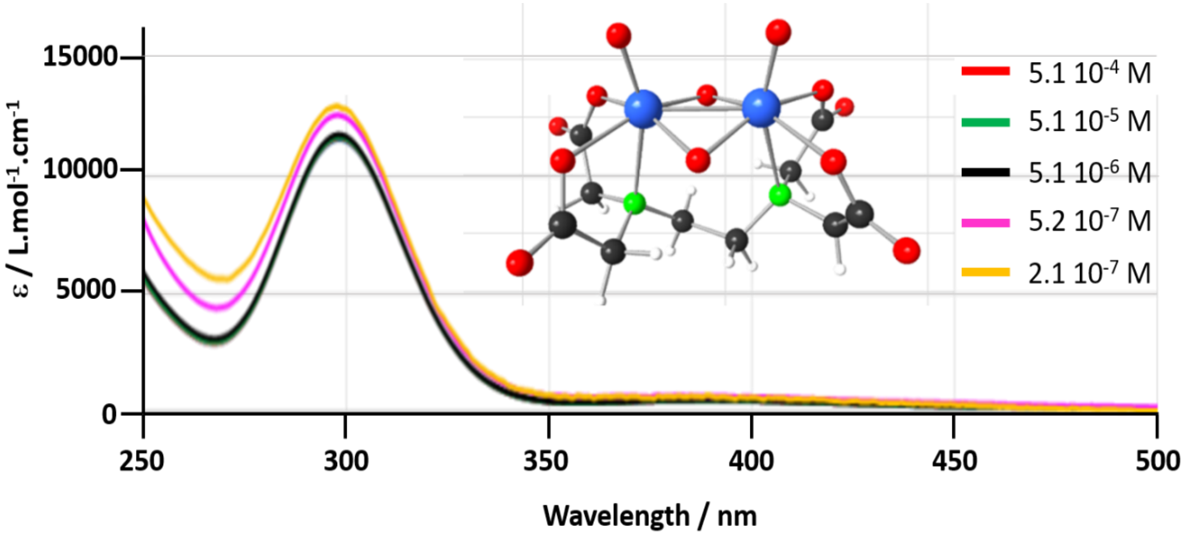


***Figure SII.10.*** *Absorption spectra of Na-$Mo_2O_4$-EDTA in diluted water solutions*

To go further, Figure SII.11 compares the spectra recorded for **Na-$Mo_2O_4$-EDTA** in water, in phosphate buffer solution (pH = 8, physiological medium) and in a sugar syrup containing 50% (w/w) of sucrose. The absorption coefficients slightly vary as a function of the medium but the spectra remain unchanged in all three media at $5.10^{-5}$ M and even lower concentrations, unambigously proving that the **$[Mo_2O_4(EDTA)]^{2-}$** complex is perfectly stable in these media.

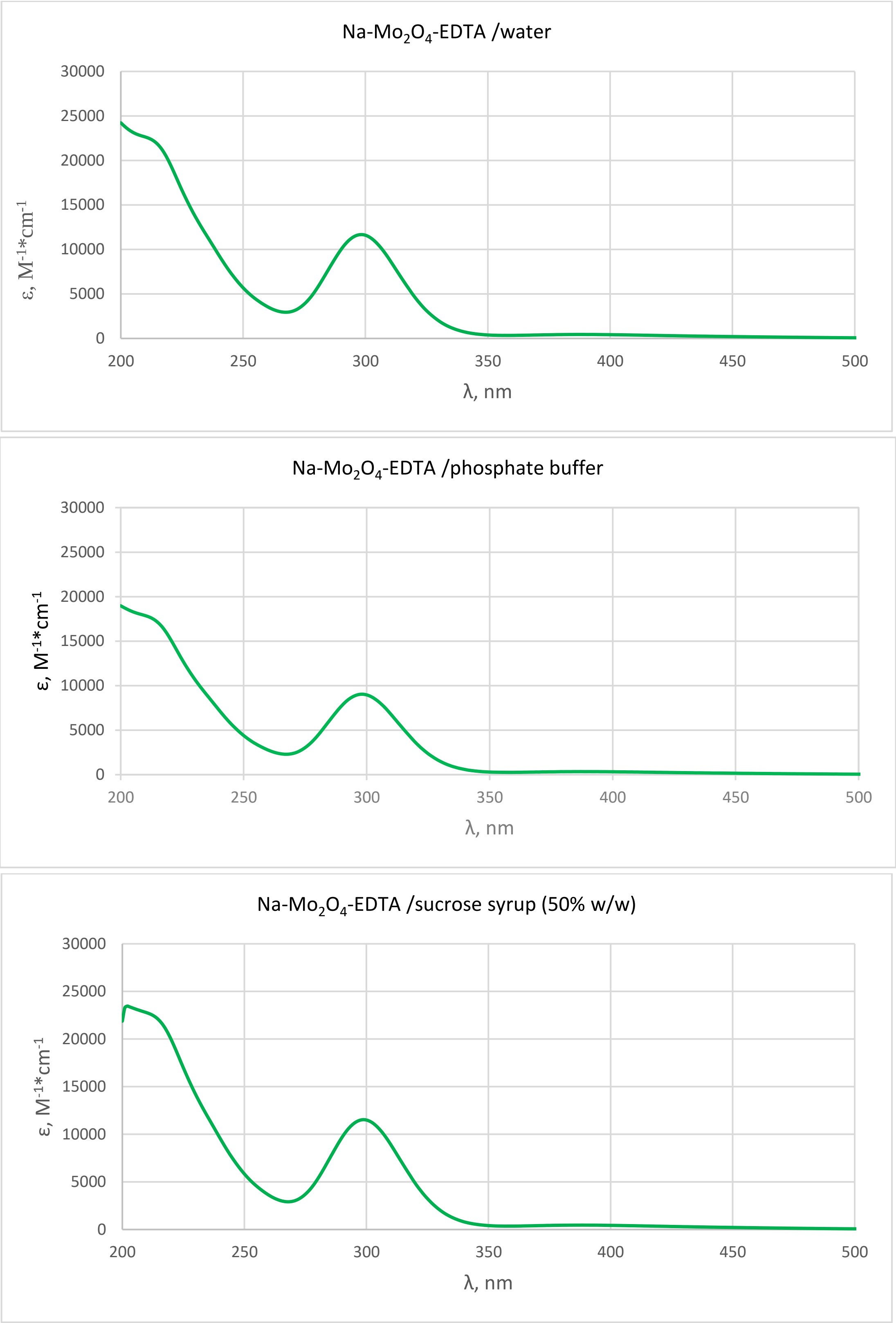


***Figure SII.11*** *UV-vis absorption spectra for 5.10$^{-5}$M* ***Na-$Mo_2O_4$-EDTA*** *in water, phosphate buffer and sucrose syrup (50% w/w).*

#### Conclusion

In summary, the chemical stability of our complexes is found in the order: **$[Mo_2O_2S_2(L\text{-}Cys)_2]^{2-}$** (decomposition starting around $4.9.10^{-6}$ M) < **$[Mo_2O_2S_2(EDTA)]^{2-}$** ($4.10^{-7}$ M) < **$[Mo_2O_4(EDTA)]^{2-}$** (< $2.10^{-7}$ M), in full agreement with DFT calculations.

**$[Mo_2O_4(EDTA)]^{2-}$** is the most stable in water. These limit concentrations are perfectly compatible with the concentration used in syrup given the honey bee hives (3 to 12 µM)
