## Supplementary material for "Food supplementation with molybdenum complexes improves honey bee health": Toxicity studies (Mice, Daphnia, Bees)

### *Supporting Information*

#### Part III. Toxicity studies

III.1- Toxicity on mice
III.2- Toxicity on *Daphnia Magna*
III.3- Acute toxicity on bees
III.4- Chronic toxicity on bees: mortality studies
III.5. Tolerance studies in beehives

### Part III. Toxicity studies

In a previous paper, we reported the biological properties of our molybdenum complexes concerning paramecia, fungi, and bacteria [1]. We examined their antioxidant characteristics (ABTS and DPPH methods) and their cytotoxicity on normal kidney cells (MDCK), pancreatic cancer cells (BxPC-3), cervical cancer cells (HeLa), and rhabdomyosarcoma cancer cells (RD). Our study revealed that these complexes either exhibited no toxicity or only minimal toxicity. In particular, a moderate bio-activity was measured for $PPh_4^+$ salts, due to this counter cation but not related to the molybdenum complexes.

To go further and to envision the use of such complexes as food additives for animals like honey bees, we need to complete these studies. First, potential toxicity was evaluated on mice for the compounds **K-$Mo_2O_2S_2$-HNTA, Na-$Mo_2O_4$-L-Cys, K-$Mo_2O_2S_2$-L-Cys, K-$Mo_2O_2S_2$-EDTA**, **Na-$Mo_2O_4$-EDTA** and **Na-$Mo_2O_4$-EDTA.** The study of the salt **$PPh_4$-$Mo_2O_4$-EDTA**, which possesses antibacterial and antifungal properties, was unfortunately not possible on mice because of the low solubility of this salt in water.

Toxicity was then evaluated on the water flea *Daphnia Magna* for **Na-$Mo_2O_4$-EDTA** and **Li-$Mo_2O_4$-EDTA** in comparison with sodium molybdate as reference as the most common commercial source of Molybdenum. This test permits to evaluate the impact that these complexes would have when present in rivers and more globally in the environment.

Finally, the acute and the chronic toxicities of these two complexes (**Na-$Mo_2O_4$-EDTA** and **Li-$Mo_2O_4$-EDTA**) was evaluated directly on honey bees.

### III.1- Toxicity on mice

#### III.1.1 Experimental procedures

**1°) Study design of acute peroral toxicity**

To study the toxicity of solutions containing the molybdenum complexes, CBA mice of both sexes, with a weight of 20±3 g, were randomly divided into 2 experimental groups and 1 control group (6 mice in each group).

In the first group, an evaluation of the tested substances with prolonged use was carried out. Mice received solutions of molybdenum complexes (concentration 50 mg/mL) at a dose of 0.5 g/kg once a day for 5 consecutive days. Each time the solution was given in a volume of 200 µL. The cumulative dose was 2.5 g/kg.

In the second group, solutions were administered in 1 day only. Each time the solution was given in a volume of 400 µL. The maximal dose of the test substances was 3 g/kg (drugs were administered three times at a dose of 1 g/kg with a break of 2 hours).

The control groups received sterile water in the same volume and with the same administration regime.

The tested solutions were dissolved in sterile water and administered to mice in the esophagus through a 1 mm diameter metal probe. The mice were deprived of food 2 hours before the procedure to empty their stomach, but had free access to water.

After the drug administration, mice were given free access to water and food. The behavior of the mice was observed daily, as well as possible changes in body weight.

Two weeks after the administration of the solutions (for group 1 counting started after the fifth injection), the mice were sacrificed by vertebral-cranial dislocation. Blood was collected from the mice, as well as organs such as the lungs, heart, kidneys, liver, spleen, stomach, small intestine and large intestine. The distribution of molybdenum was studied on the collected organs, using inductively coupled plasma atomic emission spectrophotometry (ICP-AES), and morphological changes in the organ's tissues were assessed.

**2°) Morphological Analysis**

The tissues of lungs, heart, kidneys, liver, spleen, stomach, small intestine and large intestine recovered from the necropsy were fixed in 10% formalin, embedded in paraffin, sectioned, and stained with hematoxylin and eosin (HE) for histological examination using standard techniques. After HE staining, the slides were observed and images were taken using an optical microscope (AxioImager 40, Carl Zeiss Inc., Jena, Germany).

**3°) Biodistribution of Molybdenum in CBA mice**

The blood samples were collected using a standard ocular vein blood collection technique [2] and samples of different inner organs (lungs, heart, kidneys, liver, spleen, stomach, small intestine and large intestine) from all the mice were obtained and weighed for the Mo determination. These samples were digested at 120 °C in 1 mL of a mixture of nitric acid and hydrogen peroxide (3:1 v/v) for 2 h, until the solution became transparent. Deionized water was then added to reach a volume of 5 mL before the Mo determination. The Mo concentration was measured by ICP-AES. The concentrations obtained were normalized to 1 g of weight of the tissue sample. Data are expressed as ng/g fresh tissue.

**4°) Animals and housing Conditions**

The study protocol was approved by the Ethics Committee of the Research Institute of Clinical and Experimental Lymphology – Branch of the Institute of Cytology and Genetics, Siberian Branch of Russian Academy of Sciences RICEL-branch of ICG SB RAS.

All animal procedures were carried out in accordance with the protocols approved by the Bio-ethics committee of the Siberian Branch of the Russian Academy of Sciences, recommendations for the proper use and care of laboratory animals (European Communities Council Directive 86/609/C.E.E.), and the principles of the Declaration of Helsinki.

Mice were housed in stainless steel cages containing sterile sawdust husk as bedding in ventilated animal rooms with free access to water and a commercial laboratory complete food. Animals were acclimatized for 1 week prior to the experiments.

#### III.1.2 Results

**1°) General characterization of the solution of study substances**

A concentration of 50 mg.mL$^{-1}$ is very high (around 7.5.10$^{-2}$ M). Figure SIII.1 shows the resulting solutions of the tested compounds: **K-$Mo_2O_2S_2$-EDTA** is a suspension, **Na-$Mo_2O_4$-L-Cys** and **Na-$Mo_2O_4$-EDTA** are limpid solutions, **K-$Mo_2O_2S_2$-L-Cys** and **K-$Mo_2O_2S_2$-HNTA** have a precipitate and need to be sonicated prior to giving them to mice. **Li-$Mo_2O_4$-EDTA** (not shown) is highly soluble and also gives a limpid solution.

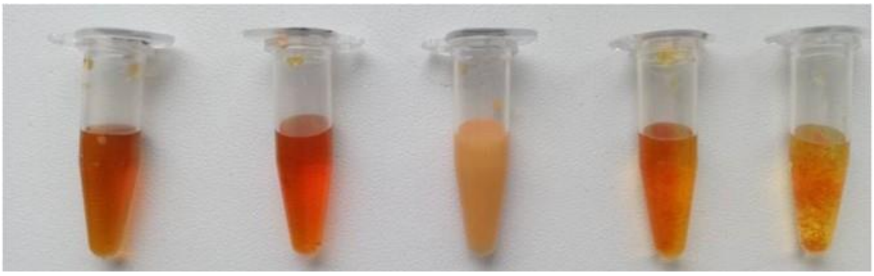

***Figure SIII.1.*** *Solutions of the studied substances, from left to right:* ***Na-$Mo_2O_4$-EDTA, Na-$Mo_2O_4$-L-Cys, K-$Mo_2O_2S_2$-EDTA, K-$Mo_2O_2S_2$-L-Cys, K-$Mo_2O_2S_2$-HNTA;*** *Conc. 50 mg/mL.*

**2°) Acute peroral toxicity**

In the first group of animals (0.5 g/kg for 5 days), the animals' state was satisfactory, the mice were mobile, and no weight loss was observed. After a one-time administration of 3 g/kg of substances **K-$Mo_2O_2S_2$-HNTA** and **Na-$Mo_2O_4$-L-Cys** mice were found to be sluggish and a loss of body weight of 5 ± 0.7% was observed for the first two days. In addition, 50% of the animals died in these first two days. Later, the remaining 50% of animals in this group began to gain weight, and to actively eat and drink. Thus, it was determined that the $LD_{50}$ for **K-$Mo_2O_2S_2$-HNTA** and **Na-$Mo_2O_4$-L-Cys** is 3 g/kg.

After the administration of **K-$Mo_2O_2S_2$-L-Cys**, **K-$Mo_2O_2S_2$-EDTA**, **Na-$Mo_2O_4$-EDTA**, and **Li-$Mo_2O_4$-EDTA,** at a dose of 3 g/kg the condition of animals was satisfactory and no weight loss was observed. $LD_{50}$ of these compounds could not be determined due to their low oral toxicity. In both treated groups (2.5 g/kg over 5 days and 3 g/kg for 1 day), the animals' state was satisfactory, the mice were mobile and no weight loss was observed. $LD_{50}$ of these compounds could not be determined due to their low oral toxicity. It was noted that the animals that received **K-$Mo_2O_2S_2$-L-Cys** and **K-$Mo_2O_2S_2$-EDTA** showed increased activity.

Table SIII.1 summarizes the results and shows that the two compounds **Na-$Mo_2O_4$-EDTA** and **Li-$Mo_2O_4$-EDTA**, are perfectly soluble, not toxic and do not have any side effect on mice.

***Table SIII.1.*** *Toxicity of molybdenum complexes on mice. NL: No Lethality in our conditions or $LD_{50}$ > 3g/kg*

| Compounds | $LD_{50}$ | Comments |
|---|---|---|
| **K-$Mo_2O_2S_2$-HNTA** | 3 g/kg | Suspension |
| **Na-$Mo_2O_4$-L-Cys** | 3 g/kg | Perfectly soluble |
| **K-$Mo_2O_2S_2$-L-Cys** | NL | Suspension / Increased activity of mice |
| **K-$Mo_2O_2S_2$-EDTA** | NL | Suspension / Increased activity of mice |
| **Na-$Mo_2O_4$-EDTA** | NL | Perfectly soluble |
| **Li-$Mo_2O_4$-EDTA** | NL | Perfectly soluble |

**3°) Morphological analysis for compounds K-$Mo_2O_2S_2$-L-Cys, Na-$Mo_2O_4$-EDTA, K-$Mo_2O_2S_2$-EDTA, K-$Mo_2O_2S_2$-HNTA and Na-$Mo_2O_4$-L-Cys.**

Morphological analyses were performed on the organs of mice from the groups that received solutions in the dose 3 g/kg since these are the only groups in which the death of animals has occurred. Mice that received solutions of **K-$Mo_2O_2S_2$-L-Cys**, **Na-$Mo_2O_4$-EDTA** and **K-$Mo_2O_2S_2$-EDTA** did not show any severe changes in organ morphology compared to control organs (Table SIII.2). These data suggest that these 3 solutions are harmless for mice after peroral injection at a dose of 3 g/kg.

***Table SIII.2.*** *Organ morphology in mice that received solutions of* ***K-Mo$_2$O$_2$S$_2$-L-Cys****,* ***Na-Mo$_2$O$_4$-EDTA*** *and* ***K-Mo$_2$O$_2$S$_2$-EDTA*** *at a dose of 3 g/kg and in control mice. There are no significant morphology changes in any of the investigated organs, which correlates with the fact that no mice died in any of these groups. Magnification: small intestine x200, large intestine x200, kidney x200, spleen x100, stomach x200, liver x200, heart x400.*

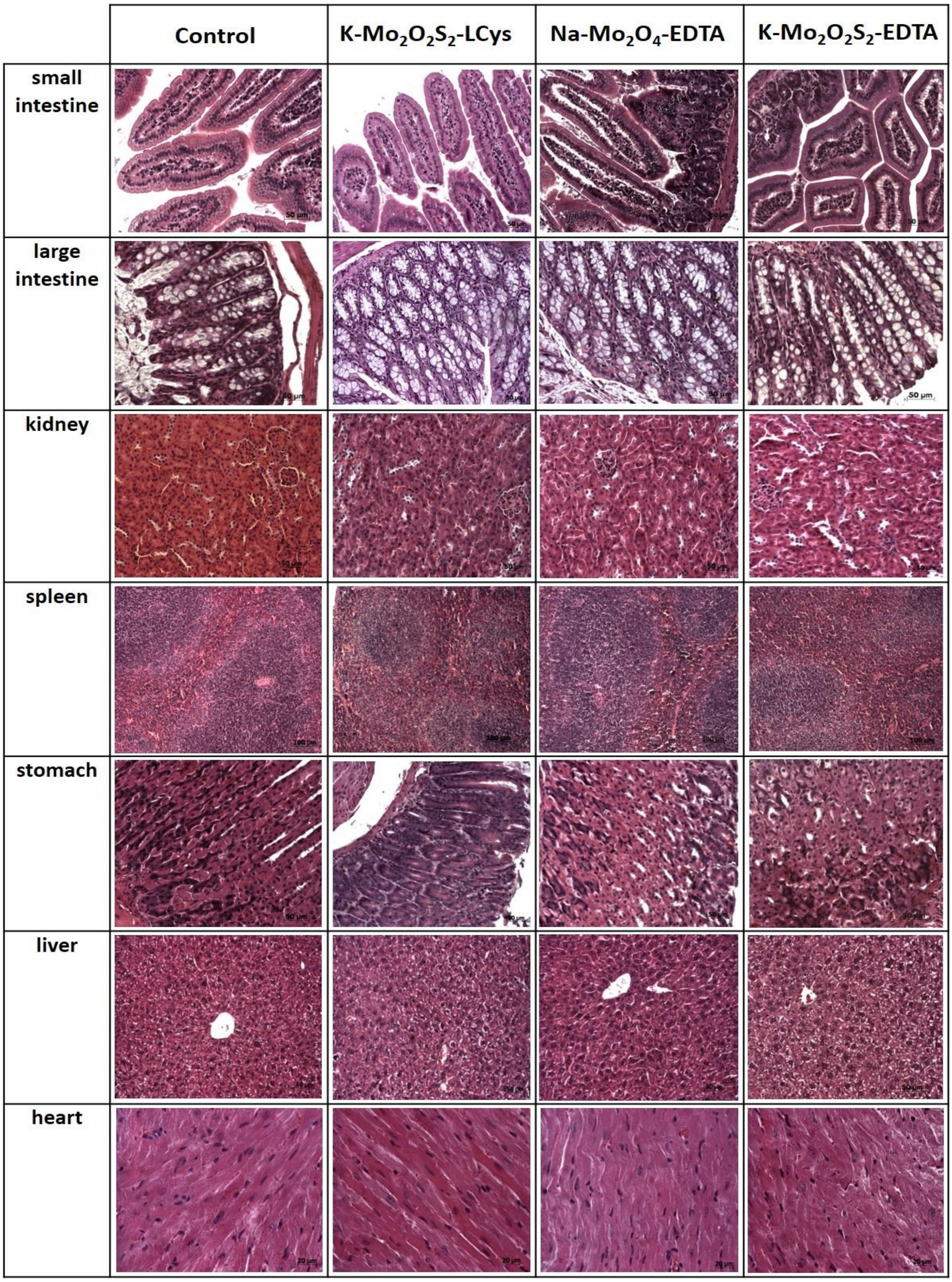

In the group of mice that received 3 g/kg of **K-$Mo_2O_2S_2$-HNTA,** we observed moderate to massive vacuolar cell dystrophy of hepatocytes with foci of necrotic cells (Figure SIII.2). In this group, 50% of the mice died within 2 days. It is apparent that mice were suffering from acute liver deficiency, which was most likely the cause of death. Liver deficiency often causes subsequent kidney disorder. Indeed, the kidneys of mice from this group demonstrate foci of nephronecrosis, which indicates that kidney function was also compromised (Figure SIII.3). The other organs of animals from this group were not affected by the solutions (Figure SIII.4).

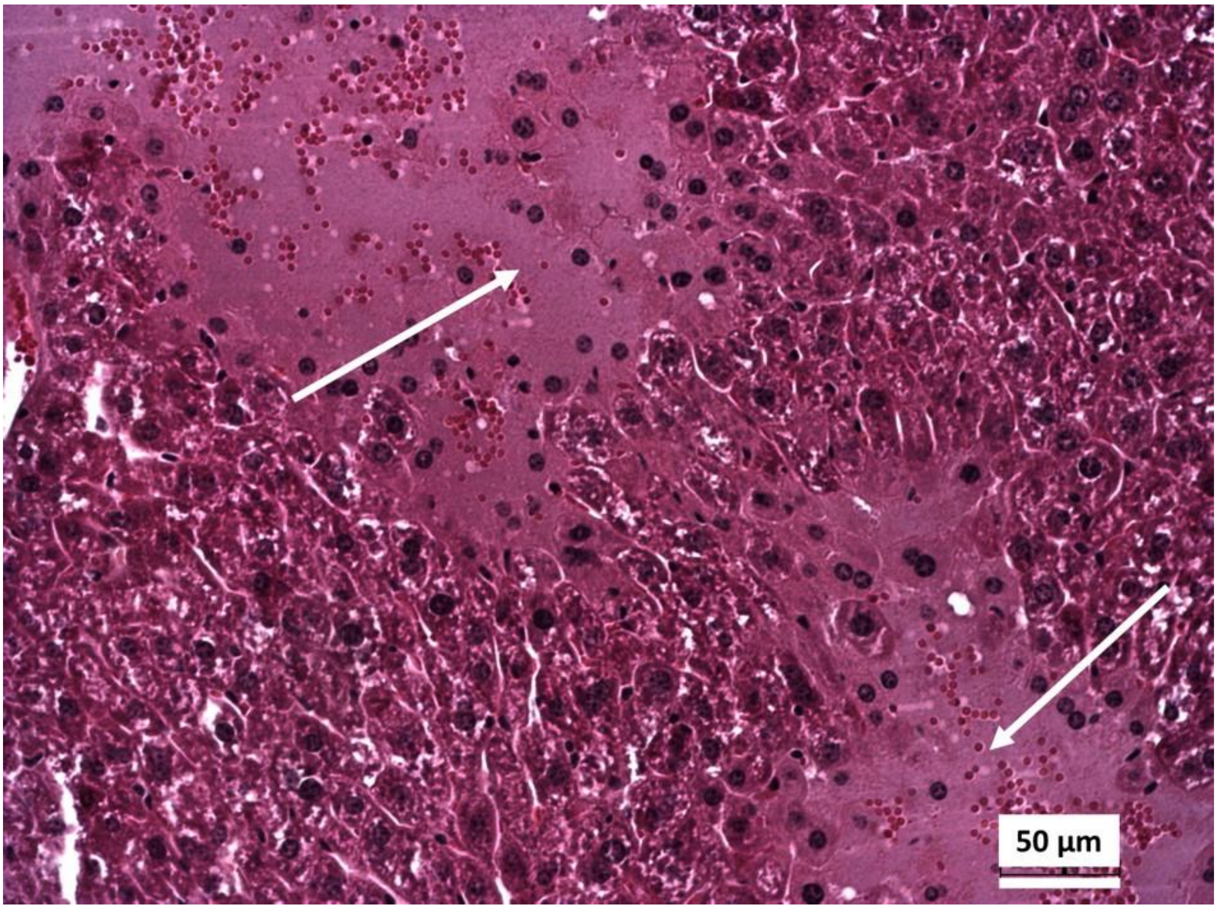


***Figure SIII.2.*** *Liver of a mouse after injection of **K-$Mo_2O_2S_2$-HNTA** at a dose of 3 g/kg. Massive necrosis foci are observed (white arrows). Hepatocytes around the foci are demonstrating moderate cell dystrophy. Magnification x200.*

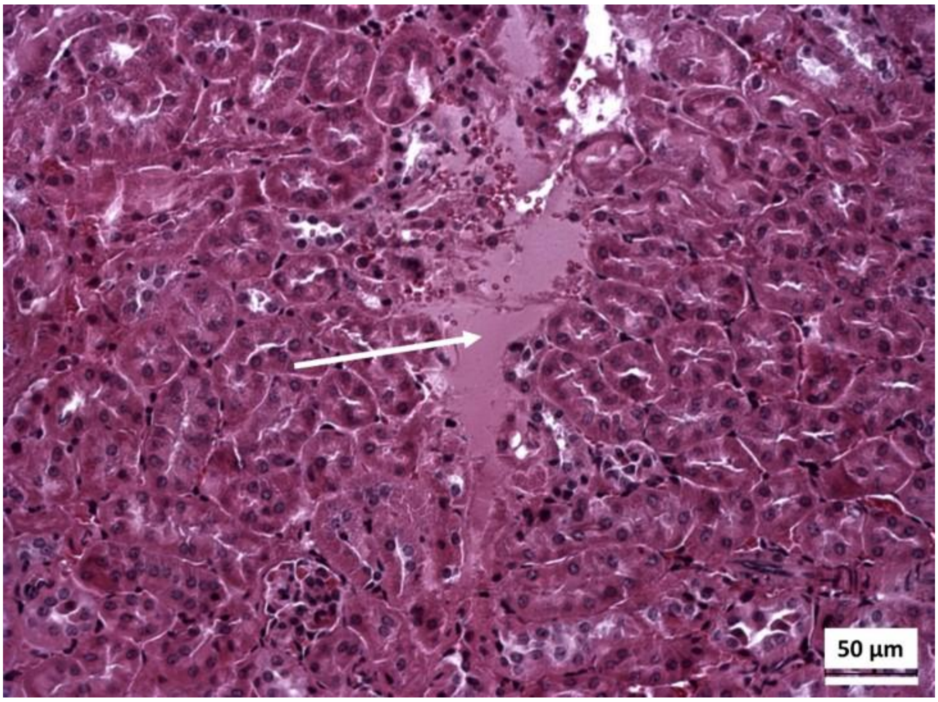


***Figure SIII.3.*** *Kidney of a mouse after injection of* ***K-$Mo_2O_2S_2$-HNTA*** *at a dose of 3 g/kg. Necrotic areas are found (white arrow). Magnification x200.*

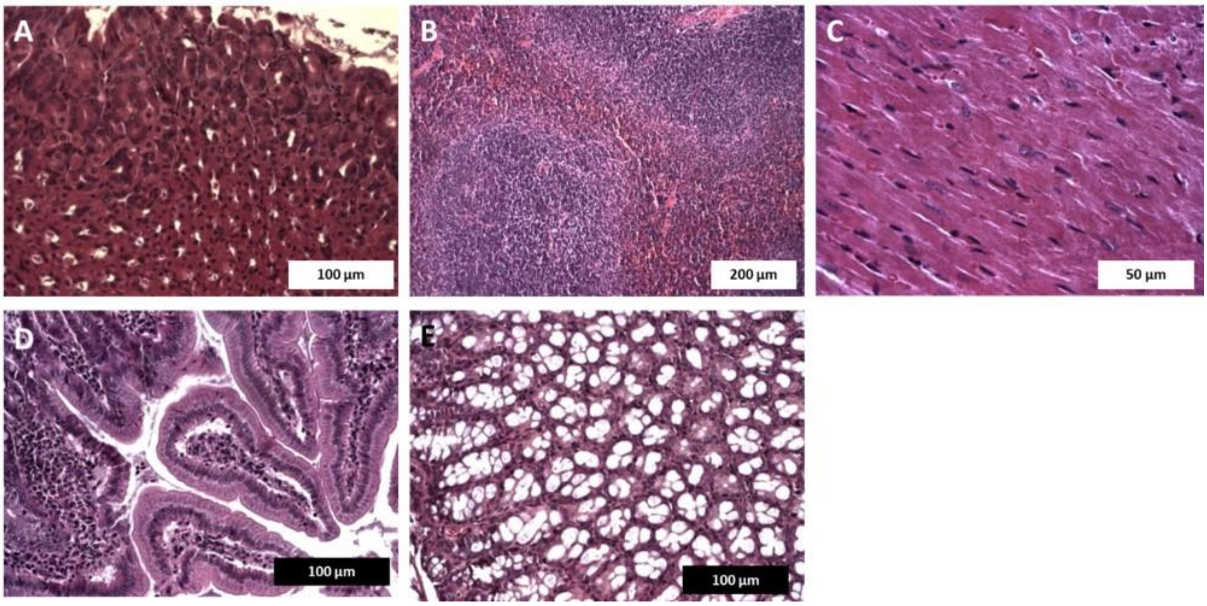


***Figure SIII.4.*** *Organs of mice after injection of* ***K-$Mo_2O_2S_2$-HNTA*** *at a dose of 3 g/kg that were not affected by the solution. A – stomach, B – spleen, C – heart, D – small intestine, E – large intestine. Magnification: stomach x200, spleen x100, heart x400, small intestine x200, large intestine x200.*

The liver of mice from the group that received 3 g/kg of **Na-$Mo_2O_4$-L-Cys** did not display any serious changes. However, the kidney had interstitial hyperemia and infiltration of protein in the tubular area (Figure SIII.5). The reason for this is most likely an increased vessel and epithelium permeability. Although this kidney disorder itself cannot be named the cause of death, deadly changes might have happened due to the increased permeability it reveals. Most likely, the brain area was affected by this change. The other organs of animals from this group were not affected by the solutions (Figure SIII.6)

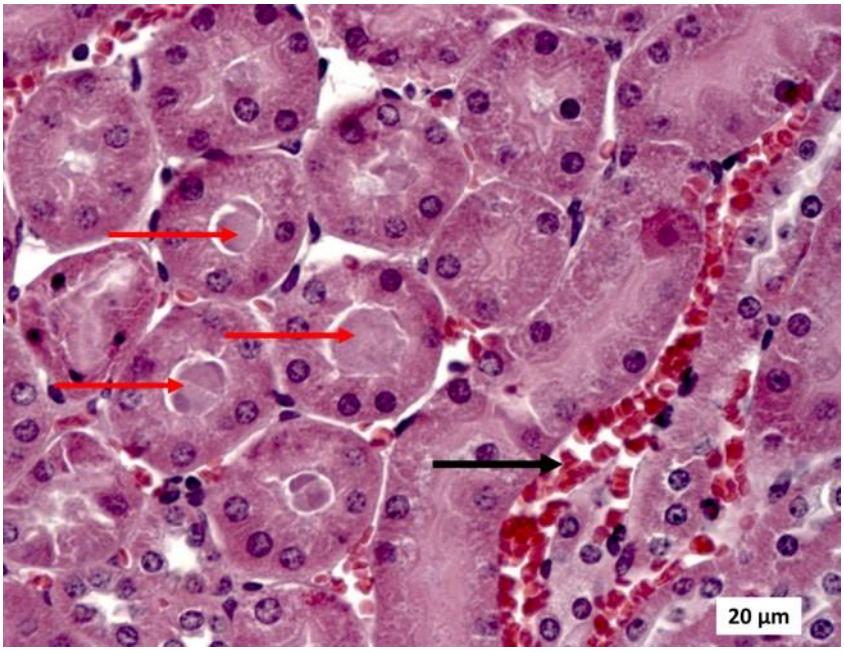


***Figure SIII.5.*** *Kidney of a mouse after receiving 3 g/kg of* ***Na-Mo$_2$O$_4$-L-Cys****. Interstitial hyperemia (black arrow) and protein infiltration of the tubular section (red arrows). Magnification X400.*

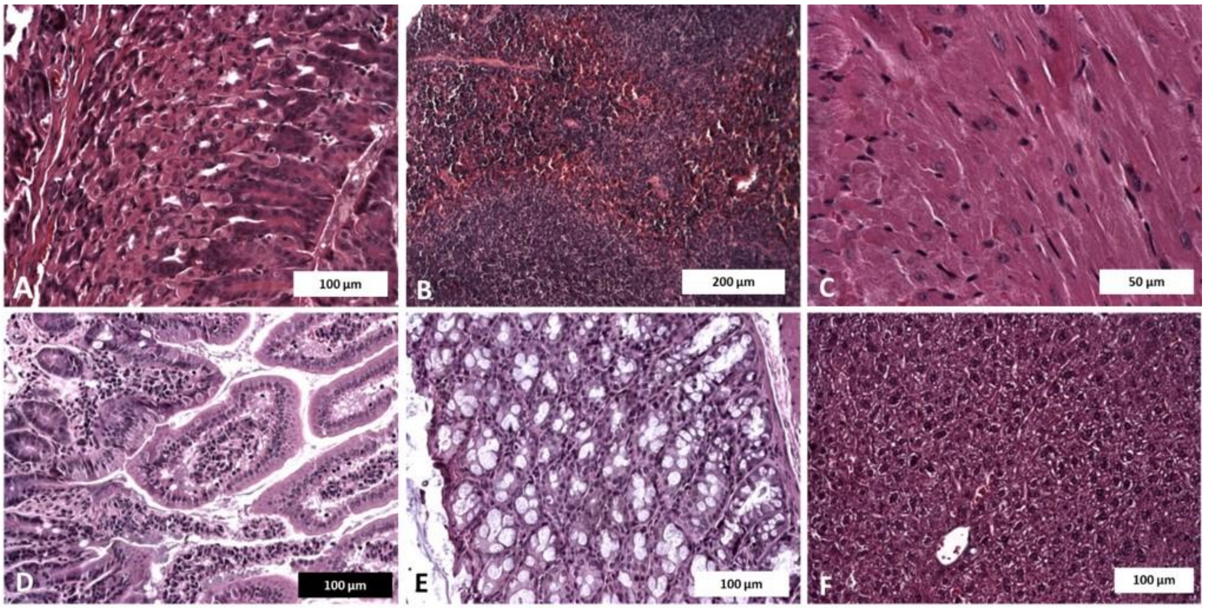


***Figure SIII.6.*** *Organs of mice that were not affected by the solution, after injection of* ***Na-Mo$_2$O$_4$-L-Cys*** *at a dose of 3 g/kg. A – stomach, B – spleen, C – heart, D – small intestine, E – large intestine, F - liver. Magnification: stomach x200, spleen x100, heart x400, small intestine x200, large intestine x200, liver x200.*

##### 4°) Analysis of molybdenum content in organs

In case of oral toxicology, the first question to be cleared is whether the substance can be resorbed at all and accumulated in organs. For the complexes and for the three groups of mice the Mo content was measured by ICP-AES. The results are given in Tables SIII.3 and SIII.4

***Table SIII.3.*** *Mo content in ppm (µg/g) measured in blood and in organs of mice that received 2.5 g/kg of complexes for 5 days (0.5 g per day). The limit of quantification (LOQ) in ppm is given into brackets*

| Complex | Blood | Spleen | Kidney | Heart | Lung | Stomach | Liver |
|---|---|---|---|---|---|---|---|
| control | <LOQ(0.05) | <LOQ(1) | <LOQ(0.5) | <LOQ(0.3) | <LOQ(0.3) | <LOQ(1) | 1.1 |
| ***K-$Mo_2O_2S_2$-L-Cys*** | <LOQ(0.05) | <LOQ(1) | <LOQ(0.5) | <LOQ(0.3) | <LOQ(0.3) | <LOQ(1) | 1.2 |
| ***K-$Mo_2O_2S_2$-HNTA*** | <LOQ(0.05) | <LOQ(1) | <LOQ(0.5) | <LOQ(0.3) | <LOQ(0.3) | <LOQ(1) | 1.2 |
| ***Na-$Mo_2O_4$-L-Cys*** | <LOQ(0.05) | <LOQ(1) | <LOQ(0.5) | <LOQ(0.3) | <LOQ(0.3) | <LOQ(1) | 1 |
| ***Na-$Mo_2O_4$-EDTA*** | <LOQ(0.05) | <LOQ(1) | <LOQ(0.5) | <LOQ(0.3) | <LOQ(0.3) | <LOQ(1) | 0.95 |
| ***K-$Mo_2O_2S_2$-EDTA*** | <LOQ(0.05) | <LOQ(1) | <LOQ(0.5) | <LOQ(0.3) | <LOQ(0.3) | <LOQ(1) | 1.7 |

***Table SIII.4.*** *Mo content in ppm (µg/g) measured in blood and in organs of mice that received 3.0 g/kg of complexes in one day. The limit of quantification (LOQ) is given into brackets*

| Complex | Blood | Spleen | Kidney | Heart | Lung | Stomach | Liver | Small Intestine | Duodenum |
|---|---|---|---|---|---|---|---|---|---|
| ***K-$Mo_2O_2S_2$-L-Cys*** | 0.64(0.05) | <LOQ(1) | 0.99(0.5) | <LOQ(0.3) | 2.0(0.3) | <LOQ(1) | 1.5(0.3) | 0.7(0.3) | 2.3(0.3) |
| ***K-$Mo_2O_2S_2$-HNTA*** | 0.24(0.05) | <LOQ(1) | 1(0.5) | <LOQ(0.3) | <LOQ(0.3) | <LOQ(1) | 1.5(0.3) | 0.2(0.3) | 0.8(0.3) |
| ***Na-$Mo_2O_4$-L-Cys*** | <LOQ(0.05) | 1.5(1) | 1.1(0.5) | 1.3(0.3) | 1.3(0.3) | 1.6(1) | 1.9(0.3) | 1.7(0.3) | 2.3(0.3) |
| ***Na-$Mo_2O_4$-EDTA*** | 0.44(0.05) | <LOQ(1) | 0.83(0.5) | <LOQ(0.3) | <LOQ(0.3) | <LOQ(1) | 1.6(0.3) | 1.4(0.3) | 0.7(0.3) |
| ***K-$Mo_2O_2S_2$-EDTA*** | 0.31(0.05) | <LOQ(1) | 0.78(0.5) | <LOQ(0.3) | 0.38(0.3) | <LOQ(1) | 1.4(0.3) | 0.6(0.3) | 1.0(0.3) |

A molybdenum content analysis in the different viscera of mice treated with a dose of 2.5 g/kg over 5 days did not reveal Mo in the studied organs. In the second group, which received 3 g/kg in one day, Mo was mainly accumulated in the blood, kidneys, liver, small intestine and large intestine. Molybdenum was also detected in the lungs of mice that received **K-$Mo_2O_2S_2$-L-Cys, Na-$Mo_2O_4$-L-Cys** and **K-$Mo_2O_2S_2$-EDTA**. The content of Mo for these three compounds in the lungs is associated with their uptake by alveolar macrophages. It should be noted that Mo was detected in the stomach after injection of **Na-$Mo_2O_4$-L-Cys** only. Thus, it is shown that the studied compounds are absorbed and end up in the organs. Since all compounds are found in the kidneys and liver, it is possible to propose that reabsorbed substances further excreted by kidneys and liver *in vivo*.

##### 5°) Morphological analysis for compound Li-$Mo_2O_4$-EDTA

The compound **Li-$Mo_2O_4$-EDTA** received a particular attention, notably due to possible effect both of the Molybdenum-based complexes but also of lithium counter cations. Morphological analysis of mice organs was performed in Group 1 (which received 0.5 g/kg once a day for 5 consecutive days) and Group 2 (3 g/kg in one day).

**Heart tissue**

There are no significant changes in the morphology of heart tissue. Blood filling of the myocardium remains within normal limits. The intermuscular stroma and cardiomyocytes does not show any pathologies. No foci of ischemic or dystrophic damage were found.

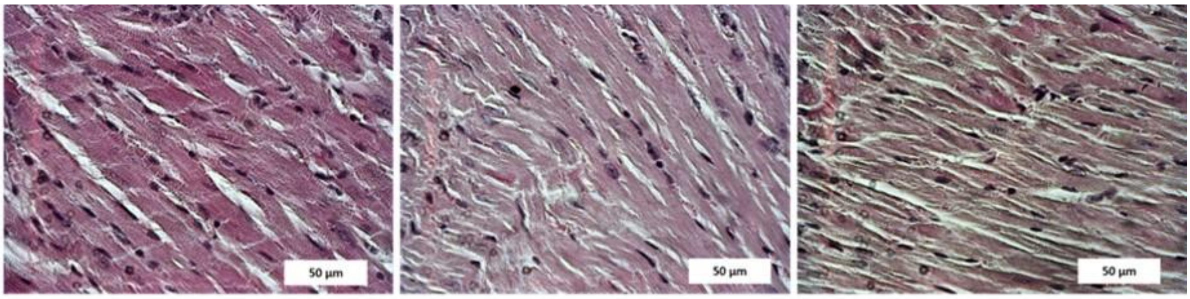


Control Group 1 Group 2

***Figure SIII.7.*** *Heart morphology of mice that received solutions of* ***Li-$Mo_2O_4$-EDTA****. Group 1 at a dosage of 2.5 g/kg (0.5 g/kg once a day for 5 consecutive days) and Group 2 at a dosage of 3 g/kg (administered three times at a dose of 1 g/kg with a break of 2 hours). Images were obtained at x400 magnification.*

**Stomach tissue**

The ratio of the layers of gastric mucosa, submucosa and muscularis externa is the same in the control and experimental groups. The condition of the gastric pit epithelial cells and gastric gland cells is not changed. No pathological changes were found in the stomach tissue.

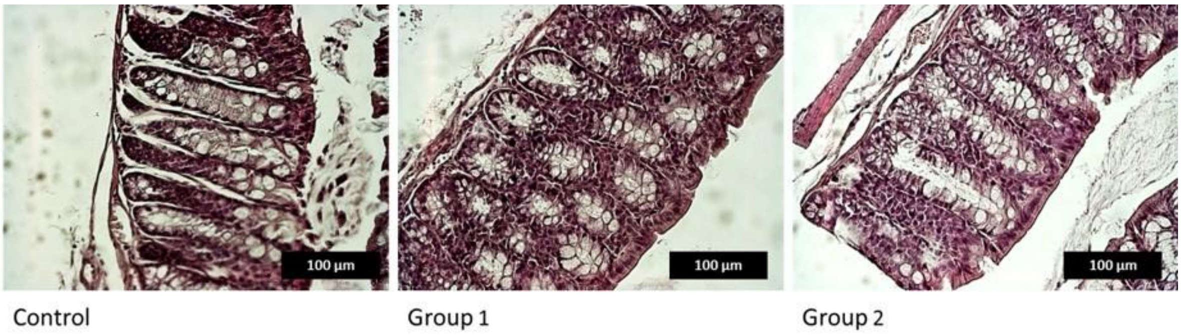


***Figure SIII.8.*** *Stomach morphology of mice that received solutions of **Li-$Mo_2O_4$-EDTA**. Group 1 at a dosage of 2.5 g/kg (0.5 g/kg once a day for 5 consecutive days) and Group 2 at a dosage of 3 g/kg (in one day). Images were obtained at x200 magnification.*

**Large intestine**

The condition of the colon wall is normal. Blood supply and functional state of the tubular glands is without any pathological changes.

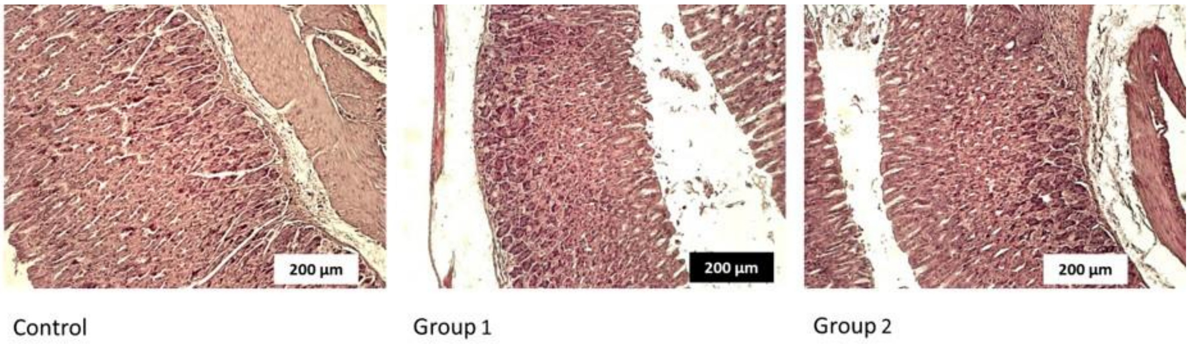


***Figure SIII.9.*** *Large intestine morphology of mice that received solutions of **Li-$Mo_2O_4$-EDTA**. Group 1 at a dosage of 2.5 g/kg (0.5 g/kg once a day for 5 consecutive days) and Group 2 at a dosage of 3 g/kg (administered in one day, in three times at a dose of 1 g/kg with a break of 2 hours). Images were obtained at x100 magnification.*

**Spleen**

The ratio of white and red pulp does not change after the injection of **Li-$Mo_2O_4$-EDTA**. The splenic nodules retain their shape. Sheathed capillaries do not show any increased hemolysis or macrophage hyperactivity.

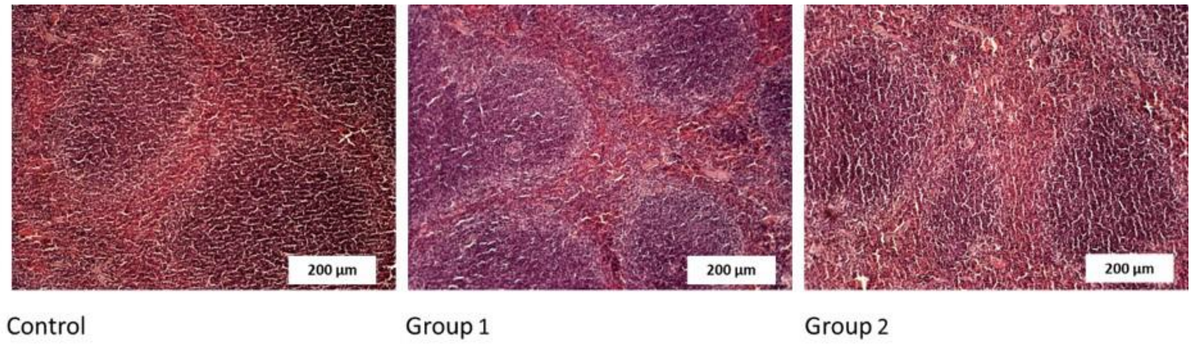


***Figure SIII.10***. *Spleen morphology of mice that received solutions of **Li-$Mo_2O_4$-EDTA**. Group 1 at a dosage of 2.5 g/kg (0.5 g/kg once a day for 5 consecutive days) and Group 2 at a dosage of 3 g/kg (administered in one day, three times at a dose of 1 g/kg with a break of 2 hours). Images were obtained at x100 magnification.*

**Kidney tissue**

No pathological changes were found in the kidneys of mice after administration of **Li-$Mo_2O_4$-EDTA**. Apparently, the substance either passes completely through the renal filter and does not linger in the kidneys, or has particles too large for renal filtration and that do not pass through the glomerulus of the nephron. No pathological changes were found in either the medulla or the cortex of the kidney.

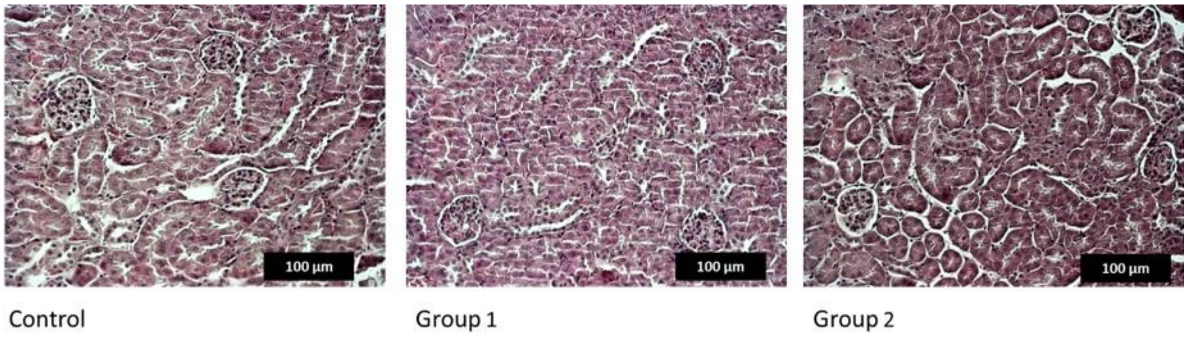


***Figure SIII.11.*** *Kidney medulla morphology of mice that received solutions of **Li-$Mo_2O_4$-EDTA.** Group 1 at a dosage of 2.5 g/kg (0.5 g/kg once a day for 5 consecutive days) and Group 2 at a dosage of 3 g/kg (administered in one day, three times at a dose of 1 g/kg with a break of 2 hours). Images were obtained at x200 magnification.*

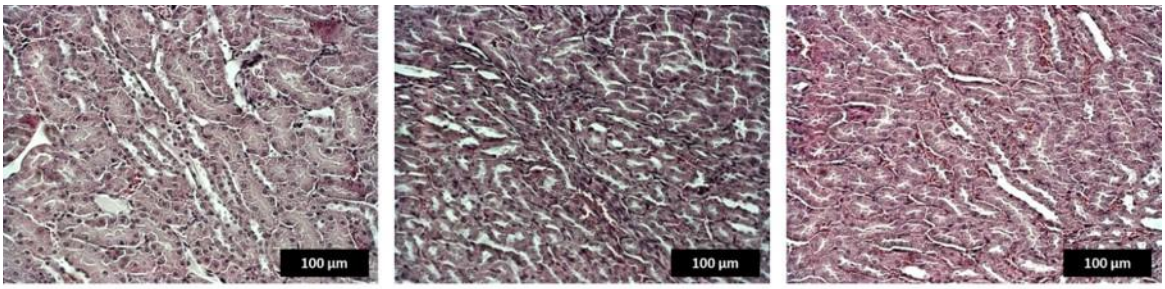


Control Group 1 Group 2

***Figure SIII.12.*** *Kidney cortex morphology of mice that received solutions of* ***Li-$Mo_2O_4$-EDTA****. Group 1 at a dosage of 2.5 g/kg (0.5 g/kg once a day for 5 consecutive days) and Group 2 at a dosage of 3 g/kg (administered in one day, three times at a dose of 1 g/kg with a break of 2 hours). Images were obtained at x200 magnification.*

**Liver tissue**

Dystrophic changes of hepatocytes are manifested in the liver tissue. It is worth noting that pathological changes are more pronounced in the area of portal channels. This suggests that the toxic effect on hepatocytes is caused by blood coming from the intestinal vessels and delivered to the liver through the portal vein.

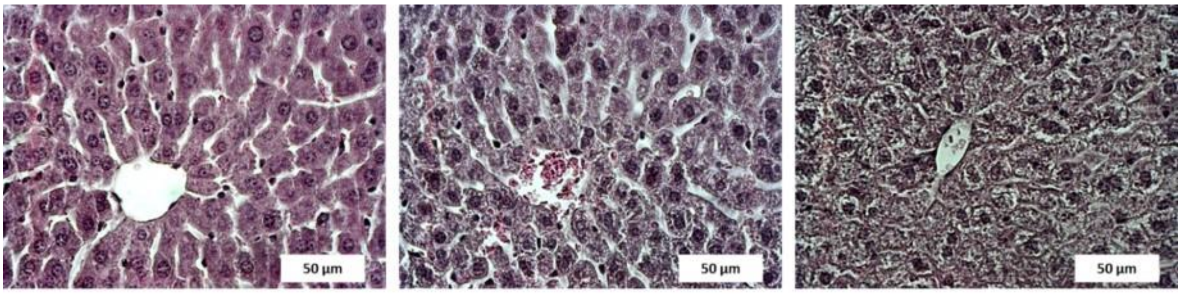


Control Group 1 Group 2

***Figure SIII.13.*** *Liver morphology of mice that received solutions of* ***Li-$Mo_2O_4$-EDTA****. Group 1 at a dosage of 2.5 g / kg (0.5 g/kg once a day for 5 consecutive days) and Group 2 at a dosage of 3 g/kg (administered in one day, three times at a dose of 1 g/kg with a break of 2 hours). Central vein area. Images were obtained at x400 magnification.*

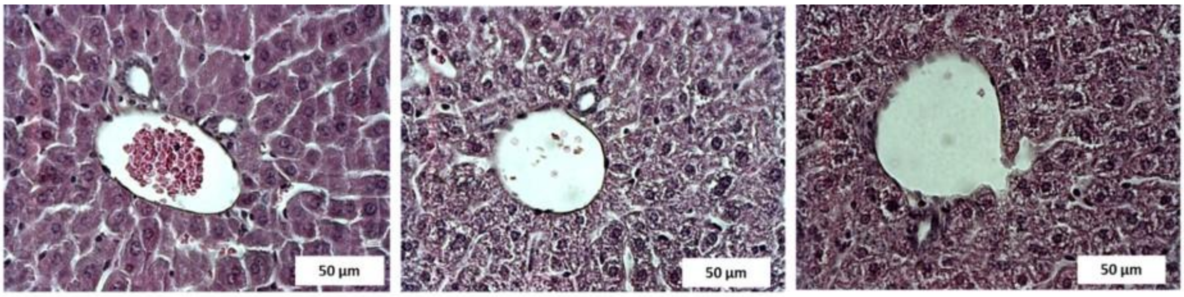


Control Group 1 Group 2

***Figure SIII.14.*** *Liver morphology of mice that received solutions of* ***Li-$Mo_2O_4$-EDTA****. Group 1 at a dosage of 2.5 g / kg (0.5 g/kg once a day for 5 consecutive days) and Group 2 at a dosage of 3 g/kg (administered in one day, three times at a dose of 1 g/kg with a break of 2 hours). Portal channels area. Images were obtained at x400 magnification.*

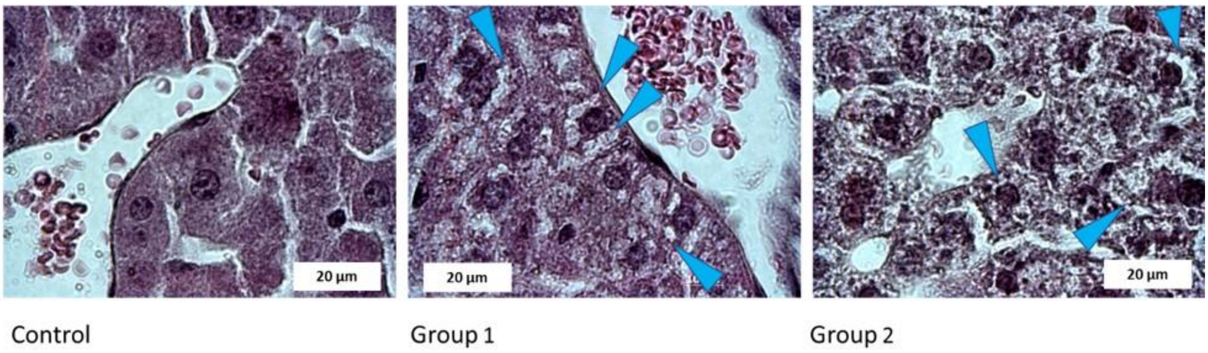


***Figure SIII.15.*** *Hepatocytes of a mouse that received solutions of* ***Li-$Mo_2O_4$-EDTA*** *Group 1 at a dosage of 2.5 g / kg (0.5 g/kg once a day for 5 consecutive days) and Group 2 at a dosage of 3 g/kg (administered in one day, three times at a dose of 1 g/kg with a break of 2 hours). Dystrophic vacuoles in cells are shown by blue arrows. Images were obtained at x1000 magnification.*

##### III.1.3 Conclusions

The determination of the acute toxicity of the compounds has shown that the most toxic compounds for mice are **K-$Mo_2O_2S_2$-HNTA** and **Na-$Mo_2O_4$-L-Cys**. $LD_{50}$ of these compounds is 3 g/kg by oral administration. $LD_{50}$ for other studied substances could not be determined due to their low toxicity by oral administration of the maximum available concentrations of synthesized compounds.

The target organs are kidneys, liver, small intestine and large intestine. In the liver and kidneys necrosis develops, in the renal tubules the accumulation of protein content was observed after the injection of the compound **Na-$Mo_2O_4$-L-Cys** at a dose of 3 g/kg. The accumulation of Molybdenum in the lungs after oral administration was noticed with **K-$Mo_2O_2S_2$-L-Cys**, **Na-$Mo_2O_4$-L-Cys** and **K-$Mo_2O_2S_2$-EDTA**. It should also be noted that the introduction of synthesized compounds within 5 days of a dose of 0.5 g/kg did not lead to their accumulation in the organs as well as to any pathological changes.

Finally, the administration of substances **Na-$Mo_2O_4$-EDTA** and **Li-$Mo_2O_4$-EDTA** did not have any significant pathological effect on most of the studied mouse organs. The only significant change was found in liver tissue when **Li-$Mo_2O_4$-EDTA** was used. The introduction of the **Li-$Mo_2O_4$-EDTA** causes hepatocyte dystrophy, which is more pronounced in the area of portal triads. This fact confirms that pathological changes are caused by the introduction of the **Li-$Mo_2O_4$-EDTA**, since blood coming from the digestive system is then delivered directly to the portal veins.

The next toxicity studies will thus be focused on these two compounds.

### III.2- Acute Toxicity on the water flea *Daphnia magna*

The molybdenum complexes explored in this study are being considered as potential supplements in bee feed. Assessing their toxicity across various living organisms, notably bees, also requires evaluating the potential environmental impact in case of release into water resources. To this end, we complemented our research with toxicity tests on aquatic organisms such as *Daphnia magna* water fleas.

Conducting toxicity tests on aquatic organisms like *Daphnia magna* before studying the effects on bees serves several purposes:

**Preliminary Safety Assessment:** Daphnia are widely used as model organisms for assessing toxicity in aquatic environments. Their reactions to different substances provide initial data on how these compounds might affect organisms in water.

**Ecological Significance:** Aquatic organisms, including Daphnia, serve as indicators of water ecosystem health. Studying their responses provides insight into the ecological safety of the substance.

**Determining Safe Dosages:** Understanding toxicity on Daphnia helps establish safe substance concentrations for bees and other organisms.

**Time and Resource Efficiency:** Preliminary testing on simpler organisms like Daphnia reduces the risk of exposing more sensitive organisms like bees in the early stages of research.

Hence, toxicity testing on aquatic organisms, such as *Daphnia magna*, represents an important stage in assessing the safety of chemical substances before conducting more intricate experiments involving higher organisms like bees.

#### Experimental procedure

The toxicity assessment of the tested compounds was conducted using *Daphnia magna* (Straus, 1820). The *Daphnia magna* used in this study were obtained from a parthenogenetic culture [3-5].

*Daphnia magna* were nourished with Chlorella vulgaris, an unicellular algae cultivated using aseptic techniques to prevent contamination by bacteria, algae, or protozoa. Chlorella vulgaris was grown in Prat's growth medium, which consisted of $KNO_3$ (1 µM), $MgSO_4{\cdot}7H_2O$ (40 µM), $K_2HPO_4{\cdot}3H_2O$ (400 µM), and $FeCl_3{\cdot}6H_2O$ (3.6 µM) in distilled water (pH adjusted to 7.0, autoclaved, and stored at 5°C).

*D. magna* were maintained in aerated aqueous straw infusion growth media supplemented with $CaCl_2$ (11.76 g/L), $NaHCO_3$ (2.59 g/L), KCl (0.23 g/L), and $MgSO_4 \cdot 7H_2O$ (4.93 g/L) to maintain a pH of approximately 7.5 ± 0.2 and ensure dissolved oxygen levels of ≥6.0 mg/L.

Juveniles were selected based on size and acclimated to fresh medium for 24 h. The *D. magna* were cultured in Costar® 24-well clear sterile multiple well plates, covered with lids to prevent contamination and evaporation while allowing gaseous exchange. Each well contained 10 daphnids in 1000 µL of each dilution of the tested compounds.

The bioassay was conducted with concentrations ranging from 0.1 to 100 µM (0.1, 1, 10, and 100 µM) to determine the $LC_{50}$ for each compound. Stock solutions were diluted with aqueous straw infusion growth media to achieve the required concentrations. The final test solutions contained up to 0.1% DMSO and had a final volume of 1 mL. A 0.1% DMSO solution in aerated medium (pH~7.5 ± 0.2; O2 ≥ 6.0 mg/L) served as the negative control.

Throughout the experiment, juvenile daphnids were incubated at 22 ± 2°C under a 16 h light/dark cycle (500–1000 lx). Mobility (viability) of the test organisms was assessed after the 24-hour exposure. The experiment was conducted in triplicate.

Daphnids were considered immobilized if they did not swim during the 15-second period following gentle agitation of the test and control solutions, even if they could still move their antennae. The percentage of viability (V, %) of *Daphnia magna* was calculated using the formula (2):

$$\mathbf{V}(\%) = \frac{\mathbf{N}_{(\mathbf{sample})}}{\mathbf{N}_{(\mathbf{control})}} \times \mathbf{100}$$

Where N represents the number of viable *Daphnia magna*. $LC_{50}$ values, which represent the median lethal concentration that kills 50% of the juvenile daphnids, were determined using the least squares fit method based on the dose-response equation.

#### Results

The toxicity study on *Daphnia magna* involved testing the biologically active compounds **Na-$Mo_2O_4$-EDTA** and **Li-$Mo_2O_4$-EDTA**, along with sodium molybdate dihydrate as a reference standard. A microscopic analysis was conducted on the control group of *D. magna* that was not exposed to any chemical compounds, revealing no pathological changes or alterations in the organisms.

The toxicity study results are presented in Table SIII.5, providing information on the impact of these compounds on *D. magna*.

***Table SIII.5***. *$LC_{50}$ values of Molybdenum compounds on Daphnia magna.*

| **Compounds** | **$LC_{50}$ ( μM)** | |
|---|---|---|
| | ***Incubation period* (24h)** | ***Incubation period* (48h)** |
| **Control** | ≥100 | ≥100 |
| **$Na_2MoO_4 \cdot 2H_2O$** | ≥100 | ≥100 |
| **Na-$Mo_2O_4$-EDTA** | ≥100 | ≥100 |
| **Li-$Mo_2O_4$-EDTA** | ≥100 | ≥100 |

The results presented in Table SIII.5 indicate that the molybdenum complexes **Na-$Mo_2O_4$-EDTA** and **Li-$Mo_2O_4$-EDTA** showed no signs of toxicity on *Daphnia magna* during the 24-hours and 48-hours incubation periods, with an $LC_{50}$≥100 μM. This absence of toxicity observed in these complexes aligns with the response observed in the control group treated with sodium molybdate.

Throughout the duration of the experiment, there were no observable adverse effects or indications of harm to the *Daphnia magna* specimens exposed to either **Na-$Mo_2O_4$-EDTA** or **Li-$Mo_2O_4$-EDTA**. The absence of toxicity at both time intervals suggests that these particular molybdenum complexes do not elicit harmful effects on *Daphnia magna* within the tested exposure periods.

**Conclusion:**

The toxicity assessment of molybdenum complexes **Na-$Mo_2O_4$-EDTA** and **Li-$Mo_2O_4$-EDTA** on the aquatic organism *Daphnia magna* revealed no signs of toxicity during the 24 and 48-hours exposure periods. These findings constitute crucial evidence confirming the absence of adverse effects of these compounds on *Daphnia magna* at the tested concentrations and timeframes.

The experiment demonstrated that the proposed molybdenum complexes did not exhibit any detrimental effects on the water fleas, *Daphnia magna*, which serve as important indicators of water ecosystem health. The absence of toxicity across different time intervals underscores the safety of these compounds for this particular aquatic organism.

These results provide valuable insights for further research and assessment of the potential use of these molybdenum complexes as supplements in bee feed. Additionally, they instill confidence in the safety of these compounds concerning their potential release into aquatic environments.

#### III.3- Evaluation of the effect of **Na-$Mo_2O_4$-EDTA** and **Li-$Mo_2O_4$-EDTA** on bees: Acute oral and contact test on honeybees (*Apis mellifera*)

#### III.3.1 Experimental procedures

The study was conducted by Testapi S.A.S., Allonnes (49), France, in accordance with OECD guidelines Nos. 213 and 214 concerning oral and contact test methods to assess the acute toxicity of plant protection products on honey bees (OECD guidelines for the testing of chemicals, "Honey bees, Acute Oral and Contact Toxicity test"; OECD 213 and 214, September 21 1998).

**1°) Reference substance / control**

Dimethoate Pestanal™ 99.8% purity provided by Sigma-Aldrich is known to be an active substance with a strong effects on honey bees.[6] It was used as toxic reference for the validation of the tests. Demineralized water mixed with sucrose solution was used as control treatment in oral tests. Demineralized water was used as control in contact tests.

**2°) Selection and preparation of honey bees for the tests**

The tests were conducted on the species *Apis mellifera L.*, on adult worker honey bees, directly collected on a hive frame and after being quickly anaesthetized with carbon dioxide. Bees were visually inspected and found to be in good health and free from disease. 10 to 12 honey bees working adults were selected and placed in holding boxes (0.6 $dm^3$, see Figure SIII.17).

Before applications, boxes were placed in incubators at a temperature of 25°±2°C, with a relative humidity between 50% and 70%. During this time, a concentrated sucrose feeding solution (500 g/L) was available *ad libitum* to bees.

**3°) Description of the number of treatments and replicates**

As the test complexes **Na-$Mo_2O_4$-EDTA** and **Li-$Mo_2O_4$-EDTA** were supposed to have low toxicity for honey bees, the oral and contact phases were directly performed as limit tests (Tables SIII.6 and SIII.7). The tests were carried out with a single high dose (expected > 100 µg/bee) and 6 replicates (3 for the toxic reference).

Considering a feeding of 2 to 8 mg of complexes per hive at the really beginning of spring (see part 5, tests in beehives) and considering that at this period the population of a hive is approximately 10,000 bees, if each bee receives a similar dose, an average dose of 0.2 to 0.8 µg of complex per bee is obtained. This value decreases if the population is larger. To evaluate the toxicity of **Na-$Mo_2O_4$-EDTA** and **Li-$Mo_2O_4$-EDTA** on bees through contact and oral absorption, the experiments were performed with 110 µg of complex per bee, which is 137 to 550 times the average value they could receive within a hive (see Tables SIII.6 and SIII.7). Untreated controls provided an assessment of « natural » mortality and informed about the quality of the selected bees.

***Table SIII.6***. *Description for oral dose-response test.*

| Modality | Product / active substance | Number of bees | Amount of diet distributed (µL/bee) | Number of replicates | Dose of active substance/bee |
|---|---|---|---|---|---|
| 1 | Water | 11 / box | 10 | 6 | 0 µg |
| 2 | Dimethoate | 11 / box | 10 | 3 | 0.10 µg |
| | | | 10 | | 0.20 µg |
| | | | 10 | | 0.35 µg |
| 3 | **Na-$Mo_2O_4$-EDTA** | 11 / box | 10 | 6 | 110 µg |
| 4 | **Li-$Mo_2O_4$-EDTA** | 11 / box | 10 | 6 | 110 µg |

***Table SIII.7.*** *Description for contact dose-response test.*

| Modality | Product / active substance | Number of bees | Amount solution distributed (µL/bee) | Number of replicates | Dose of active substance/bee |
|---|---|---|---|---|---|
| 1 | Water | 10 / box | 1 | 6 | 0 µg |
| 2 | Dimethoate | 10 / box | 1 | 3 | 0.10 µg |
| | | | 1 | | 0.20 µg |
| | | | 1 | | 0.35 µg |
| 3 | **Na-$Mo_2O_4$-EDTA** | 10 / box | 1 | 6 | 110 µg |
| 4 | **Li-$Mo_2O_4$-EDTA** | 10 / box | 1 | 6 | 110 µg |

**4°) Preparation of solutions**

**Solutions of Na-$Mo_2O_4$-EDTA and Li-$Mo_2O_4$-EDTA**

Oral limit test: the high concentration oral solutions of **Na-$Mo_2O_4$-EDTA** or **Li-$Mo_2O_4$-EDTA** were prepared by sampling 22 mg of complexes diluted in 2 mL syrup (sucrose 50%) to obtain a 2 mL solution at 11 µg of complex per µL. 10 µL were given to the bees (Table SIII.6), which corresponds to 110 µg/bee.

Contact limit test: the high concentration solutions of **Na-$Mo_2O_4$-EDTA** or **Li-$Mo_2O_4$-EDTA** were prepared by sampling 220 mg of complex diluted in 2 mL syrup (sucrose 500 g/L) to obtain a 2 mL solution equivalent to 110 µg /µL. 1 µL was applied on the bees (Table SIII.7) which corresponds to 110 µg/bee.

**Dimethoate** (toxic reference) was prepared from an analytical standard named Dimethoate Pestanal™. This product was dispersed in demineralized water only or finally in sucrose solution (depending on the test) in order to obtain concentrations described as "low" "medium" and "high" as recommended in the guideline.

For the oral test, several dilutions in syrup were performed to obtain 0.01; 0.02; 0.035 µg dimethoate/µL in order to obtain final doses of 0.1, 0.2 and 0.35 µg dimethoate/bee as feeding solutions which are distributed at 10 µL per bee (110 µL for a box containing 11 bees).
For the contact tests, several dilutions in demineralized water were performed to obtain 0.1; 0.20; 0.3 µg dimethoate/µL in order to reach final doses of 0.1 µg/bee; 0.20 µg/bee and 0.3 µg/bee as contact solutions which are distributed at 1 µL per bee.

##### 5°) Exposure conditions

###### Equipment

Tools and specific equipment used in this study present characteristics in conformity with the requirements of referenced methodologies (OECD guidelines for the testing of chemicals, "Honey bees, Acute Oral and Contact Toxicity test"; OECD 213 and 214, September 21 1998). Holding boxes were plastic made, and their volume was 0.6 $dm^3$ (see Figure SIII.17) In order to avoid possible contamination within boxes as well as feeders (syringes), all materials were removed after use. Equipment use was managed under specific Standard Operating Procedures (SOP). Test chambers were maintained in darkness, except for bee operated procedures, which were conducted in subdued light. Temperature was maintained from 23 to 27°C and relative humidity ranged between 49% and 89%.

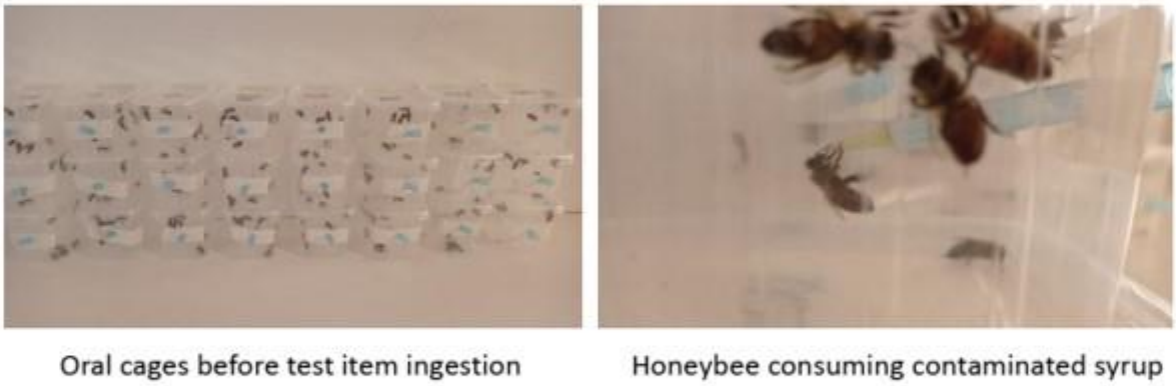


*Figure SIII.17 : Holding Boxes used for this study.*

###### Mode of application

The applications were performed in a short period of time. Treatments begin with the control units, then the units with complexes and finally the toxic reference item in the respective boxes.

###### Oral test

On the application day, feeders (syringes) were removed from each holding box and bees starve for one to two hours. Then bees receive the sucrose solution containing products at the different concentrations. This contaminated syrup was distributed at a mean of 10 µL per bee (110 µL per feeder per box of 11 bees). The amount of contaminated diet consumed was monitored up to 6 hours after administration. As soon as the contaminated syrup has been eaten, the syringe was filled with non-treated sucrose solution. If some syrup was remaining after 6 hours, the amount was measured. Bees were fed *ad libitum* with sucrose solution syrup until the last observation.

**Contact test**

In order to apply substances, bees were put through a light anaesthesia with carbon dioxide. The small volume of the holding boxes allowed doing this anaesthesia with a very low gas flow and leads bees under lethargy for few minutes.

Substances (water in the control and solutions of Mo-based complexes or toxic reference modality) were manually applied in only once. A micro syringe allowed laying out 1 µL of solution on the bee thorax (see Figure SIII.18)

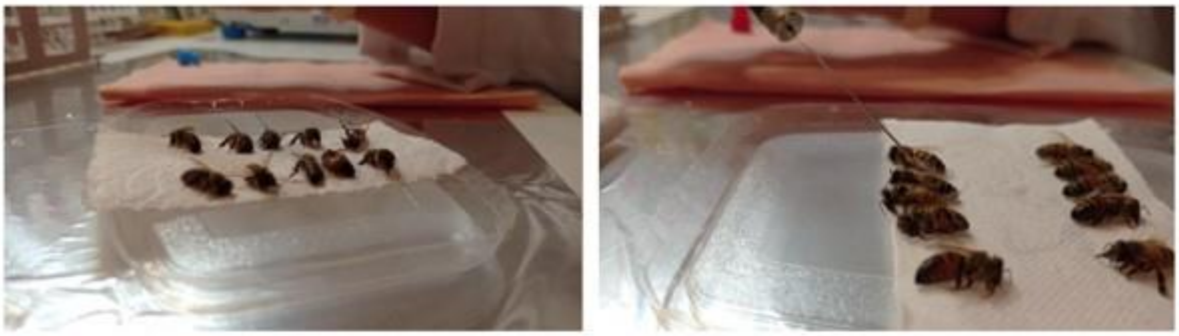

Contact anesthetised honeybees before application of solution (1µL)

Application of 1µL of solution

*Figure SIII.18 : Treatment of bees with 1 µL of solution on the thorax of sleeping bees with $CO_2$.*

**6°) Observations**

Following substance applications, bees were kept in temperature and relative humidity-controlled conditions. Main assessments consisted in counting dead bees. In each holding box, dead bees were counted at T+4 hours, T+24 hours, and T+48 hours at least. If the test item mortality increases for more than 10% from T+24h to T+48h, further assessments were recorded every 24h until T+96h, as long as the control percentage mortality remains below 10% and the test item mortality increased. The mortality is expressed in percentage of the initial population before application of modalities.

**7°) Data analysis**

Data analysis were performed on Excel software. In order to correct the mortality recorded in the test substance treatments by the mortality of the control group, the following Abbott formula was used [7].

$$M = \frac{(\%P - \%T) \times 100}{100 - \%T}$$

By using mortality percentages referring to the trial methodology: **M** was the corrected mortality expressed in percent of initial population; **% P** was the percentage of mortality induced by the product; **% T** was the percentage of mortality in the controls. Each test and control treatment were represented by a minimum of three boxes.

Data on mortality were analyzed with appropriate statistical methods. A $X^2$ test was performed on raw mortality data to highlight any difference between the tested modalities (**Na-$Mo_2O_4$-EDTA** or **Li-$Mo_2O_4$-EDTA**) and the water control [8].

#### III.3.2 Results

##### Validity criteria

The percentages of mortality recorded in the control and the toxic reference treatments validated the laboratory design with data from OECD guidelines :

- Mean mortality in the control was less than 10% of the initial population:
  o Oral dose-response test, 3 % at T+24h, 5 % at T+48h
  o Contact dose-response test, 3 % at T+24h, 7 % at T+48h
- The mortality in the toxic reference at T+24h matched the range of LD50, 0.1-0.35 µg dimethoate per bee:
  o Oral dose-response test, 24h-$LD_{50}$= 0.192 µg per bee
  o Contact dose-response test, 24h-$LD_{50}$= 0.159 µg per bee.

##### Results for Oral tests

The doses chosen were 110 µg **Na-$Mo_2O_4$-EDTA** or **Li-$Mo_2O_4$-EDTA** per bee. Bees were fed with 10 µL of syrup each. Results are given in Table SIII.8.

***Table SIII.8.*** *Mean mortality rates in the oral tests.*

| | | | T+24H | | T+48H | |
|---|---|---|---|---|---|---|
| Treatment | Dose consumed (µg/bee)[a] | Number of replicats | % of mortality | % of corrected mortality[b] | % of mortality | % of corrected mortality[b] |
| Control | - | 6 | 3 | - | 5 | - |
| **Na-$Mo_2O_4$-EDTA** | 110 | 6 | 2 | 0 | 8 | 3 |
| **Li-$Mo_2O_4$-EDTA** | 110 | 6 | 2 | 0 | 9 | 5 |
| Dimethoate | 0.1 | 3 | 27 | 25 | 76 | 75 |
| | 0.2 | 3 | 60 | 58 | 100 | 100 |
| | 0.35 | 3 | 85 | 84 | 100 | 100 |

a. Comsumed dose expressd after 6 hours' exposure
b. Expressed as a corrected value according to the Abott formula

The $X^2$ test show no differences between mortality in test items modalities (either **Na-$Mo_2O_4$-EDTA** or **Li-$Mo_2O_4$-EDTA**) and the control modality, p=0.47 for MoNa and p=0.30 for MoLi.

From this phase, conclusion about the **oral 48h-$LD_{50}$ values were:**

- >**110 µg Na-$Mo_2O_4$-EDTA per bee**.
- >**110 µg Li-$Mo_2O_4$-EDTA per bee**

#### Results for contact tests

The doses chosen were: 1 µL per bee (110 µg **Na-$Mo_2O_4$-EDTA** or **Li-$Mo_2O_4$-EDTA** per bee). One µL solution was applied per bee thorax.

***Table SIII.9.*** *Mean mortality rates in the contact tests.*

| | | | T+24H | | T+48H | | T+72H | |
|---|---|---|---|---|---|---|---|---|
| Treatment | Dose consumed (µg/bee) | Number of replicats | % of mortality | % of corrected mortality[a] | % of mortality | % of corrected mortality[a] | % of mortality | % of corrected mortality[a] |
| Control | - | 6 | 3 | - | 7 | - | 15 | - |
| **Na-$Mo_2O_4$-EDTA** | 110 | 6 | 2 | 0 | 3 | 0 | 15 | 0 |
| **Li-$Mo_2O_4$-EDTA** | 110 | 6 | 0 | 0 | 2 | 0 | 7 | 0 |
| Dimethoate | 0.1 | 3 | 40 | 38 | 53 | 50 | 53 | 45 |
| | 0.2 | 3 | 53 | 52 | 97 | 96 | 97 | 96 |
| | 0.35 | 3 | 97 | 97 | 90 | 89 | 97 | 96 |

a. Expressed as a corrected value according to the Abott formula

At T+48h, the $X^2$ test show no differences between mortality in test items modalities (either **Na-$Mo_2O_4$-EDTA** or **Li-$Mo_2O_4$-EDTA**) and the control modality, p=0.40 for **Na-$Mo_2O_4$-EDTA** and p=0.17 for **Li-$Mo_2O_4$-EDTA.**

Conclusion about the **oral 48h-$LD_{50}$ values were:**

- >**110 µg Na-$Mo_2O_4$-EDTA per bee**.
- **>110 µg Li-$Mo_2O_4$-EDTA per bee.**

### III.4- Chronic oral toxicity on bees : mortality studies

The objective of this study was to observed if a chronical treatment with Molybdenum complexes **Li-$Mo_2O_4$-EDTA** and **Na-$Mo_2O_4$-EDTA** solubilized in a sugar syrup could alter bees survival.

#### III.4.1 Experimental procedure

The study was conducted by the EGCE team at CNRS, Gif-sur-Yvette (91), France. The experimental plan was inspired by OECD directive No 245 (OECD (2017), *Test No. 245: Honey Bee (Apis Mellifera L.), Chronic Oral Toxicity Test (10-Day Feeding)*, OECD Guidelines for the Testing of Chemicals, Section 2, Éditions OCDE, Paris, https://doi.org/10.1787/9789264284081-en), with the difference that the experiment was carried out until the last bee died and not for only 10 days as requested by the OECD directive 245.

##### 1°) Animals

This study was performed in Autumn 2020 for **Li-$Mo_2O_4$-EDTA** and Spring 2021 for **Na-$Mo_2O_4$-EDTA** on *Apis mellifera* workers from the CNRS Apiary in Gif-sur-Yvette, France. The study was conducted on controlled age bees. The bees were sampled before their emergence by placing a capped brood comb in an incubator at 35°C and 50% humidity. The day of their emergence, bees were collected, counted and placed in groups of 50 individuals into plexiglass cages of Pain-type [9] (see Figure SIII.19) in which they were kept during the whole experiment.

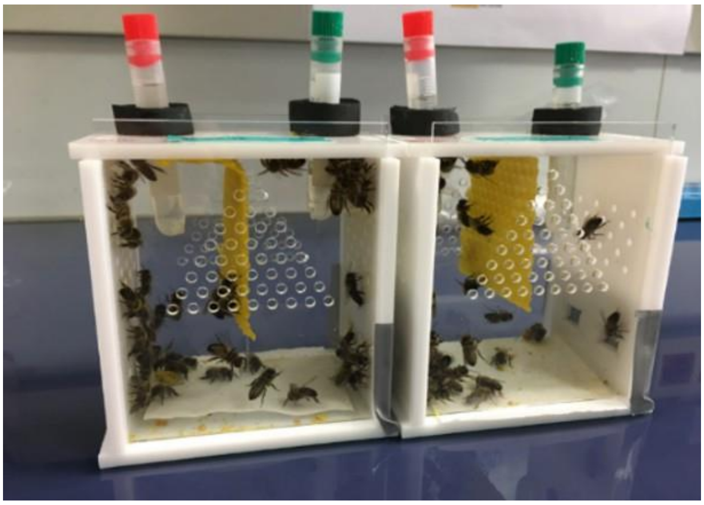

***Figure SIII.19** : "Pain"-type Cage used in this experiment.*

The cages contained a wax strip, a paper filter used to collect the faeces and 2 pierced tubes providing food and water to the bees inside the cage. The cages were kept at 35°C and 50% of relative humidity. During the first 8 days after emergence, bees had at their disposal a sugar solution (sucrose 50% w/w), water and pollen *ad libitum*. After 8 days, only water and sugar solution were provided *ad libitum* to the bees. To reduce as much as possible the impact on bees of possible temperature or humidity variations inside the incubator, the cages were moved with the incubator at every inspection (up/down; right/left; front/back).

**2°) Molybdenum treatments**

In this experiment, 3 different concentrations were used: 2, 20 and 400 mg/L for both **Na-$Mo_2O_4$-EDTA** or **Li-$Mo_2O_4$-EDTA** complexes. The right mass of complexes was dissolved in sucrose solution (50% w/w). The sucrose solution without complexes was used in the control cages. Three cages of 50 individuals were prepared per treatment for both complexes.

Oral exposure was used for all treatments, as it is the administration mode used in colonies. Bees were allowed to drink the molybdenum-enriched sugar solution *ad libitum* during all the experiment, which lasted until the last honey bee worker died.

Three times a week (Mondays, Wednesdays and Fridays), sucrose and water feeders were cleaned and refill, after that, dead bees were counted and took off from the cage. Then dead bees were stored in Eppendorf tubes labelled depending on their age, cage and treatment and placed at -20°C. Death rate, water and sugar consumption were reported every time.

**3°) Statistical analysis**

The analyses were performed with R (version 1.4.1106). The effect of **Na-$Mo_2O_4$-EDTA** and **Li-$Mo_2O_4$-EDTA** on honey bee survival was estimated by a survival analysis, based on Cox models using the function coxph from the package coxme. The quality of the model was estimated using the likelihood ratio.

#### III.4.2 Results

The results of all replicates are given in figures SIII.20 and SIII.21, respectively for **Na-$Mo_2O_4$-EDTA** and **Li-$Mo_2O_4$-EDTA**.

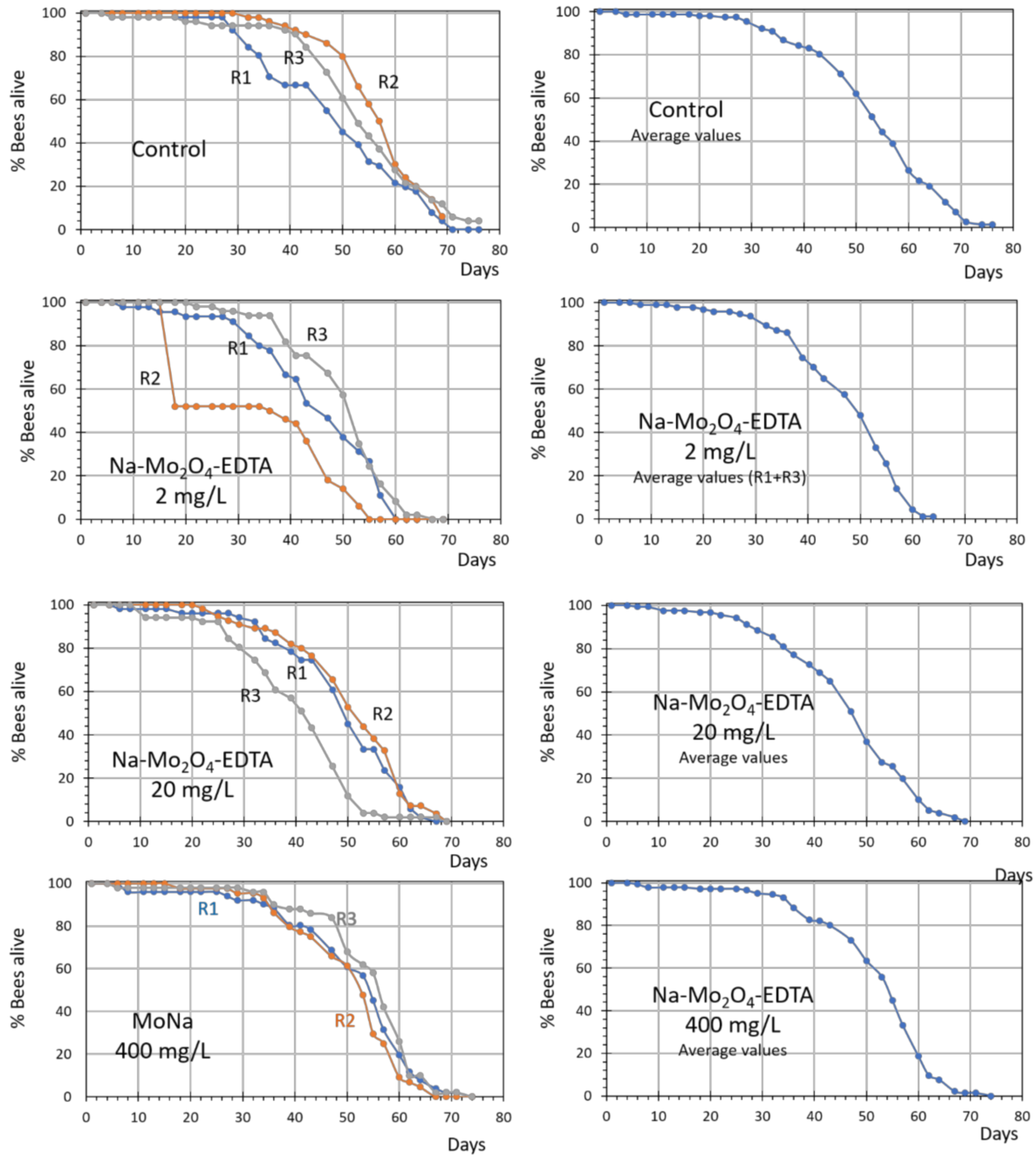


***Figure SIII.20 :** Survival curves obtained in control group and groups fed with **Na-Mo$_2$O$_4$-EDTA** at 2, 20 or 400 mg/L. The three replicates (cages of 50 individuals) are shown in the left column (R1, R2 and R3 curves), while the figure in the right column correspond to the average survival curve. Note that for the experiment at 2 mg/L, the replicate R2 is not considered in the average curve (see text).*

It should be noted that one cage of the second replicate (R2) treated with **Na-Mo$_2$O$_4$-EDTA** at 2 mg/L showed a sudden, strong mortality around day 15 due to the accidental obturation of the sugar feeder. As mortality was totally unrelated to treatment, this cage was not considered in average curves and for further analysis.

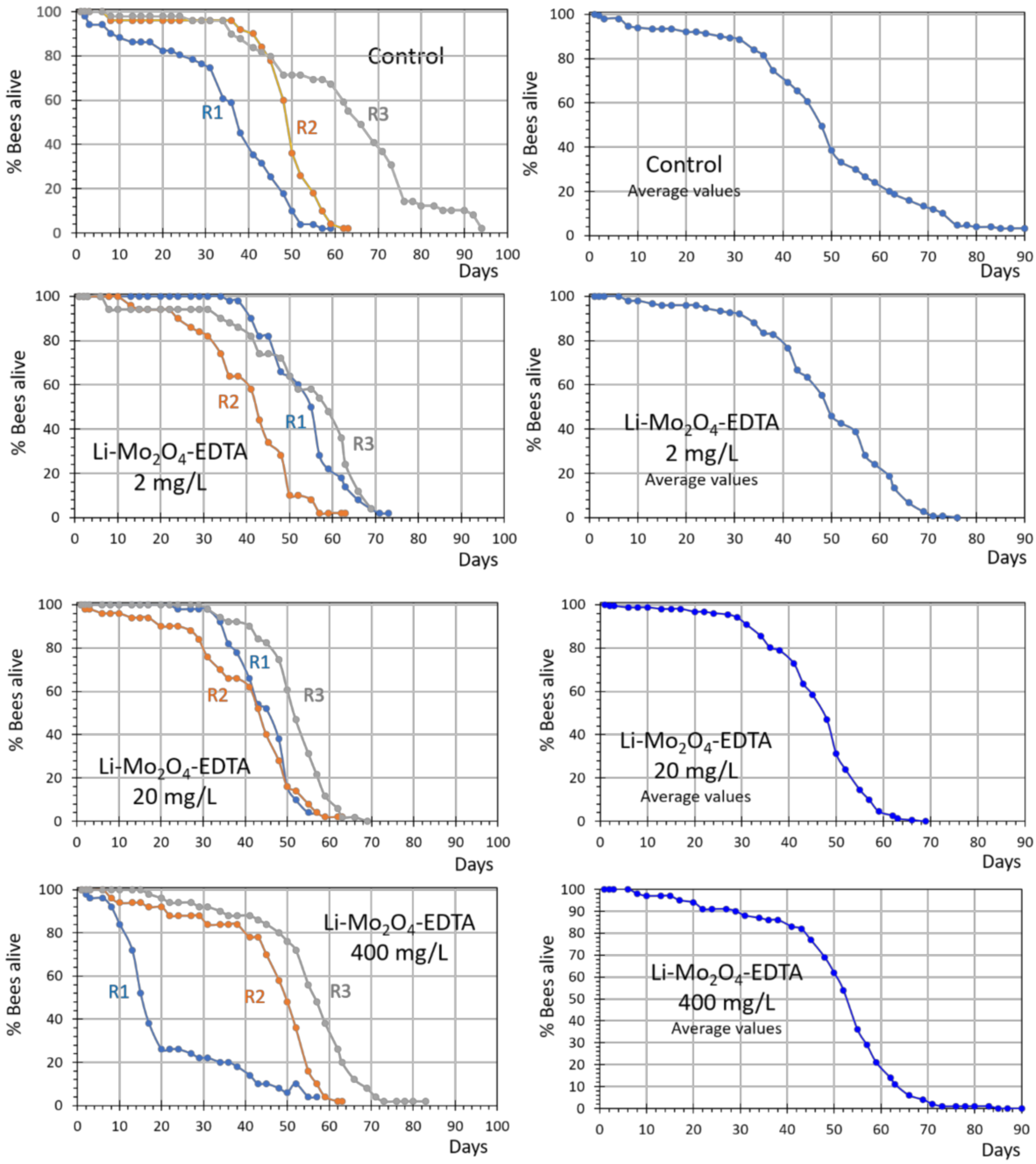


***Figure SIII.21 :** Survival curves obtained in control group and groups fed with **Li-$Mo_2O_4$-EDTA** at 2, 20 or 400 mg/L. The three replicates (cages of 50 individuals) are shown in the left column, while the figure in the right column correspond to the average survival curve. Note that for the experiment at 400 mg/L, the replicate R1 is not considered in the average curve (see text).*

Similarly, in the case of **Li-$Mo_2O_4$-EDTA**, a cage from the first replicate (R1) treated with **Li-$Mo_2O_4$-EDTA** at 400 mg/L showed an important and abnormal mortality rate during the first two weeks of the experiment. The origin of this behavior is not determined, but does not fit with observation in the two other groups. This cage was therefore not included in the average curve nor in the analysis.

Figures SIII.22 and SIII.23 presents the average curves obtained for the different doses and the controls for the **Na-$Mo_2O_4$-EDTA** and **Li-$Mo_2O_4$-EDTA** experiments, respectively.

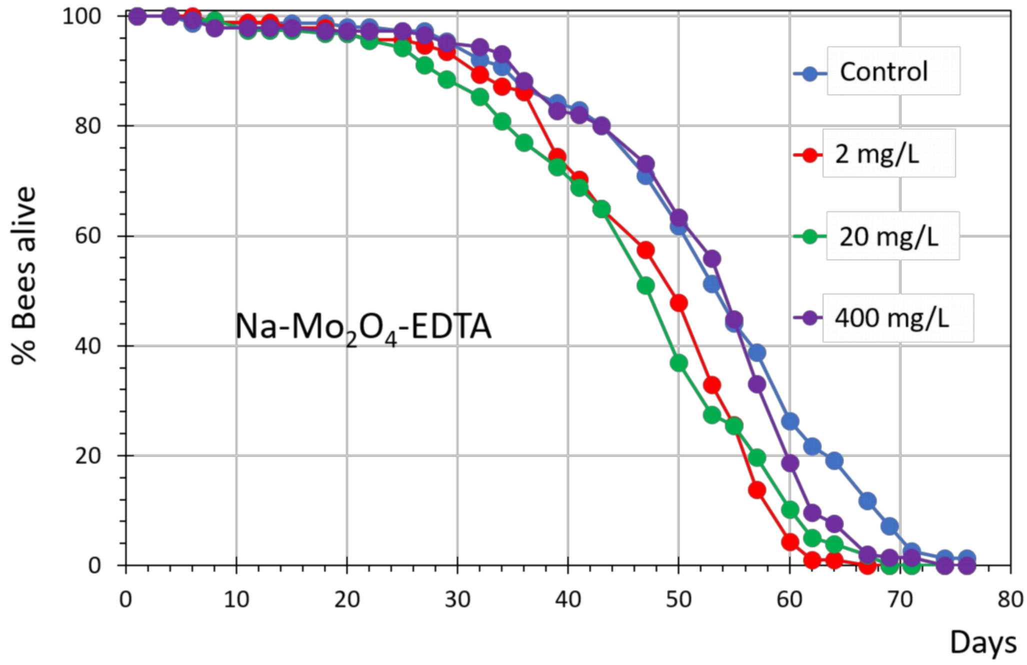


***Figure SIII.22 :*** *Superimposition od the average survival curves obtained in control group and groups fed with **Na-$Mo_2O_4$-EDTA** at 2, 20 or 400 mg/L.*

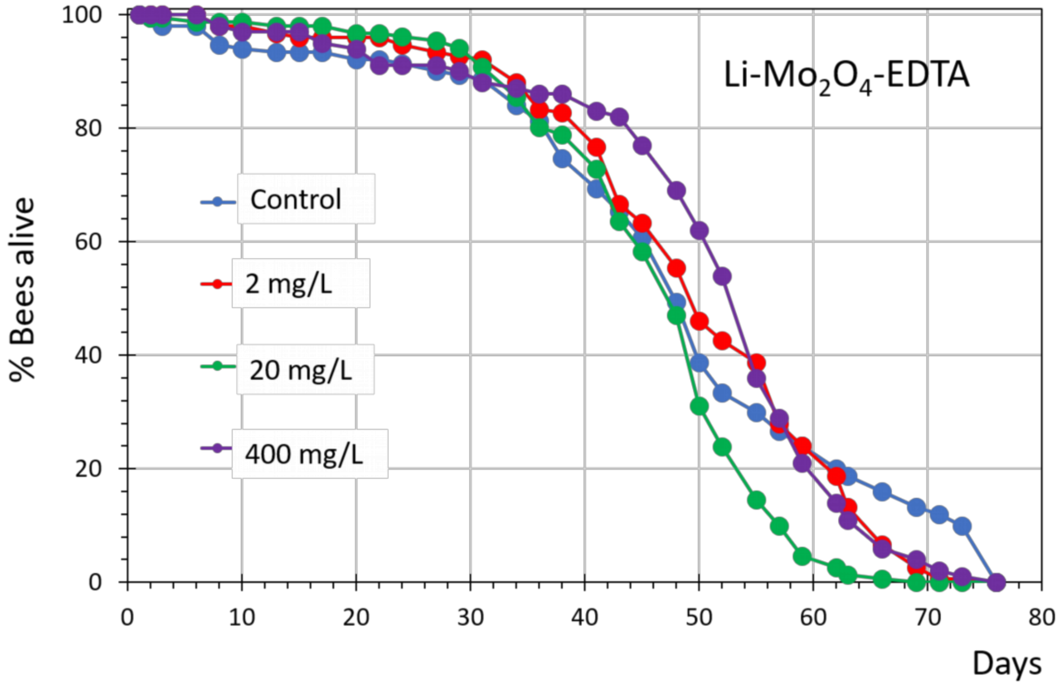


***Figure SIII.23 :*** *Superimposition od the average survival curves obtained in control group and groups fed with **Li-$Mo_2O_4$-EDTA** at 2, 20 or 400 mg/L.*

For both of complexes, three distinct periods can be observed on the mortality curves. The first phase, before 40 days, shows highly stable populations, with very little mortality. In the second phase, between 40 and 60 days, mortality starts and the death rate increases. Then, after 60-70 days, when populations have been strongly reduced, the death rate decreases again.

The OECD protocol No 245 typically evaluates mortality effects during the first period, i.e. during the first 10 days. Here, the conclusion of this protocol is that no effect of any of the doses of the two complexes can be seen on honey bee mortality in the first 10 days (Table SIII.10)

***Table SIII.10 :*** *Results of the Cox model for bees fed with molybdenum complexes (**Na-$Mo_2O_4$-EDTA** or **Li-$Mo_2O_4$-EDTA**) during 10 days. Each treatment is compared with the control group.*

| **Treatment** | **Concentration** | **β** | 95% CI | **HR** | **p-value** |
|---|---|---|---|---|---|
| **Na-$Mo_2O_4$-EDTA** | 2 mg/L | -0.647 | [0.154 ;1.782] | 0.52 | 0.60 |
| | 20 mg/L | -0.730 | [0.142 ;1.64] | 0.48 | 0.55 |
| | 400 mg/L | 0.447 | [0.628 ;3.896] | 1.56 | 0.62 |
| ***Li-$Mo_2O_4$-EDTA*** | 2 mg/L | -1.210 | [0.102 ;0.868] | 0.29 | 0.28 |
| | 20 mg/L | -1.536 | [0.069 ;0.669] | 0.22 | 0.16 |
| | 400 mg/L | 0.064 | [0.748 ;4.807] | 1.07 | 0.94 |

***Table SIII.11 :*** *Results of the Cox model for bees fed with molybdenum complexes* **Na-$Mo_2O_4$-EDTA** or ***Li-$Mo_2O_4$-EDTA*** *during 74 and 97 days respectively. Each treatment is compared with the control group.*

| **Treatment** | **Concentration** | **β** | 95% CI | **HR** | **p-value** |
|---|---|---|---|---|---|
| **Na-$Mo_2O_4$-EDTA** | 2 mg/L | 0.635 | [1.44 ;2.47] | 1.89 | 0.018 |
| | 20 mg/L | 0.841 | [1.78 ;3.02] | 2.32 | 0.002 |
| | 400 mg/L | 0.06 | [0.82 ;1.38] | 1.07 | 0.81 |
| ***Li-$Mo_2O_4$-EDTA*** | 2 mg/L | 0.841 | [1.06 ;5.08] | 2.32 | 0.28 |
| | 20 mg/L | 1.116 | [1.39 ;6.70] | 3.05 | 0.16 |
| | 400 mg/L | 0.497 | [0.75 ;3.60] | 1.64 | 0.53 |

We nevertheless continued the experiment throughout the lifetime of all the bees of the cages (see Table SIII-11). We found that generally, the average curves obtained for control batches and for bees fed with 2, 20 and 400 mg/L solutions of both complexes are very close (see Figures SIII.22 and SIII.23). Because of the high numbers of individuals tested in each case, statistical analysis indicates significant effects for some doses. For **Na-$Mo_2O_4$-EDTA,** the lower doses of 2 mg/L (HR : 1.89, CI : 1.44-2.47, P=0.018) and 20 mg/L (HR : 2.32, CI : 1.78-3.02,

P=0.002) induced slightly, but significantly, higher mortality than the control, but the higher dose of 400 mg/L was undistinguishable from the control (HR : 1.56, CI : 0.82-1.38, P = 0.81). For **Li-$Mo_2O_4$-EDTA,** no difference in mortality appeared between the treated and control groups (Table SIII.11). Generally, given the relatively low treatment effects and the high variability existing between individual cages treated in the same manner (see Figure SIII.21), **we conclude from these experiments that there is no notable positive effect, nor notable excess mortality linked to chronic feeding with the two complexes, even at high concentration**.

##### III.4.3 Conclusion

In this study we have estimated the effect of 2 molybdenum complexes, **Na-$Mo_2O_4$-EDTA** or **Li-$Mo_2O_4$-EDTA**, on honey bee survival at short-term (10 days) and long-term (>60 days) under Laboratory conditions. The conclusion of this study is that feeding honey bees with the complexes does not induce any marked positive or deleterious impact.

### III.5- Tolerance study of complex Na-$Mo_2O_4$-EDTA in beehives

To complement the laboratory studies, a tolerance study was carried out in an experimental apiary in France involving 60 bee hives in total. This study was conducted according to European and national regulatory requirements and according to the guidance on the assessment of the efficacy of feed additives, EFA Journal 2018, 16-5), 5274 (https://doi.org/10.2903/j/efsa/2018.5274). The objective of this study was to determine if an overdosage of the complex **Na-$Mo_2O_4$-EDTA** in syrup given to beehives leads to higher mortality of the bees during a two months experiment.

At D-15 to D0, the honey bee colonies and the sister queens were selected to get homogeneous groups of colonies. At D0 and D7 all colonies were fed with 2 L of syrup. The 60 beehives were divided into 5 groups randomly distributed:

- 12 beehives fed with 2x2L of syrup only (control group)
- 12 beehives fed with 2x2L of syrup containing **Na-$Mo_2O_4$-EDTA at 1 mg/L** ("4 mg" group)
- 12 beehives fed with 2x2L of syrup containing **Na-$Mo_2O_4$-EDTA at 2 mg/L** ("8 mg" group)
- 12 beehives fed with 2x2L of syrup containing **Na-$Mo_2O_4$-EDTA at 4 mg/L** ("16 mg" group)
- 12 beehives fed with 2x2L of syrup containing **Na-$Mo_2O_4$-EDTA at 20 mg/L** ("80 mg" group)

This experiment was also used for tracking the complex within the hives The experimental details are given in Part V of the supporting information. The number of dead bees has been tracked using collection box (Gary traps) disposed at the front of 8 hives from each modality tested. The collections boxes have been checked every 3 days to avoid degradation of individuals and consumption by predators from D0 until the end of the experiment at D56.

#### Results

The results are given in Figure SIII.24 for the 5 groups of beehives, while raw data are given in Table SIII.12.

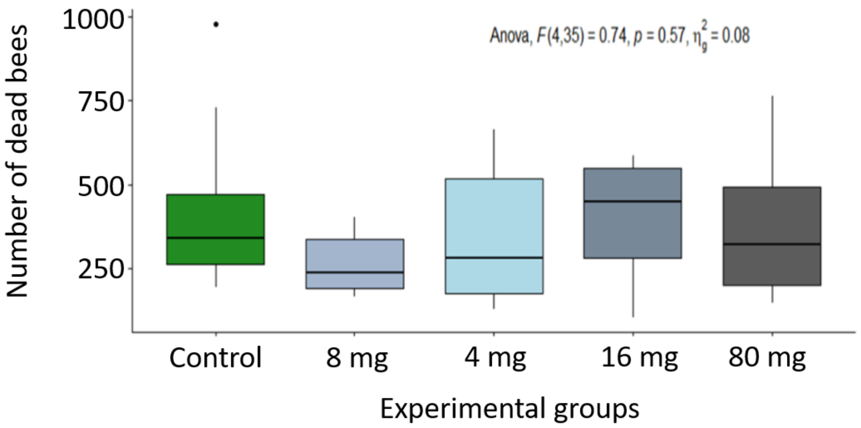


***Figure SIII.24.*** *Mean total number of dead bees recorded every 3 days during the experiment and according to the experimental groups. No significant differences were observed between the groups (p-value = 0.57).*

***Table SIII.11 :*** *Results of bees mortality from D0 to D56 for each group of beehives.*

| Date | Group | Dead bees | | |
|---|---|---|---|---|
| | | Sum of the 8 hives sampled | mean | SD |
| D0 | Control | 0 | 0 | 0 |
| | 4 mg | 0 | 0 | 0 |
| | 8 mg | 0 | 0 | 0 |
| | 16 mg | 0 | 0 | 0 |
| | 80 mg | 0 | 0 | 0 |
| D4 | Control | 319 | 46 | 23 |
| | 4 mg | 290 | 36 | 28 |
| | 8 mg | 431 | 54 | 37 |
| | 16 mg | 364 | 46 | 20 |
| | 80 mg | 306 | 38 | 30 |
| D7 | Control | 135 | 17 | 13 |
| | 4 mg | 133 | 17 | 10 |
| | 8 mg | 111 | 14 | 14 |
| | 16 mg | 183 | 23 | 10 |
| | 80 mg | 78 | 10 | 10 |
| D11 | Control | 315 | 39 | 38 |
| | 4 mg | 161 | 20 | 13 |
| | 8 mg | 156 | 20 | 15 |
| | 16 mg | 148 | 19 | 11 |
| | 80 mg | 131 | 16 | 15 |
| D13 | Control | 64 | 8 | 15 |
| | 4 mg | 61 | 8 | 8 |
| | 8 mg | 24 | 3 | 2 |
| | 16 mg | 49 | 6 | 4 |
| | 80 mg | 25 | 3 | 3 |
| D18 | Control | 206 | 26 | 47 |
| | 4 mg | 49 | 6 | 7 |
| | 8 mg | 42 | 5 | 5 |
| | 16 mg | 128 | 16 | 13 |
| | 80 mg | 62 | 8 | 10 |
| D21 | Control | 167 | 21 | 29 |
| | 4 mg | 50 | 6 | 4 |
| | 8 mg | 44 | 6 | 2 |
| | 16 mg | 215 | 27 | 20 |
| | 80 mg | 530 | 66 | 115 |

| | | | | |
|---|---|---|---|---|
| D26 | Control | 304 | 38 | 53 |
| | 4 mg | 302 | 38 | 72 |
| | 8 mg | 88 | 11 | 6 |
| | 16 mg | 455 | 57 | 57 |
| | 80 mg | 266 | 33 | 46 |
| D28 | Control | 93 | 12 | 11 |
| | 4 mg | 223 | 28 | 68 |
| | 8 mg | 80 | 10 | 11 |
| | 16 mg | 132 | 17 | 13 |
| | 80 mg | 24 | 3 | 2 |
| D33 | Control | 230 | 29 | 16 |
| | 4 mg | 140 | 18 | 9 |
| | 8 mg | 123 | 15 | 7 |
| | 16 mg | 482 | 60 | 65 |
| | 80 mg | 129 | 16 | 15 |
| D35 | Control | 124 | 16 | 16 |
| | 4 mg | 91 | 11 | 12 |
| | 8 mg | 63 | 8 | 8 |
| | 16 mg | 393 | 49 | 44 |
| | 80 mg | 107 | 13 | 26 |
| D39 | Control | 610 | 76 | 85 |
| | 4 mg | 614 | 77 | 82 |
| | 8 mg | 378 | 47 | 57 |
| | 16 mg | 1947 | 243 | 168 |
| | 80 mg | 586 | 73 | 89 |
| D42 | Control | 161 | 20 | 22 |
| | 4 mg | 75 | 9 | 9 |
| | 8 mg | 46 | 6 | 4 |
| | 16 mg | 167 | 21 | 14 |
| | 80 mg | 586 | 73 | 89 |
| D46 | Control | 241 | 30 | 39 |
| | 4 mg | 570 | 71 | 136 |
| | 8 mg | 125 | 16 | 12 |
| | 16 mg | 785 | 98 | 182 |
| | 80 mg | 322 | 40 | 50 |
| D49 | Control | 285 | 36 | 21 |
| | 4 mg | 417 | 52 | 60 |
| | 8 mg | 217 | 27 | 19 |
| | 16 mg | 834 | 104 | 179 |
| | 80 mg | 367 | 46 | 44 |

| | | | | |
|---|---|---|---|---|
| D56 | Control | 207 | 26 | 14 |
| | 4 mg | 173 | 22 | 17 |
| | 8 mg | 187 | 23 | 12 |
| | 16 mg | 394 | 49 | 41 |
| | 80 mg | 210 | 26 | 20 |

**Conclusion**

The results of this study indicate that no significant effect on the bees mortality by feeding colonies by the complex **Na-$Mo_2O_4$-EDTA** in syrup with a global dose in the 8-80 mg range per hive.
