## Supplementary material for "Food supplementation with molybdenum complexes improves honey bee health": Tests in beehives

### *Supporting Information*

#### Part IV. Tests in beehives and analyses of honey

IV.1- Tests in the apiary of Institute of Zoology, Moldova: 2013-2019
IV.2- Tests in France in operating conditions: 2019.
IV.3- Tests in California, USA: 2019-2020
IV.4- Tests in Greece: 2020-2021.
IV.5- General conclusions
V.6- Raw data

#### Introduction

As a follow-up to studies on the toxicity, stability and uptake of certain molybdenum complexes in bees, a selection of complexes was first tested in the academic apiary of the Institute of Zoology, Moldavian State University on *Apis melliferra carpatica* bees in forest areas in the Chisinau region, Moldova. The aims of these studies, presented in **part IV.1**, was to evidence a positive action on bee colonies of our complexes, to identify the best candidate and to find the main morphological parameters that can be affected by our complexes.

The experimental conditions used in the studies presented in this first part are perfect for an academic apiary or for small beekeeping exploitations, but not very well suited to large-scale operations as seen in professional beekeeping in Europe or in the USA, for example. Consequently, in **part IV.2**, new tests were carried out in France under real beekeeping conditions with a single feeding to show that professional use is possible and that the effects measured are significant. In addition, in a second test campaign, we wanted to see whether feeding with our complexes would make it possible to dispense with certain, often toxic, drug treatments against *Varroa*. These two questions will be assessed in the two test campaigns presented in this part. Nevertheless, the downside of testing under real operating conditions is that a smaller number of parameters can be monitored.

Bee colonies suffer an increased mortality rate during the autumn-winter period. In **part IV.3**, two test campaigns are carried out in California, USA, a region with an important annual mortality rate to evidence a protective effect of our complexes during the wintering period.

Finally, in **part IV.4**, a test campaign was carried out in Greece to see whether over-dosing with **Li-$Mo_2O_4$-EDTA** compound could have deleterious effects on colonies. Activity against *Varroa* and fungus *Nosema* are also reported in this part.

A general conclusion is given in part **IV.5** and all raw data are given in part **IV.6**.

#### IV.1- Tests in the apiary of Institute of Zoology, Moldova (2013-2019)

For this study, we focused on complexes $[Mo_2O_4(EDTA)]^{2-}$, $[Mo_2O_2S_2(EDTA)]^{2-}$ and $[Mo_2O_2S_2(L\text{-}cys)_2]^{2-}$, which appear the most stable ones among a larger series of complexes (See Part II of the Supporting Information). Initially, we found the $PPh_4^+$ salt of complex $[Mo_2O_4(EDTA)]^{2-}$ interesting, since we previously demonstrated that the $PPh_4^+$ cation can provide an additional antibacterial property [1]. The compound denoted **$PPh_4$-$Mo_2O_4$-EDTA** was the first tested in beehives in 2013 in comparison with a control batch and a commercial reference product based on Spirulina (Apispir). The first encouraging results were the subject of a first patent in 2016 [2].

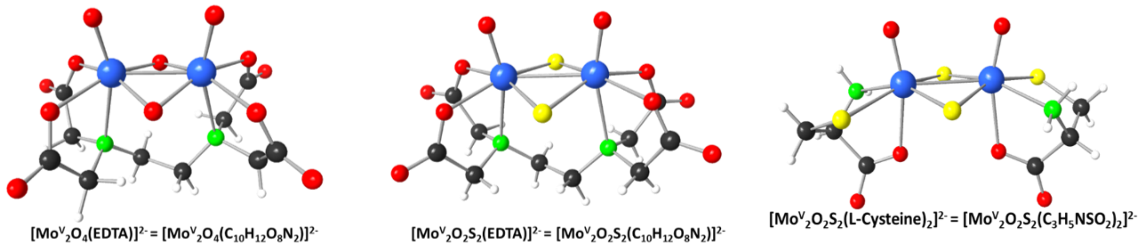


***Figure SIV.1.*** *Structures of the three complexes tested in beehives*

Subsequently (2016), we tested an analogous compound with sulfide-bridging ligands instead of oxo-bridging ligands, *i.e.* $[Mo_2O_2S_2(EDTA)]^{2-}$ as $PPh_4^+$ salt (denoted **$PPh_4$-$Mo_2O_2S_2$-EDTA**) and complex $[Mo_2O_2S_2(L\text{-}cys)_2]^{2-}$ as potassium salt (**K-$Mo_2O_2S_2$-LCys**), which also showed good chemical stability and little toxicity (see Parts II and III, SI). These tests enabled us to show that complex $[Mo_2O_4(EDTA)]^{2-}$ is the most effective. All other tests were then carried out with this complex in the form of $PPh_4^+$ salt (**$PPh_4$-$Mo_2O_4$-EDTA**), $Na^+$ salt (**Na-$Mo_2O_4$-EDTA**) and $Li^+$ salt (**Li-$Mo_2O_4$-EDTA**), in varying doses (2018 and 2019).

All these tests are grouped together and described below, with the number of hives involved, the protocols followed, the overall results and the measurements obtained for each hive, along with a statistical study for each parameter measured. The results of honey analyses carried out over 1 test campaigns are also presented.

##### IV.1.1 Testing protocols

The biological *in natura* tests were performed on domestic European honeybee families of *Apis mellifera carpatica* breed in an experimental apiary situated in Ghidighici forest district (Moldova) by the Apiculture team of the Institute of Zoology of the Republic of Moldova.

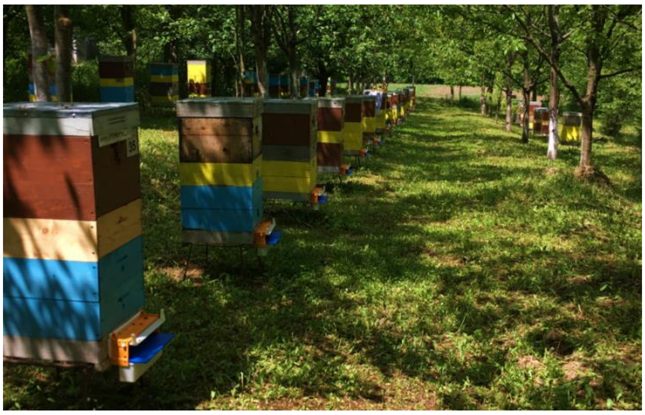

***Figure SIV.2.*** *Beehives of the experimental apiary in Moldova*

To administer the complexes to bees, the latter are added into two types of honeybees' feeding sugary products: candy (70% sucrose, 30% honey) and syrup (50% sucrose, 50% water). The dose of the complex, whatever its nature, in the candy is 0.4-1.2 mg/kg and in the syrup is 0.2-0.6 mg/L. The candy and syrup are used to feed the bees in the early spring period of their activity when there is pollen and nectar deficiency in nature.

The experiment is done simultaneously on at least 2 groups of the same number of beehives (between 10-16 beehives for each). The first group is the control group, where the bees receive simple candy and sugar syrup. The other groups are dedicated to the tested and/or commercial reference compounds. In these groups, the bees receive the candy and the syrup enriched with the complexes or reference molecules.

The feeding of bees is done by adding the candy/syrup in a container situated in the interior upper part of the hive. The volume of syrup is adjusted depending on the number of frames with bees at the initial moment, more precisely 200 g of candy per frame and 100 mL of syrup per frame.

For the first step of feeding, the candies are given only once. Then for the main feeding, determined volumes of syrup are repeatedly used to feed the bees every two days for a period of two weeks, which translates by approximately **a total of 2 to 6 mg of each compound** for each beehive.

From the beginning of the test until the end (after the second harvest), all the bee families are monitored for several key morpho-productive parameters that reflect the colony development level and vital aspects of bees. The studied parameters are: hygienic behaviour, queen’s prolificity (fecundity), beebread production, colony strength (=adult population), honey production and wax production. The monitoring of these parameters agrees with the *Zootechnical norms regarding the evaluation of bee families, the breeding and certification of the beekeeping parent material* approved by Government Decision Nr 306 on 28/04/2011 of the Republic of Moldova. Amended HG214 of 12.04.23, MO126/13.04.23 art. 273; in force 26.08.23 [3]. More details are given below.

**Assessment of hygienic behaviour**

The ability to fight contagious pathogens is determined by measuring the hygienic behavior of the bee family. To identify bee families with pronounced hygienic behavior, standard tests are used, whereby the brood on a compact surface is artificially killed in order to determine the speed and accuracy with which the bees identify and eliminate the dead brood. The evaluation is carried out in May-July, twice on the same family of bees, under different environmental conditions and at different time intervals. The brood is killed in the capped stage (*pupa*) by puncturing with a fine needle through the cap of cells on a portion of honeycomb in the family nest, on a square surface of 5 x 5 cm (100 cells, see Figure SIV.3) marked in the corners with matches. After 24 hours, the number of cells from which the dead brood was removed is estimated. The ratio between the removed and the initially killed offspring on the marked surface of the honeycomb, expressed as a percentage, represents disease resistance.

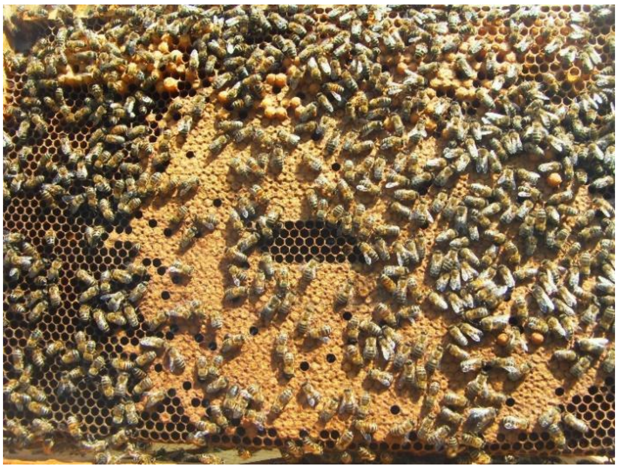

***Figure SIV.3.*** *View of the brood inside the hive. Evaluation of resistance to diseases is made on such a brood at this stage of development when the cells are capped (pupa stage of the bees). Some brood is killed in the capped stage with a fine needle.*

**Assessing the brood quantity and the prolificity of the queen**

Prolificity of the queen bee (eggs/24 hours) is determined at the end of spring (May 20-31) by dividing the number of cells with brood stages to 12 (the duration of the development of the brood in days), resulting in the number of eggs laid within 24 hours. The number of cells in the nest is measured by using the Netz frame (a frame with standard Langstroth size, whose surface is divided into 32 equals squares). The number of squares (5 x 5 $cm^2$) occupied by brood is multiplied by 100, resulting in the total number of cells with the capped brood.

**Assessing the strength of the colony**

The strength of the bee colony is represented by the number of adult bees in the nest at the time of appreciation. The assessment is carried out three times a year: at the spring review (March-April), at the end of spring (20-31 May) and at the autumn review (September). Following these three assessments, the average strength of the bee colony is determined. The quantity of bees (kg) is determined by multiplying the number of intervals between frames, uniformly occupied with bees, with the coefficient 0.25 for the standard Dadant frame (435x300 mm) and 0.2 for the standard Langstroth frame (435x230 mm) as defined by Zootechnical norms [3].

**Evaluation of honey production**

Honey production is determined for each family of bees by summing the quantity of honey-merchandise, extracted during the harvest season, with the amount of honey accumulated in the nest and left (at the autumn revision) as bee food for the winter period. The quantity of honey-merchandise is determined at each extraction, by weighing honeycombs before and after extraction (with a precision of 0.1 kg), the difference in weight constituting the quantity of honey-merchandise extracted. The amount of honey left in the nest for bee feeding is determined by the autumn (September) revision, by weighing the honeycomb frames and substracting (from their total weight) the total weight of the standard honeycomb frames: for the Dadant type frame (435x300 mm) - 0.6 kg, for Langstroth frame (435x230 mm) - 0.5 kg.

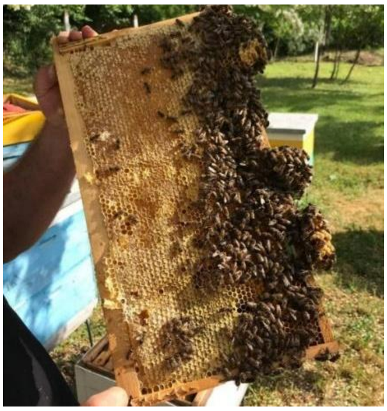

***Figure SIV.4.*** *Honeycomb frame with honey supply stored in capped cells*

**Assessing the degree of *Varroa* infestation on bees.**

*Varroa* is considered to be one of multiple stress factors contributing to the high levels of bee losses around the world.

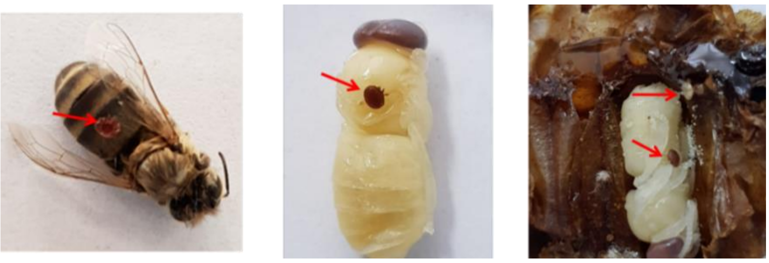

***Figure SIV.5.*** *Varroa destructor (red arrows) is an external parasitic mite that feeds on the honey bees* Apis cerana *and Apis mellifera.*

Evaluation of infestation rate involves counting the number of *Varroa* mites on a sample of adult bees. The value provides an index for monitoring the level of parasitism in the colony.

The method uses powdered sugar and a transparent glass "shaker" jar with a capacity of 1kg. The lid is made of galvanized steel mesh of the 3mm mesh type. The mesh allows *Varroa* mites to pass through, but retains the bees.

The method is implemented in the following stages: shake 40 to 50 g of bees from an outer frame of a bee colony into the jar (approximately 400 bees), add 100 g of powdered sugar and roll the shaker on itself for 1 minute to cover all the bees with powdered sugar. The powdered sugar loosens the *Varroa* mites from the bee's body, and they fall off. Leave to stand for 1 minute: the bees' delousing behavior reinforces the fall of the *Varroa* mites. The powdered sugar is sieved to retain and count the *Varroa* mites. The result is expressed as the number of mites per 10 g of bees (approximately 100 bees), and can be expressed as a percentage %. This rate is indicative of *Varroa* infestation in the colony. A rate between 1 and 2% indicates low Varroa infestation. A moderate rate is between 2 and 5%, in which case colony treatment should be scheduled. A rate above 5% requires emergency treatment of the colony.

The degree of brood infestation with *Varroa* mites is determined in each family by examining 100 uncapped cells with drone brood, located in a compact area on the honeycombs, where the mites were counted. Indeed, it is known that in the drone brood, the rate of cell infestation is higher than in worker brood because drone brood cells are larger and the post-capping stage is longer which allows the mite to produce more offspring per cycle. The total number of found mites in these cells constitutes the degree of infestation. If up to 20 mites are found per 100 cells with drone brood, the infestation is considered low. A rate of 20 to 30 mites in cells with brood drone determine moderate infestation. A rate with more than 30 mites of brood cells is considered strong.

**Statistics.**
Even if the colonies are of comparable size at the start of the test, the variations in the results obtained in the hives are sometimes very important. Apiary size is limited. Consequently, the number of hives per batch is limited to between 10 and 16, depending on the number of batches to be tested. More modality to be tested, less the number of hives for each modality. A compromise must be found. For the results obtained (raw data provided at the end of Part IV of this Supporting Information), standard deviations are calculated. Shapiro and F tests were first applied to see if the data follow a normal law. As the function of the results Student (T-test), U-test from Mann-Withney or Kruskall-Wallis tests were applied to address the significance of the results obtained. These tests are not detailed in this part of the supporting Information. Nevertheless, the most significant results are discussed in the main teste with the corresponding p-values.

##### IV.1.2 Results

##### 1°) Test campaigns 2013 and 2016

The aim of the first test campaigns of 2013 and 2016 was to identify the most promising complex among the three complexes selected because of their high stability and their low or absence of toxicity (see part II of the Supporting Information). For the first test performed in 2013, we focused on the complex **PPh$_4$-Mo$_2$O$_4$-EDTA**, which was patented in 2016 [2], while the test campaign of 2016 was dedicated to its sulfurated analogue **PPh$_4$-Mo$_2$O$_2$S$_2$-EDTA** to analyze the effect the nature of the molybdenum cluster core and **K-Mo$_2$O$_2$S$_2$-LCys** to study of the effect of the nature of the ligand. A total of 2 mg for each compound was tested through the protocol described above. Note that for the two formers, since the molecular masses are similar it corresponds approximatively to the same amount of Mo. For the third complex, the molecular mass is almost two times lower. The concentration in Mo is thus twice higher.

For a syrup containing 0.2 mg/L the molecular concentrations are:

- $2.9*10^{-7}$ mol/L for **K-$Mo_2O_2S_2$-Cys** (Mm = 694.58 g/mol)
- $1.5*10^{-7}$ mol/L for **$PPh_4$-$Mo_2O_2S_2$-EDTA** (Mm = 1309.04 g/mol)
- $1.5*10^{-7}$ mol/L for **$PPh_4$-$Mo_2O_4$-EDTA** (Mm = 1294.9 g/mol)

The results for each parameter are shown in the table SIV.1 below, while the details of the data are given in Tables SIV.9-SIV.14.

***Table SIV.1*** *Results obtained for the main parameters monitored for the 2013 and 2016 test campaigns* (mean ± se). *For the tested complexes, variation to the control group is given in %*

| | **2013** | | **2016** | | |
|---|---|---|---|---|---|
| | **Reference** | **$PPh_4$-$Mo_2O_4$-EDTA** | **Reference** | **K-$Mo_2O_2S_2$-LCys** | **$PPh_4$-$Mo_2O_2S_2$-EDTA** |
| **Number of bee colonies** | 16 | 16 | 12 | 12 | 12 |
| **Hygienic behaviour, %** | 88.40 ± 0.4 | 92.20 ± 0.4<br>+4.3% | 86.3 ± 1.8 | 92.2 ± 1.7<br>+6.8% | 92.0 ± 1.3<br>+6.6% |
| **Queen bee's prolificity, eggs/24h** | 1590 ± 20 | 1757 ± 15<br>+10.5% | 1133 ± 67 | 1245 ± 41<br>+9.0% | 1248 ± 33<br>+9.3% |
| **Capped brood quantity, hundreds of cells** | 190.8 ± 2.4 | 210.8 ± 1.7<br>+9.5% | 143.0 ± 3.4 | 149.4 ± 5.0<br>+4.5% | 149.8 ± 3.9<br>+4.8% |
| **Viability of the brood, %** | 89.30±0.30 | 91.30±0.20<br>+2.2% | | | |
| **Colony strength, kg of bees** | 3.20 ± 0.02 | 3.58 ± 0.05<br>+11.9% | 1.12 ± 0.05 | 1.13 ± 0.04<br>+0.9% | 1.13 ± 0.04<br>+0.9% |
| **Beebread quantity, hundreds of cells** | 90.5 ± 1.8 | 110.20±2.70<br>+21.8% | 89.3 ± 2.9 | 90.4 ± 4.2<br>+1.2% | 94.1 ± 3.0<br>+5.4% |
| **Wax quantity, number fabricated honeycombs** | 0.28±0.08 | 0.39 ± 0.01<br>+39% | 0.29±0.01 | 0.32±0.01<br>+10.3% | 0.33±0.01<br>+13.7% |
| **Honey quantity, kg** | 11.62 ± 0.40 | 13.94 ± 0.36<br>+19.6% | 3.97 ± 0.13 | 4.20 ± 0.17<br>+6.0% | 4.40 ± 0.21<br>+10.6% |

As shown in Table SIV.1, we observe a beneficial effect on the colonies, which were fed in spring with any of the three compounds **K-$Mo_2O_2S_2$-Cys**, **$PPh_4$-$Mo_2O_2S_2$-EDTA**, **$PPh_4$-$Mo_2O_4$-EDTA**. For all compounds, despite being introduced in very small quantities, only 2 mg, the ability of the bees for hygien increases by 6.8%, the prolificity of the queen bee by 10.5% (167 extra eggs per day compared to the reference), the capped brood by 9.5%, the strength of the colony by 11.9%, the bee bread quantity by 21.8%, the honey production by 19.6% and the wax production even by 39%.

For all complexes, the followed parameters are higher than those measured for the group control. The comparison between **PPh$_4$-Mo$_2$O$_2$S$_2$-EDTA** and **PPh$_4$-Mo$_2$O$_4$-EDTA**, which only differ from the chemical environment of the Mo atoms in the clusters $[Mo_2O_2E_2]^{2+}$ (E = S or O), clearly shows that there's no advantage in using sulfur complex over oxo complex. In addition, toxicity studies on the sulfur complex showed hyperactivity in mice, whereas no effect was measured for the oxo analogue (see Part III, Supporting Information). Compound **PPh$_4$-Mo$_2$O$_4$-EDTA** therefore appears to be the better of the two.

The results obtained for the second sulfur complex, **K-Mo$_2$O$_2$S$_2$-LCys**, are also very limited compared with the other two, despite a concentration twice higher. It is likely that at this low concentration the complex is destroyed (see Part II, Supporting Information), which is not the case for **PPh$_4$-Mo$_2$O$_4$-EDTA**, and that the resulting product is probably less available.

The conclusion of these two campaigns is that we should focus on the **PPh$_4$-Mo$_2$O$_4$-EDTA** complex, which achieves the best results for almost all parameters, sometimes by far. For this compound, the hygienic behavior is improved by only +4.3 %, but the increased prolificity of the queen bee by +10.5 % provoked an increase of the number of capped brood (+9.5%), a slightly better viability of the brood (+2.2%) and a stronger colony (+11.9%). More bees in better shape logically induces an increase of the bee bread stocks (+21.8%), and an increase of the honey production (+19.6%). Note also that the quantity of wax is increased by +39% in comparison with the control group.

##### 2°) Test campaign 2018

New test campaigns were organized in 2017, 2018 and 2019 to evaluate the influence of the counter cations ($PPh_4^+$, $Na^+$ and $Li^+$) associated with the complex **$[Mo_2O_4(EDTA)]^{2-}$**, evaluate the impact on the *Varroa* infestation and optimize the efficiency of these complexes.

Unfortunately, the 2017 campaign could not be carried out in good conditions due to heavy snowfalls in April on the test apiary. We therefore present the results obtained in 2018 on 3 salts of the **$[Mo_2O_4(EDTA)]^{2-}$** complex.

- **PPh$_4$-Mo$_2$O$_4$-EDTA** previously tested
- **Li-Mo$_2$O$_4$-EDTA**, as Lithium is known to have an action against *Varroa* mites [4]
- **Na-Mo$_2$O$_4$-EDTA** for which the $Na^+$ cation is, *a priori*, without action and which will enable us to measure only the effect of the **$[Mo_2O_4(EDTA)]^{2-}$** complex only.

For the 2018 test campaign, the same experimental conditions as those used in 2013 and 2016 were used: a total of about 2 mg of each compound per beehive, which corresponds to molar concentrations in the syrup (0.2 mg/L) of $1.5*10^{-7}$ M for **PPh$_4$-Mo$_2$O$_4$-EDTA** and around 2.5 to $3*10^{-7}$ M for **Na-Mo$_2$O$_4$-EDTA** or **Li-Mo$_2$O$_4$-EDTA**, respectively.

The parameters which were followed are the same but in addition, for this 2018 test campaign, we evaluated the effect on *Varroa* infestation.

The results obtained for each parameter are shown in the table SIV.2 below, while the details of the data are given in Tables SIV.15 -SIV.18.

***Table SIV.2*** *Results obtained for the main parameters monitored for the 2018 test campaigns (mean ± se). For the tested complexes, the variation in relation to the reference group is given in %.*

| | **Control** | **$PPh_4$-$Mo_2O_4$-EDTA** | **Na-$Mo_2O_4$-EDTA** | **Li-$Mo_2O_4$-EDTA** |
|---|---|---|---|---|
| **Number of colonies** | 10 | 10 | 10 | 10 |
| **Hygienic behaviour, %** | 89.0 ± 1.3 | 94.8 ± 1.3<br>+6.5% | 90.4 ± 1.3<br>+1.6% | 91.6 ± 1.0<br>+2.9% |
| **Queen bee's prolificity, eggs/24h** | 1432 ± 113 | 1533 ± 56<br>+7% | 1622 ± 33<br>+13.3% | 1600 ± 52<br>+11.7% |
| **Capped brood quantity, hundreds of cells** | 171.9 ± 14.7 | 184.0 ± 6.8<br>+7.0% | 194.6 ± 4.0<br>+13.2% | 192.0 ± 6.2<br>+11.7% |
| **Colony strength, kg** | 2.41 ± 0.06 | 2.46 ± 0.07<br>+2.0% | 2.44 ± 0.04<br>+1.2% | 2.54 ± 0.04<br>+5.4% |
| **Beebread quantity, hundreds of cells** | 131.0 ± 3.3 | 131.0 ± 4.3<br>+0.0% | 120.5 ± 4.7<br>-8.0% | 123.8 ± 4.3<br>-5.5% |
| **Wax quantity, № fabricated honeycombs** | 0.30 ± 0.01 | 0.31 ± 0.01<br>+3.3% | 0.32 ± 0.01<br>+6.6% | 0.35 ± 0.01<br>+16.7% |
| **Varroa infestation. Number of Varroa / 10 g of bees** | 1.78 ± 0.14 | 1.40 ± 0.12<br>-21.3% | 1.54 ± 0.12<br>-13.5% | **1.02 ± 0.07<br>-42.7%** |
| **Honey quantity, kg** | 20.79 ± 3.49 | 23.50 ± 2.31<br>+13.0% | **31.10 ± 3.00<br>+49.6%** | **29.70 ± 3.75<br>+42.9%** |

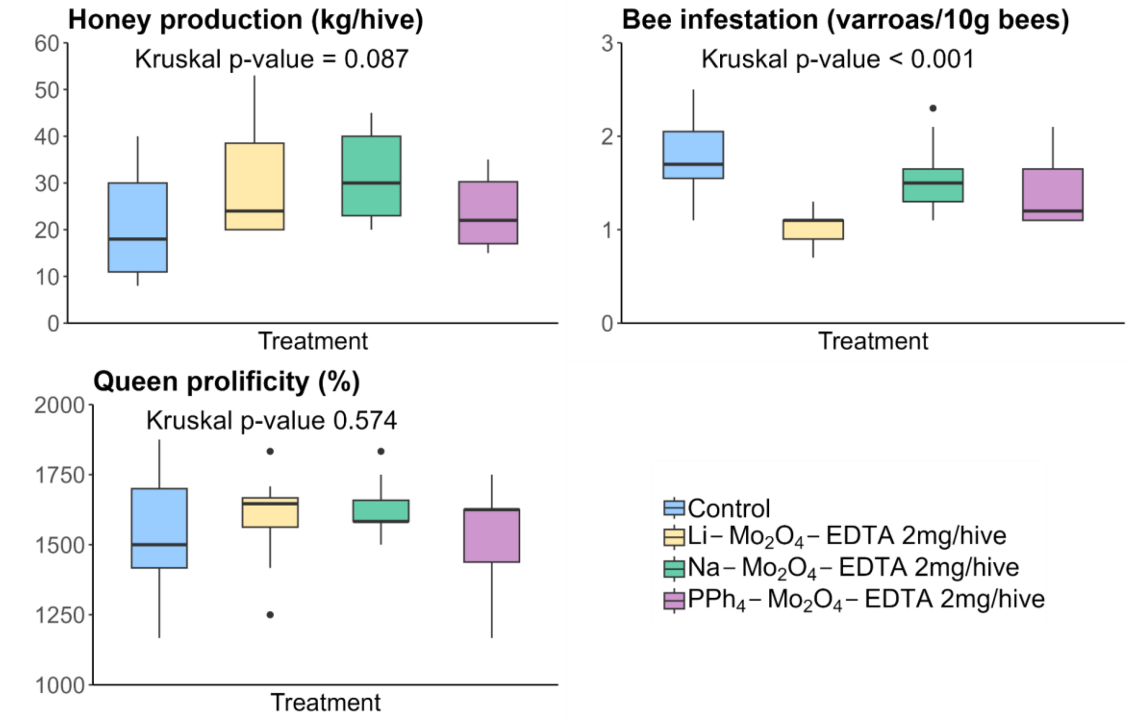


***Figure SIV.6**. Comparisons of the effects observed with Mo-based complexes vs the group control as reference. Test campaigns of 2018. Significant effects are observed for honey production and bee infestation with Varroa.*

As shown in Table SIV.2, the three salts of the complex $[Mo_2O_4(EDTA)]^{2-}$, i.e. $PPh_4$-$Mo_2O_4$-EDTA, Li-$Mo_2O_4$-EDTA, and Na-$Mo_2O_4$-EDTA exhibit positive effects in the colonies in comparison with the group control but the two alkali salts of the complex appear more interesting.

The prolificity of the queen bee is increased by +13.3 % (190 extra eggs per day) for Na-$Mo_2O_4$-EDTA and not far from the result obtained with Li-Mo2O4-EDTA (+11.7%), while the value obtained with $PPh_4$-$Mo_2O_4$-EDTA is lower (+7% only this year *vs* +10.5 % in 2013).

The main result of this test campaign comes from the effect against the *Varroa* infestation. All three complexes show a reduction in the degree of Varroa infestation, but the effect is limited with the **$PPh_4$-$Mo_2O_4$-EDTA** and **Na-$Mo_2O_4$-EDTA** salts. On the contrary, the lithium salt **Li-$Mo_2O_4$-EDTA** shows a remarkable activity, since the infestation rate is reduced by over 42.9% on worker bees. Lithium cation is known to have a deleterious effect on Varroa [4], but Ziegelmann et al. reported effects of lithium salts at high concentrations in excess of $2.10^{-3}$M. **Here, we are dealing with Lithium concentrations of the order of $10^{-7}$ M, *i.e.* 4 orders of magnitude lower**. This result suggests a synergistic effect between the Mo complex and the $Li^+$ cation and the next campaign will be focused on this point.

Finally, the production of honey only increases of +13% with **$PPh_4$-$Mo_2O_4$-EDTA** but this value is not significant, while it rises to +42.9 % for **Li-$Mo_2O_4$-EDTA** and up to **+49.6 %** for **Na-$Mo_2O_4$-EDTA**, which translate by an average production of 31.10 ± 3.00 kg per hive *vs* 20.79 ± 3.49 kg for the group control. This increase due to only 2 mg of **Na-$Mo_2O_4$-EDTA** is likely only due to the complex **$[Mo_2O_4(EDTA)]^{2-}$** if we consider that $Na^+$ cation does not play a role, especially at this low concentration**.**

**Analysis of honey**

Of course, such an increase in honey production legitimately raises the question of the quality of the honey produced. To answer this question, we compiled average samples of honey for each group, and these were analyzed by the "Famille Michaud Apiculteurs" laboratory, Gan, France, an independent laboratory accredited by the INAO, by measuring sugar levels and a number of other important parameters required by European regulations (directives 2001/110/CE and 2014/63/UE). These parameters are hydroxymethylfurfural content, humidity, pH, electrical conductivity, amylase content and sugars content. The honey samples were collected in spring to be as close as possible to the feeding with Mo complexes.

For each honey sample, the Mo, Li, Na and P contents were also determined by ICP-MS (Limit of Quantification, LOQ = 0.01 mg/kg). The results are gathered in the Table SIV.3 below.

***Table SIV.3.*** *Results of the analyses performed on honey samples collected in the 4 groups on the 2018 test campaign. The limit of quantification (LOQ) is usually of 0.1 ppm for ICP-MS measurements and 0.1 % for HPLC study of sugars.*

| **Parameter** | **control** | **$PPh_4$-$Mo_2O_4$-EDTA** | **Na-$Mo_2O_4$-EDTA** | **Li-$Mo_2O_4$-EDTA** | **Specifications EU regulation.** |
|---|---|---|---|---|---|
| **Mo, mg/kg** | < LOQ | < LOQ | < LOQ | < LOQ | - |
| **Na, mg/kg** | 9.700 | 6.100 | 9.700 | 8.800 | |
| **Li, mg/kg** | 0.043 | 0.024 | 0.034 | 0.028 | |
| **P, mg/kg** | 60.000 | 50.700 | 55.500 | 60.600 | |
| **Hydroxymethylfurfural, mg/kg** | 2.67 | <2 | <2 | 2.07 | ≤ 40 mg/kg |
| **Humidity, %** | 17.2 | 15.9 | 15.7 | 15.7 | ≤ 20% |
| **Electric conductivity, µS/cm** | 252 | 189 | 239 | 256 | - |
| **pH** | 4.10 | 4.06 | 4.06 | 4.04 | - |

| | | | | | |
|---|---|---|---|---|---|
| **Amylase, *Schade* units** | 21.50 | 18.50 | 22.50 | 23.60 | ≥ 8 DZ Schade |
| **Glucose, %** | 26.0 | 25.8 | 26.3 | 26.4 | - |
| **Fructose, %** | 42.1 | 42.7 | 42.5 | 42.7 | - |
| **Isomaltose, %** | < LOQ | < LOQ | < LOQ | < LOQ | - |
| **Saccharose, %** | 1.8 | 3.8 | 3.0 | 2.4 | ≤ 5% |
| **Turanose, %** | 2.4 | 2.2 | 2.2 | 2.2 | - |
| **Melezitose, %** | < LOQ | < LOQ | < LOQ | < LOQ | - |
| **Maltose, %** | 1.3 | 1.6 | 1.7 | 1.7 | - |
| **Erlose, %** | 1.9 | 2.7 | 2.1 | 2.1 | - |
| **Trehalose, %** | < LOQ | < LOQ | < LOQ | < LOQ | - |
| **Ratio Fructose/Glucose** | 1.62 | 1.66 | 1.62 | 1.60 | - |
| **Total disaccharides, %** | 3.10 | 5.4 | 4.70 | 4.10 | - |
| **Other oligosaccharides, %** | < LOQ | < LOQ | < LOQ | < LOQ | - |

As shown in table SIV.3, the honey produced by the hives that received molybdenum supplements is similar in composition to the honey produced by the control group. Physico-chemical parameters comply with European Union specifications. In particular, the distribution of sugars is very similar between batches, and sucrose levels remain below 5% in all cases, in line with European specifications. It means that feeding bees with Mo complexes does not alter the enzymatic mechanisms involved in the production of honey.

Moreover, the presence of Mo was not detected in any honey, which shows that Mo complexes are consumed by bees and do not pass into the honey produced.

**3°) Test campaign 2019**

For the next campaign, we focused on the **Li-$Mo_2O_4$-EDTA** complex for its anti-varroa effect. This complex is tested with two different dosages (total of 2 and 6 mg per hive with a sugar syrup at 0.2 or 0.6 mg/L), a control batch, and a batch with hydrated lithium acetate as reference for lithium ions.

- **Li-$Mo_2O_4$-EDTA** : 2 kg of candy at 0.4 mg/kg (0.8 mg in total) then 6-7 L of syrup at 0.2 mg/L ($3.2x10^{-7}$ mol/L in complex ; $6.4x10^{-7}$ mol/L in $Li^+$). Global dose : 2 mg/hive.

- **Li-$Mo_2O_4$-EDTA** : 2 kg of candy at 1.2 mg/kg (2.4 mg in total) then 6-7 L of syrup at 0.6 mg/L ($9.6x10^{-7}$ mol/L in complex ; $1.92x10^{-6}$ mol/L in $Li^+$). Global dose : 6 mg/hive.

- **$LiCH_3COO.H_2O$**: 2 kg of candy at 0.4 mg/kg (0.8 mg in total) then 6-7 L of syrup at 0.2 mg/L ($2.36 \times 10^{-6}$ mol/L in $Li^+$). Global dose : 2 mg/hive

The experimental protocol is the same as for previous campaigns with a focus on the rate of *Varroa* infestation for bees and brood. In particular, for the last two groups, we focused only on these parameters.

The results obtained for each parameter are shown in the table SIV.4 below, while the details of the data are given in Tables SIV.19-SIV.22.

***Table SIV.4*** *Results obtained for the main parameters monitored for the 2019 test campaigns (mean ± se). For the complexes tested, the variation in relation to the control is given in %.*

| | **control** | **Li-$Mo_2O_4$-EDTA** 2mg | **Li-$Mo_2O_4$-EDTA** 6 mg | **$LiCH_3COO$** 2 mg |
|---|---|---|---|---|
| **Number of colonies** | 10 | 10 | 10 | 10 |
| **Hygienic behaviour, %** | 86.00 ± 1.07 | 94.00 ± 1.20 +9.3% | 96.40 ± 1.00 +12.0% | 89.8±1.4 +1.04% |
| **Queen bee's prolificity, eggs/24h** | 1629 ± 30 | 1746 ± 29 +7.2% | 1825 ±38 +12.0% | 1596±41 -2.1% |
| **Capped brood quantity, hundreds of cells** | 195.5 ± 3.61 | 209.5 ± 3.45 +7.2% | 219.0 ±4,6 +12.0% | 191.5±4.9 -2.1% |
| **Colony strength, kg** | 3.39 ± 0.10 | 3.90 ± 0.11 +15.0% | 4.05 ±0.08 +19.4% | 3.32±0.09 - 2.1% |
| **Beebread quantity, hundreds of cells** | 114.0 ± 4.5 | 140.0 ± 3.0 +22.8% | 137.5 ±3.4 +20.2% | 121.0±3.0 +6.1% |
| **Wax quantity, kg** | 0.27 ± 0.01 | 0.34 ± 0.01 +25.9% | 0.35 ±0.01 +29.6% | 0.29 ±0.01 +7.4% |
| **Varroa infestation. Number of Varroa / 10 g of bees** | 2.30 ± 0.40 | **1.22 ± 0.20 -47.0%** | **0.89 ± 0.16 -61.3%** | 1.60 ± 0.15 -30.4% |
| **Degree of brood infestation, %** | 28.20 ± 4.9 | **12.00 ± 1.38 -57.4%** | **5.2 ± 1.3 -81.6%** | 18.3 ± 2.6 -35.1% |
| **Honey quantity, kg** | 21.0 ± 0.8 | 24.0 ± 0.5 +14.3% | 24.7± 0.5 +17.6% | 19.10 ±0.96 -10.0% |

The results of this test campaign in the hives confirm the effects of the complexes measured in previous campaigns, particularly in protecting bees against *Varroa* infestation.

With an overall dose of 2mg of **Li-$Mo_2O_4$-EDTA** per hive, the rate of *Varroa* infestation of worker bees fell by 47%, in line with last year's figures of 43%. This effect is increased to -61.3% when the dose of complex is multiplied by 3. On the other hand, lithium acetate, used as a reference, also showed a significant reduction in the infestation rate to -30.4% by using 2mg per hive. This rate is lower than those obtained with the **Li-$Mo_2O_4$-EDTA** complex. Furthermore, the Lithium concentration in the LiOAc group is 3.68 times higher than in the **Li-$Mo_2O_4$-EDTA**-2mg batch and 1.23 times lower than in the **Li-$Mo_2O_4$-EDTA**-6mg group. In other words, the effects of LiOAc are lower than those measured for **Li-$Mo_2O_4$-EDTA**-2mg, despite the higher $Li^+$ content, suggesting that the complex specifically has a protective action against Varroa, in synergy, where possible, with the $Li^+$ cations. Besides, the effect of LiOAc appears negative or negligible on the development of the colonies and on the production of honey.

Interestingly, the rate of Varroa infestation on the brood that was measured during this test campaignwas very high, *i.e.* 28.20% in the control batch. After feeding with the reference compound LiOAc (2mg/hive), this rate dropped by 35.1% compared to the control group, demonstrating the action of lithium cations. The **Li-$Mo_2O_4$-EDTA** complex was still much better, reducing Varroa infestation on brood by -57.4% with 2 mg and up to -81.6% with 6 mg of **Li-$Mo_2O_4$-EDTA** per hive.

Beyond this result, it also indicates that the complex introduced into the feeding syrup is used not only to feed the bees already present, but also the larvae and unborn bees, with a probable effect on several generations of bees.

#### IV.2- Test campaign in France (2019)

Test campaigns in beehives are highly dependent on the colonies, their environment, and climatic conditions. It is also reasonable to assume that they may also depend on feeding protocols and bee species. In previous campaigns, the complexes are given to the bees every two days during a period of two weeks. This protocol can be easily applied in academic apiaries or for non-professional beekeepers. Therefore, a first question arises on the efficiency of our complexes in classical professional conditions used in beekeeping in Europe.

To assess this point, a test campaign was carried out in France on experimental apiary, using *Apis mellifera "buckfast"* bees widely used by professional beekeepers in real operating conditions, in the Paris region, at Gif-sur-Yvette (department number 91, see Figure SI.1 in part 1 of the SI) on 22 hives. The feeding conditions are those used by professional beekeepers, and in accordance with French regulations, the honey produced by the test groups (if so) is not sold but left to the bees or destroyed. The complex used was **Li-$Mo_2O_4$-EDTA**.

##### V.2.1 Test campaign in Gif-sur-Yvette. France. 2019

For this test campaign, 22 hives were selected and split into two batches: 11 control hives and 11 test hives. In early spring, 0.5 L of sugar syrup (50% sucrose. w/w) was introduced into each colony on April 1. All the colonies are controlled to be equivalent at the start of the experiment, and honey supers are added when necessary for honey production. In spring and at the end of the experiment, the quantities of honey produced is measured in the beehive's honey supers only by weighting them and comparing them with the mass of those that are empty and whose frames are already built (5.5±0.1 kg in average). The masses given in the next tables take into account the additions of this honey supers to keep a value corresponding to the initial conditions: body of the beehive + colony of bees + ressources.

For the test hives, 2 mg of **Li-$Mo_2O_4$-EDTA** complex were introduced in only one time into the syrup for each hive of the test group to match the total amount of complex given in hives in Moldavia. For these colonies placed in real operating conditions, little intervention was carried out. We monitored colony mass as a function of time and the quantity of honey produced on July 15. The results are gathered in Table SIV.5. while the details of the following for each hive is given in Tables SIV.23 and SIV.24 and the comparison of mean colony masses for the control and **Li-$Mo_2O_4$-EDTA** groups during the experiment is given in Figure SIV.7.

***Table SIV.5***. *Results (mean ± se) obtained for the Gif-sur-Yvette Campaign (2019)*

| | **Number of hives** | **Initial colony weight[a] (1st April). kg** | **Final colony weight (colony+ressources).[b] 15th july. kg Variation. %** | **Honey production/hive. spring[c]** | **Total Honey production/hive (15th july) spring+summer[c]** |
|---|---|---|---|---|---|
| **Control Group** | 11 | 24.41±1.00 | 54.16±3.98 (+29.75 kg. +122%) | 4.36±1.03 | 19.13±3.73 |
| **Li-$Mo_2O_4$-EDTA** 2mg / 0.5 L syrup | 11 | 25.07±0.63 | 66.94±3.91 (+41.87 kg. +167%) | 4.21±0.56 | 30.33±3.43 |
| **Variation *vs* control. %** | - | +2.7% | +23.6% | -3.4% | **+58.7%** |

a. Initial colony weight of the hives, which consist only of a body of hive at the start of the experiment; [b]. Mass of the hive at the end of the experiment. This mass takes into account the mass of the added honey supers (5.50 kg) to consider only the body of the hives, the weight of the colony and its ressources (honey. beebread); [c]. This mass is estimated by the mass of the honey supers considering that an empty honey supers has a mass of 5.5 kg.

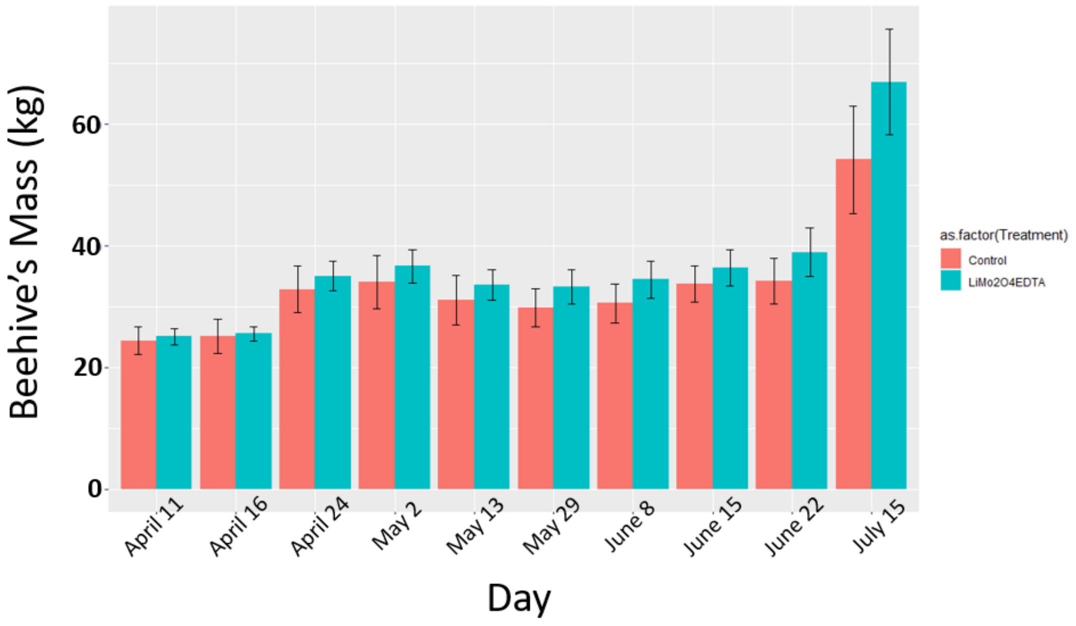


***Figure SIV.7*** *Comparison of mean colony weights for the control and Li groups during the experiment.*

In this experiment carried out under real beekeeping conditions, we found that just 2mg of **Li-$Mo_2O_4$-EDTA** complex per hive introduced in a single dose in 0.5L of syrup on April 1 resulted in a +23.6% increase in hive weight on July 15 compared with the reference batch, and a very significant +58.7% increase in the quantity of honey produced (30.3 kg on average for the test batch versus 19.1 kg for the control hives on July 15). This result can be compared with the production increase observed during the 2018 campaign in Moldavia with 2 mg of the **Li-$Mo_2O_4$-EDTA** complex (+47%) administered over a two-week period. However, as shown in Figure SIV.7 and Tables SIV.23 and SIV.24. the differences in colony weight and honey production between the two groups are very limited for one and a half months and start to become significant two months after start-up. This might suggest that either the syrup is consumed over several weeks by the colonies or the impact of the product is greater on eggs, brood and young bees to explain such a delay or both. In fact, it takes 21 days from egg to young bee and a further month from young bee to forager bee. Further experiments would be needed to confirm this hypothesis.

Nevertheless, this experiment validates the use of our complexes in a limited number of feedings. **Our complexes used as food complement in syrup are thus perfectly compatible with professional beekeeping**.

#### IV.3- Tests in California. USA : winters 2019 and 2020

In previous test campaigns, we were able to measure significant effects on colony development and protection. The autumn-winter period, known as the wintering period, is a very important period in the life cycle of colonies. During this period, the population is reduced and colony activity is minimal. It is during this period that colony losses can be very significant.

We therefore felt it essential to evaluate the effects of our complexes on winter mortality. The tests were carried out on hives located in the San Francisco area. Two test campaigns have been carried out for winter 2019-2020 and 2020-2021.

##### IV.3.1 Wintering campaign 2019-2020

For this first campaign, 151 beehives were chosen, spread over 6 different apiaries in California - south of San Francisco - either in hill or in valley environment. The bees were of different types: Wildflower Meadows ™ (Italian VSH), Beeweaver. Californian Carniolan bee (Pope Canyon), Californian Tom bee and Californian Italian bee (Sam).

The hives are Langstroth 2-body type (15 frames vs. 10 for Dadant hives in Europe). On each apiary, two populations of hives are randomly divided into those serving as controls and those receiving the **Li-$Mo_2O_4$-EDTA** complex (76 controls, 75 test hives).

Between October 25 and 28 2019, all the colonies are fed with 1 US Gallon (3.78 liters) of 65% sugar syrup introduced into the hive (see Figure SIV.8. right). For the test hives, the **Li-$Mo_2O_4$-EDTA** product is syringed at the same time as a concentrated aqueous solution (8 g/L. see Figure SIV.8. left) in sugar syrup into the frame feeder. 0.5 mL of this solution is introduced and dispersed. Each test hive thus receives a single overall dose of 4 mg **Li-$Mo_2O_4$-EDTA**. Control hives receive only the sugar syrup.

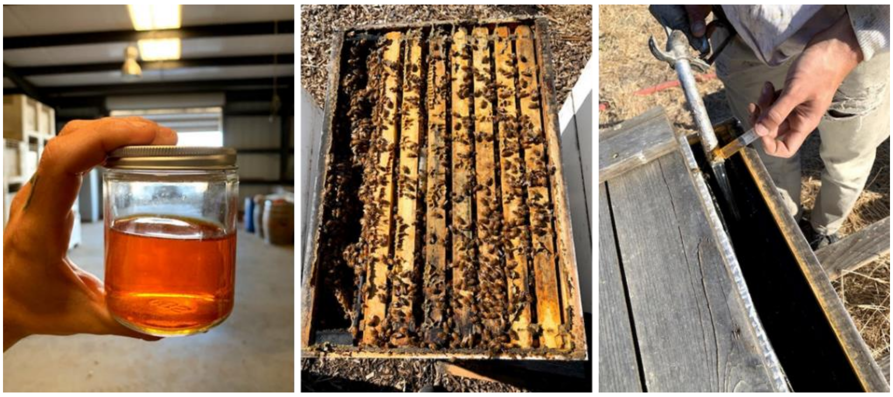

***Figure SIV.8***. *Aqueous solution of **Li-$Mo_2O_4$-EDTA** at 8 g/L ; top view of a colony ; view of the introduction of the complex with the syrup in test colonies.*

The surviving colonies are counted on January 7th 2020. A total of 73 colonies out of the initial 151 died, giving an overall mortality of 48.3%. Interestingly, the mortality in Hill apiaries is 55% in average *vs* 44% in Valley. The mortality also differs from the bee types. For Californian Carniolan bee (Pope Canyon) and Californian Tom bee types, the mortality rate is 81.8% (27/33) and 87.5% (7/8) respectively. while it is lower for Californian Italian bee (Sam). i.e. 65.38% (17/26) and much lower with the Wildflower (Italian VSH). Beeweaver type bees: 26.67% (12/45) and 25.64% (10/39) respectively.

More in detail, for the control group, 47 colonies were lost out of 76. *i.e*. 61.8% winter mortality. which is a classic result in this region. In contrast, the mortality in test hives treated with 4 mg **Li-$Mo_2O_4$-EDTA** amounted to 26 colonies lost out of the initial 75, *i.e.* 34.7% winter mortality, a drop -43.8% compared with control hives. The results are summarized in Table SIV.6.

***Table SIV.6****. Results of the wintering feeding on the mortality in California. USA. 2019-2020*

| Group | Feeding | Number of hive | Colonies lost 7$^{th}$ January 2020 | %Colonies lost |
|---|---|---|---|---|
| Control | 25-28$^{th}$ October 2019 | 76 | 47 | 61.8% |
| **Li-$Mo_2O_4$-EDTA. 4 mg** | | 75 | 26 | 34.7% |

**IV.3.2 Wintering campaign 2020-2021**

For this second campaign, 220 beehives were involved, spread over 11 different apiaries (from 8 to 24 beehives par apiary) in California, south of San Francisco either in hill or in valley environments. Taking into account the previous results, the bees were of Wildflower (Italian VSH) or Beeweaver types only.

The hives are Langstroth 2-body type (15 frames vs. 10 for Dadant hives in Europe). Each apiary is randomly divided into 4 batches of hives. for a total of:

- **Group 1:** 55 control hives fed only with sugar syrup on September 20 2020 and on October 12. 2020
- **Group 2:** 55 hives receiving **Li-$Mo_2O_4$-EDTA** on September 20 2020 (8mg in 1 US Gallon) and suger syrup on October 12. 2020
- **Group 3**: 55 hives receiving sugar syrup on September 20 2020 and **Li-$Mo_2O_4$-EDTA** (8mg in 1 US Gallon) and suger syrup on October 12 2020
- **Group 4:** 55 hives receiving **Li-$Mo_2O_4$-EDTA** on September 20 2020 and on October 12 2020. (4mg in 1 US Gallon each time)

1 apiary contains only Wildflower bees. 2 apiaries contain only Beeweaver bees. The remaining 10 apiaries contain both Wildflower and Beeweaver bees, equally divided between the 4 batches.

The **Li-$Mo_2O_4$-EDTA** product is syringed into the frame feeder between September 20. 2020 and/or October 12. 2020 as a concentrated aqueous solution (0.8 g/L) in sugar syrup. 10 mL of this solution are introduced and dispersed in 1 US Galon of 65% sugar syrup introduced into the hive. Each test hive thus receives an overall dose of 8 mg **Li-$Mo_2O_4$-EDTA** for wintering. Control hives receive only the sugar syrup.

Dead colonies are counted on December 31. 2020 (see Table SIV.7). A total of 28 colonies had died by 12/31/2020, i.e. 12.73% of the total. The mortality rate is identical for Wildflower (21/165. 12.72%) and Beeweaver (7/55. 12.72%) bees. Mortality by batch was as follows:

- Group 1: 15 hives / 55 hives. i.e. 27.3% losses
- Group 2: 0 hives / 55 hives. i.e. 0% losses
- Group 3: 10 hives / 55 hives. i.e. 18.2% losses
- Group 4: 3 hives / 55 hives. 5.5% loss.

***Table SIV.7***. *Results of the wintering feeding on the mortality in California. USA. 2019-2020*

| Group | Feeding | | Number of hive | Colonies lost 31th December 2020 | %Colonies lost (variation *vs* control group) |
|---|---|---|---|---|---|
| | September 2020 | October 2020 | | | |
| Control. Group 1 | Sugar syrup only | Sugar syrup only | 55 | 15 | 27.3% |
| **Group 2** | **Li-$Mo_2O_4$-EDTA** 8mg in sugar syrup | Sugar syrup only | 55 | 0 | 0% (-100%) |
| **Group 3** | Sugar syrup only | **Li-$Mo_2O_4$-EDTA** 8mg in sugar syrup | 55 | 10 | 18.2% (-33.3%) |
| **Group 4** | **Li-$Mo_2O_4$-EDTA** 4mg in sugar syrup | **Li-$Mo_2O_4$-EDTA** 4mg in sugar syrup | 55 | 3 | 5.5% (-79.9%) |

The results show that feeding in October, as in the previous year's campaign, reduces mortality by 33% compared with the control batch. This compares with -44% the previous year. The dose effect, 4 or 8 mg/hive, was not significant.

On the other hand. **earlier feeding when the queen begins to produce eggs to form winter bees gives exceptional results**. Feeding in September (group 2) or September + October (group 4) greatly reduces colony mortality. No colonies in group 2 were lost out of 55 hives. and only 3 colonies out of 55 were lost with group 4, **representing a reduction in winter mortality of around 80% compared with the control batch**.

#### IV.4- Test campaign in Greece: 2020-2021.

##### IV.4.1 Introduction / objectives of the study

In previous campaigns, colonies received quantities ranging from a few mg to 30 mg of Mo-complexes over several weeks/months. Obviously, the question arises as to the maximum quantities that can be used. Is there a limit beyond which doses become deleterious? The aim of this test campaign, carried out over a wintering period in Greece, is to answer this question.

For this campaign, we focused on the **Li-$Mo_2O_4$-EDTA** complex, for which we know that lithium is not an innocent cation, unlike the sodium cation.

The experiment was conducted in collaboration with the Department of Apiculture- ELGO 'DIMITRA', in Nea Moudania, Greece (see Figure SIV.9).

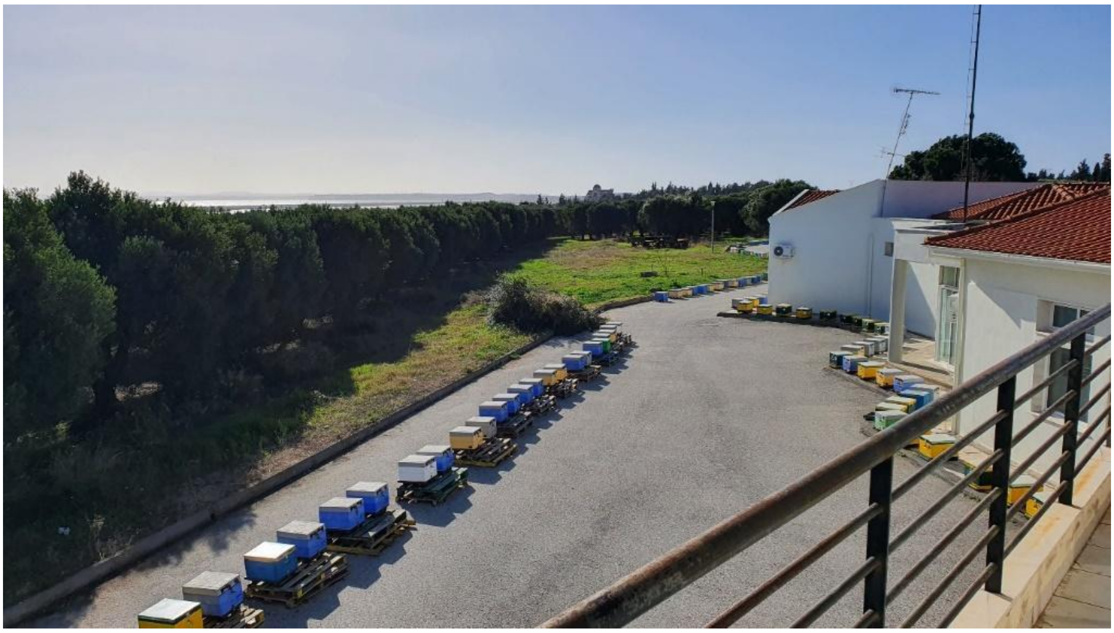

***Figure SIV.9.*** *Apiary of the Department of Apiculture, Nea Moudania, Greece.*

The aim of this pilot study was to assess the performance of the honey bee colonies after the addition of the product **Li-$Mo_2O_4$-EDTA** in their food supply. The specific objectives of the study were:

- To determine the impact of feeding with **Li-$Mo_2O_4$-EDTA** on the development of colonies, during the winter and early spring (also overwintering ability)
- To examine the impact of **Li-$Mo_2O_4$-EDTA** on the mite *Varroa destructor* load after the end of the two periods on the different groups.
- To examine the impact of **Li-$Mo_2O_4$-EDTA** on the fungus *Vairimorpha spp.* load after the end of the two periods on the different groups.

##### IV.4.2 Experimental design

The experimental design was split in two parts. Thirty (30) beehives were used in total during the period from December 2020 until April 2021 as follows:

**1st part- winter 2020-2021.** During the period December 2020 till march 2021, 20 beehives (called Group "MoLi") received candy supplemented with the product **Li-$Mo_2O_4$-EDTA,** while 10 beehives were used as a control group and they were supplied with plain candy. The 20 treated colonies received about 4 Kg of "sugar candy" each containing 10 mg of **Li-$Mo_2O_4$-EDTA** per kg of candy (therefore 40 mg of **Li-$Mo_2O_4$-EDTA** in total). Note that during this period 1 colony from the control group was lost, decreasing the number of hives of this group to 9.

**2nd part- spring 2021.** In a second period, from March 2021 to April 2021, the "treated colonies" from the group "MoLi" from the 1st part were divided in two groups : 10 beehives were supplied with **Li-$Mo_2O_4$-EDTA** in a sugar syrup (Group MoLi-A), while the other 10 and the 10 initial control beehives received only sugar syrup (Group MoLi-B). The **Li-$Mo_2O_4$-EDTA** product was supplied to the colonies at a concentration of 10 mg / liter of syrup and a total of 4 liters was given to the colonies.

Therefore, one group of beehives received 40 + 40 = 80 mg of **Li-$Mo_2O_4$-EDTA** from December 2020 till March/April 2021 (Group "MoLi-A"); one group of beehives received 40 mg of **Li-$Mo_2O_4$-EDTA** from December 2020 till March/April 2021 (group "MoLi-B") and one group received only sugar (group control). The scheme SIV.1 summarizes these three groups of hives.

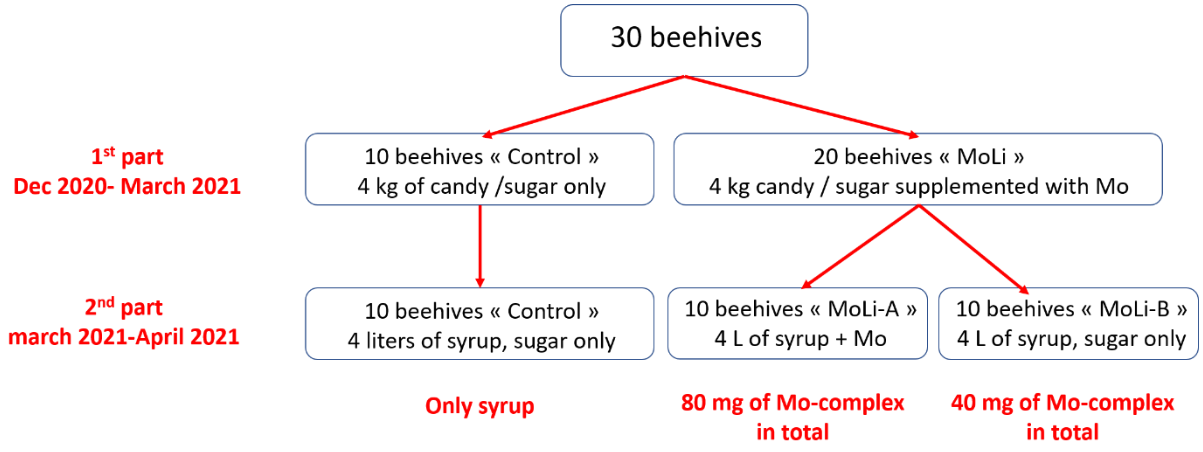


*Scheme SIV.1. Schematic view of the three groups involved in the test campaign in Greece.*

##### IV.4.3 Description of the protocol

In order to assess the development (also strength) of the colonies and their overwintering ability, a measurement of the population and the brood was necessary. Both traits were assessed based on a well-defined protocol [6, 7]. The protocol enables the user to follow the development of the colonies, to detect any reduction in population and count the dead bees (collected in front of the hives in traps) and brood and accurately assess the dynamics of the colonies, also showing indirectly the health of the colonies. Brood area (number of brood cells) and population (number of adult bees) were assessed three times during the experiment: just before the experiment started, at the end of the winter, and at the end of the spring (end of experiment).

At the same time, the direct assessment of the health of the colonies was performed by sampling and measuring the presence/ infestation levels of two main parasites: *Varroa destructor* mite and *Vairimorpha spp mirosporidium* (previous *Nosema* spp) based on international protocols, also used in COLOSS association *Varroa* Task Force.

In particular, the number of dead *Varroa* mites were monitored at the bottom of the bee colonies, continuously and till after the feeding of the colonies stopped in April 2021. After this monitoring, a 'critical' treatment occurred to ensure an accurate and efficient removal of the mites, using oxalic acid by trickling application. The efficacy of the **Li-$Mo_2O_4$-EDTA** treatment was assessed by the formula E= Number of mites before the critical treatment X 100/ (Number of dead mites after the critical treatment + Number of mites before the critical treatment).

*Vairimorpha* infestation on adult bees was assessed based on laboratory method suggested by the Office international des épizooties (OIE) [8], using 30-60 adult bees from the outer frames of the colonies. The outcome is expressed as the number of *Vairimorpha* spores/ bee.

##### IV.4.4 Raw data

All the raw data obtained for brood cells, bee population, dead bees, Varroa and Vairimorpha counts for the two periods of the experiments are listed in tables SIV.25 to SIV.32. The results are discussed in paragraph IV.4.5.

##### IV.4.5 Results and discussion

###### 1°) Colony dynamics

The evolution of the number of brood cells during the experiment is given in figure SIV.10. At the beginning of the experiment the number of brood cells is 0 for the hives of the control group and 1015 in average for the other hives belonging to the group “MoLi”. In March 2021, these numbers increased to 16363 for control hives and 12810 in average for the hives of the MoLi group (20 beehives).

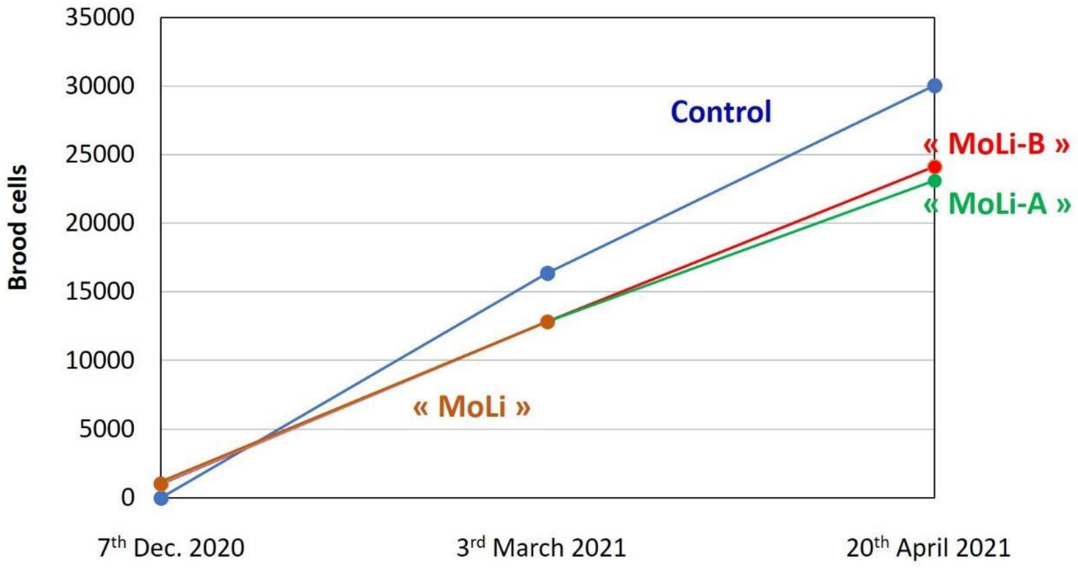


***Figure SIV.10.*** *Number of brood cells at the starting of the experiment, at the end of winter (March) and at the end of the experiment (April) for the control group and the "MoLi", "MoLi-A" and "MoLi-B" groups.*

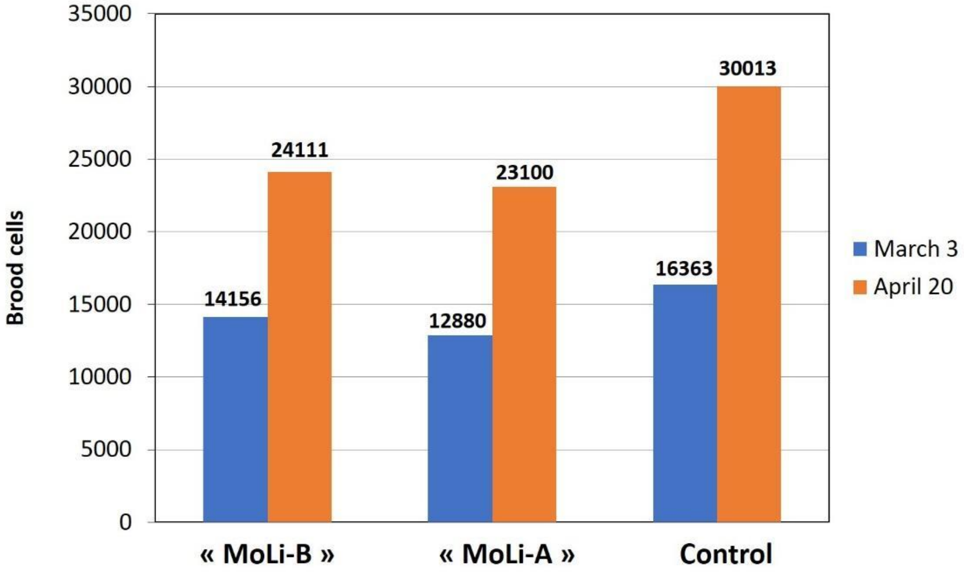


***Figure SIV.11.*** *Number of brood cells at the end of winter march and at the end of the experiment for the control, "MoLi-A", and "MoLi-B" groups.*

During the second phase of the experiment (see Figures SIV.10 and SIV.11), the differences between the three groups control, MoLi-A and MoLi-B increases again to reach respectively 30013, 23100 and 24111 brood cells per hive in average (not significant).

This result suggests that the winter treatment with 40 mg of complex **Li-$Mo_2O_4$-EDTA** in candy bags (MoLi group) do not favor the development of brood cells in the colonies. Besides, the effect of continuation of feeding with syrup supplemented with **Li-$Mo_2O_4$-EDTA** in March-April (group MoLi-A) seems to increase again the difference with the control group but the difference with the MoLi-B is not so significant in March (see Figure SIV.11).

The same pattern appears also on the population of bees (number of adult bees, see Figure SIV.12), although the difference is not significant (at least till the time of the last measurement). However, it is important to note that the amount of brood in one colony is represented as number of adult bees 21 days later.

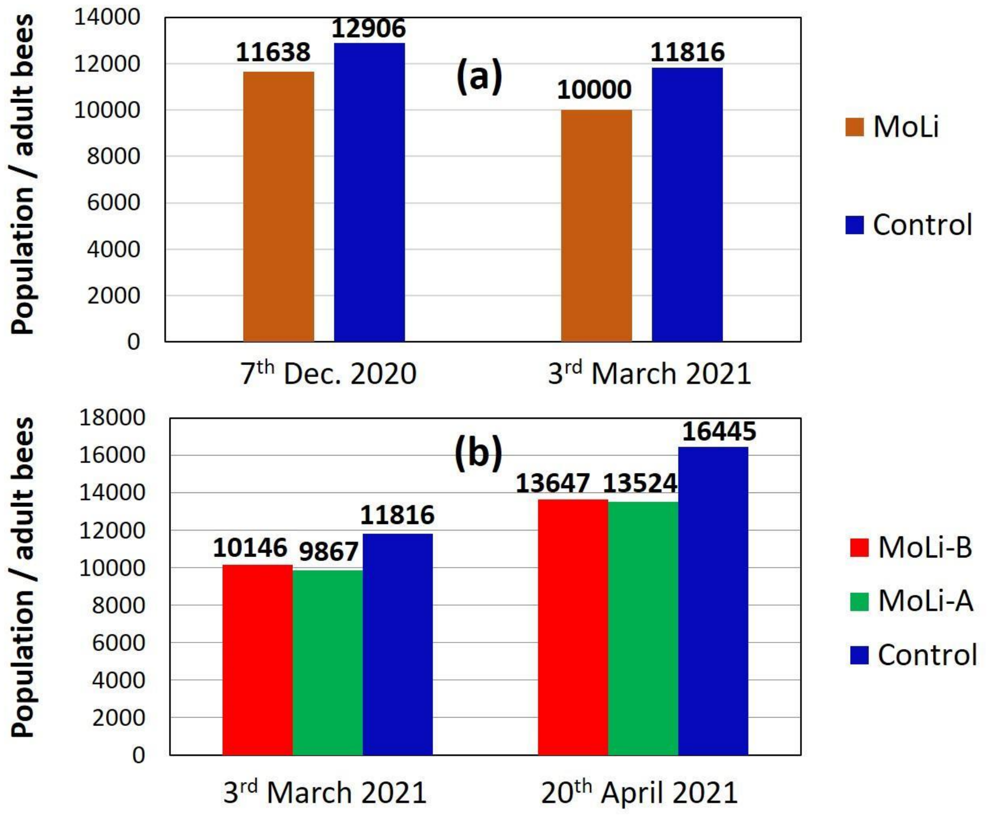


***Figure SIV.12.*** *Adult bees population of the colonies during the course of the experiment for the control, MoLi, MoLi-A and MoLi-B groups.*

**2°) Toxicity indications**

In this pilot study, we did not look into differences in detoxification enzyme profile or other mechanisms showing intoxication of the bees (e.g. heat shock proteins). We only observe the number of dead bees at the entrance of the colonies (using simple traps). The variations of dead bees for the different groups during the two periods of the experiments are given in Figures SIV.13a and SIV.13b, respectively.

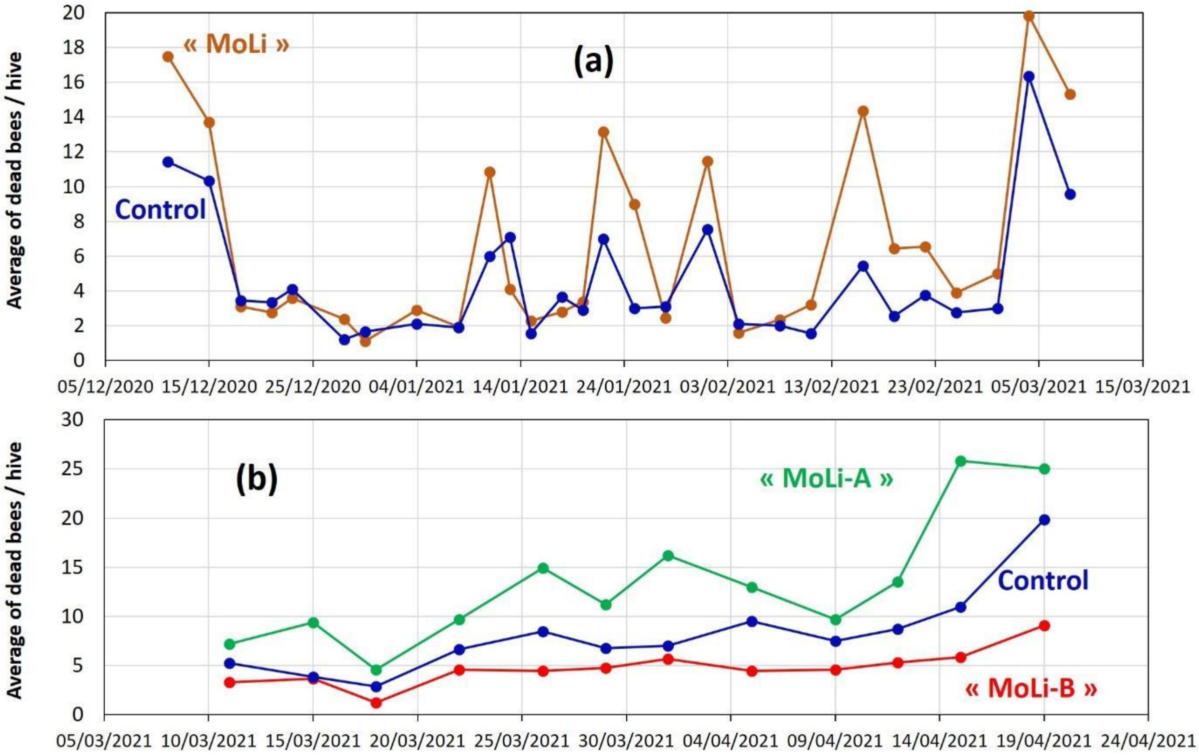


***Figure SIV.13.*** *(a) Dead bees in front of the colonies, while the colonies were fed with sugar candy (winter period) for the control and the MoLi groups; (b) Dead bees in front of the colonies while the colonies were fed with sugar syrup (spring period) for the control and the MoLi-A (the group fed with supplemented candy and supplemented syrup in green) and the MoLi-B (group fed with supplemented candy and pure syrup in spring period in red) groups. The number of dead bees is given as an average value per hive to take into account the number of hives, which differs in each group (9 or 8 for control, 20 for MoLi, 9 for MoLi-B, 10 for MoLi-A)*

Interestingly, as shown in Figure SIV.13a, the number of dead bees per hive appears significantly higher on the colonies fed with **Li-$Mo_2O_4$-EDTA** as a food supplement during the first period of the experiment (P=0.001). The above difference with the control group was increasing while the feeding continued in the spring period (MoLi-A group). Conversely, we can note that in the group fed **Li-$Mo_2O_4$-EDTA** during the winter but plain syrup during spring (MoLi-B group), the dead bees immediately stop increasing and the evolution of dead bees per hive during the second period is significantly lower than the control group but not significantly lower. However, the group continue to be fed with **Li-$Mo_2O_4$-EDTA** (Moli-A group) continue to show significantly higher number of dead bees compared to control and Group Moli-B (P=0.001).

Feeding bee colonies with high quantity of complex **Li-$Mo_2O_4$-EDTA** during long periods has deleterious effects on the colonies, even if the increase of mortality remains relatively low. This result contrasts with the chronic and acute toxicity studies (see part III of the supporting information), which evidenced no toxicity of this complex on bees in laboratory conditions, even at high concentration. It suggests either that the process is more complex in beehives or that a shock administration might be better tolerated by the bees than a long exposure in natural conditions in hives.

##### 3°) *Varroa* infestation levels

First, it is important to note that the colonies involved in the experiment did not have a heavy *Varroa* load and the number of fallen mites in the control colonies was low, which does not favour the demonstration of a significant effect.

The figure SIV.14 shows the average number of *Varroa* fallen per hive during the winter period. At the starting of the experiment, the number of Varroa appears higher in hives receiving the Mo complex in candy. After one week, the curve showing the number of *Varroa* fallen per hive become almost systematically lower than for the control group, which indicates a lower varroa load for hives fed with sugar supplemented by the complex **Li-$Mo_2O_4$-EDTA**, in agreement with previous studies in hives, notably in Moldova.

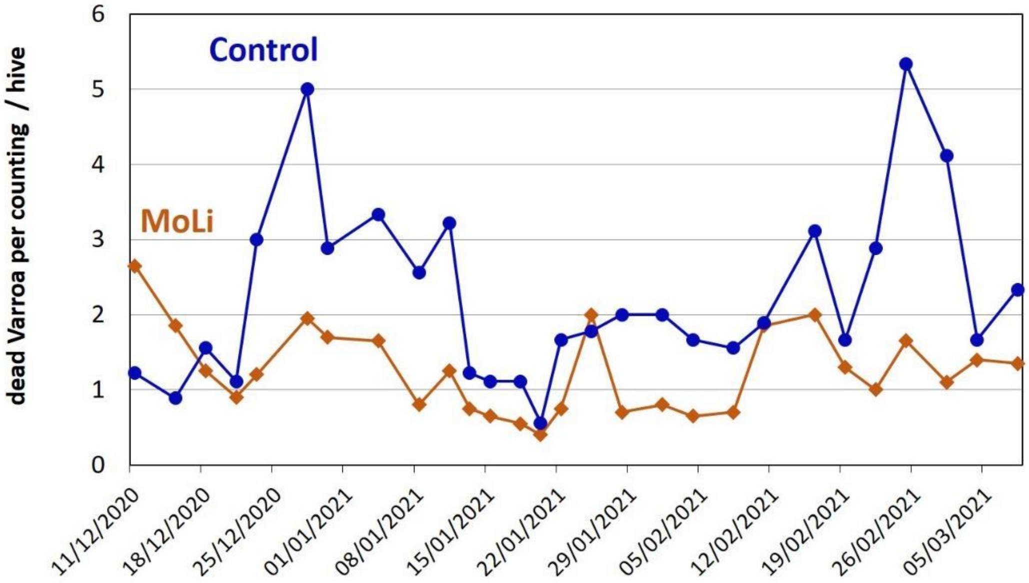


***Figure SIV.14.*** *Variation of the average number of dead Varroa mites per hive at the bottom of the colonies during the winter treatment for the control group (9 hives) and the group MoLi (20 hives).*

The figure SIV.15 shows the variation of the average number of varroa fallen per hive during the spring period. During this period, colonies are fed with syrup (4 L). Control and MoLi-B groups received only syrup with sugar, while the MoLi-A group received syrup containing the complex **Li-$Mo_2O_4$-EDTA** at 10 mg/L (40 mg in total). The number of Varroa fallen is clearly lower for the group MoLi-B even after the critical treatment by oxalic acid occurred in April, which demonstrates a lower *Varroa* load for this group in general.

Concerning the group MoLi-A, the average number of varroa fallen is intermediate between the control group and the group MoLi-B. In a previous paragraph, we evidenced that the spring feeding with 40 mg of **Li-$Mo_2O_4$-EDTA** becomes deleterious for bees. In terms of *Varroa* load, this additional treatment does not appear to have any effect.

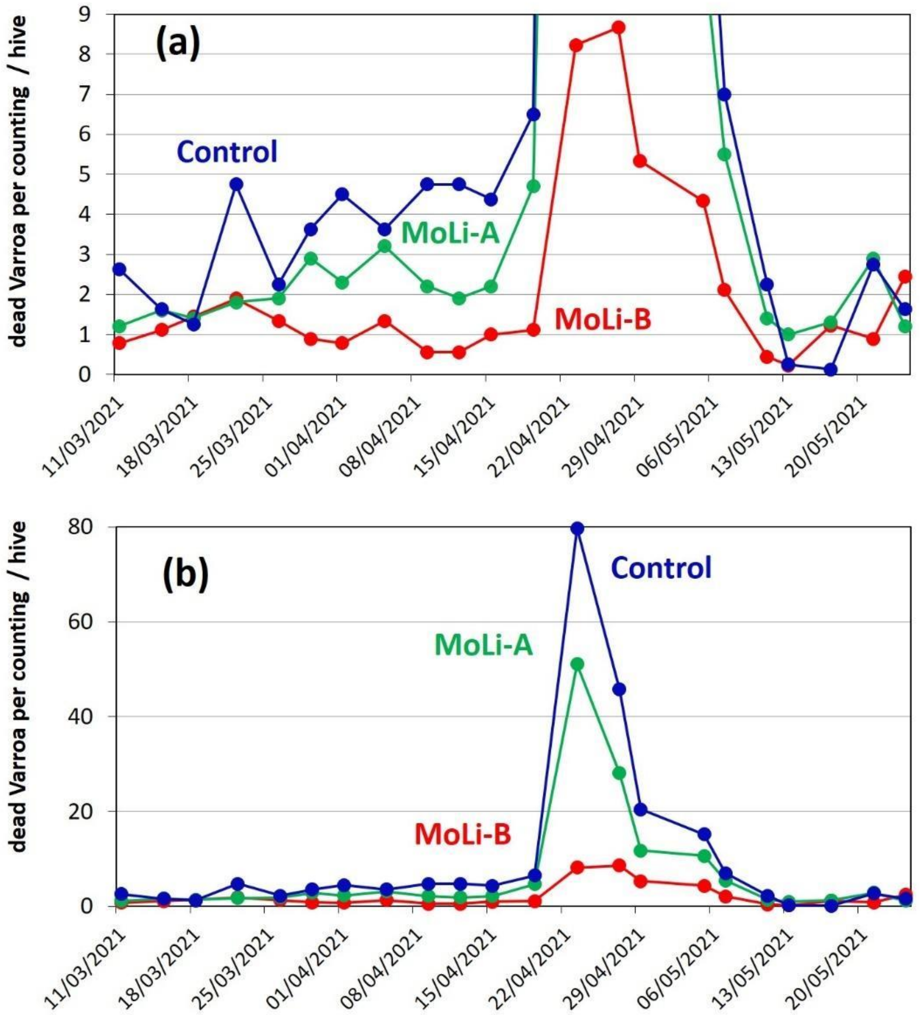


***Figure SIV.15.*** *Variation of the average number of dead Varroa mites per hive at the bottom of the colonies during the spring treatment with syrup for the control group (8 hives) and the group MoLi-B (9 hives), or syrup plus Li-$Mo_2O_4$-EDTA for the group MoLi-A (10 hives).*

Finally, the efficacy of the treatments can be estimated by E = (Number of mites before the critical treatment X 100) / (Number of dead mites after the critical treatment + Number of mites before the critical treatment). The Table SIV.8 gathers the values calculated for the three groups. The efficacy appears to be significantly higher for the group MoLi-B compared to the control group and the MoLi-A group (P=0.03); however, MoLi-B group appears to have lower *Varroa* loads in general.

***Table SIV.8****. Efficacy parameter against Varroa load*

| Group | Number of mites before the critical treatment | Number of dead mites after the critical treatment | Efficacy E of the treatment (%) |
|---|---|---|---|
| Control | 357 | 1402 | 20.3 |
| MoLi-A | 273 | 1150 | 19.2 |
| MoLi-B | 115 | 305 | **27.4** |

##### 4°) *Vairimorpha* infestation level

The figure SIV.16 represents the average number of *Vairimorpha* spores per bee at the starting of the experiment, at the end of winter and at the end of the experiment. At the beginning of the experiment, the number of spores is similar in the two groups Control and MoLi. Furthermore, as seen in Table SIV.32, the hive n°19 of the control group displays an enormous quantity of Nosema spores (6 millions). Considering the low number of hives, this hive alone distorts the averages calculated for this group, and it is considered as an extreme value. This hive is not considered in the Figure SIV.16, or in the analysis

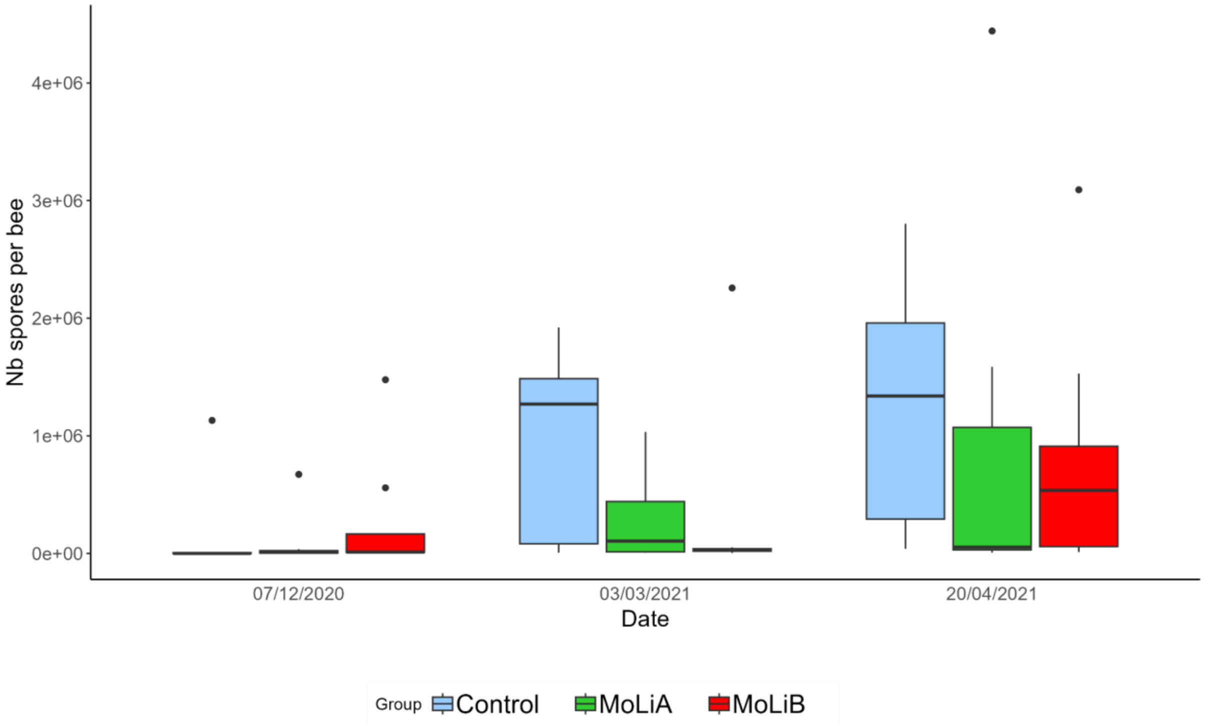


***Figure SIV.16.*** *Number of Vairimorpha spores per bee, as monitored before and after the winter feeding of the colonies with sugar candy; and after the spring feeding of the colonies with syrup.*

During winter period (Figure SIV.16 left and middle), feeding the colonies with Li-$Mo_2O_4$-EDTA (group MoLi) seems to keep the infestation of *Vairimorpha* microsporidium low (difference with the control colonies was found not significant with P=0.54). Later in the season, the Li-$Mo_2O_4$-EDTA groups also start increasing on *Vairimorpha* numbers and the difference with the control group was again not significant (P =0.7, Fig. SIV.16 right).

#### Conclusions of the test campaign in Greece

From this short-term experiment, has become obvious that the complex Li-$Mo_2O_4$-EDTA product, administered to honey bee colonies in a concentration of 10 mg in 1 kg of candy sugar during the winter period (40 mg in total), might have some potentials against Varroa and *Vairimorpha*, even during the spring period in case of *Varroa*.

However, continuing to feed the colonies with this syrup complex at a dose of 10 mg/L (40 mg in total) has apparent adverse effects on adult bees, whose mortality rate increases (we did not distinguish whether the dead bees were foragers or nursing bees). A lower concentration could give better results and have no harmful effects, as we have shown in other test campaigns in Moldova, France and the USA. It is possible that the product Li-$Mo_2O_4$-EDTA and in particular the lithium cations, accumulate in the bees (in accordance with the studies presented in part VI of the SI) to explain these deleterious effects observed with a feeding of 80 mg of complex per hive.

#### IV.5 General conclusions of the tests performed in hives.

In this part, we presented the test campaigns performed in Moldavia, in France, in the USA and in Greece. The scale of these tests is unprecedented. They were carried out on different bee species, in different environments, under different experimental conditions, at different times of the year and in 4 different countries. The results obtained in these different campaigns agree with each other.

The first step was to identify the most effective complexes in the hives: **Na-$Mo_2O_4$-EDTA** and **Li-$Mo_2O_4$-EDTA**

For these two complexes, we have highlighted:

- Greater colony growth (Moldavia).
- Better protection of brood and bees against Varroa destructor, both as syrup form (Moldavia, France) and as candy form (Greece), particularly for the complex **Li-$Mo_2O_4$-EDTA**.
- A possible effect against *Vairimorpha* in hives treated with supplemented candy (Greece).
- A -significant increase in the production of wax and honey (Moldova, France) without altering its quality and without any residue of the complexes.
- A significant reduction in colony mortality during the winter (USA).
- An estimation of the optimal doses of complexes, with evidence of deleterious doses for complex **Li-$Mo_2O_4$-EDTA** (Greece).

#### IV.6 Raw data.

***Table SIV.9*** *SDetails of the main parameters monitored for the* ***control group*** *during the* ***2013 test campaign****.*

| sSample | Hive numbering | Colony strenght | | Capped brood quantity, hundreds of cells | Queen bee's prolificity, eggs/24h | Beebread quantity, hundreds of cells | Honey quantity | | Wax | | Hygienic behaviour, % |
|---|---|---|---|---|---|---|---|---|---|---|---|
| | | Intervale | kg | | | | Number of frames | kg | number fabricated honeycomb | kg | |
| **1** | 2 | 12.5 | 3.13 | 190 | 1583 | 95 | 7.0 | 11.5 | 2.3 | 0.28 | 90.0 |
| **2** | 3 | 13.2 | 3.30 | 200 | 1667 | 100 | 7.5 | 13.5 | 2.7 | 0.32 | 90.0 |
| **3** | 8 | 12.5 | 3.13 | 195 | 1675 | 80 | 8.0 | 10.5 | 2.2 | 0.26 | 87.5 |
| **4** | 9 | 12.6 | 3.15 | 180 | 1500 | 85 | 7.0 | 10.0 | 2.3 | 0.28 | 87.5 |
| **5** | 10 | 13.0 | 3.25 | 205 | 1708 | 95 | 7.5 | 12.8 | 2.5 | 0.30 | 87.5 |
| **6** | 11 | 12.4 | 3.10 | 180 | 1500 | 97 | 7.2 | 10.5 | 2.0 | 0.24 | 90.0 |
| **7** | 12 | 12.8 | 3.20 | 190 | 1583 | 78 | 8.0 | 9.5 | 2.2 | 0.26 | 86.5 |
| **8** | 13 | 13.0 | 3.25 | 195 | 1625 | 90 | 7.0 | 13.5 | 2.5 | 0.30 | 87.5 |
| **9** | 14 | 13.2 | 3.30 | 190 | 1583 | 100 | 7.2 | 13.0 | 2.6 | 0.31 | 87.5 |
| **10** | 15 | 13.5 | 3.38 | 190 | 1583 | 90 | 6.5 | 13.4 | 2.4 | 0.29 | 89.0 |
| **11** | 17 | 12.4 | 3.10 | 185 | 1542 | 93 | 7.0 | 11.0 | 2.0 | 0.24 | 90.5 |
| **12** | 42 | 12.5 | 3.13 | 195 | 1625 | 100 | 8.0 | 13.5 | 2.7 | 0.32 | 88.5 |
| **13** | 21 | 12.5 | 3.13 | 193 | 1608 | 84 | 7.5 | 10.5 | 2.2 | 0.26 | 89.5 |
| **14** | 24 | 12.6 | 3.15 | 180 | 1500 | 92 | 7.2 | 10.5 | 2.5 | 0.30 | 88.0 |
| **15** | 1 | 13.0 | 3.25 | 210 | 1750 | 80 | 6.8 | 13.2 | 2.5 | 0.30 | 91.5 |
| **16** | 25 | 13.0 | 3.25 | 175 | 1458 | 88 | 6.5 | 9.0 | 2.0 | 0.24 | 84.0 |
| **M±m** | | | **3.20±0.02** | **190.80±2.40** | **1590.00±20.00** | **90.50±1.80** | | **11.62±0.40** | | **0.28±0.08** | **88.40±0.40** |
| **σ** | | | **0.08** | **9.50** | **79.00** | **7.40** | | **1.61** | | **0.03** | **1.80** |
| **Min** | | | **3.10** | **175.00** | **1458.00** | **78.00** | | **9.00** | | **0.24** | **84.00** |
| **Max** | | | **3.38** | **210.00** | **1750.00** | **100.00** | | **13.50** | | **0.32** | **91.50** |
| **m=σ/sqrt(16)** | | | **0.02** | **2.38** | **19.75** | **1.85** | | **0.40** | | **0.01** | **0.45** |

***Table SIV.10** Details of the main parameters monitored for the group **$PPh_4$-$Mo_2O_4$-EDTA** during the **2013 test campaign**.*

| Sample | Hive numbering | Colony strenght | | Capped brood quantity, hundreds of cells | Queen bee's prolificity, eggs/24h | Beebread quantity, hundreds of cells | Honey quantity | | Wax | | Hygienic behaviour, % |
|---|---|---|---|---|---|---|---|---|---|---|---|
| | | Intervale | kg | | | | Number of frames | kg | number fabricated honeycomb | kg | |
| **1** | 5 | 14.3 | 3.58 | 220 | 1833 | 110 | 8.5 | 14.8 | 3.2 | 0.38 | 92.5 |
| **2** | 22 | 15.0 | 3.75 | 206 | 1717 | 120 | 8.5 | 14.7 | 3.2 | 0.38 | 93.9 |
| **3** | 29 | 14.5 | 3.63 | 200 | 1667 | 90 | 8.3 | 12.3 | 2.8 | 0.34 | 91.5 |
| **4** | 47 | 14.3 | 3.58 | 205 | 1708 | 125 | 8.5 | 16.0 | 3.5 | 0.42 | 93.0 |
| **5** | 48 | 13.8 | 3.45 | 210 | 1750 | 120 | 9.2 | 15.4 | 4.0 | 0.48 | 93.0 |
| **6** | 49 | 14.7 | 3.68 | 215 | 1792 | 124 | 8.7 | 15.5 | 4.0 | 0.48 | 94.0 |
| **7** | 50 | 15.0 | 3.75 | 206 | 1717 | 115 | 8.7 | 14.8 | 3.5 | 0.42 | 92.0 |
| **8** | 52 | 15.0 | 3.75 | 218 | 1817 | 125 | 8.5 | 14.5 | 3.5 | 0.42 | 95.0 |
| **9** | 54/51 | 12.8 | 3.20 | 205 | 1708 | 95 | 9.0 | 12.0 | 2.7 | 0.32 | 89.5 |
| **10** | 56 | 14.0 | 3.50 | 210 | 1750 | 110 | 8.5 | 13.5 | 2.9 | 0.35 | 91.0 |
| **11** | 58 | 14.2 | 3.55 | 220 | 1833 | 105 | 8.4 | 12.8 | 3.0 | 0.36 | 91.9 |
| **12** | 60 | 13.2 | 3.30 | 210 | 1750 | 100 | 8.5 | 11.0 | 2.8 | 0.34 | 91.0 |
| **13** | 62 | 15.0 | 3.75 | 212 | 1767 | 110 | 8.5 | 15.0 | 4.0 | 0.48 | 93.5 |
| **14** | 63 | 13.5 | 3.38 | 203 | 1692 | 108 | 8.0 | 13.7 | 3.0 | 0.36 | 91.0 |
| **15** | 65 | 15.8 | 3.95 | 225 | 1875 | 110 | 8.2 | 14.5 | 3.0 | 0.36 | 92.5 |
| **16** | 78 | 13.8 | 3.45 | 208 | 1733 | 96 | 9.5 | 12.5 | 2.9 | 0.35 | 91.0 |
| **M±m** | | | **3.58±0.05** | **210.8±1.70** | **1757.00±15.00** | **110.20±2.70** | | **13.94±0.36** | | **0.39±0.01** | **92.20±0.40** |
| **σ** | | | **0.19** | **7.00** | **58.00** | **11.00** | | **1.45** | | **0.05** | **1.40** |
| **Min** | | | **3.20** | **200.00** | **1667.00** | **90.00** | | **11.00** | | **0.32** | **89.50** |
| **Max** | | | **3.95** | **225.00** | **1875.00** | **125.00** | | **16.00** | | **0.48** | **95.00** |
| **m=σ/sqrt(16)** | | | **0.05** | **1.75** | **14.50** | **2.75** | | **0.36** | | **0.01** | **0.35** |

***Table SIV.11** Viability of larvae for the groups **control** and **$PPh_4$-$Mo_2O_4$-EDTA** during the **2013 test campaign**.*

| Sample | Group Control | | $PPh_4$-$Mo_2O_4$-EDTA | |
|---|---|---|---|---|
| | Hive numbering | Viability % | Hive numbering | Viabil % |
| **1** | 2 | 90.2 | 5 | 91.4 |
| **2** | 3 | 88.8 | 22 | 93.4 |
| **3** | 8 | 89.4 | 29 | 90.7 |
| **4** | 9 | 88.9 | 47 | 90.2 |
| **5** | 10 | 91.5 | 48 | 91.8 |
| **6** | 11 | 87.8 | 49 | 91.9 |
| **7** | 12 | 88.7 | 50 | 91.3 |
| **8** | 13 | 88.7 | 52 | 92.2 |
| **9** | 14 | 87.8 | 54 | 90.2 |
| **10** | 15 | 88.9 | 56 | 90.8 |
| **11** | 17 | 88.7 | 58 | 91.3 |
| **12** | 42 | 90.4 | 60 | 91.0 |
| **13** | 21 | 87.4 | 62 | 91.1 |
| **14** | 24 | 90.7 | 63 | 90.7 |
| **15** | 1 | 90.9 | 65 | 91.7 |
| **16** | 25 | 90.3 | 78 | 91.3 |
| **M±m** | | **89.30±0.30** | | **91.30±0.20** |
| **σ** | | **1.20** | | **0.80** |
| **Min** | | **87.40** | | **90.20** |
| **Max** | | **91.50** | | **93.40** |
| **m=σ/sqrt(16)** | | **0.30** | | **0.20** |

***Table SIV.12*** *Details of the main parameters monitored for the* ***control group*** *during the* ***2016 test campaign****.*

| Sample | Hive numbering | Colony strenght | | Capped brood quantity, hundreds of cells | Honey quantity, kg | Beebread quantity, hundreds of cells | Wax, | | Queen bee's prolificity, eggs/24h | Hygienic behaviour, % | |
|---|---|---|---|---|---|---|---|---|---|---|---|
| | | Intervale | kg | | | | number fabricated honeycomb | kg | | Cells over 50 | % |
| **1** | 14 | 3.4 | 0.85 | 141 | 3.3 | 84 | 2.3 | 0.28 | 475 | 42 | 84 |
| **2** | 36 | 4.9 | 1.23 | 155 | 5.0 | 95 | 2.5 | 0.30 | 1292 | 38 | 76 |
| **3** | 37 | 5.1 | 1.28 | 135 | 4.1 | 73 | 2.6 | 0.31 | 1125 | 47 | 94 |
| **4** | 30 | 3.4 | 0.85 | 132 | 4.1 | 88 | 2.1 | 0.25 | 1100 | 42 | 84 |
| **5** | 18 | 5.1 | 1.28 | 156 | 3.8 | 105 | 3.0 | 0.36 | 1300 | 47 | 94 |
| **6** | 44 | 4.8 | 1.20 | 159 | 3.8 | 90 | 2.5 | 0.30 | 1325 | 41 | 82 |
| **7** | 5 | 5.0 | 1.25 | 155 | 3.6 | 106 | 3.0 | 0.36 | 1292 | 48 | 96 |
| **8** | 22 | 3.3 | 0.83 | 145 | 3.9 | 84 | 1.8 | 0.22 | 1208 | 39 | 78 |
| **9** | 47 | 4.4 | 1.10 | 140 | 4.2 | 80 | 1.9 | 0.23 | 1167 | 43 | 86 |
| **10** | 48 | 5.0 | 1.25 | 134 | 3.8 | 97 | 2.7 | 0.32 | 1117 | 44 | 88 |
| **11** | 49 | 4.6 | 1.15 | 145 | 3.6 | 79 | 2.4 | 0.29 | 1208 | 45 | 90 |
| **12** | 50 | 4.5 | 1.13 | 119 | 4.4 | 91 | 2.1 | 0.25 | 992 | 42 | 84 |
| **M±m** | | | **1.12±0.05** | **143±3.46** | **3.97±0.13** | **89.30±2.94** | **2.41±0.11** | **0.29±0.01** | **1133.00±66.48** | | **86.30±1.82** |
| **σ** | | | **0.17** | **11.97** | **0.44** | **10.17** | **0.39** | **0.04** | **230.01** | | **6.31** |
| **m=σ/sqrt(12)** | | | **0.05** | **3.46** | **0.13** | **2.94** | **0.11** | **0.01** | **66.48** | | **1.82** |

.

***Table SIV.13** Details of the main parameters monitored for the **group K-$Mo_2O_2S_2$-LCys** during the **2016 test campaign**.*

| Sample | Hive numbering | Colony strenght | | Capped brood quantity, hundreds of cells | Honey quantity, kg | Beebread quantity, hundreds of cells | Wax | | Queen bee's prolificity, eggs/24h | Hygienic behaviour, % | |
|---|---|---|---|---|---|---|---|---|---|---|---|
| | | Intervale | kg | | | | number fabricated honeycomb | kg | | Cells over 50 | % |
| **1** | 39 | 5.2 | 1.30 | 165 | 4.7 | 70 | 2.8 | 0.34 | 1375 | 48 | 96 |
| **2** | 25 | 4.0 | 1.00 | 128 | 3.9 | 104 | 2.7 | 0.32 | 1067 | 44 | 88 |
| **3** | 16 | 4.0 | 1.00 | 147 | 3.2 | 98 | 2.5 | 0.30 | 1225 | 43 | 86 |
| **4** | 40 | 5.0 | 1.25 | 162 | 3.9 | 68 | 2.5 | 0.30 | 1350 | 45 | 90 |
| **5** | 19 | 4.5 | 1.13 | 139 | 4.1 | 103 | 2.7 | 0.32 | 1158 | 47 | 94 |
| **6** | 21 | 4.2 | 1.05 | 148 | 4.0 | 100 | 2.6 | 0.31 | 1233 | 40 | 80 |
| **7** | 46 | 5.1 | 1.28 | 166 | 4.8 | 84 | 3.0 | 0.36 | 1383 | 48 | 96 |
| **8** | 23 | 4.9 | 1.23 | 138 | 4.4 | 95 | 2.8 | 0.34 | 1150 | 46 | 92 |
| **9** | 24 | 3.7 | 0.93 | 130 | 4.0 | 83 | 2.0 | 0.24 | 1083 | 48 | 96 |
| **10** | 25 | 4.6 | 1.13 | 171 | 5.1 | 99 | 3.0 | 0.36 | 1425 | 47 | 94 |
| **M±m** | | | **1.13±0.04** | **149.40±4.97** | **4.20±0.17** | **90.40±4.22** | **2.70±0.09** | **0.32±0.01** | **1244.90±41.42** | | **91.20±1.66** |
| **σ** | | | **0.13** | **15.72** | **0.55** | **13.34** | **0.29** | **0.03** | **130.90** | | **5.26** |
| **m=σ/sqrt(10)** | | | **0.04** | **4.97** | **0.17** | **4.22** | **0.09** | **0.01** | **41.42** | | **1.66** |

***Table SIV.14*** *Details of the main parameters monitored for the* ***group $PPh_4$-$Mo_2O_2S_2$-EDTA*** *during the* ***2016 test campaign.***

| Sample | Hive numbering | Colony strenght | | Capped brood quantity, hundreds of cells | Honey quantity, kg | Beebread quantity, hundreds of cells | Wax | | Queen bee's prolificity, eggs/24h | Hygienic behaviour, % | |
|---|---|---|---|---|---|---|---|---|---|---|---|
| | | Intervale | kg | | | | number fabricated honeycomb | kg | | Cells over 50 | % |
| **1** | 26 | 4.8 | 1.20 | 152 | 5.2 | 103 | 3.0 | 0.36 | 1267 | 49 | 98 |
| **2** | 27 | 3.9 | 0.98 | 138 | 4.1 | 89 | 2.7 | 0.32 | 1150 | 45 | 90 |
| **3** | 28 | 3.8 | 0.95 | 144 | 4.5 | 94 | 2.6 | O.31 | 1200 | 45 | 90 |
| **4** | 29 | 4.9 | 1.23 | 152 | 3.6 | 100 | 2.9 | 0.35 | 1267 | 48 | 96 |
| **5** | 17 | 5.1 | 1.28 | 150 | 5.7 | 80 | 2.8 | 0.34 | 1250 | 43 | 86 |
| **6** | 31 | 4.8 | 1.20 | 154 | 3.9 | 104 | 2.9 | 0.35 | 1283 | 47 | 94 |
| **7** | 8 | 5.2 | 1.30 | 128 | 4.7 | 84 | 2.7 | 0.32 | 1067 | 49 | 98 |
| **8** | 33 | 4.1 | 1.03 | 175 | 4.4 | 85 | 2.7 | 0.32 | 1458 | 45 | 90 |
| **9** | 34 | 3.9 | 0.98 | 158 | 3.9 | 108 | 2.5 | 0.30 | 1317 | 44 | 88 |
| **10** | 10 | 4.7 | 1.18 | 147 | 3.9 | 94 | 2.5 | 0.30 | 1225 | 45 | 90 |
| **M±m** | | | **1.13±0.04** | **149.80±3.93** | **4.40±0.21** | **94.10±3.01** | **2.70±0.05** | **0.33±0.01** | **1248.40±32.68** | | **92.00±1.33** |
| **σ** | | | **0.13** | **12.41** | **0.66** | **9.51** | **0.17** | **0.02** | **103.28** | | **4.21** |
| **m=σ/sqrt(10)** | | | **0.04** | **3.93** | **0.21** | **3.01** | **0.05** | **0.01** | **32.68** | | **1.33** |

***Table SIV.15*** *Details of the main parameters monitored for the* ***group Control*** *during the* ***2018 test campaigns.***

| Sample | Hive numbering | Queen bee's prolificity, eggs/24h | Capped brood quantity, hundreds of cells | Colony strenght | | Beebread quantity, hundreds of cells | Wax | | Degree of infestation / 10 g of bees[a] | Hygienic behaviour, % | Honey. kg |
|---|---|---|---|---|---|---|---|---|---|---|---|
| | | | | Intervale | kg | | number fabricated honeycomb | kg | | | |
| **1** | 1 | 1167 | 140 | 8.6 | 2.15 | 135 | 2.0 | 0.24 | 1.1 | 88 | 13.9 |
| **2** | 2 | 1700 | 204 | 9.2 | 2.30 | 140 | 3.0 | 0.36 | 1.3 | 90 | 32 |
| **3** | 3 | 1875 | 225 | 9.3 | 2.33 | 125 | 2.5 | 0.30 | 2.3 | 84 | 8 |
| **4** | 44 | 1417 | 170 | 9.5 | 2.37 | 130 | 2.2 | 0.26 | 1.7 | 88 | 18 |
| **5** | 5 | 1664 | 200 | 10.5 | 2.63 | 120 | 2.4 | 0.29 | 2.5 | 86 | 40 |
| **6** | 6 | 1500 | 180 | 8.8 | 2.20 | 150 | 2.2 | 0.26 | 1.7 | 86 | 31 |
| **7** | 19 | 1458 | 175 | 9.5 | 2.38 | 115 | 2.3 | 0.28 | 1.7 | 88 | 27 |
| **8** | 8 | 1708 | 205 | 10.5 | 2.65 | 140 | 2.5 | 0.30 | 1.9 | 88 | 18 |
| **9** | 33 | 500 | 60 | 11.0 | 2.75 | 130 | 3.0 | 0.36 | 1.5 | 98 | 10 |
| **10** | 10 | 1333 | 160 | 9.5 | 2.37 | 125 | 2.5 | 0.30 | 2.1 | 94 | 10 |
| **M±m** | | **1432.20 ±112.60** | **171.90 ±14.71** | | **2.41 ±0.06** | **131.00 ±3.32** | | **0.30 ±0.01** | **1.78 ±0.14** | **89.00 ±1.31** | **20.79 ±3.49** |
| **σ** | | **387.60** | **46.50** | | **0.19** | **10.48** | | **0.03** | **0.43** | **4.13** | **11.04** |
| **m=σ/sqrt(10)** | | **122.66** | **14.72** | | **0.06** | **3.32** | | **0.01** | **0.14** | **1.31** | **3.49** |

a. number of mites Varroa per 10 g of bees, which approximatively corresponds to 100 bees. Results in %

***Table SIV.16*** *Details of the main parameters monitored for the* ***group $PPh_4$-$Mo_2O_4$-EDTA*** *during the* ***2018 test campaign.***

| Sample | Hive numbering | Queen bee's prolificity, eggs/24h | Capped brood quantity, hundreds of cells | Colony strenght | | Beebread quantity, hundreds of cells | Wax | | Degree of infestation / 10 g of bees[a] | Hygienic behaviour, % | Honey. kg |
|---|---|---|---|---|---|---|---|---|---|---|---|
| | | | | Intervale | kg | | number fabricated honeycomb | kg | | | |
| **11** | 11 | 1625 | 195 | 9.7 | 2.43 | 120 | 2.5 | 0.30 | 1.7 | 98 | 32 |
| **12** | 31 | 1167 | 140 | 8.5 | 2.12 | 145 | 2.0 | 0.24 | 1.1 | 98 | 16 |
| **13** | 13 | 1500 | 180 | 10.2 | 2.55 | 135 | 2.5 | 0.30 | 2.1 | 98 | 22 |
| **14** | 14 | 1333 | 160 | 8.5 | 2.13 | 140 | 2.0 | 0.24 | 1.9 | 90 | 20 |
| **15** | 15 | 1667 | 200 | 10.5 | 2.63 | 135 | 3.0 | 0.36 | 1.1 | 94 | 25 |
| **16** | 16 | 1625 | 195 | 10.8 | 2.70 | 150 | 3.0 | 0.36 | 1.1 | 86 | 32 |
| **17** | 17 | 1417 | 170 | 9.2 | 2.30 | 140 | 2.5 | 0.30 | 1.1 | 94 | 35 |
| **18** | 18 | 1625 | 195 | 10.2 | 2.55 | 120 | 3.0 | 0.36 | 1.3 | 98 | 15 |
| **19** | 7 | 1625 | 195 | 10.0 | 2.50 | 115 | 2.5 | 0.30 | 1.5 | 94 | 16 |
| **20** | 20 | 1750 | 210 | 10.8 | 2.70 | 110 | 3.0 | 0.36 | 1.1 | 98 | 22 |
| **M±m** | | **1533.40 ±56.55** | **184.00 ±6.78** | | **2.46 ±0.07** | **131.00 ±4.33** | | **0.31 ±0.01** | **1.40 ±0.12** | **94.80 ±1.31** | **23.50±2.31** |
| **σ** | | **178.70** | **21.44** | | **0.21** | **13.70** | | **0.05** | **0.38** | **4.13** | **7.31** |
| **m=σ/sqrt(10)** | | **56.55** | **6.78** | | **0.07** | **4.34** | | **0.02** | **0.12** | **1.31** | **2.31** |

a. number of mites Varroa per 10 g of bees, which approximatively corresponds to 100 bees. Results in %

***Table SIV.17*** *Details of the main parameters monitored for the* ***group Na-$Mo_2O_4$-EDTA*** *during the* ***2018 test campaign.***

| Sample | Hive numbering | Queen bee's prolificity, eggs/24h | Capped brood quantity, hundreds of cells | Colony strenght | | Beebread quantity, hundreds of cells | Wax | | Degree of infestation / 10 g of bees[a] | Hygienic behaviour, % | Honey. kg |
|---|---|---|---|---|---|---|---|---|---|---|---|
| | | | | Intervale | kg | | number fabricated honeycomb | kg | | | |
| **21** | 39 | 1583 | 190 | 6.6 | 2.50 | 135 | 2.5 | 0.30 | 1.3 | 90 | 40 |
| **22** | 47 | 1500 | 180 | 6.4 | 2.13 | 145 | 2.0 | 0.24 | 1.1 | 88 | 20 |
| **23** | 23 | 1750 | 210 | 7.0 | 2.38 | 105 | 2.5 | 0.30 | 1.7 | 94 | 40 |
| **24** | 24 | 1583 | 190 | 7.1 | 2.55 | 100 | 3.0 | 0.36 | 1.5 | 98 | 45 |
| **25** | 25 | 1583 | 190 | 6.5 | 2.50 | 115 | 3.0 | 0.36 | 2.1 | 92 | 23 |
| **26** | 26 | 1583 | 190 | 6.7 | 2.50 | 105 | 3.0 | 0.36 | 1.3 | 94 | 30 |
| **27** | 27 | 1633 | 196 | 6.8 | 2.50 | 120 | 2.5 | 0.30 | 2.3 | 88 | 20 |
| **28** | 28 | 1500 | 180 | 7.1 | 2.50 | 125 | 2.5 | 0.30 | 1.5 | 88 | 23 |
| **29** | 29 | 1667 | 200 | 7.0 | 2.50 | 120 | 3.0 | 0.36 | 1.5 | 84 | 40 |
| **30** | 35 | 1833 | 220 | 6.4 | 2.30 | 135 | 2.5 | 0.30 | 1.1 | 88 | 30 |
| **M±m** | | **1621.50 ±33.16** | **194.60 ±3.98** | | **2.44 ±0.04** | **120.50 ±4.68** | | **0.32 ±0.01** | **1.54 ±0.12** | **90.40 ±1.29** | **31.10 ±3.00** |
| **σ** | | **104.80** | **12.60** | | **0.13** | **14.80** | | **0.04** | **0.39** | **4.08** | **9.49** |
| **m=σ/sqrt(10)** | | **33.16** | **3.99** | | **0.04** | **4.68** | | **0.01** | **0.12** | **1.29** | **3.00** |

a. number of mites Varroa per 10 g of bees, which approximatively corresponds to 100 bees. Results in %

***Table SIV.18*** *Details of the main parameters monitored for the* ***group Li-$Mo_2O_4$-EDTA*** *during the* ***2018 test campaign.***

| Sample | Hive numbering | Queen bee's prolificity, eggs/24h | Capped brood quantity, hundreds of cells | Colony strenght | | Beebread quantity, hundreds of cells | Wax | | Degree of infestation / 10 g of bees[a] | Hygienic behaviour, % | Honey. kg |
|---|---|---|---|---|---|---|---|---|---|---|---|
| | | | | Intervale | kg | | number fabricated honeycomb | kg | | | |
| **31** | 12 | 1417 | 170 | 9.0 | 2.75 | 115 | 2.5 | 0.30 | 1.3 | 90 | 20 |
| **32** | 32 | 1625 | 195 | 10.2 | 2.55 | 130 | 3.0 | 0.36 | 1.1 | 96 | 22 |
| **33** | 9 | 1667 | 200 | 9.5 | 2.38 | 140 | 3.0 | 0.36 | 0.7 | 92 | 20 |
| **34** | 34 | 1250 | 150 | 9.8 | 2.45 | 138 | 2.5 | 0.30 | 1.1 | 94 | 20 |
| **35** | 30 | 1708 | 205 | 10.0 | 2.50 | 120 | 3.0 | 0.36 | 1.3 | 90 | 40 |
| **36** | 36 | 1625 | 195 | 10.0 | 2.50 | 135 | 3.0 | 0.36 | 1.1 | 88 | 20 |
| **37** | 37 | 1667 | 200 | 10.0 | 2.50 | 125 | 3.0 | 0.36 | 0.7 | 92 | 34 |
| **38** | 38 | 1542 | 185 | 9.5 | 2.38 | 100 | 2.5 | 0.30 | 1.1 | 86 | 53 |
| **39** | 21 | 1667 | 200 | 11.0 | 2.75 | 105 | 3.5 | 0.42 | 0.9 | 96 | 26 |
| **40** | 40 | 1833 | 220 | 10.5 | 2.63 | 130 | 3.0 | 0.36 | 0.9 | 92 | 42 |
| **M±m** | | **1600.10 ±51.68** | **192.00 ±6.20** | | **2.54 ±0.04** | **123.80 ±4.30** | | **0.35 ±0.01** | **1.02 ±0.07** | **91.60 ±1.01** | **29.70 ±3.76** |
| **σ** | | **163.32** | **19.60** | | **0.13** | **13.60** | | **0.03** | **0.21** | **3.20** | **11.87** |
| **m=σ/sqrt(10)** | | **51.68** | **6.20** | | **0.04** | **4.30** | | **0.01** | **0.07** | **1.01** | **3.76** |

a. number of mites Varroa per 10 g of bees, which approximatively corresponds to 100 bees. Results in %

***Table SIV.19*** *Details of the main parameters monitored for the group* ***Control*** *during the* ***2019 test campaign.***

| Sample | Hive numbering | Colony strenght | | Capped brood quantity hundreds of cells. | Queen bee's prolificity, eggs/24h | Amount of food accumulated in the nest | | Wax | | Hygienic behaviour, % | | Larvae viability at 5 days % | Anti-Varroa properties | | |
|---|---|---|---|---|---|---|---|---|---|---|---|---|---|---|---|
| | | Inter vals | kg | | | Hone y kg | Beebread quantity, hundreds of cells | number honey-comb | kg | Cells over 50 | % | | Fallen mites in 24 h | Bee infestat ion rate,% | Degree of brood infestat ion |
| **1** | 1 | 14.5 | 3.62 | 200 | 1667 | 21 | 120 | 2.6 | 0.31 | 45 | 90 | 88 | 21 | 5.3 | 32 |
| **2** | 2 | 12.0 | 3.00 | 195 | 1625 | 15 | 100 | 2.0 | 0.24 | 44 | 88 | 86 | 8 | 1.3 | 18 |
| **3** | 4 | 15.0 | 3.75 | 195 | 1625 | 23 | 100 | 2.5 | 0.30 | 42 | 84 | 88 | 6 | 0.9 | 40 |
| **4** | 6 | 12.8 | 3.20 | 190 | 1583 | 21 | 105 | 1.8 | 0.22 | 40 | 82 | 88 | 9 | 1.9 | 52 |
| **5** | 7 | 13.0 | 3.25 | 190 | 1583 | 18 | 130 | 2.5 | 0.30 | 44 | 88 | 90 | 20 | 2.3 | 38 |
| **6** | 8 | 13.2 | 3.30 | 180 | 1500 | 20 | 115 | 2.0 | 0.24 | 43 | 86 | 86 | 11 | 2.0 | 24 |
| **7** | 9 | 14.0 | 3.50 | 210 | 1750 | 22 | 135 | 2.2 | 0.26 | 44 | 88 | 88 | 14 | 1.9 | 8 |
| **8** | 61 | 14.5 | 3.62 | 215 | 1792 | 22 | 130 | 2.0 | 0.24 | 44 | 88 | 90 | 10 | 3.4 | 16 |
| **9** | 12 | 15.3 | 3.82 | 200 | 1667 | 24 | 110 | 2.7 | 0.32 | 45 | 90 | 86 | 17 | 2.8 | 36 |
| **10** | 14 | 11.2 | 2.80 | 180 | 1500 | 20 | 95 | 1.9 | 0.23 | 40 | 80 | 80 | 10 | 1.4 | 18 |
| **M±m** | | | **3.39 ±0.10** | **195.50 ±3.61** | **1629.20 ±30.13** | **21.00 ±0.82** | **114.00 ±4.52** | | **0.27 ±0.01** | | **86.00 ±1.07** | **87.00 ±0.91** | **13.00 ±1.63** | **2.30 ±0.40** | **28.20 ±4.29** |
| **σ** | | | **0.33** | **11.41** | **95.20** | **2.59** | **14.29** | | **0.04** | | **3.37** | **2.86** | **5.16** | **1.27** | **13.57** |
| **m=σ/sqrt(10)** | | | **0.10** | **3.61** | **30.13** | **0.82** | **4.52** | | **0.01** | | **1.07** | **0.91** | **1.63** | **0.40** | **4.29** |

***Table SIV.20*** *Details of the main parameters monitored for the group* ***Li-Mo$_2$O$_4$-EDTA (2mg)*** *during the* ***2019 test campaign.***

| Sample | Hive numbering | Colony strenght | | Capped brood quantity hundreds of cells. | Queen bee's prolificity, eggs/24h | Amount of food accumulated in the nest | | Wax | | Hygienic behaviour, % | | Larvae viability at 5 days % | Anti-Varroa properties | | |
|---|---|---|---|---|---|---|---|---|---|---|---|---|---|---|---|
| | | Inter vals | kg | | | Hone y kg | Beebread quantity, hundreds of cells | number honey-comb | kg | Cells over 50 | % | | Fallen mites in 24 h | Bee infestat ion rate,% | Degree of brood infestat ion |
| **1** | 28 | 15 | 3.75 | 220 | 1833 | 24 | 150 | 3.0 | 0.36 | 48 | 96 | 88 | 20 | 1.3 | 10 - 20 |
| **2** | 24 | 14 | 3.50 | 195 | 1625 | 21 | 135 | 2.8 | 0.34 | 46 | 92 | 86 | 18 | 1.0 | 11 - 22 |
| **3** | 34 | 16 | 4.00 | 210 | 1750 | 26 | 125 | 2.8 | 0.34 | 50 | 100 | 90 | 19 | 0.8 | 14 - 28 |
| **4** | 38 | 15 | 3.75 | 200 | 1667 | 23 | 135 | 2.9 | 0.35 | 44 | 88 | 88 | 24 | 0.3 | 19 - 38 |
| **5** | 42 | 18 | 4.50 | 220 | 1833 | 25 | 150 | 3.1 | 0.37 | 50 | 100 | 92 | 15 | 1.0 | 17 - 34 |
| **6** | 16 | 16 | 4.00 | 210 | 1750 | 22 | 145 | 2.9 | 0.35 | 46 | 92 | 86 | 5 | 0.8 | 8 - 16 |
| **7** | 45 | 16 | 4.00 | 215 | 1792 | 24 | 140 | 2.4 | 0.29 | 46 | 92 | 90 | 15 | 1.9 | 4 - 8 |
| **8** | 46 | 17 | 4.25 | 220 | 1833 | 26 | 150 | 3.0 | 0.36 | 48 | 96 | 90 | 8 | 1.8 | 14 - 28 |
| **9** | 56 | 16 | 4.00 | 215 | 1792 | 24 | 140 | 2.7 | 0.32 | 46 | 92 | 88 | 8 | 0.9 | 13 - 26 |
| **10** | 52 | 13 | 3.25 | 190 | 1583 | 22 | 125 | 2.5 | 0.30 | 47 | 94 | 84 | 14 | 2.4 | 10 - 20 |
| **M±m** | | | **3.90±0.11** | **209.50±3.45** | **1745.80±28.77** | **24.00±0.54** | **140.00±3.02** | | **0.34±0.01** | | **94.00±1.20** | **88.20±0.76** | **15.00±1.91** | **1.22±0.20** | **12.00±1.38** |
| **σ** | | | **0.35** | **10.90** | **90.90** | **1.70** | **9.55** | | **0.03** | | **3.80** | **2.39** | **6.04** | **0.63** | **4.37** |
| **m=σ/sqrt(10)** | | | **0.11** | **3.45** | **28.77** | **0.54** | **3.02** | | **0.01** | | **1.20** | **0.76** | **1.91** | **0.20** | **1.38** |

***Table SIV.21*** *Details of the main parameters monitored for the group* ***$LiCH_3COO$*** *during the* ***2019 test campaign.***

| Sample | Hive numbering | Colony strenght | | Capped brood quantity hundreds of cells. | Queen bee's prolificity. eggs/24h | Amount of food accumulated in the nest | | Wax | | Hygienic behaviour, % | | Larvae viability at 5 days % | Anti-Varroa properties | | |
|---|---|---|---|---|---|---|---|---|---|---|---|---|---|---|---|
| | | Intervals | kg | | | Honey kg | Beebread quantity. hundreds of cells | number honey-comb | kg | Cells over 50 | % | | Fallen mites in 24 h | Bee infestation rate.% | Degree of brood infestation |
| **1** | **21** | 14 | 3.5 | 180 | 1500 | 16 | 105 | 2.3 | 0.28 | 44 | 88 | 84 | 27 | 1.0 | 35 |
| **2** | **23** | 12.5 | 3.12 | 180 | 1500 | 16 | 115 | 2.2 | 0.26 | 41 | 82 | 80 | 20 | 1.7 | 14 |
| **3** | **31** | 13 | 3.25 | 165 | 1375 | 15 | 110 | 2.5 | 0.30 | 42 | 84 | 84 | 18 | 1.84 | 20 |
| **4** | **26** | 14.5 | 3.62 | 200 | 1667 | 22 | 125 | 2.6 | 0.31 | 48 | 96 | 86 | 17 | 0.83 | 8 |
| **5** | **27** | 14 | 3.5 | 195 | 1625 | 23 | 130 | 2.5 | 0.30 | 48 | 96 | 82 | 15 | 1.9 | 20 |
| **6** | **56** | 13.5 | 3.37 | 200 | 1667 | 20 | 120 | 2.6 | 0.31 | 45 | 90 | 82 | 12 | 1.3 | 14 |
| **7** | **30** | 13 | 3.25 | 210 | 1750 | 22 | 120 | 2.8 | 0.34 | 44 | 88 | 80 | 16 | 1.5 | 10 |
| **8** | **32** | 12.5 | 3.12 | 200 | 1667 | 18 | 135 | 2.5 | 0.30 | 46 | 92 | 80 | 15 | 1.8 | 18 |
| **9** | **35** | 15 | 3.75 | 210 | 1750 | 22 | 120 | 2.2 | 0.26 | 46 | 92 | 82 | 11 | 1.7 | 28 |
| **10** | **44** | 11 | 2.75 | 175 | 1458 | 17 | 130 | 1.9 | 0.23 | 45 | 90 | 78 | 21 | 2.4 | 16 |
| **M±m** | | | **3.32 ±0.09** | **191.5±4.9** | **1596±41** | **19.10 ±0.96** | **121.0±3.0** | | **0.29±0.01** | | **89.8±1.4** | **81.8±0.7** | **17.2±1.5** | **1.60±0.15** | **18.3±2.6** |
| **σ** | | | **0.29** | **15.5** | **129** | **3.03** | **9.4** | | **0.03** | | **4.6** | **2.4** | **4.7** | **0.46** | **8.1** |
| **m=σ/sqrt(10)** | | | **0.05** | **4.9** | **40.82** | **1.04** | **2.97** | | **0.01** | | **1.46** | **0.76** | **1.49** | **0.15** | **2.56** |

***Table SIV.22*** *Details of the main parameters monitored for the group* ***Li-$Mo_2O_4$-EDTA (6mg)*** *during the* ***2019 test campaign.***

| Sample | Hive numbering | Colony strenght | | Capped brood quantityhundreds of cells. | Queen bee's prolificity. eggs/24h | Amount of food accumulated in the nest | | Wax | | Hygienic behaviour, % | | Larvae viability at 5 days % | Anti-Varroa properties | | |
|---|---|---|---|---|---|---|---|---|---|---|---|---|---|---|---|
| | | Intervals | kg | | | Honey kg | Beebread quantity. hundreds of cells | number honey-comb | kg | Cells over 50 | % | | Fallen mites in 24 h | Bee infestation rate.% | Degree of brood infestation |
| **1** | **39** | | 4.0 | 215 | 1792 | 26 | 125 | 3.2 | 0.38 | 45 | 90 | 88 | 25 | 0.56 | 3 |
| **2** | **19** | | 3.75 | 200 | 1667 | 24 | 120 | 2.6 | 0.31 | 48 | 96 | 86 | 18 | 1.0 | 8 |
| **3** | **47** | | 4.25 | 230 | 1917 | 22 | 135 | 3.0 | 0.36 | 47 | 94 | 90 | 19 | 0.8 | 9 |
| **4** | **49** | | 4.0 | 215 | 1792 | 23 | 130 | 2.9 | 0.35 | 50 | 100 | 84 | 24 | 1.52 | 5 |
| **5** | **51** | | 4.5 | 245 | 2042 | 28 | 145 | 3.5 | 0.42 | 50 | 100 | 90 | 15 | 0.8 | 4 |
| **6** | **53** | | 3.75 | 210 | 1750 | 25 | 135 | 2.5 | 0.30 | 48 | 96 | 88 | 5 | 1.0 | 2 |
| **7** | **29** | | 4.25 | 220 | 1833 | 25 | 150 | 3.0 | 0.36 | 48 | 96 | 90 | 15 | 1.9 | 13 |
| **8** | **59** | | 4.25 | 235 | 1958 | 26 | 155 | 3.4 | 0.41 | 50 | 100 | 92 | 8 | 0.8 | 0 |
| **9** | **10** | | 4.0 | 220 | 1833 | 24 | 140 | 2.8 | 0.34 | 49 | 98 | 90 | 8 | 0.0 | 1 |
| **10** | **15** | | 3.75 | 200 | 1667 | 24 | 140 | 2.4 | 0.29 | 47 | 94 | 84 | 148 | 0.56 | 7 |
| **M±m** | | | **4.05±0.08** | **219.0±4.6** | **1825±38** | **24.7±0.5** | **137.5±3.4** | | **0.35±0.01** | | **96.4±1.0** | **88.2±0.8** | | **0.89±0.16** | **5.2±1.3** |
| **σ** | | | **0.26** | **14.5** | **121** | **1.70** | **10.9** | | **0.04** | | **6.7** | **2.7** | | **0.52** | **4.05** |
| **m=σ/sqrt(10)** | | | **0.08** | **4.59** | **38.3** | **0.53** | **3.45** | | **0.01** | | **2.12** | **0.85** | | **0.16** | **1.42** |

***Table SIV.23.*** *Details of the mass and the quantity of honey for beehives of the group* ***Control*** *of the experiment (France, Gif-sur-Yvette Campaign 2019)*

| Sample | Hive Numbering | Mass 1th April. kg | Mass 16th April. kg | Mass 24th April. kg | Mass 2nd May. kg | Mass 13th May. kg | Spring Honey. kg 13th May | Mass 29th May. kg | Mass 8th June. kg | Mass 15th June. kg | Mass 22th June. kg | Mass 15th July. kg | Honey total. kg |
|---|---|---|---|---|---|---|---|---|---|---|---|---|---|
| 1 | D | 22.79 | 22.95 | 28.80 | 29.1 | 26.8 | 2.30 | 25.60 | 25.80 | 29.40 | 27.00 | 34.75 | 2.55 |
| 2 | F | 26.70 | 27.00 | 36.95 | 38.4 | 35.2 | 3.70 | 35.25 | 36.80 | 39.55 | 41.80 | 71.75 | 30.10 |
| 3 | 195 | 24.25 | 25.45 | 34.10 | 34.45 | 30.6 | 3.45 | 29.65 | 29.80 | 34.60 | 36.05 | 47.8 | 7.30 |
| 4 | 25 | 21.05 | 21.10 | 25.60 | 25.9 | 23.9 | 1.85 | 23.65 | 24.40 | 27.85 | 27.35 | 37.6 | 2.10 |
| 5 | E | 20.20 | 20.15 | 23.60 | 23.75 | 21.7 | 1.10 | 22.05 | 23.20 | 27.70 | 26.65 | 38.35 | 7.15 |
| 6 | B | 24.40 | 23.80 | 30.45 | 31.1 | 28.3 | 3.70 | 28.10 | 28.35 | 31.10 | 31.40 | 50.05 | 18.15 |
| 7 | 44D | 23.05 | 23.50 | 32.80 | 34.3 | 31.4 | 3.10 | 31.40 | 32.25 | 35.50 | 36.20 | 62.45 | 28.30 |
| 8 | C | 22.90 | 24.20 | 36.00 | 37.8 | 34.4 | 3.80 | 34.35 | 34.55 | 36.10 | 35.70 | 64.85 | 31.60 |
| 9 | 8 | 29.75 | 32.15 | 40.70 | 43.85 | 41.15 | 10.85 | 35.35 | 36.90 | 40.85 | 43.10 | 69.05 | 30.95 |
| 10 | 35 | 22.90 | 23.30 | 31.50 | 32.6 | 29.35 | 2.85 | 29.50 | 30.85 | 33.40 | 34.35 | 62.05 | 32.55 |
| 11 | I | 30.55 | 33.05 | 41.00 | 43.6 | 39.3 | 11.25 | 33.30 | 33.45 | 35.65 | 36.05 | 57.1 | 19.70 |
| **M±m** | | 24.41 ±1.00 | 25.15 ±1.24 | 32.86 ±1.70 | 34.08 ±1.97 | 31.10 ±1.82 | 4.36 ±1.03 | 29.84 ±1.40 | 30.58 ±1.44 | 33.79 ±1.33 | 34.15 ±1.69 | 54.16 ±3.98 | 19.13 ±3.73 |
| **σ** | | 3.16 | 3.93 | 5.38 | 6.24 | 5.77 | 3.26 | 4.42 | 4.54 | 4.20 | 5.34 | 12.59 | 11.79 |
| **m=σ/sqrt(10)** | | 1.00 | 1.24 | 1.70 | 1.97 | 1.82 | 1.03 | 1.40 | 1.44 | 1.33 | 1.69 | 3.98 | 3.73 |

At the initial time, the eleven colonies of the control group are estimated to be comparable. Nevertheless, two of them appears a bit stronger at the starting (samples 9 and 11). Beehive's honey supers have been generally added around the 24th may, excepted for samples 9 and 11 for which honey supers have been added earlier. In summer, additional honey supers have been added at the end of June, excepted for samples 1 and 4, for which it was not necessary.

***Table SIV.24.*** *Details of the mass and the quantity of honey for beehives of the group* ***Li-Mo₂O₄-EDTA*** *of the experiment. (France, Gif-sur-Yvette Campaign 2019)*

| Sample | Hive Numbering | Mass 1th April. kg | Mass 16th April. kg | Mass 24th April. kg | Mass 2nd May. kg | Mass 13th May. kg | Spring Honey. kg 13th May | Mass 29th May. kg | Mass 8th June. kg | Mass 15th June. kg | Mass 22th June. kg | Mass 15th July. kg | Honey total. kg |
|---|---|---|---|---|---|---|---|---|---|---|---|---|---|
| 12 | 44G | 24.50 | 26.05 | 33.50 | 35.7 | 32.65 | 2.65 | 31.55 | 33.00 | 35.05 | 35.00 | 74.45 | 36.70 |
| 13 | G | 25.50 | 25.80 | 38.05 | 40.05 | 36.8 | 7.00 | 37.00 | 37.55 | 39.15 | 46.40 | 76.6 | 38.75 |
| 14 | H | 24.20 | 24.60 | 32.30 | 33 | 29.5 | 4.90 | 27.60 | 27.65 | 29.60 | 30.60 | 37.9 | 5.35 |
| 15 | 26 | 26.00 | 25.80 | 39.10 | 40.9 | 37.4 | 7.25 | 37.85 | 40.45 | 43.45 | 48.55 | 86.05 | 46.65 |
| 16 | 91 | 25.05 | 25.75 | 33.85 | 35.4 | 33.5 | 2.30 | 33.30 | 34.20 | 37.30 | 37.20 | 64.55 | 30.85 |
| 17 | 51 | 22.70 | 23.70 | 33.55 | 35.5 | 32.6 | 4.00 | 32.30 | 32.50 | 32.35 | 34.10 | 69.65 | 33.00 |
| 18 | C2 | 21.05 | 22.25 | 28.90 | 30.1 | 27.5 | 1.70 | 27.25 | 28.30 | 31.95 | 33.40 | 55.7 | 20.85 |
| 19 | B2 | 24.55 | 24.60 | 31.50 | 32.8 | 30.4 | 3.70 | 29.80 | 31.25 | 33.25 | 36.75 | 68.15 | 34.65 |
| 20 | 15 | 27.90 | 27.75 | 36.35 | 37.05 | 33.1 | 2.70 | 32.80 | 34.85 | 37.15 | 38.70 | 71.75 | 33.80 |
| 21 | 2 | 28.20 | 28.40 | 40.85 | 43.75 | 40.3 | 4.50 | 40.65 | 42.10 | 42.10 | 42.60 | 56.9 | 18.25 |
| 22 | 9 | 26.10 | 26.35 | 37.85 | 39.1 | 36.1 | 5.60 | 36.05 | 37.20 | 38.85 | 45.15 | 74.65 | 34.75 |
| **M±m** | | 25.07 ±0.63 | 25.55 ±0.52 | 35.07 ±1.10 | 36.67 ±1.21 | 33.62 ±1.14 | 4.21 ±0.56 | 33.29 ±1.28 | 34.46 ±1.39 | 36.38 ±1.32 | 38.95 ±1.78 | 66.94 ±3.91 | 30.33 ±3.43 |
| **σ** | | 1.98 | 1.66 | 3.47 | 3.83 | 3.60 | 1.77 | 4.06 | 4.39 | 4.17 | 5.62 | 12.37 | 10.85 |
| **m=σ/sqrt(10)** | | 0.63 | 0.52 | 1.10 | 1.21 | 1.14 | 0.56 | 1.28 | 1.39 | 1.32 | 1.78 | 3.91 | 3.43 |

At the initial time, the eleven colonies of the group are estimated to be comparable. Beehive's honey supers have been generally added around the 24th may. In summer, additional honey supers have been added at the end of June, excepted for sample 14, for which it was not necessary.

**Table SIV.25** Evolution of Brood cells number in the different modalities of the experiment. Greece campaign.

| Group | Hive number | Dates (period 1) 07/12/2020 | Dates (period 1) 03/03/2021 | Group | Dates (period 2) 03/03/2021 | Dates (period 2) 20/04/2021 |
|---|---|---|---|---|---|---|
| MoLi (20 hives at the beginning; 19 hives during the experiment) | 2 | 0 | 19600 | MoLi-B (9 hives) | 19600 | 25900 |
| | 13 | 0 | 7700 | | 7700 | 18900 |
| | 19 | 0 | 12600 | | 12600 | 25900 |
| | 101 | 0 | 9100 | | 9100 | 35000 |
| | 103 | 0 | 9100 | | 9100 | 32200 |
| | 105 | 0 | 24500 | | 24500 | 27300 |
| | 106 | 0 | 20300 | | 20300 | 23800 |
| | 112 | 0 | 17500 | | 17500 | 16800 |
| | 7 | 0 | - | | - | - |
| | 401 | 0 | 7000 | | 7000 | 11200 |
| | | | | Average | 14156 | 24111 |
| | 18 | 6300 | 18200 | MoLi-A (10 hives) | 18200 | 29400 |
| | 16 | 0 | 8400 | | 8400 | 28000 |
| | 15 | 4200 | 13300 | | 13300 | 23800 |
| | 14 | 2100 | 17500 | | 17500 | 34300 |
| | 12 | 0 | 11900 | | 11900 | 30100 |
| | 11 | 2800 | 14700 | | 14700 | 17500 |
| | 404 | 0 | 7000 | | 7000 | 11900 |
| | 10 | 0 | 4200 | | 4200 | 11200 |
| | 402 | 0 | 9800 | | 9800 | 18200 |
| | 8 | 4900 | 23800 | | 23800 | 26600 |
| Average | | 1015 | 12810 | Average | 12880 | 23100 |
| Control (9 hives at the beginning; 8 hives | 27 | 0 | 16100 | Control (8 hives) | 16100 | 35000 |
| | 26 | 0 | - | | - | - |
| | 25 | 0 | 18200 | | 18200 | 37100 |
| | 24 | 0 | 25200 | | 25200 | 35000 |
| | 23 | 0 | 17500 | | 17500 | 26600 |
| | 22 | 0 | 21000 | | 21000 | 29400 |

| | | | | | | |
|---|---|---|---|---|---|---|
| during the experiment) | 21 | 0 | 6300 | | 6300 | 22400 |
| | 20 | 0 | 11200 | | 11200 | 24500 |
| | 19 | 0 | 15400 | | 15400 | 30100 |
| Average | | 0 | 16363 | Average | 16363 | 30013 |

**Table SIV.26** Evolution of population of adult bees in the different modalities of the experiment.

| | | Dates (period 1) | | | Dates (period 2) | |
|---|---|---|---|---|---|---|
| Group | Hive number | 07/12/2020 | 03/03/2021 | Group | 03/03/2021 | 20/04/2021 |
| MoLi (20 hives at the beginning; 19 hives during the experiment) | 2 | 13570 | 13800 | MoLi-B (9 hives) | 13800 | 16100 |
| | 13 | 9660 | 7820 | | 7820 | 7820 |
| | 19 | 11730 | 10580 | | 10580 | 13340 |
| | 101 | 10580 | 8510 | | 8510 | 16100 |
| | 103 | 9430 | 8970 | | 8970 | 17710 |
| | 105 | 15410 | 12650 | | 12650 | 16100 |
| | 106 | 15180 | 12420 | | 12420 | 15410 |
| | 112 | 13110 | 11040 | | 11040 | 9890 |
| | 7 | 7590 | - | | - | - |
| | 401 | 6440 | 5520 | | 5520 | 10350 |
| | - | - | - | Average | 10146 | 13647 |
| | 18 | 13800 | 11960 | MoLi-A (10 hives) | 11960 | 16330 |
| | 16 | 17250 | 14720 | | 14720 | 15180 |
| | 15 | 12880 | 9430 | | 9430 | 10350 |
| | 14 | 14490 | 12190 | | 12190 | 19090 |
| | 12 | 10580 | 7820 | | 7820 | 20700 |
| | 11 | 13340 | 10350 | | 10350 | 9890 |
| | 404 | 6900 | 5290 | | 5290 | 5290 |
| | 10 | 4600 | 3680 | | 3680 | 6440 |
| | 402 | 9660 | 7130 | | 7130 | 9890 |
| | 8 | 16560 | 16100 | | 16100 | 22080 |

| | | | | | | |
|---|---|---|---|---|---|---|
| Average | | 11638 | 9999 | Average | 9867 | 13524 |
| Control<br><br>(9 hives at the beginning; 8 hives during the experiment) | 27 | 15410 | 10810 | Control<br><br>(8 hives) | 10810 | 17710 |
| | 26 | 17480 | - | | - | - |
| | 25 | 11730 | 9660 | | 9660 | 16100 |
| | 24 | 15180 | 17020 | | 17020 | 17480 |
| | 23 | 13340 | 15180 | | 15180 | 15180 |
| | 22 | 13570 | 13800 | | 13800 | 19550 |
| | 21 | 7590 | 9660 | | 9660 | 14260 |
| | 20 | 10810 | 8740 | | 8740 | 14260 |
| | 19 | 11040 | 9660 | | 9660 | 17020 |
| Average | | 12906 | 11816 | Average | 11816 | 16445 |

**Table SIV.27** Count of dead bees in front of the hive for the different methods during the first period of the experiment (7 dec. - 5 march 2021).

| | | Dates | | | | | | | | | |
|---|---|---|---|---|---|---|---|---|---|---|---|
| Group | Hive number | Dec. 11 | Dec. 15 | Dec. 18 | Dec. 21 | Dec. 23 | Dec. 28 | Dec. 30 | Jan. 4 | Jan. 8 | Jan. 11 |
| MoLi<br>(20 hives) | 2 | 1 | 3 | 1 | 0 | 2 | 0 | 0 | 0 | 0 | 4 |
| | 13 | 0 | 6 | 1 | 1 | 3 | 1 | 0 | 0 | 0 | 3 |
| | 19 | 6 | 6 | 1 | 3 | 2 | 0 | 1 | 3 | 0 | 5 |
| | 101 | 4 | 6 | 0 | 4 | 8 | 4 | 0 | 4 | 3 | 25 |
| | 103 | 7 | 0 | 0 | 0 | 0 | 0 | 0 | 1 | 2 | 3 |
| | 105 | 11 | 1 | 0 | 0 | 2 | 1 | 0 | 3 | 3 | 56 |
| | 106 | 3 | 0 | 1 | 1 | 2 | 1 | 1 | 2 | 0 | 12 |
| | 112 | 11 | 1 | 1 | 0 | 7 | 1 | 0 | 1 | 1 | 4 |
| | 7 | 7 | 10 | 4 | 2 | 6 | 5 | 2 | 4 | 3 | 4 |
| | 401 | 13 | 14 | 5 | 7 | 7 | 3 | 1 | 2 | 0 | 3 |
| | 18 | 60 | 21 | 6 | 3 | 3 | 3 | 3 | 3 | 5 | 20 |
| | 16 | 30 | 24 | 5 | 1 | 7 | 4 | 1 | 3 | 3 | 14 |
| | 15 | 70 | 58 | 7 | 4 | 4 | 3 | 4 | 5 | 4 | 12 |
| | 14 | 27 | 68 | 8 | 11 | 2 | 6 | 1 | 3 | 2 | 13 |

| | | | | | | | | | | | |
|---|---|---|---|---|---|---|---|---|---|---|---|
| | 12 | 8 | 5 | 0 | 0 | 1 | 2 | 2 | 14 | 5 | 7 |
| | 11 | 28 | 12 | 3 | 1 | 2 | 3 | 2 | 2 | 0 | 8 |
| | 404 | 11 | 3 | 4 | 2 | 3 | 3 | 1 | 1 | 2 | 1 |
| | 10 | 13 | 19 | 5 | 6 | 6 | 3 | 3 | 3 | 3 | 6 |
| | 402 | 22 | 4 | 1 | 1 | 2 | 2 | 0 | 3 | 2 | 8 |
| | 8 | 18 | 13 | 9 | 8 | 3 | 3 | 0 | 1 | 1 | 9 |
| Sum | | 350 | 274 | 62 | 55 | 72 | 48 | 22 | 58 | 39 | 217 |
| Average/hive | | 17.5 | 13.7 | 3.1 | 2.75 | 3.6 | 2.4 | 1.1 | 2.9 | 1.95 | 10.85 |
| Control (9 hives) | 27 | 5 | 0 | 7 | 5 | 0 | 1 | 3 | 2 | 5 | 3 |
| | 26 | 37 | 5 | 8 | 9 | 6 | 3 | 2 | 1 | 0 | 9 |
| | 25 | 7 | 1 | 3 | 2 | 3 | 1 | 1 | 3 | 1 | 6 |
| | 24 | 5 | 5 | 1 | 0 | 1 | 0 | 2 | 0 | 0 | 8 |
| | 23 | 7 | 1 | 3 | 4 | 2 | 0 | 3 | 1 | 0 | 10 |
| | 22 | 6 | 26 | 3 | 7 | 4 | 2 | 1 | 3 | 3 | 3 |
| | 21 | 8 | 4 | 0 | 0 | 3 | 1 | 0 | 2 | 0 | 4 |
| | 20 | 22 | 46 | 5 | 3 | 16 | 3 | 2 | 4 | 8 | 6 |
| | 19 | 6 | 5 | 1 | 0 | 2 | 0 | 1 | 3 | 0 | 5 |
| Sum | | 103 | 93 | 31 | 30 | 37 | 11 | 15 | 19 | 17 | 54 |
| Average/hive | | 11.44 | 10.33 | 3.44 | 3.33 | 4.11 | 1.22 | 1.67 | 2.11 | 1.89 | 6.00 |

| | | Dates | | | | | | | | | |
|---|---|---|---|---|---|---|---|---|---|---|---|
| Group | Hive number | Jan. 13 | Jan. 15 | Jan. 18 | Jan. 20 | Jan. 22 | Jan. 25 | Jan. 28 | Feb. 1 | Feb. 4 | Feb. 4 |
| MoLi (20 hives) | 2 | 0 | 3 | 0 | 1 | 6 | 0 | 1 | 1 | 0 | 1 |
| | 13 | 0 | 3 | 0 | 2 | 21 | 4 | 0 | 1 | 0 | 0 |
| | 19 | 3 | 1 | 7 | 5 | 8 | 34 | 1 | 3 | 2 | 0 |
| | 101 | 9 | 0 | 3 | 3 | 12 | 5 | 3 | 6 | 1 | 9 |
| | 103 | 3 | 4 | 0 | 3 | 21 | 1 | 0 | 5 | 1 | 0 |
| | 105 | 5 | 0 | 2 | 6 | 22 | 2 | 2 | 6 | 2 | 0 |
| | 106 | 1 | 0 | 2 | 0 | 5 | 5 | 2 | 0 | 3 | 0 |
| | 112 | 1 | 1 | 0 | 2 | 1 | 0 | 3 | 0 | 0 | 0 |
| | 7 | 7 | 4 | 2 | 5 | 21 | 3 | 3 | 37 | 0 | 2 |
| | 401 | 2 | 6 | 4 | 3 | 12 | 15 | 2 | 5 | 3 | 3 |

| | | | | | | | | | | | |
|---|---|---|---|---|---|---|---|---|---|---|---|
| | 18 | 4 | 0 | 5 | 0 | 22 | 9 | 2 | 12 | 1 | 1 |
| | 16 | 3 | 0 | 9 | 8 | 11 | 14 | 3 | 20 | 3 | 6 |
| | 15 | 9 | 4 | 4 | 12 | 10 | 18 | 0 | 22 | 4 | 2 |
| | 14 | 20 | 6 | 4 | 4 | 12 | 11 | 1 | 70 | 3 | 3 |
| | 12 | 4 | 5 | 2 | 2 | 13 | 5 | 0 | 3 | 1 | 3 |
| | 11 | 4 | 0 | 2 | 3 | 12 | 9 | 9 | 6 | 3 | 4 |
| | 404 | 2 | 2 | 1 | 0 | 10 | 3 | 0 | 9 | 0 | 5 |
| | 10 | 0 | 4 | 0 | 6 | 15 | 9 | 5 | 5 | 1 | 3 |
| | 402 | 5 | 0 | 4 | 3 | 7 | 8 | 4 | 3 | 3 | 0 |
| | 8 | 0 | 3 | 5 | 0 | 22 | 25 | 8 | 15 | 1 | 5 |
| Sum | | 82 | 46 | 56 | 68 | 263 | 180 | 49 | 229 | 32 | 47 |
| Average/hive | | 4.10 | 2.30 | 2.80 | 3.40 | 13.15 | 9.00 | 2.45 | 11.45 | 1.60 | 2.35 |
| Control (9 hives) | 27 | 0 | 0 | 0 | 1 | 7 | 2 | 0 | 2 | 1 | 3 |
| | 26 | 48 | 2 | 12 | 4 | 7 | 2 | 0 | 36 | 3 | 5 |
| | 25 | 1 | 2 | 2 | 2 | 5 | 6 | 7 | 2 | 0 | 3 |
| | 24 | 3 | 0 | 3 | 3 | 3 | 2 | 5 | 3 | 2 | 1 |
| | 23 | 1 | 2 | 0 | 0 | 2 | 3 | 1 | 4 | 2 | 2 |
| | 22 | 0 | 4 | 4 | 4 | 1 | 0 | 1 | 3 | 0 | 1 |
| | 21 | 1 | 3 | 2 | 2 | 12 | 4 | 2 | 7 | 3 | 0 |
| | 20 | 7 | 0 | 3 | 3 | 3 | 5 | 8 | 8 | 6 | 3 |
| | 19 | 3 | 1 | 7 | 7 | 23 | 3 | 4 | 3 | 2 | 0 |
| Sum | | 64 | 14 | 33 | 26 | 63 | 27 | 28 | 68 | 19 | 18 |
| Average/hive | | 7.11 | 1.56 | 3.67 | 2.89 | 7.00 | 3.00 | 3.11 | 7.56 | 2.11 | 2.00 |

| | | Dates | | | | | | | |
|---|---|---|---|---|---|---|---|---|---|
| Group | Hive number | Feb. 11 | Feb. 16 | Feb. 19 | Feb. 22 | Feb. 25 | March 1 | March 4 | March 8 |
| MoLi (20 hives ; 19 hives after the lost of hive N°7 in march) | 2 | 2 | 24 | 0 | 3 | 2 | 3 | 7 | 11 |
| | 13 | 0 | 2 | 0 | 0 | 2 | 0 | 7 | 8 |
| | 19 | 3 | 8 | 6 | 5 | 2 | 4 | 15 | 9 |
| | 101 | 5 | 5 | 2 | 1 | 1 | 3 | 17 | 10 |
| | 103 | 3 | 3 | 3 | 6 | 7 | 1 | 19 | 2 |
| | 105 | 1 | 17 | 9 | 4 | 1 | 5 | 10 | 5 |

| | | | | | | | | | |
|---|---|---|---|---|---|---|---|---|---|
| | 106 | 4 | 14 | 3 | 4 | 3 | 9 | 11 | 3 |
| | 112 | 0 | 7 | 4 | 1 | 3 | 4 | 9 | 4 |
| | 7 | 5 | 13 | 6 | 10 | 5 | - | - | - |
| | 401 | 3 | 4 | 5 | 15 | 4 | 2 | 30 | 10 |
| | 18 | 3 | 22 | 9 | 8 | 8 | 7 | 27 | 8 |
| | 16 | 2 | 20 | 14 | 7 | 2 | 5 | 25 | 18 |
| | 15 | 4 | 11 | 11 | 8 | 4 | 3 | 26 | 46 |
| | 14 | 5 | 10 | 9 | 10 | 4 | 3 | 34 | 41 |
| | 12 | 3 | 15 | 8 | 6 | 5 | 6 | 17 | 12 |
| | 11 | 5 | 17 | 6 | 7 | 6 | 4 | 23 | 36 |
| | 404 | 3 | 5 | 3 | 5 | 1 | 12 | 25 | 30 |
| | 10 | 6 | 4 | 8 | 10 | 4 | 5 | 22 | 21 |
| | 402 | 4 | 56 | 3 | 11 | 5 | 4 | 26 | 2 |
| | 8 | 3 | 30 | 20 | 10 | 9 | 15 | 27 | 15 |
| Sum | | 64 | 287 | 129 | 131 | 78 | 95 | 377 | 291 |
| Average/hive | | 3.2 | 14.35 | 6.45 | 6.55 | 3.09 | 5.00 | 19.84 | 15.32 |
| Control (9 hives) | | 1 | 7 | 4 | 2 | 0 | 1 | 18 | 19 |
| | | 3 | 28 | 5 | 4 | 3 | | | |
| | | 3 | 2 | 2 | 8 | 3 | 2 | 15 | 6 |
| | | 1 | 1 | 1 | 3 | 4 | 2 | 19 | 10 |
| | | 2 | 2 | 2 | 2 | 5 | 7 | 17 | 12 |
| | | 1 | 3 | 3 | 2 | 4 | 4 | 20 | 15 |
| | | 0 | 1 | 1 | 6 | 2 | 3 | 14 | 3 |
| | | 3 | 2 | 2 | 0 | 2 | 4 | 28 | 10 |
| | | 0 | 3 | 3 | 7 | 2 | 4 | 16 | 11 |
| Sum | | 14 | 49 | 23 | 34 | 25 | 27 | 147 | 86 |
| Average/hive | | 1.56 | 5.44 | 2.56 | 3.78 | 2.78 | 3.00 | 16.33 | 9.56 |

**Table SIV.28** Count of dead bees in front of the hive during the second period of the experiment (11 march -19 april 2021).

| | | Dates | | | | | | | | | | | |
|---|---|---|---|---|---|---|---|---|---|---|---|---|---|
| Group | Hive number | March 11 | March 15 | March 18 | March 22 | March 26 | March 29 | April 1 | April 5 | April 9 | April 12 | April 15 | April 19 |
| MoLi-B (9 hives. the colony n°7 was lost) | 2 | 2 | 3 | 2 | 24 | 3 | 1 | 2 | 6 | 8 | 10 | 6 | 32 |
| | 13 | 1 | 3 | 0 | 2 | 2 | 1 | 3 | 0 | 0 | 3 | 10 | 3 |
| | 19 | 5 | 6 | 1 | 0 | 3 | 1 | 6 | 2 | 5 | 0 | 4 | 17 |
| | 101 | 3 | 5 | 1 | 4 | 4 | 3 | 0 | 7 | 7 | 2 | 2 | 6 |
| | 103 | 2 | 0 | 1 | 3 | 7 | 24 | 12 | 10 | 10 | 7 | 9 | 4 |
| | 105 | 0 | 0 | 2 | 2 | 2 | 2 | 9 | 2 | 2 | 5 | 4 | 3 |
| | 106 | 1 | 2 | 1 | 2 | 1 | 5 | 11 | 8 | 4 | 12 | 1 | 1 |
| | 112 | 1 | 2 | 1 | 2 | 6 | 0 | 3 | 2 | 1 | 4 | 2 | 1 |
| | 7 | | | | | | | | | | | | |
| | 401 | 15 | 12 | 2 | 2 | 12 | 6 | 5 | 3 | 4 | 5 | 15 | 15 |
| Sum | | 30 | 33 | 11 | 41 | 40 | 43 | 51 | 40 | 41 | 48 | 53 | 82 |
| Average/hive | | 3.33 | 3.67 | 1.22 | 4.56 | 4.44 | 4.78 | 5.67 | 4.44 | 4.56 | 5.33 | 5.89 | 9.11 |
| MoLi-A (10 hives) | 18 | 6 | 8 | 2 | 7 | 8 | 2 | 5 | 7 | 6 | 17 | 10 | 21 |
| | 16 | 7 | 10 | 6 | 9 | 12 | 7 | 6 | 15 | 15 | 9 | 18 | 28 |
| | 15 | 9 | 9 | 4 | 8 | 22 | 12 | 9 | 18 | 6 | 25 | 26 | 23 |
| | 14 | 13 | 8 | 5 | 12 | 13 | 8 | 12 | 16 | 28 | 21 | 60 | 60 |
| | 12 | 4 | 3 | 4 | 2 | 6 | 6 | 7 | 18 | 3 | 11 | 2 | 18 |
| | 11 | 6 | 13 | 0 | 5 | 15 | 4 | 8 | 13 | 8 | 6 | 22 | 20 |
| | 404 | 7 | 19 | 3 | 3 | 18 | 11 | 25 | 17 | 7 | 22 | 28 | 15 |
| | 10 | 5 | 7 | 3 | 19 | 7 | 7 | 7 | 12 | 5 | 7 | 10 | 14 |
| | 402 | 6 | 6 | 12 | 17 | 42 | 41 | 70 | 8 | 13 | 10 | 45 | 40 |
| | 8 | 9 | 11 | 7 | 15 | 6 | 14 | 13 | 6 | 6 | 7 | 37 | 11 |
| Sum | | 72 | 94 | 46 | 97 | 149 | 112 | 162 | 130 | 97 | 135 | 258 | 250 |
| Average/hive | | 7.20 | 9.40 | 4.60 | 9.70 | 14.90 | 11.20 | 16.20 | 13.00 | 9.70 | 13.50 | 25.80 | 25.00 |
| | 27 | 5 | 4 | 4 | 9 | 4 | 9 | 2 | 16 | 15 | 10 | 13 | 15 |

| | | | | | | | | | | | | | |
|---|---|---|---|---|---|---|---|---|---|---|---|---|---|
| Control (8 hives. the colony n°26 was lost) | 26 | | | | | | | | | | | | |
| | 25 | 8 | 1 | 5 | 11 | 17 | 7 | 8 | 13 | 11 | 7 | 24 | 12 |
| | 24 | 1 | 3 | 2 | 6 | 9 | 9 | 9 | 8 | 7 | 6 | 4 | 20 |
| | 23 | 9 | 2 | 3 | 5 | 6 | 10 | 15 | 4 | 6 | 9 | 8 | 27 |
| | 22 | 3 | 6 | 0 | 6 | 3 | 7 | 11 | 5 | 4 | 5 | 9 | 19 |
| | 21 | 5 | 4 | 2 | 5 | 10 | 4 | 2 | 10 | 6 | 6 | 5 | 13 |
| | 20 | 7 | 7 | 3 | 6 | 8 | 3 | 5 | 11 | 4 | 15 | 9 | 17 |
| | 19 | 4 | 4 | 4 | 5 | 11 | 5 | 4 | 9 | 7 | 12 | 16 | 36 |
| Sum | | 42 | 31 | 23 | 53 | 68 | 54 | 56 | 76 | 60 | 70 | 88 | 159 |
| Average/hive | | 5.25 | 3.88 | 2.88 | 6.63 | 8.50 | 6.75 | 7.00 | 9.50 | 7.50 | 8.75 | 11.00 | 19.88 |

**Table SIV.29** Count of varroa fallen during the first period of the experiment (7 dec. 2020 till 8 march 2021).

| | | Dates | | | | | | | | | |
|---|---|---|---|---|---|---|---|---|---|---|---|
| Group | Hive number | Dec. 11 | Dec. 15 | Dec. 18 | Dec. 21 | Dec. 23 | Dec. 28 | Dec. 30 | Jan. 4 | Jan. 8 | Jan. 11 |
| MoLi (20 hives) | 2 | 1 | 1 | 1 | 2 | 0 | 2 | 3 | 2 | 4 | 4 |
| | 13 | 0 | 3 | 1 | 0 | 2 | 0 | 0 | 1 | 0 | 1 |
| | 19 | 3 | 6 | 4 | 3 | 3 | 1 | 4 | 7 | 1 | 3 |
| | 101 | 2 | 4 | 1 | 1 | 1 | 3 | 0 | 2 | 0 | 1 |
| | 103 | 4 | 7 | 1 | 0 | 2 | 2 | 2 | 1 | 0 | 0 |
| | 105 | 24 | 6 | 6 | 4 | 6 | 6 | 3 | 1 | 1 | 6 |
| | 106 | 2 | 0 | 0 | 0 | 0 | 1 | 0 | 2 | 0 | 0 |
| | 112 | 5 | 0 | 0 | 0 | 1 | 1 | 0 | 0 | 0 | 1 |
| | 7 | 1 | 3 | 1 | 1 | 0 | 2 | 3 | 2 | 0 | 0 |
| | 401 | 2 | 0 | 0 | 0 | 1 | 0 | 0 | 1 | 0 | 2 |
| | 18 | 0 | 0 | 0 | 0 | 2 | 9 | 1 | 1 | 2 | 0 |
| | 16 | 0 | 0 | 2 | 1 | 0 | 1 | 0 | 0 | 0 | 0 |
| | 15 | 1 | 1 | 0 | 0 | 0 | 1 | 1 | 1 | 1 | 2 |
| | 14 | 0 | 0 | 0 | 0 | 0 | 1 | 2 | 2 | 1 | 1 |
| | 12 | 0 | 1 | 3 | 4 | 0 | 0 | 3 | 3 | 0 | 1 |

| | | | | | | | | | | | |
|---|---|---|---|---|---|---|---|---|---|---|---|
| | 11 | 2 | 0 | 0 | 0 | 0 | 0 | 0 | 0 | 0 | 2 |
| | 404 | 0 | 0 | 0 | 0 | 0 | 0 | 0 | 0 | 1 | 0 |
| | 10 | 0 | 0 | 1 | 0 | 1 | 0 | 3 | 0 | 0 | 0 |
| | 402 | 6 | 4 | 0 | 0 | 3 | 7 | 5 | 5 | 2 | 0 |
| | 8 | 0 | 1 | 4 | 2 | 2 | 2 | 4 | 2 | 3 | 1 |
| Sum | | 53 | 37 | 25 | 18 | 24 | 39 | 34 | 33 | 16 | 25 |
| Average/hive | | 2.65 | 1.85 | 1.25 | 0.90 | 1.20 | 1.95 | 1.70 | 1.65 | 0.80 | 1.25 |
| Control (9 hives) | 27 | 0 | 0 | 0 | 0 | 0 | 2 | 1 | 2 | 0 | 0 |
| | 26 | 3 | 0 | 0 | 0 | 3 | 24 | 11 | 6 | 9 | 12 |
| | 25 | 2 | 2 | 8 | 4 | 11 | 12 | 7 | 11 | 10 | 13 |
| | 24 | 0 | 0 | 1 | 0 | 0 | 2 | 1 | 0 | 0 | 0 |
| | 23 | 0 | 0 | 0 | 0 | 2 | 1 | 0 | 3 | 0 | 1 |
| | 22 | 0 | 0 | 0 | 0 | 1 | 3 | 2 | 1 | 1 | 1 |
| | 21 | 1 | 0 | 1 | 3 | 6 | 0 | 0 | 0 | 2 | 2 |
| | 20 | 2 | 0 | 0 | 1 | 1 | 0 | 0 | 0 | 0 | 0 |
| | 19 | 3 | 6 | 4 | 2 | 3 | 1 | 4 | 7 | 1 | 0 |
| Sum | | 11 | 8 | 14 | 10 | 27 | 45 | 26 | 30 | 23 | 29 |
| Average/hive | | 1.22 | 0.89 | 1.56 | 1.11 | 3.00 | 5.00 | 2.89 | 3.33 | 2.56 | 3.22 |

| | | Dates | | | | | | | | | |
|---|---|---|---|---|---|---|---|---|---|---|---|
| Group | Hive number | Jan. 13 | Jan. 15 | Jan. 18 | Jan. 20 | Jan. 22 | Jan. 25 | Jan. 28 | Feb. 1 | Feb. 4 | Feb. 4 |
| MoLi (20 hives) | 2 | 0 | 0 | 1 | 0 | 0 | 1 | 0 | 0 | 0 | 0 |
| | 13 | 0 | 0 | 0 | 0 | 1 | 0 | 0 | 0 | 1 | 0 |
| | 19 | 3 | 3 | 3 | 0 | 6 | 9 | 5 | 1 | 3 | 2 |
| | 101 | 0 | 0 | 0 | 0 | 0 | 2 | 0 | 2 | 1 | 1 |
| | 103 | 0 | 0 | 0 | 0 | 0 | 0 | 0 | 0 | 2 | 1 |
| | 105 | 2 | 2 | 1 | 1 | 2 | 1 | 2 | 1 | 0 | 1 |
| | 106 | 3 | 0 | 0 | 0 | 1 | 1 | 0 | 0 | 1 | 0 |
| | 112 | 0 | 0 | 0 | 0 | 1 | 6 | 0 | 2 | 1 | 0 |
| | 7 | 0 | 3 | 2 | 0 | 0 | 1 | 2 | 1 | 0 | 2 |
| | 401 | 0 | 0 | 0 | 2 | 0 | 5 | 0 | 3 | 0 | 0 |
| | 18 | 1 | 3 | 2 | 3 | 0 | 5 | 0 | 1 | 0 | 1 |

| | | | | | | | | | | | |
|---|---|---|---|---|---|---|---|---|---|---|---|
| | 16 | 0 | 0 | 0 | 0 | 1 | 1 | 0 | 0 | 0 | 0 |
| | 15 | 0 | 0 | 0 | 0 | 0 | 2 | 0 | 1 | 2 | 0 |
| | 14 | 0 | 1 | 1 | 0 | 0 | 1 | 2 | 0 | 0 | 0 |
| | 12 | 2 | 1 | 1 | 1 | 1 | 0 | 1 | 0 | 1 | 2 |
| | 11 | 0 | 0 | 0 | 0 | 1 | 0 | 0 | 0 | 0 | 0 |
| | 404 | 0 | 0 | 0 | 0 | 0 | 2 | 0 | 0 | 1 | 1 |
| | 10 | 1 | 0 | 0 | 0 | 0 | 0 | 0 | 0 | 0 | 2 |
| | 402 | 3 | 0 | 0 | 0 | 1 | 0 | 0 | 1 | 0 | 1 |
| | 8 | 0 | 0 | 0 | 1 | 0 | 3 | 2 | 3 | 0 | 0 |
| Sum | | 15 | 13 | 11 | 8 | 15 | 40 | 14 | 16 | 13 | 14 |
| Average/hive | | 0.75 | 0.65 | 0.55 | 0.40 | 0.75 | 2.00 | 0.70 | 0.80 | 0.65 | 0.70 |
| Control (9 hives) | 27 | 0 | 2 | 0 | 0 | 0 | 0 | 2 | 1 | 1 | 0 |
| | 26 | 0 | 1 | 0 | 0 | 2 | 1 | 0 | 5 | 0 | 5 |
| | 25 | 0 | 1 | 5 | 1 | 6 | 3 | 5 | 6 | 9 | 6 |
| | 24 | 4 | 0 | 0 | 1 | 1 | 2 | 3 | 1 | 1 | 0 |
| | 23 | 2 | 0 | 0 | 1 | 0 | 1 | 0 | 1 | 0 | 1 |
| | 22 | 0 | 1 | 1 | 2 | 0 | 0 | 2 | 2 | 1 | 0 |
| | 21 | 2 | 0 | 1 | 0 | 0 | 0 | 1 | 0 | 0 | 0 |
| | 20 | 0 | 2 | 0 | 0 | 0 | 0 | 0 | 1 | 0 | 0 |
| | 19 | 3 | 3 | 3 | 0 | 6 | 9 | 5 | 1 | 3 | 2 |
| Sum | | 11 | 10 | 10 | 5 | 15 | 16 | 18 | 18 | 15 | 14 |
| Average/hive | | 1.22 | 1.11 | 1.11 | 0.56 | 1.67 | 1.78 | 2.00 | 2.00 | 1.67 | 1.56 |

| | | Dates | | | | | | | |
|---|---|---|---|---|---|---|---|---|---|
| Group | Hive number | Feb. 11 | Feb. 16 | Feb. 19 | Feb. 22 | Feb. 25 | March 1 | March 4 | March 8 |
| MoLi (20 hives) | 2 | 2 | 2 | 0 | 0 | 4 | 1 | 0 | 0 |
| | 13 | 0 | 0 | 1 | 0 | 0 | 0 | 1 | 0 |
| | 19 | 0 | 3 | 3 | 2 | 3 | 6 | 3 | 2 |
| | 101 | 0 | 0 | 0 | 0 | 2 | 3 | 0 | 1 |
| | 103 | 2 | 2 | 1 | 1 | 0 | 0 | 0 | 0 |
| | 105 | 4 | 4 | 5 | 1 | 1 | 1 | 4 | 3 |
| | 106 | 4 | 4 | 1 | 1 | 2 | 1 | 2 | 1 |

| | | | | | | | | | |
|---|---|---|---|---|---|---|---|---|---|
| | 112 | 0 | 0 | 0 | 0 | 0 | 1 | 0 | 1 |
| | 7 | 1 | 1 | 2 | 1 | 1 | | | |
| | 401 | 5 | 5 | 0 | 0 | 4 | 0 | 1 | 2 |
| | 18 | 3 | 3 | 2 | 0 | 0 | 2 | 5 | 6 |
| | 16 | 0 | 0 | 0 | 0 | 1 | 0 | 3 | 1 |
| | 15 | 2 | 2 | 2 | 1 | 0 | 1 | 0 | 0 |
| | 14 | 2 | 2 | 0 | 2 | 0 | 0 | 1 | 2 |
| | 12 | 1 | 1 | 2 | 1 | 3 | 0 | 0 | 1 |
| | 11 | 1 | 1 | 2 | 2 | 1 | 2 | 2 | 1 |
| | 404 | 5 | 5 | 3 | 0 | 3 | 0 | 2 | 3 |
| | 10 | 0 | 0 | 2 | 1 | 1 | 0 | 1 | 0 |
| | 402 | 2 | 2 | 0 | 3 | 0 | 3 | 3 | 1 |
| | 8 | 3 | 3 | 0 | 4 | 7 | 1 | 0 | 2 |
| Sum | | 37 | 40 | 26 | 20 | 33 | 22 | 28 | 27 |
| Average/hive | | 1.85 | 2.00 | 1.30 | 1.00 | 1.65 | 1.10 | 1.40 | 1.35 |
| Control (9 hives) | 27 | 3 | 3 | 1 | 0 | 2 | 3 | 2 | 2 |
| | 26 | 4 | 4 | 0 | 5 | 11 | | | |
| | 25 | 4 | 7 | 7 | 10 | 19 | 21 | 5 | 6 |
| | 24 | 0 | 1 | 0 | 1 | 1 | 3 | 3 | 0 |
| | 23 | 1 | 0 | 1 | 4 | 2 | 2 | 0 | 1 |
| | 22 | 2 | 7 | 2 | 1 | 5 | 1 | 2 | 3 |
| | 21 | 0 | 2 | 0 | 2 | 2 | 1 | 1 | 2 |
| | 20 | 3 | 1 | 1 | 1 | 3 | 0 | 1 | 3 |
| | 19 | 0 | 3 | 3 | 2 | 3 | 6 | 1 | 4 |
| Sum | | 17 | 28 | 15 | 26 | 48 | 37 | 15 | 21 |
| Average/hive | | 1.89 | 3.11 | 1.67 | 2.89 | 5.33 | 4.11 | 1.67 | 2.33 |

**Table SIV.30** Count of varroa fallen by ice sugar method on 8th december 2020 and 3rd march 2021.

| Group | Hive number | Dec. 8 | March 3 |
|---|---|---|---|
| MoLi (20 hives) | 2 | 0.4 | 0.2 |
| | 13 | 0.4 | 0.2 |
| | 19 | 0.8 | 1 |
| | 101 | 0.2 | 0.4 |
| | 103 | 0.4 | 0.2 |
| | 105 | 0 | 1.4 |
| | 106 | 0.4 | 0.6 |
| | 112 | 0.6 | 0.4 |
| | 7 | 0.8 | 1 |
| | 401 | 1 | 0.6 |
| | 18 | 1.3 | 0.6 |
| | 16 | 0.6 | 0.6 |
| | 15 | 0 | 0.8 |
| | 14 | 1.6 | 0.2 |
| | 12 | 0.6 | 0.6 |
| | 11 | 1.6 | 0.8 |
| | 404 | 0 | 1.2 |
| | 10 | 0 | 0.2 |
| | 402 | 3.6 | 3.2 |
| | 8 | 0.6 | 1 |
| Average/hive | | 0.75 | 0.76 |
| Control (9 hives) | 27 | 0 | 0.6 |
| | 26 | 1.2 | |
| | 25 | 9.4 | 4.8 |
| | 24 | 0 | 0.2 |
| | 23 | 0 | 0.2 |
| | 22 | 0 | 0.2 |
| | 21 | 0 | 0.4 |
| | 20 | 0 | 0.4 |
| | 19 | 0 | 1 |
| Average/hive | | 1.18 | 0.98 |

**Table SIV.31** Count of varroa fallen during the second period of the experiment (1 march 2020 till 24 may 2021). Note that a critical treatment with oxalic acid took place on 20th avril 2021.

| | | Dates | | | | | | | |
|---|---|---|---|---|---|---|---|---|---|
| Group | Hive number | March 11 | March 15 | March 18 | March 22 | March 26 | March 29 | April 1 | April 5 |
| MoLi-B (9 hives; the colony n°7 was lost) | 2 | 1 | 1 | 0 | 1 | 1 | 1 | 0 | 2 |
| | 13 | 0 | 0 | 0 | 1 | 0 | 2 | 0 | 0 |
| | 19 | 2 | 3 | 1 | 1 | 1 | 1 | 1 | 1 |
| | 101 | 0 | 1 | 1 | 2 | 1 | 0 | 1 | 2 |
| | 103 | 1 | 2 | 3 | 2 | 0 | 1 | 3 | 1 |
| | 105 | 2 | 0 | 2 | 3 | 2 | 0 | 0 | 1 |
| | 106 | 0 | 0 | 0 | 2 | 2 | 1 | 2 | 1 |
| | 112 | 0 | 2 | 1 | 1 | 1 | 1 | 0 | 1 |
| | 7 | | | | | | | | |
| | 401 | 1 | 1 | 5 | 4 | 4 | 1 | 0 | 1 |
| Sum | | 7 | 10 | 13 | 17 | 12 | 8 | 7 | 7 |
| Average/hive | | 0.78 | 1.11 | 1.44 | 1.89 | 1.33 | 0.89 | 0.78 | 0.78 |
| MoLi-A (10 hives) | 18 | 1 | 0 | 2 | 4 | 0 | 1 | 5 | 1 |
| | 16 | 1 | 0 | 1 | 1 | 1 | 2 | 2 | 1 |
| | 15 | 2 | 5 | 3 | 1 | 2 | 1 | 5 | 2 |
| | 14 | 1 | 1 | 1 | 0 | 2 | 3 | 3 | 1 |
| | 12 | 2 | 0 | 2 | 3 | 3 | 1 | 1 | 2 |
| | 11 | 0 | 0 | 0 | 1 | 1 | 1 | 1 | 0 |
| | 404 | 1 | 2 | 0 | 2 | 0 | 7 | 1 | 1 |
| | 10 | 0 | 1 | 1 | 1 | 1 | 1 | 1 | 0 |
| | 402 | 2 | 4 | 4 | 4 | 5 | 8 | 3 | 2 |
| | 8 | 2 | 3 | 0 | 1 | 4 | 4 | 1 | 2 |
| Sum | | 12 | 16 | 14 | 18 | 19 | 29 | 23 | 12 |
| Average/hive | | 1.2 | 1.6 | 1.4 | 1.8 | 1.9 | 2.9 | 2.3 | 1.2 |
| Control | 27 | 2 | 3 | 1 | 5 | 0 | 1 | 1 | 2 |
| | 26 | | | | | | | | |

| | | | | | | | | | |
|---|---|---|---|---|---|---|---|---|---|
| (8 hives. the colony n°26 was lost) | 25 | 7 | 6 | 4 | 14 | 5 | 20 | 23 | 16 |
| | 24 | 5 | 0 | 2 | 3 | 2 | 1 | 3 | 2 |
| | 23 | 0 | 0 | 0 | 1 | 1 | 0 | 0 | 1 |
| | 22 | 3 | 0 | 1 | 5 | 2 | 4 | 4 | 6 |
| | 21 | 1 | 3 | 0 | 4 | 1 | 1 | 2 | 1 |
| | 20 | 0 | 0 | 0 | 3 | 3 | 1 | 1 | 1 |
| | 19 | 3 | 1 | 2 | 3 | 4 | 1 | 2 | 0 |
| Sum | | 21 | 13 | 10 | 38 | 18 | 29 | 36 | 29 |
| Average/hive | | 2.63 | 1.63 | 1.25 | 4.75 | 2.25 | 3.63 | 4.50 | 3.63 |

| | | Dates | | | | | | | |
|---|---|---|---|---|---|---|---|---|---|
| Group | Hive number | April 9 | April 12 | April 15 | April 19 | April 23 | April 27 | April 29 | May 5 |
| MoLi-B (9 hives. the colony n°7 was lost) | 2 | 0 | 0 | 1 | 1 | 4 | 1 | 1 | 1 |
| | 13 | 0 | 0 | 1 | 1 | 6 | 4 | 2 | 1 |
| | 19 | 0 | 0 | 0 | 1 | 5 | 2 | 1 | 2 |
| | 101 | 0 | 1 | 1 | 2 | 8 | 3 | 1 | 1 |
| | 103 | 1 | 1 | 1 | 2 | 9 | 6 | 4 | 2 |
| | 105 | 2 | 0 | 0 | 0 | 7 | 5 | 3 | 5 |
| | 106 | 1 | 1 | 2 | 1 | 11 | 8 | 5 | 3 |
| | 112 | 1 | 2 | 2 | 0 | 3 | 1 | 0 | 0 |
| | 7 | | | | | | | | |
| | 401 | 0 | 0 | 1 | 2 | 21 | 48 | 31 | 24 |
| Sum | | 5 | 5 | 9 | 10 | 74 | 78 | 48 | 39 |
| Average/hive | | 0.56 | 0.56 | 1.00 | 1.11 | 8.22 | 8.67 | 5.33 | 4.33 |
| MoLi-A (10 hives) | 18 | 1 | 6 | 2 | 8 | 75 | 44 | 25 | 30 |
| | 16 | 1 | 1 | 1 | 5 | 27 | 19 | 14 | 13 |
| | 15 | 1 | 2 | 0 | 2 | 25 | 12 | 5 | 3 |
| | 14 | 2 | 3 | 0 | 4 | 43 | 21 | 12 | 9 |
| | 12 | 1 | 2 | 1 | 2 | 58 | 33 | 25 | 21 |
| | 11 | 2 | 1 | 0 | 1 | 20 | 8 | 3 | 1 |
| | 404 | 2 | 2 | 0 | 3 | 33 | 15 | 10 | 7 |

| | | | | | | | | | |
|---|---|---|---|---|---|---|---|---|---|
| | 10 | 0 | 0 | 1 | 1 | 12 | 7 | 3 | 2 |
| | 402 | 12 | 1 | 8 | 20 | 182 | 97 | 14 | 12 |
| | 8 | 0 | 1 | 9 | 1 | 36 | 25 | 7 | 9 |
| Sum | | 22 | 19 | 22 | 47 | 511 | 281 | 118 | 107 |
| Average/hive | | 2.2 | 1.9 | 2.2 | 4.7 | 51.1 | 28.1 | 11.8 | 10.7 |
| Control (8 hives. the colony n°26 was lost) | 27 | 1 | 2 | 0 | 8 | 55 | 24 | 12 | 8 |
| | 26 | | | | | | | | |
| | 25 | 19 | 21 | 14 | 24 | 368 | 207 | 87 | 52 |
| | 24 | 2 | 1 | 5 | 3 | 66 | 29 | 13 | 11 |
| | 23 | 2 | 0 | 1 | 0 | 14 | 10 | 4 | 5 |
| | 22 | 6 | 2 | 1 | 5 | 53 | 44 | 18 | 14 |
| | 21 | 1 | 3 | 4 | 4 | 37 | 21 | 11 | 20 |
| | 20 | 5 | 4 | 7 | 2 | 16 | 11 | 6 | 2 |
| | 19 | 2 | 5 | 3 | 6 | 29 | 20 | 13 | 10 |
| Sum | | 38 | 38 | 35 | 52 | 638 | 366 | 164 | 122 |
| Average/hive | | 4.75 | 4.75 | 4.38 | 6.50 | 79.75 | 45.75 | 20.50 | 15.25 |

| | | Dates | | | | | |
|---|---|---|---|---|---|---|---|
| Group | Hive number | May 7 | May 11 | May 13 | May 17 | May 21 | May 24 |
| MoLi-A (9 hives. the colony n°7 was lost) | 2 | 1 | 0 | 0 | 0 | 1 | 1 |
| | 13 | 0 | 0 | 0 | 0 | 1 | 0 |
| | 19 | 1 | 0 | 0 | 0 | 1 | 13 |
| | 101 | 0 | 0 | 0 | 3 | 1 | 1 |
| | 103 | 1 | 0 | 0 | 1 | 0 | 0 |
| | 105 | 3 | 0 | 0 | 1 | 0 | 2 |
| | 106 | 2 | 1 | 0 | 3 | 0 | 3 |
| | 112 | 0 | 1 | 1 | 1 | 2 | 1 |
| | 7 | | | | | | |
| | 401 | 11 | 2 | 1 | 2 | 2 | 1 |
| Sum | | 19 | 4 | 2 | 11 | 8 | 22 |
| Average/hive | | 2.11 | 0.44 | 0.22 | 1.22 | 0.89 | 2.44 |

| | | | | | | | |
|---|---|---|---|---|---|---|---|
| MoLi-B (10 hives) | 18 | 13 | 5 | 1 | 2 | 5 | 3 |
| | 16 | 9 | 1 | 3 | 1 | 2 | 1 |
| | 15 | 2 | 1 | 0 | 2 | 7 | 0 |
| | 14 | 5 | 0 | 0 | 2 | 0 | 0 |
| | 12 | 12 | 2 | 0 | 1 | 2 | 1 |
| | 11 | 1 | 2 | 1 | 2 | 3 | 1 |
| | 404 | 3 | 0 | 1 | 1 | 4 | 3 |
| | 10 | 1 | 0 | 1 | 1 | 1 | 2 |
| | 402 | 5 | 3 | 2 | 1 | 3 | 1 |
| | 8 | 4 | 0 | 1 | 0 | 2 | 0 |
| Sum | | 55 | 14 | 10 | 13 | 29 | 12 |
| Average/hive | | 5.5 | 1.4 | 1 | 1.3 | 2.9 | 1.2 |
| Control (8 hives. the colony n°26 was lost) | 27 | 4 | 1 | 0 | 0 | 1 | 0 |
| | 26 | | | | | | |
| | 25 | 20 | 6 | 0 | 0 | 6 | 1 |
| | 24 | 5 | 1 | 0 | 0 | 2 | 1 |
| | 23 | 2 | 1 | 0 | 0 | 1 | 0 |
| | 22 | 8 | 2 | 2 | 1 | 1 | 1 |
| | 21 | 10 | 2 | 0 | 0 | 0 | 1 |
| | 20 | 1 | 2 | 0 | 0 | 3 | 7 |
| | 19 | 6 | 3 | 0 | 0 | 8 | 2 |
| Sum | | 56 | 18 | 2 | 1 | 22 | 13 |
| Average/hive | | 7.00 | 2.25 | 0.25 | 0.13 | 2.75 | 1.63 |

**Table SIV.32** Count of Nosema spores / bee during the frist and the second period of the experiment for the different groups of hives.

| | | Dates (period 1) | | | Dates (period 2) | |
|---|---|---|---|---|---|---|
| Group | Hive number | 07/12/2020 | 03/03/2021 | Group | 03/03/2021 | 20/04/2021 |
| MoLi (20 hives at the beginning; 19 hives during the experiment) | 2 | 0 | 36000 | MoLi-B (9 hives) | 36000 | 705000 |
| | 13 | 1476923 | 2257212 | | 2257212 | 3092000 |
| | 19 | 0 | 84000 | | 84000 | 825000 |
| | 101 | 12000 | 51428 | | 51428 | 369231 |
| | 103 | 12000 | 21724 | | 21724 | 1528846 |
| | 105 | 35000 | 24000 | | 24000 | 702000 |
| | 106 | 9000 | 36000 | | 36000 | 12000 |
| | 112 | 6000 | 3000 | | 3000 | 42000 |
| | 7 | | | | | |
| | 401 | 558000 | 11739 | | 11739 | 63000 |
| | - | | - | Average | 280567 | 815453 |
| | 18 | 36000 | 135000 | MoLi-A (10 hives) | 135000 | 1350000 |
| | 16 | 0 | 168000 | | 168000 | 28928 |
| | 15 | 6000 | 12857 | | 12857 | 39000 |
| | 14 | 12000 | 75000 | | 75000 | 24000 |
| | 12 | 672000 | 888462 | | 888462 | 4442308 |
| | 11 | 0 | 6207 | | 6207 | 6000 |
| | 404 | 27000 | 6207 | | 6207 | 1586538 |
| | 10 | 3000 | 15000 | | 15000 | 66000 |
| | 402 | 3000 | 1032692 | | 1032692 | 30000 |
| | 8 | 12000 | 532500 | | 532500 | 237000 |
| Average | | 151575 | 284054 | Average | 287193 | 780977 |
| Control (9 hives at the beginning; 8 hives | 27 | 1131000 | 1269231 | Control (8 hives) | 1269231 | 2803462 |
| | 26 | - | - | | | |
| | 25 | 0 | 27000 | | 27000 | 1338462 |
| | 24 | 6000 | 1471154 | | 1471154 | 180000 |
| | 23 | 6000 | 6000 | | 6000 | 39000 |

| | | | | | | |
|---|---|---|---|---|---|---|
| during the experiment) | 22 | 0 | 1921154 | | 1921154 | 2000000 |
| | 21 | 0 | 1500000 | | 1500000 | 1920000 |
| | 20 | 0 | 138000 | | 138000 | 403846 |
| | 19 | **6624000** | **6166250** | | **6166250** | **9230770** |
| Average | | 661381 | 1562349 | Average | 1562349 | 2239443 |
| Average without hive n°19 | | 119325 | 904648 | Average without hive n°19 | 904648 | 1240681 |
