## Supplementary material for "Food supplementation with molybdenum complexes improves honey bee health": Tracking Mo in hives

### *Supporting Information*

#### Part V. Tracking Mo in hives after the feeding

- **V.1** Objectives of the study
- **V.2** Materials and methods
- **V.3** Results
  - V.3.1 Molybdenum-content in honey
  - V.3.2 Molybdenum-content in bee larvae
  - V.3.3 Molybdenum-content in worker bees
  - V.3.4 Molybdenum-content in wax
- **V.4 Conclusions**

### Part V. Tracking Mo in hives after the feeding

#### V.1 Objectives of the study

In part IV, we demonstrated that feeding hives with **Na-$Mo_2O_4$-EDTA** or **Li-$Mo_2O_4$-EDTA** can have positive effects notably on colony growth, on hygienic behavior and on infection with *Varroa* mites. These effects are observed over several months, whereas feeding is carried out either all at once or over a short period of time. These results therefore raise questions about the mode of action and persistence of these complexes in bee colonies.

In this study, we will focus on **Na-$Mo_2O_4$-EDTA**. In order to better understand the behaviour of this complex in the hives after the feeding, the quantity of the element molybdenum has been tracked during 2 months in worker bees, in larvae, in the wax, in the "honey" stored in the brood frames, and in the bee bread prepared by bees from pollen stored in the brood frames. This study aimed to answer the following questions:

- How do bees consume the syrup containing **Na-$Mo_2O_4$-EDTA**?
- How does it pass into adult bees and larvae?
- What is the external source of molybdenum?
- Why has molybdenum supplementation with **Na-$Mo_2O_4$-EDTA** a positive effect on the bees?

#### V.2 Materials and methods

##### V.2.1 Animals

| | |
|---|---|
| Taxonomic Group: | Hymenoptera, Apidae |
| Species: | *Apis mellifera mellifera* L. |
| Source: | APINOV apiary SAS in La Rochelle, France. |
| Experimental unit: | The experimental unit considered is a healthy colony of honey bees (*Apis mellifera*). All colonies are composed with sister queens of the year laying eggs (same genetic line), 2 frames of brood, 1 frame of honey and 1 embossed wax frame. The size of the frame corresponds to Dadant or Langstrot standards. We evaluate the population at the start of the study to be between 6000 and 9000 honey bees. |

##### V.2.2 Husbandry conditions

The good health of a colony can be defined as the presence of a queen, and the absence of illness and/or intoxication.

Hives assigned to the study were arranged in such a way as to avoid any form of bee drift (for example: the entrance to the hives were turned following the 4 cardinal points).

Each beehive was fed by directly pouring syrup in the feeder of the hive. Each hive received 2L of syrup with the corresponding **Na-$Mo_2O_4$-EDTA** concentration (1-20 $mg.L^{-1}$) at the beginning of the experiment (D0) and again 2L after 7 days (D7). So, each beehive received 4 liters of syrup in one week and 4-80 mg of **Na-$Mo_2O_4$-EDTA** in total.

##### V.2.3 Diets

Each beehive has been fed with a syrup containing **Na-$Mo_2O_4$-EDTA** or with syrup without **Na-$Mo_2O_4$-EDTA**. The syrup was prepared in different concentrations:

- Syrup without **Na-$Mo_2O_4$-EDTA** (**control**)
- Syrup with the recommended dose of **Na-$Mo_2O_4$-EDTA** (**RD**): 2 $mg.L^{-1}$
- 2 times less the recommended dose (**RD/2**): 1 $mg.L^{-1}$
- 2 times the recommended dose (**2RD**): 4 $mg.L^{-1}$
- 10 times the recommended dose (**10RD**): 20 $mg.L^{-1}$

##### V.2.6 Experimental syrup diet

The control diet is composed of the sugar syrup Apistar® (Icko Apiculture; saccharose 34%, glucose 33% and fructose 33%, density: 1.45), widely used in beekeeping.

To prepare the experimental syrup diets, the **Na-$Mo_2O_4$-EDTA** complex was dissolved in the Apistar® syrup, according to each target concentration. The control and experimental syrup diets were visually identical and prepared in 500 litre tanks (Figure SV.1). The syrup was homogenised during 2 hours using a circulation pump.

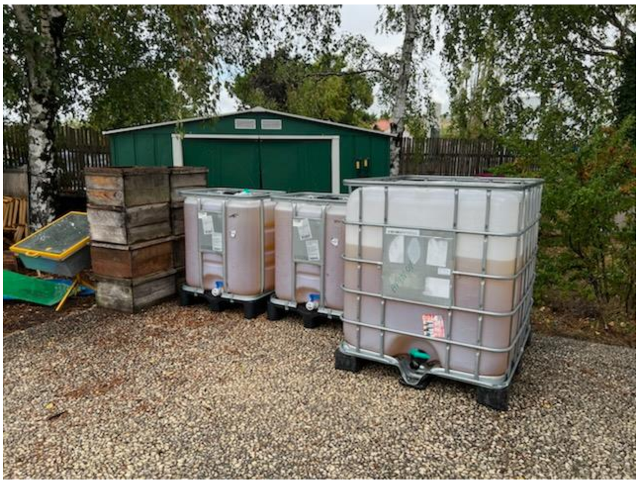

*Figure SV.1. Pictures of the 500 liters tanks used for the preparation of syrups containing*

***Na-$Mo_2O_4$-EDTA***

The concentration of **Na-$Mo_2O_4$-EDTA** within the syrups was controlled by ICP-MS by the LEAV Laboratory, La Roche sur Yon, France. The results are gathered in Table SV.1. The discrepancies between the calculated and the measured values of molybdenum concentration in the syrup are due to mistakes made on the big volume of the tanks. For the sake of simplicity, in the rest of this document we will use the target values to designate the various batches.

***Table SV.1*** *Concentration of Mo in syrups*

| Syrup name | Total $Na\text{-}Mo_2O_4$-EDTA concentration in the hive (mg) | $Na\text{-}Mo_2O_4$-EDTA concentration in the syrup (mg/L) | Molybdenum concentration in the syrup (mg/kg) | |
|---|---|---|---|---|
| | | | Calculated | Measured |
| **Control**[(1)] | 0mg | 0mg/L | 0ppm | <0.050 |
| **RD/2**[(2)] | 4mg | 1mg/L | 0.2ppm | 0.20 |
| **RD**[(3)] | 8mg | 2mg/L | 0.4ppm | 0.344 |
| **2RD**[(4)] | 16mg | 4mg/L | 0.8ppm | 0.64 |
| **10RD**[(5)] | 80mg | 20mg/L | 4ppm | 4.5 |

[(1)] Syrup without « **$Na\text{-}Mo_2O_4$-EDTA** » ; [(2)] 2 times less than the recommended dose; [(3)] Syrup corresponding to the recommended dose ; [(4)]2 times the recommended dose ; [(5)]10 times the recommended dose.

Once the syrups prepared, 2L were firstly applied in each hive at D0, and 2 additional liters added after 7 days (D7), for a total of 4L at the corresponding concentration per beehive. Thus, the concentration at the recommended dose (RD) in the beehive is about **8 mg.beehive$^{-1}$**, and 4, 16, 80 mg.beehive$^{-1}$, respectively for RD/2, 2RD and 10RD syrups.

##### V.2.7 Study design

###### 1°) Schedule

The sampling of the worker bees, the larvae, the wax, honey (food reserve, see Figure SV.2) and bee bread has begun in May 2022 and has ended at the end of June 2022. The analysis of molybdenum levels in the samples has started in September 2022.

###### 2°) Sampling process

Food reserve samples were collected from a total of 40 hives (8 hives for each of the 5 **$Na\text{-}Mo_2O_4$-EDTA** syrup concentrations: control, RD/2, RD, 2RD, and 10RD), and wax, bee bread, larvae and working bee samples were collected from a total of 16 hives each (8 hives for each of the 2 **$Na\text{-}Mo_2O_4$-EDTA** syrup concentrations: control and RD), all randomly chosen.

**$Na\text{-}Mo_2O_4$-EDTA** syrup concentrations were given to 22 hives for the control and the RD, 10 hives for the 2RD and RD/2, and 8 hives for the 10RD modality, all randomly chosen and numbered.

Honey (food reserve), wax, pollen, and larvae samples: for each hive, one sample was composed of 3 sampling points taken on 2 different frames. Each sample amounted to:

- 50 g of honey (food reserve) in 20 cm$^2$ of wax. In the beehive, bees store honey on the outer parts of the frames (red circle on the Figure SV.2). Honey and wax samples were collected in that area. Note that honey (food reserve) was taken from the brood frames for the tests, whereas beekeepers take it from the honey super, which reduces the risk of taking honey that is still in the syrup state). The honey/food reserve samples were manually extracted from the wax and wax samples were obtained after washing samples with water to remove honey/food sample.
- 20 cm$^2$ of comb containing beebread,

- 20 $cm^2$ of comb containing 6-8 days old larvae.

Worker samples: For each hive, one sample was composed of 50 g of worker bees, following the AFNOR standard XPX 43-909 (Active biomonitoring of the environment using honeybees). Worker bees were taken directly from the frame (see Figure SV.2).

For all samples collected, the molybdenum content was analyzed using the ICP-MS method according to ISO/DIS 17294-1:2022 by the accredited laboratory LEAV in La Roche sur Yon, France. For each ICP-MS measurement the uncertainty level was ± 0.02 ppm ($\mu g_{Mo}$/g).

***Table SV.2*** *Summary of the study design and samples collected*

| Beekeeping matrices | Number of hives sampled by modality | Date of sampling | Quantity sampled per hive | Analysis |
|---|---|---|---|---|
| Honey | Control: 8<br>RD/2: 8<br>RD: 8<br>2RD: 8<br>10RD: 8 | D0, D28, D42 and D56 | 50 g | Molybdenum residues ICP-MS |
| Wax | Control: 8<br>RD: 8 | D0, D56 | 20 $cm^2$ | |
| Beebread | Control: 8<br>RD: 8 | D0, D56 | 20 $cm^2$ | |
| Larvae | Control: 8<br>RD: 8 | D0, D14, D28, D56 | 20 $cm^2$ | |
| Workers | Control: 8<br>RD: 8 | D0, D14, D28, D56 | 50 g | |

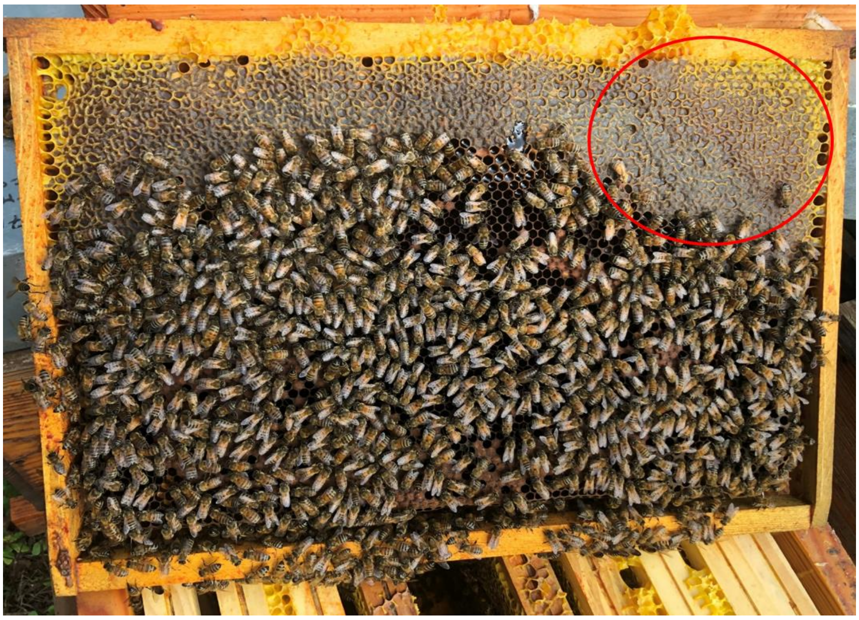

***Figure SV.2*** *Picture of a typical frame in beehives. The red circle indicates the zone where food reserves are stored (usually honey).*

#### V.3 Results

##### V.3.1 Molybdenum-content in honey

The Table SV.3 gathers the results of Mo analyses in honey/food reserve samples. Figure SV.3 represents the average molybdenum content found in honey/food reserve with the standard deviation, while Figure SV.4 presents the comparison of Mo content between the group treated with **Na-$Mo_2O_4$-EDTA** at the recommended dose (RD) and the control group.

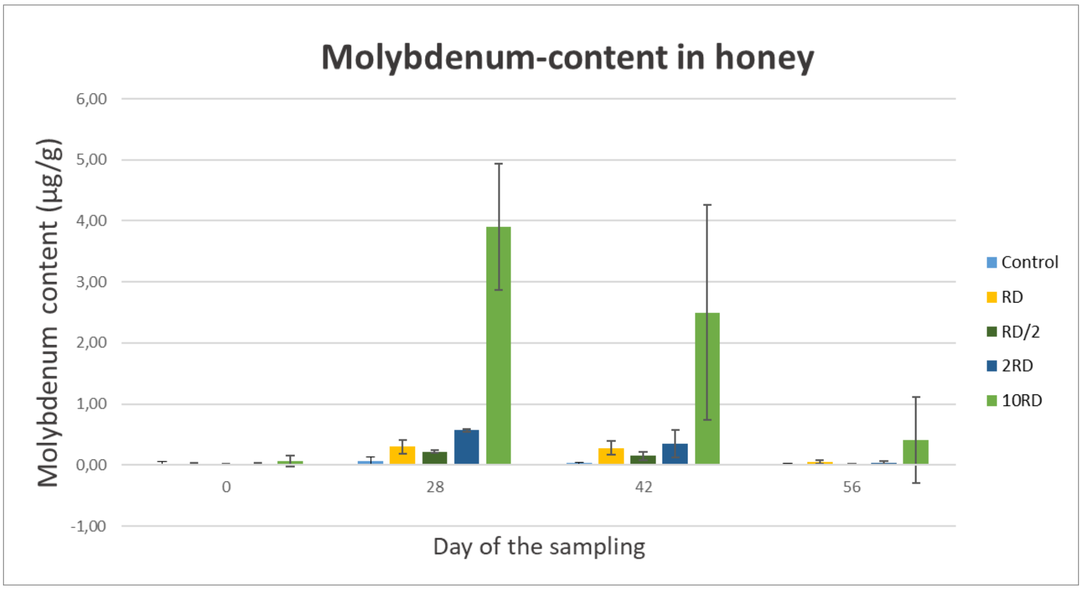


***Figure SV.3.*** *Molybdenum content in honey/food reserve in µg/g sampled at D0, D28, D42 and D56 for control and the following* **Na-$Mo_2O_4$-EDTA** *concentrations: RD/2, RD, 2RD and 10RD.*

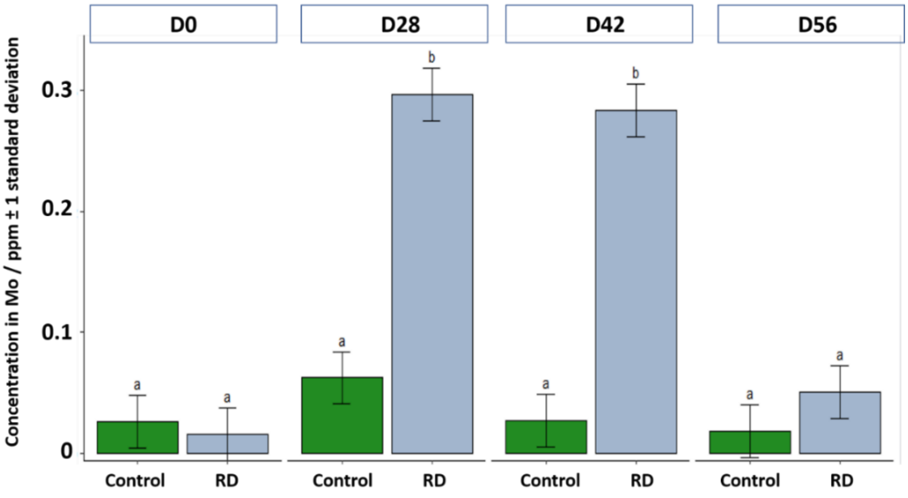


***Figure SV.4.*** *Molybdenum content in honey/food reserve in µg/g sampled at D0, D28, D42 and D56 for the control group and the group fed with* **Na-$Mo_2O_4$-EDTA** *at the recommended dose (RD – 8 mg/beehive).*

At D0, the level of molybdenum in honey/food reserve was below 0.1 ppm and sometimes below the limit of quantification (<0.01 ppm), except for hive 284 of the 10RD batch, which was abnormally high at the beginning of the experiment (see Table SV.3).

Samples of honey/food reserve were taken at D28, D42 and D56. At D28 and D42 the molybdenum content of honey/food reserve matches well to the level of molybdenum in the syrup given to beehives in batches RD/2, RD, 2RD and 10RD, while the molybdenum level remained low in the control batch. It evidences that the collected samples did not correspond to honey produced by the colony, but rather to the treatment syrup collected by the bees and stored around the brood (to feed the colony).

At D56, the level of molybdenum was below 0.1 ppm in all batches, except for 3 beehives (248, 251, 255) in the 10RD batch. This suggests that these samples corresponded now to honey produced by the bees and not to the stored syrup.

From this data, **we can conclude that after feeding the colonies, the syrup is stored within the beehives and consumed by the colonies over 1 month and a half**. This slow consumption permits to feed several generations of bees and allows a prolonged effect of **Na-$Mo_2O_4$-EDTA** within the colonies (It takes 21 days for a bee to develop from the egg stage to emergence as an adult).

***Table SV.3.*** *Molybdenum-content in honey/food reserve in ppm (µg$_{Mo}$/g). For each ICP-MS measurement, uncertainty is ± 0.02 ppm (µg$_{Mo}$/g). LOQ: Limit Of Quantification*

| Batch (Molybdenum content in syrup) | Hive number | D0 | D28 | D42 | D56 |
|---|---|---|---|---|---|
| Control (only syrup) | 259 | 0.011 | 0.022 | 0.019 | 0.016 |
| | 266 | < 0.010 (LOQ) | 0.021 | 0.053 | 0.015 |
| | 263 | 0.018 | 0.100 | 0.020 | 0.023 |
| | 264 | < 0.010 (LOQ) | 0.019 | 0.013 | 0.021 |
| | 265 | 0.020 | 0.018 | 0.014 | 0.012 |
| | 256 | 0.022 | 0.024 | 0.036 | 0.016 |
| | 247 | 0.017 | 0.200 | 0.042 | 0.019 |
| | 330 | 0.098 | 0.093 | 0.018 | 0.020 |
| | **Average** | **0.026** | **0.062** | **0.027** | **0.018** |
| RD/2 (Molybdenum 0.20 ppm) | 269 | < 0.010 (LOQ) | 0.26 | 0.22 | 0.038 |
| | 261 | 0.013 | 0.20 | 0.19 | 0.011 |
| | 253 | < 0.010 (LOQ) | 0.21 | 0.18 | 0.012 |
| | 327 | < 0.010 (LOQ) | 0.24 | 0.11 | 0.012 |
| | 320 | < 0.010 (LOQ) | 0.21 | 0.14 | 0.011 |
| | 315 | < 0.010 (LOQ) | 0.20 | 0.039 | 0.013 |
| | 283 | 0.014 | 0.21 | 0.084 | 0.018 |

| | | | | | |
|---|---|---|---|---|---|
| | 321 | 0.014 | 0.24 | 0.22 | 0.021 |
| | **Average** | **0.011** | **0.22** | **0.15** | **0.017** |
| RD (Molybdenum 0.34 ppm) | 260 | < 0.010 (LOQ) | 0.33 | 0.35 | 0.11 |
| | 262 | < 0.010 (LOQ) | 0.25 | 0.29 | 0.037 |
| | 258 | < 0.010 (LOQ) | 0.36 | 0.32 | 0.015 |
| | 252 | < 0.010 (LOQ) | 0.37 | 0.31 | 0.045 |
| | 250 | < 0.010 (LOQ) | 0.36 | 0.30 | 0.071 |
| | 329 | < 0.010 (LOQ) | 0.33 | 0.021 | 0.081 |
| | 344 | < 0.010 (LOQ) | 0.044 | 0.33 | 0.014 |
| | 328 | 0.050 | 0.33 | 0.35 | 0.029 |
| | **Average** | **0.015** | **0.30** | **0.28** | **0.050** |
| 2RD (Molybdenum 0.64 ppm) | 270 | 0.020 | 0.57 | 0.33 | 0.088 |
| | 267 | < 0.010 (LOQ) | 0.57 | 0.45 | 0.020 |
| | 257 | < 0.010 (LOQ) | 0.58 | 0.53 | 0.010 |
| | 325 | 0.014 | 0.53 | 0.45 | 0.017 |
| | 318 | < 0.010 (LOQ) | 0.56 | 0.025 | 0.014 |
| | 313 | 0.042 | 0.59 | 0.64 | 0.059 |
| | 322 | < 0.010 (LOQ) | 0.61 | 0.017 | 0.015 |
| | 270 | 0.020 | 0.57 | 0.33 | 0.011 |
| | **Average** | **0.017** | **0.57** | **0.35** | **0.029** |
| 10RD (Molybdenum 4.5 ppm) | 255 | 0.017 | 4.2 | 4.1 | 2.0 |
| | 251 | 0.074 | 3.9 | 1.7 | 0.20 |
| | 249 | 0.064 | 1.6 | 4.4 | 0.049 |
| | 248 | 0.017 | 4.9 | 4.1 | 0.85 |
| | 331 | 0.014 | 4.3 | 1.6 | 0.067 |
| | 310 | 0.013 | 4.6 | 3.8 | 0.015 |
| | 312 | 0.014 | 3.4 | 0.52 | 0.059 |
| | 284 | 0.28 | 4.4 | 0.042 | 0.054 |
| | **Average** | **0.061** | **3.9** | **2.5** | **0.41** |

##### V.3.2 Molybdenum-content in bee larvae

The Mo-contents in bee larvae at D0, D14, D28 and D56 are gathered in Table SV.4, while Figure SV.5 presents the comparison of Mo content of larvae between the group treated with **Na-$Mo_2O_4$-EDTA** at the recommended dose RD and the control group.

***Table SV.4.*** *Molybdenum-content in bee larvae in ppm ($\mu g_{Mo}/g$). For each ICP-MS measurement the uncertainty is ± 0.02 ppm ($\mu g_{Mo}/g$). nd: not determined because of lack of larvae when samples were collected.*

| Batch | Hive number | D0 | D14 | D28 | D56 |
|---|---|---|---|---|---|
| | 259 | 0.18 | 0.23 | 0.23 | 0.25 |
| | 266 | 0.28 | 0.11 | 0.13 | 0.17 |
| | 264 | 0.30 | 0.17 | 0.21 | 0.21 |
| | 265 | 0.35 | 0.15 | 0.19 | 0.19 |
| Control | 256 | 0.30 | 0.12 | 0.24 | 0.16 |
| | 263 | nd | 0.17 | 0.23 | nd |
| | 330 | 0.23 | 0.26 | 0.25 | nd |
| | 247 | 0.19 | 0.13 | nd | 0.098 |
| | **Average** | **0.26** | **0.17** | **0.211** | **0.18** |
| | 260 | 0.23 | 0.33 | 0.44 | 0.22 |
| | 258 | 0.24 | 0.23 | 0.32 | 0.19 |
| | 252 | 0.22 | 0.22 | 0.40 | 0.19 |
| | 250 | 0.25 | 0.19 | 0.30 | 0.18 |
| | 329 | 0.34 | 0.19 | 0.28 | 0.13 |
| RD | 344 | 0.20 | 0.28 | 0.28 | 0.19 |
| | 328 | 0.17 | 0.29 | 0.24 | 0.42 |
| | 262 | 0.24 | 0.18 | nd | 0.29 |
| | **Average**<br>**(variation *vs* control)** | **0.24**<br>**(-8%)** | **0.24**<br>**(+42%)** | **0.323**<br>**(+53%)** | **0.226**<br>**(+26%)** |

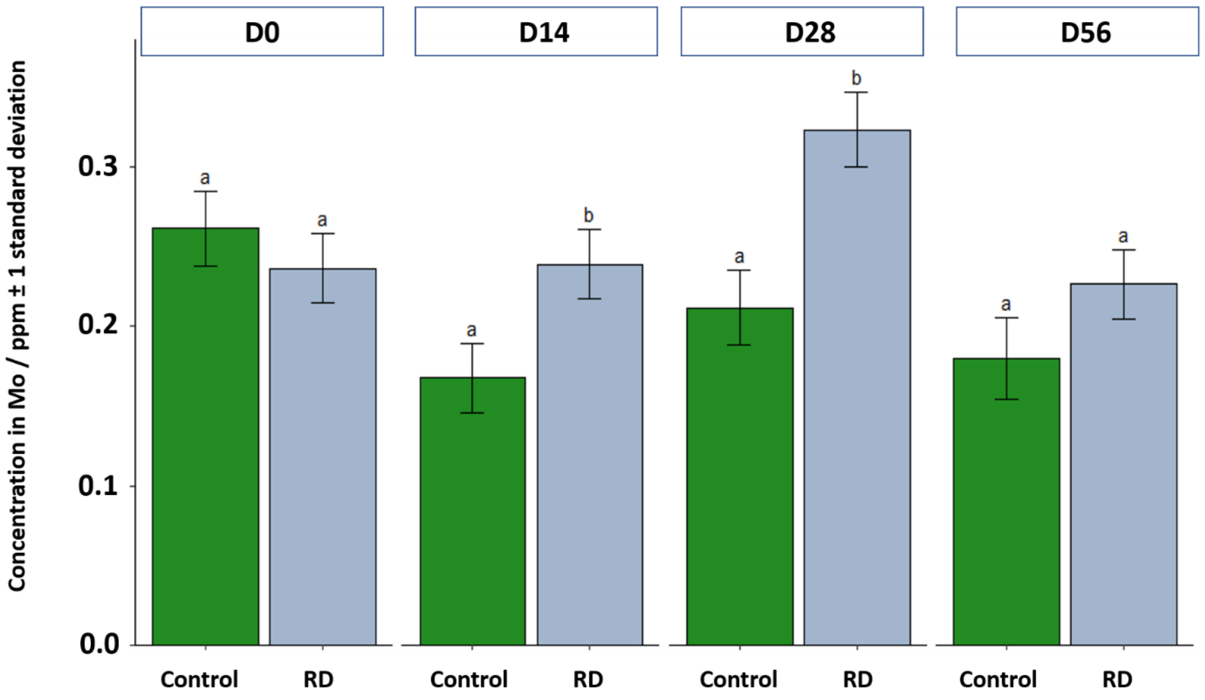


***Figure SV.5:*** *Molybdenum content in larvae in µg/g sampled at D0, D14, D28, D56 for the control group and the group treated with* **Na-$Mo_2O_4$-EDTA** *at the recommended dose (RD – 8 mg/beehive).*

At D0, the level of molybdenum in the control group and in the 8mg/beehive was very similar: 0.24-0.26 ppm. After two weeks at D14, the level decreased in the control group (0.17 on average), while it remained stable in the 8 mg/beehive group (0.24 ppm on average) resulting in a difference of +42% for molybdenum level in the 8 mg/beehive group (RD).

This difference increased again at D28 with reaching +53% molybdenum in the 8mg/beehive group compared to the control group. This suggest that the syrup stored within the beehive is given to the larvae and that later molybdenum is assimilated.

At D56, the difference between the two groups decreased according to the results obtained for honey.

In conclusion, it is clear that **the syrup containing molybdenum is consumed and assimilated by bee larvae, which provokes a significant increase, especially at D28 ($F_{1,14}$ = 8.98, p-value < 0.01), of the level of molybdenum in larvae during at least 2 months**.

##### V.3.3 Molybdenum-content in worker bees

The Mo-contents in worker bees at D0, D14, D28 and D56 are gathered in Table SV.5, while Figure SV.6 presents the comparison of Mo content of worker bees between the group treated with **Na-$Mo_2O_4$-EDTA** at the recommended dose and the control group.

***Table SV.5.*** *Mo-content in worker bees in ppm ($\mu g_{Mo}/g$). For each measurement with the ICP-MS method the uncertainty is ±0.02 ppm ($\mu g_{Mo}/g$).*

| Batch | Hive number | D0 | D14 | D28 | D56 |
|---|---|---|---|---|---|
| Control | 259 | 0.37 | 0.27 | 0.33 | 0.22 |
| | 266 | 0.36 | 0.28 | 0.29 | 0.26 |
| | 263 | 0.49 | 0.42 | 0.39 | 0.36 |
| | 264 | 0.30 | 0.75 | 0.34 | 0.30 |
| | 265 | 0.41 | 0.26 | 0.23 | 0.23 |
| | 256 | 0.30 | 0.30 | 0.28 | 0.21 |
| | 247 | 0.35 | 0.42 | 0.34 | 0.16 |
| | 330 | 0.38 | 0.24 | 0.27 | 0.18 |
| | **Average** | **0.37** | **0.367** | **0.309** | **0.24** |
| RD | 260 | 0.46 | 0.78 | 0.53 | 0.38 |
| | 262 | 0.3 | 0.63 | 0.50 | 0.27 |
| | 258 | 0.52 | 0.63 | 0.64 | 0.26 |
| | 252 | 0.56 | 0.73 | 0.63 | 0.20 |
| | 250 | 0.32 | 0.49 | 0.53 | 0.22 |
| | 329 | 0.38 | 0.50 | 0.55 | 0.27 |
| | 344 | 0.44 | 0.54 | 0.37 | 0.22 |
| | 328 | 0.39 | 0.64 | 0.70 | 0.25 |
| | **Average (variation *vs* control)** | **0.421 (+14%)** | **0.617 (+68%)** | **0.556 (+80%)** | **0.259 (+8%)** |

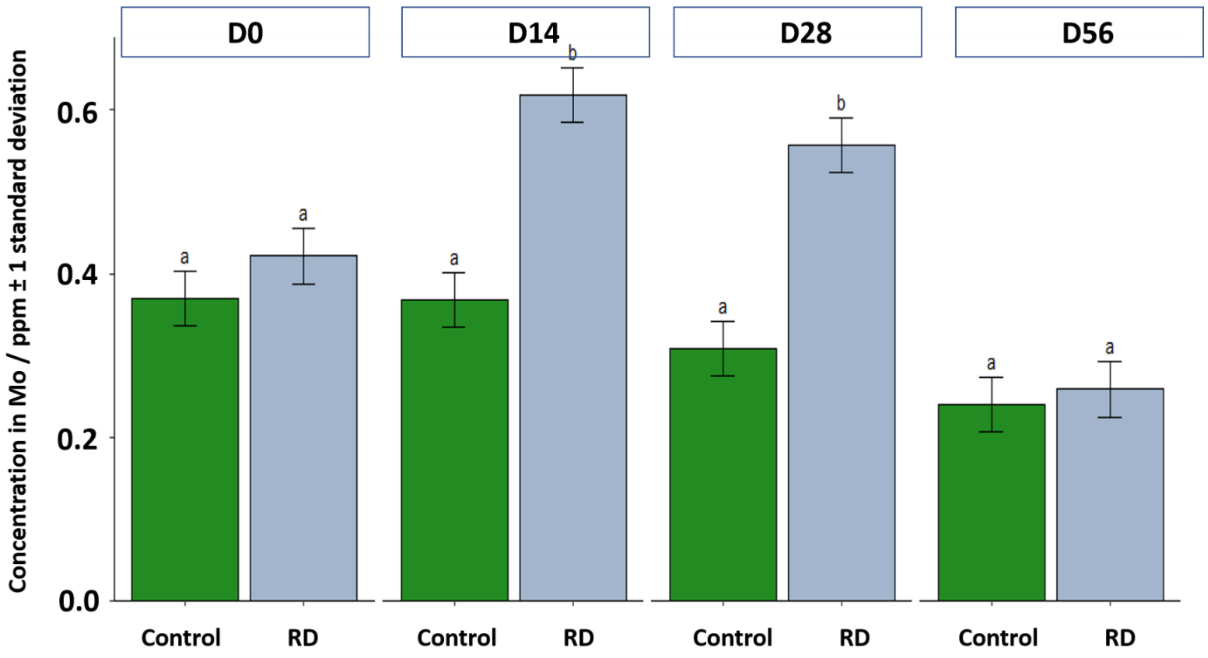


***Figure SV.6.*** *Molybdenum content in worker bees in µg/g sampled at D0, D14, D28, D56 for the control group and the group treated with* **Na-$Mo_2O_4$-EDTA** *at the recommended dose (RD – 8 mg/beehive).*

At D0, the level of molybdenum in the control group and in the 8mg/beehive group was of the same order: around 0.4 ppm.

After two weeks at D14, this level remained stable in the control group, while it increased by +68% in the 8mg/beehive group (0.62 ppm). This strong difference indicates that workers consumed the syrup and assimilated molybdenum from the syrup.

At D28, the level of molybdenum decreased in the control group (0.31 ppm), but remained high in the 8mg/beehive group (0.56 ppm), representing +80% compared to the control. Interestingly, some of these workers may have been larvae at D14, for which we already saw an increase in molybdenum content. Therefore, the level of Mo in these D28 workers may have been acquired at both larval and adult stage.

At D56, the levels of molybdenum in the control and in the 8mg/beehive groups were again of the same order, and showed lower values than at D0 in both groups.

In conclusion, it is clear that the syrup containing molybdenum is consumed by worker bees, which provokes an important and significant increase in the level of molybdenum in bees, especially at D14 and D28 ($F_{3,42} = 8.95$, p-value < 0.001)). This effect disappears after 2 months. **It confirms that feeding with Na-$Mo_2O_4$-EDTA-containing syrup has an effect on bees during 2 months, and therefore on several generations of bees.**

##### V.3.4 Molybdenum-content in wax

Material flows are supposed to be slow for wax and wax contents should not vary rapidly. Moreover, we evidenced that the feeding syrup is found until D42 in brood cells. Some wax samples were then collected at D0 and at D56. The Mo-contents in wax at D0 and D56 are presented in Table SV.6 and Figure SV.7 for hives treated with **Na-$Mo_2O_4$-EDTA** at the recommended dose and control hives.

***Table SV.6.*** *Mo-content in wax in ppm (μg$_{Mo}$/g). For each ICP-MS measurement the uncertainty is ±0.02 ppm (μg$_{Mo}$/g).*

| Batch | Hive number | D0 | D56 |
|---|---|---|---|
| Control | 259 | 0.21 | 0.20 |
| | 266 | 0.13 | 0.12 |
| | 264 | 0.36 | 0.12 |
| | 265 | 0.14 | 0.12 |
| | 256 | 0.15 | 0.08 |
| | 263 | 0.33 | 0.15 |
| | 330 | 0.28 | 0.08 |
| | 247 | 0.34 | 0.05 |
| | **Average** | **0.24** | **0.12** |
| RD | 260 | 0.25 | 0.21 |
| | 258 | 0.21 | 0.09 |
| | 252 | 0.21 | 0.06 |
| | 250 | 0.29 | 0.17 |
| | 329 | 0.17 | 0.06 |
| | 344 | 0.20 | 0.12 |
| | 328 | 0.26 | 0.43 |
| | 262 | 0.15 | 0.14 |
| | **Average** | **0.22** | **0.16** |

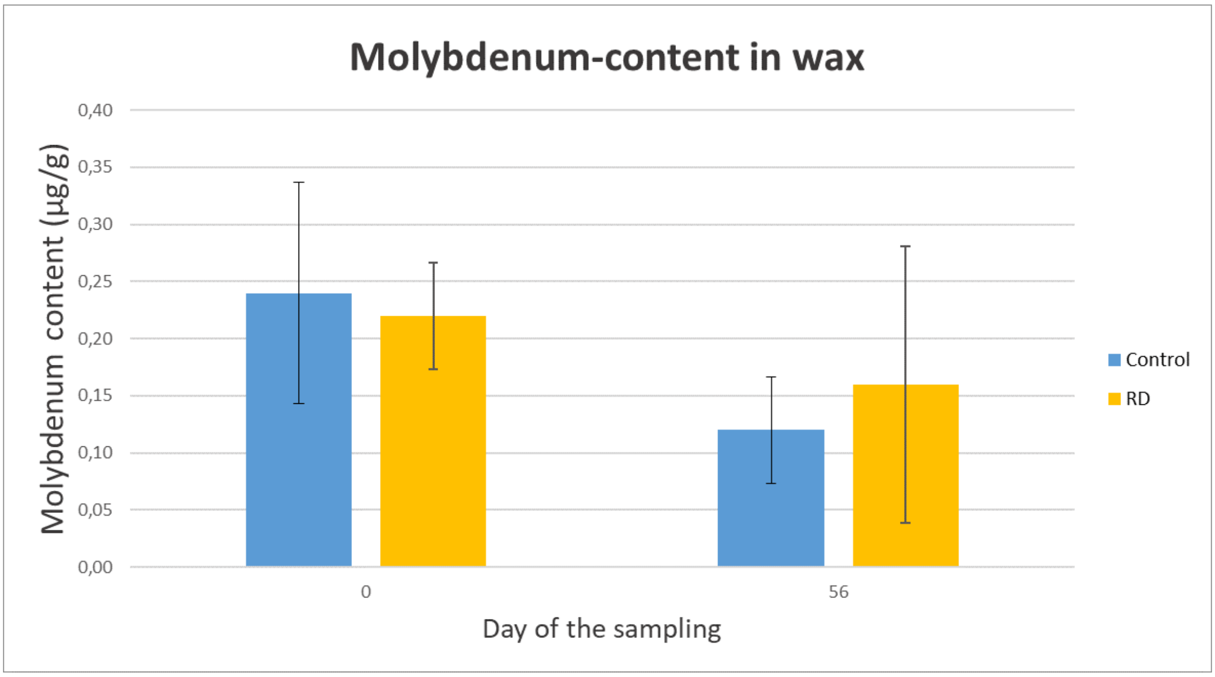

***Figure SV.7:*** *Molybdenum content in wax in µg/g sampled at D0 and D56 from control hives and hives treated with* **Na-$Mo_2O_4$-EDTA** *at the recommended dose (RD – 8 mg/beehive).*

Samples of wax were collected in the control and 8mg/beehive group at D0 and D56. The level of molybdenum was determined by ICP-MS to verify if our product **Na-$Mo_2O_4$-EDTA** passes into wax.

At D0, the analyses reveal that wax naturally contains some molybdenum. The level was similar for the two groups: 0.24-0.22 ppm. At D56, the levels were lower, 0.12 and 0.16 ppm respectively, but still very similar between the two groups.

We can conclude that the complex **Na-$Mo_2O_4$-EDTA** does not pass into the wax after the feeding. The fact that the **Na-$Mo_2O_4$-EDTA** molecule is anionic and highly soluble in water (>1000 g/L) and poorly soluble in octanol (<2.5 mg/L) is coherent with a low lipophilicity and therefore a low affinity for wax.

##### V.3.5 Molybdenum-content in bee bread

The bee bread is produced by bees within the beehives. It corresponds to fermented pollen and it is used to feed bee larvae as well as workers. Bee bread constitutes a source of minerals and trace elements for bees.

The Mo-contents in bee bread at D0 and D56 are shown in Table SV.7 and Figure SV.8 for hives treated with **Na-$Mo_2O_4$-EDTA** at the recommended dose and control hives.

Beebread samples were collected from beehives of the control and of the 8mg/beehive group at D0 and D56. Unfortunately, at D56 (in July), because of the drought around the apiary, the sources of pollen were scarce, and it was not possible to find bee bread in all the beehives. This explains the missing values in Table SV.7 at D56 for some beehives.

***Table SV.7*** *Mo-content in bee bread in ppm (μg$_{Mo}$/g). For each ICP-MS measurement the uncertainty is ±0.02 ppm (μg$_{Mo}$/g).*

| Batch | Hive number | D0 | D56 |
|---|---|---|---|
| Control | 259 | 0.19 | - |
| | 266 | 0.16 | 0.095 |
| | 264 | 0.40 | - |
| | 265 | 0.21 | - |
| | 256 | 0.19 | 0.15 |
| | 263 | 0.39 | - |
| | 330 | 0.29 | - |
| | 247 | 0.55 | - |
| | **Average** | **0.30** | **0.12** |
| RD | 260 | 0.42 | - |
| | 258 | 0.49 | 0.11 |
| | 252 | 0.39 | - |
| | 250 | 0.48 | 0.11 |
| | 329 | 0.17 | - |
| | 344 | 0.30 | - |
| | 328 | 0.28 | - |
| | 262 | 0.24 | 0.09 |
| | **Average** | **0.35** | **0.10** |

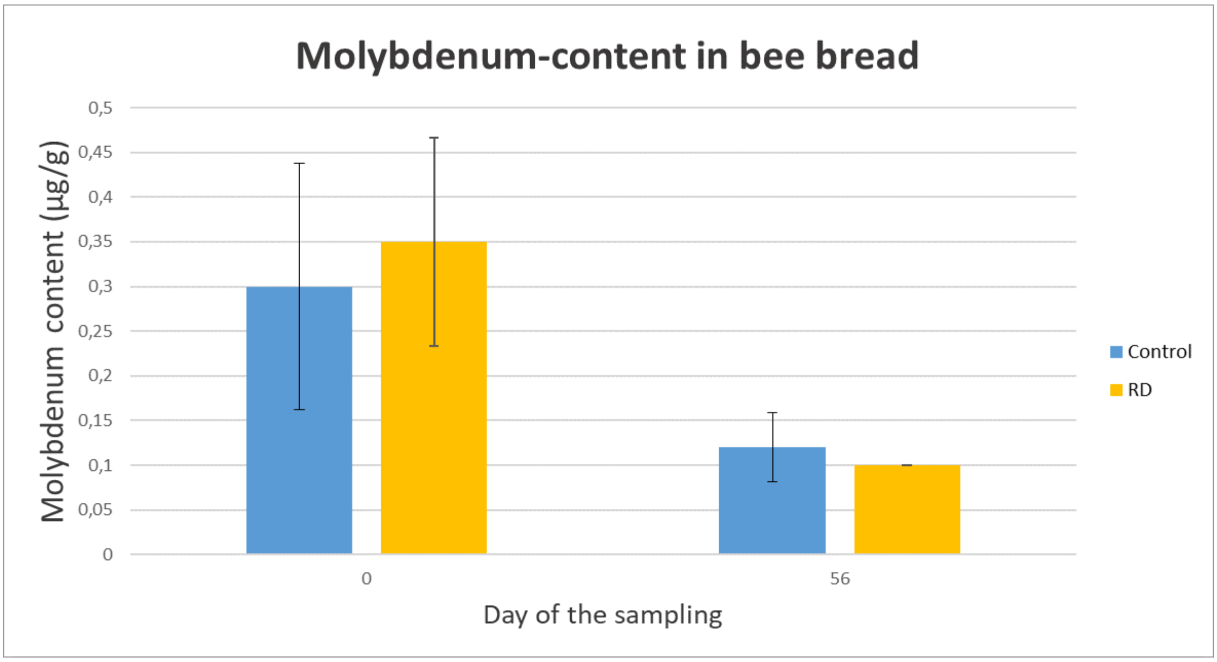


***Figure SV.8.*** *Molybdenum content in bee-bread in µg/g sampled at D0 and D56 from control hives and hives treated with* **Na-$Mo_2O_4$-EDTA** *at the recommended dose (RD – 8 mg/beehive).*

Interestingly, the entry of molybdenum inside the beehive thanks to the pollens and through bee bread at D0 is found in the 0.30-0.35 ppm range for control and 8mg/beehive groups. This value is of the same order as the Mo level in larvae and worker bees at D0.

At D56, the quantity of pollen collected fell dramatically because of the drought and the level of molybdenum decreased drastically to 0.10-0.12 ppm.

This lower level of molybdenum in bee bread at D56 compared to D0 is coherent with the decrease in molybdenum level observed in the bee larvae and worker bees of the control groups from D0 to D56. **It indicates that some climatic conditions, like drought (high temperature and low rainfall), when the sources of the trace element molybdenum become rarer in pollen, decrease the availability of molybdenum for bees. Feeding with Na-$Mo_2O_4$-EDTA can balance this deficiency.**

#### V.4 Conclusions

Thanks to the monitoring of molybdenum levels after feeding hives with **Na-$Mo_2O_4$-EDTA** supplemented sugar syrup, we conclude that:

- **Na-$Mo_2O_4$-EDTA** does not pass into the wax after the feeding. It is due to the anionic state and the high solubility in water of the **Na-$Mo_2O_4$-EDTA** molecule, and is coherent with a low lipophilicity and therefore a low affinity for wax.

- The molybdenum content in honey/food reserves corresponds to the level of molybdenum found in the **Na-$Mo_2O_4$-EDTA** syrup given to the beehives. After 2 months of supplementation, the levels of molybdenum decrease drastically, meaning that the syrup stored in the combs is now replaced by real honey.

After feeding the colonies, **Na-$Mo_2O_4$-EDTA** is stored within the beehive and consumed by the colonies for about 1 month and a half.

- After supplementation with **Na-$Mo_2O_4$-EDTA**, the level of molybdenum in the larvae and in the workers increases during 2 months and then returns to basal values, which means that the syrup is consumed by the bees and the larvae for 2 months. The syrup is stored in the hive for 1.5-2 months and is then replaced by real honey. Therefore, the supplementation of the hives with the syrup can benefit several generations of bees. Workers consumed directly the syrup with **Na-$Mo_2O_4$-EDTA** in the feeder, inducing a fast increase in their molybdenum values. The larvae are fed indirectly with **Na-$Mo_2O_4$-EDTA**. The working bees stored the syrup with **Na-$Mo_2O_4$-EDTA** in brood cells and larvae could also assimilate the **Na-$Mo_2O_4$-EDTA**.

- The bee bread is a compound made by bees out of pollen inside the hives. molybdenum is naturally found in the bee bread, which suggests that pollen is bees' natural source of molybdenum. This explains why molybdenum contents in bee bread were not impacted by **Na-$Mo_2O_4$-EDTA** supplementation.

- During the tested period from May to July, pollen became scarcer and gradually poorer in molybdenum. Indeed, at the end of the study, the quantity of pollen collected fell dramatically because of the drought, and it was not possible to find bee bread in all the hives. At the same time, the level of molybdenum also decreased drastically in the bee bread. Its show that molybdenum is less available naturally for bees when drought appears. Supplementation with **Na-$Mo_2O_4$-EDTA** undoubtedly makes up for this deficiency and allows bees to be supplemented with molybdenum without being hit by the drought consequences.
