## Supplementary material for "Food supplementation with molybdenum complexes improves honey bee health": Assimilation of Mo-complexes by honeybees

k) Hellenic Agriculture Org. "DIMITRA", Institute of Animal Science, Department of Apiculture, 63200 Nea Moudania, Greece,

### *Supporting Information*

#### Part VI. Assimilation of Mo by bees : ICP-MS/ICP-OES studies

### Part VI. Assimilation of Mo by bees : ICP-MS/ICP-OES studies

Following the mortality studies described in part III.4 of this « supporting information », dead bees were isolated according to their date of death and stored in a freezer at -20°C. This large number of bee samples provides an opportunity to evaluate the assimilation of Mo complexes by the bees, and in particular to see the variation in the concentration of the element Mo, but also Na for the complex **Na-$Mo_2O_4$-EDTA** and Li for the complex **Li-$Mo_2O_4$-EDTA**, as a function of feeding duration and complex concentration in the syrup. This is the aim of Part VI.

#### VI.1 Experimental procedures

##### 1°) Animal preparation

Dead bees from experiment III.4 (chronic toxicity on bees) were collected as the experiment progressed and stored in the freezer at -20°C. Each replicate involves 50 bees and each dose was tested 3 times, i.e. 150 bees per control or per dose of molybdenum complex. The dead bees were then divided into 3 batches (Periods 1, 2, 3) according to the duration of feeding to form batches of 30 to 50 individuals as given in Table SVI.1. For simplicity, the periods were defined identically for all the concentrations of **Na-$Mo_2O_4$-EDTA** (and control) on the one hand, and for all concentrations of **Li-$Mo_2O_4$-EDTA** complexes (and control) on the other (see Figures SVI.1 and SVI.2).

***Table SVI.1****. Time of treatment for the 3 batches used for ICP-MS analysis. Periods were determined in order to have around 40-50 individuals per analysis and in order to analyze the 3 distinct phases observed in the experiment detailed in Part III.4.*

| Complexes | Period 1 | Period 2 | Period 3 |
|---|---|---|---|
| Na-$Mo_2O_4$-EDTA | **Days 22 - 47** | **Days 47-55** | **Days 55-74** |
| Li-$Mo_2O_4$-EDTA | **Days 17 - 41** | **Days 41-55** | **Days 55-71** |

After cryopreservation bees were placed on a bed of dry ice to separate the three tagma: head, thorax and abdomen. The samples were stored at -80°C.

##### 2°) ICP-MS analysis

The determination of Mo and Na contents (**Na-$Mo_2O_4$-EDTA** treatment) or Mo and Li contents (**Li-$Mo_2O_4$-EDTA** treatment) was performed by ICP-MS. The samples of head, thorax or abdomen were first freeze-dried, then ground, then mineralized before being analyzed by ICP-MS or by ICP-OES depending on the levels of Mo, Li, or Na. These experiments were performed by the UT2A Laboratory in Pau (www.ut2a.fr/fr/)

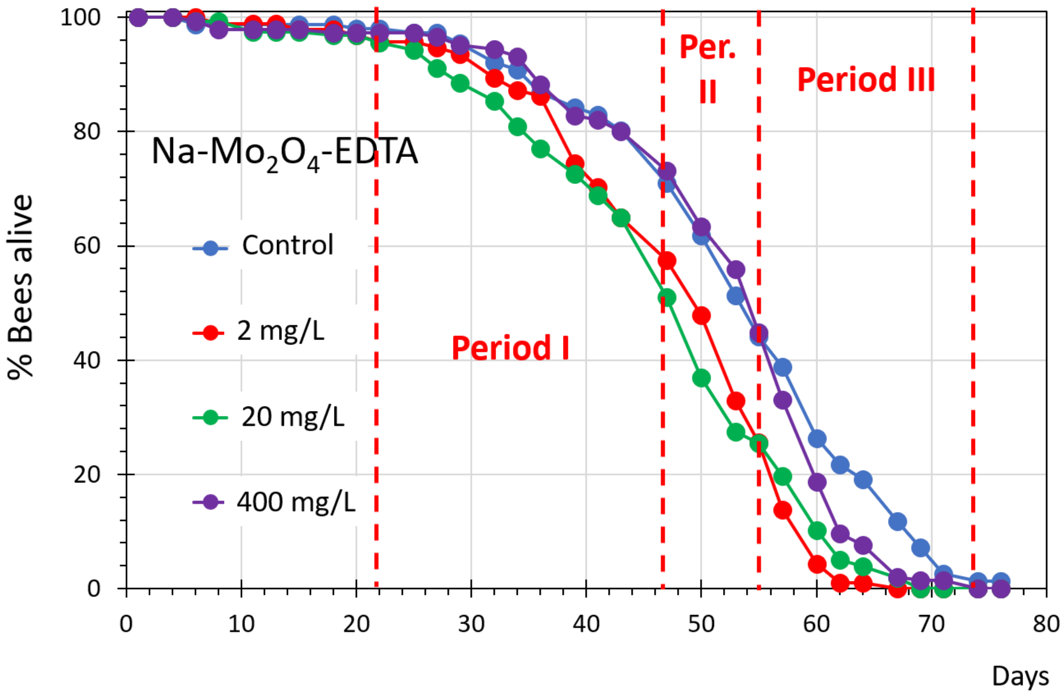


***Figure SVI.1.*** *Mortality curves obtained with complex* ***Na-Mo₂O₄-EDTA*** *(see Part III.4 of supporting information) highlighting the 3 periods defined to follow Mo and Na contents in bees.*

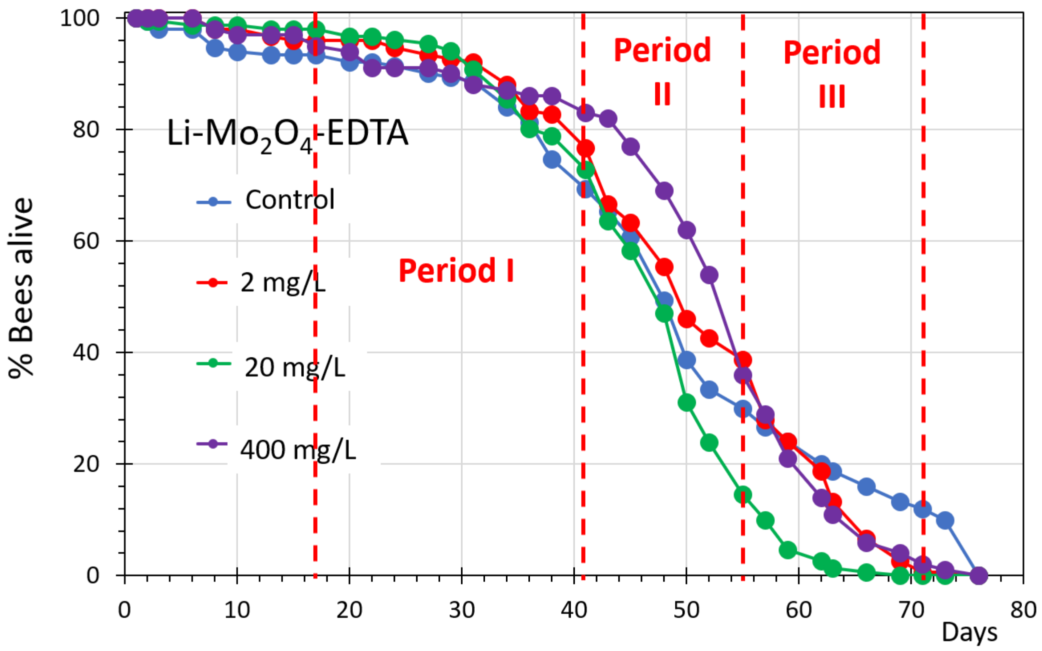


***Figure SVI.2.*** *Mortality curves obtained with complex* ***Li-Mo₂O₄-EDTA*** *(see Part III.4 of supporting information) highlighting the 3 periods defined to follow Mo and Li contents in bees.*

#### VI.2 Results and discussion

The results are gathered in Tables SVI.2 and SVI.3 for **Na-$Mo_2O_4$-EDTA** and **Li-$Mo_2O_4$-EDTA**, respectively.

Concerning **Na-$Mo_2O_4$-EDTA**, the level of Na is naturally high in bees and feeding with solution of **Na-$Mo_2O_4$-EDTA** does not provokes any significant increase in the level of Na found in bees excepted with a concentration of 400 mg/L. The Mo-content was relatively low in the head, thorax and abdomen of control bees compared to the value expected from Part I (0.4 ppm, see Part I of this SI). The head and abdomen appear richer in Mo than the thorax.

Table SVI.2 unambiguously evidences that the level of Mo increases in head, thorax and abdomen when the concentration of **Na-$Mo_2O_4$-EDTA** increases in the feeding syrup but for each concentration the level seems to reach a plateau which does not depend on the period and thus on the duration of feeding.

It is important to note that the level of Mo reaches very extremely high levels in the abdomen (maximum: 1600 ppm). This result is explained in particular by the fact that in cages, bees defecate very little (they usually do so in flight), so that these high levels may not be located in abdomen tissues, but rather in the faeces.

***Table SVI.2***. *Mo and Na contents in ppm (µg/g of dried bee) in the head, thorax and abdomen of bees treated with the complex* ***Na-$Mo_2O_4$-EDTA*** *in mortality tests*

| Modality | Periods | Head | | Thorax | | Abdomen | |
|---|---|---|---|---|---|---|---|
| | | Mo (ppm) | Na (ppm) | Mo (ppm) | Na (ppm) | Mo (ppm) | Na (ppm) |
| Control | I | 0.20 ± 0.01 | 720 ± 40 | 0.12 ± 0.01 | 350 ± 5 | 0.18 ± 0.02 | 177 ± 2 |
| | II | 0.20 ± 0.01 | 650 ± 50 | 0.13 ± 0.01 | 375 ± 3 | 0.21 ± 0.03 | 200 ± 10 |
| | III | 0.17 ± 0.01 | 680 ± 30 | 0.11 ± 0.01 | 330 ± 10 | 0.19 ± 0.01 | 160 ± 20 |
| **Na-$Mo_2O_4$-EDTA** 2mg/L | I | 0.70 ± 0.01 | 728 ± 9 | 0.6 ± 0.1 | 380 ± 10 | 14 ± 1 | 141 ± 4 |
| | II | 0.63 ± 0.01 | 710 ± 10 | 0.9 ± 0.3 | 400 ± 60 | 17 ± 1 | 176 ± 1 |
| | III | 0.42 ± 0.01 | 710 ± 6 | 0.7 ± 0.1 | 370 ± 30 | 12 ± 3 | 190 ± 10 |
| **Na-$Mo_2O_4$-EDTA** 20mg/L | I | 2.02 ± 0.01 | 761 ± 8 | 2.06 ± 0.01 | 340 ± 20 | 71 ± 9 | 150 ± 10 |
| | II | 1.33 ± 0.01 | 661 ± 3 | 2.12 ± 0.04 | 430 ± 20 | 80±20 | 210±20 |
| | III | 2.15 ± 0.75 | 760 ± 10 | 2.08 ± 0.08 | 409 ± 8 | 77±4 | 250±20 |
| **Na-$Mo_2O_4$-EDTA** 400mg/L | I | 30 ± 1 | 830 ± 5 | 51 ± 5 | 500 ± 60 | 1230±300 | 410±60 |
| | II | 28 ± 1 | 834 ± 4 | 80 ± 25 | 530 ± 20 | 1600±200 | 530±10 |
| | III | 23 ± 1 | 1003 ± 4 | 43 ± 15 | 540 ± 6 | 1500±100 | 510±50 |

***Table SVI.3**. Mo and Li contents in ppm (µg/g of dried bee) in the head, thorax and abdomen of bees treated with the complex Li-$Mo_2O_4$-EDTA in mortality tests*

| Modality | Periods | **Head** | | **Thorax** | | **Abdomen** | |
|---|---|---|---|---|---|---|---|
| | | Mo (ppm) | Li (ppm) | Mo (ppm) | Li (ppm) | Mo (ppm) | Li (ppm) |
| Control | I | 0.66 ± 0.07 | 0.06 ± 0.01 | 0.27 ± 0.01 | 0.03 ± 0.01 | 0.68 ± 0.01 | 0.20 ± 0.01 |
| | II | 0.46 ± 0.02 | 0.04 ± 0.01 | 0.26 ± 0.01 | 0.03 ± 0.01 | 0.66 ± 0.01 | 0.15 ± 0.01 |
| | III | 0.40 ± 0.01 | 0.04 ± 0.01 | 0.23 ± 0.01 | 0.03 ± 0.01 | 0.75 ± 0.01 | 0.17 ± 0.01 |
| **Li-$Mo_2O_4$-EDTA** 2mg/L | I | 1.86 ± 0.01 | 1.06 ± 0.01 | 3.07 ± 0.02 | 1.31 ± 0.02 | 134 ± 5 | 10.0 ± 0.09 |
| | II | 1.48 ± 0.01 | 0.84 ± 0.02 | 2.31 ± 0.04 | 0.93 ± 0.01 | 127 ± 1 | 11 ± 2 |
| | III | 1.62 ± 0.04 | 1.11 ± 0.02 | 2.00 ± 0.04 | 1.23 ± 0.01 | 121 ± 3 | 9 ± 2 |
| **Li-$Mo_2O_4$-EDTA** 20mg/L | I | 4.36 ± 0.04 | 1.77 ± 0.01 | 7.81 ± 0.04 | 2.27 ± 0.02 | 180 ± 10 | 12 ± 1 |
| | II | 3.72 ± 0.01 | 2.35 ± 0.01 | 5.50 ± 0.07 | 2.53 ± 0.02 | 235.6 ± 0.4 | 17 ± 4 |
| | III | 3.39 ± 0.02 | 1.58 ± 0.01 | 3.45 ± 0.07 | 1.80 ± 0.02 | 209 ± 6 | 16.7 ± 0.6 |
| **Li-$Mo_2O_4$-EDTA** 400mg/L | I | 31.9 ± 0.1 | 10.89 ± 0.08 | 63.1 ± 0.4 | 16.3 ± 0.1 | 2600 ± 300 | 190 ± 20 |
| | II | 37.6 ± 0.2 | 14.0 ± 0.1 | 58.6 ± 0.2 | 18.87 ± 0.05 | 3600 ± 200 | 263 ± 9 |
| | III | 35.03 ± 0.03 | 16.5 ± 0.2 | 104.0 ± 0.3 | 24.9 ± 0.1 | 3520 ± 10 | 259.6 ± 0.4 |

Concerning **Li-$Mo_2O_4$-EDTA**, the natural level of Lithium is very low (below 0.1 ppm) in the head and thorax and around 0.2 ppm in the abdomen of control bees. For this series of bees, the natural Mo-content in the head, thorax and abdomen was relatively high: 0.7 ppm in the head and abdomen and around 0.3 ppm in the thorax. The head and abdomen appear again about two times richer in Mo than the thorax.

Table SVI.3 unambiguously evidences that the levels of Mo and Li both increase in the head, thorax and abdomen when the concentration of **Li-$Mo_2O_4$-EDTA** increases in the feeding syrup. As observed with **Na-$Mo_2O_4$-EDTA,** for each concentration of **Li-$Mo_2O_4$-EDTA** the level seems to reach a plateau which does not depend on the period and thus on the duration of feeding.

As for **Na-$Mo_2O_4$-EDTA**, the levels of Mo and Li reached very high levels in the abdomen. As observed above, such high levels may be due to concentration in the faeces and not in the abdomen tissue.

Figures SVI.3-SVI.5 present the correlation of the Mo-content versus Li-contents in each tagma. Note that such correlation curves are not possible for **Na-$Mo_2O_4$-EDTA** since the level of Na is naturally very high in bees.

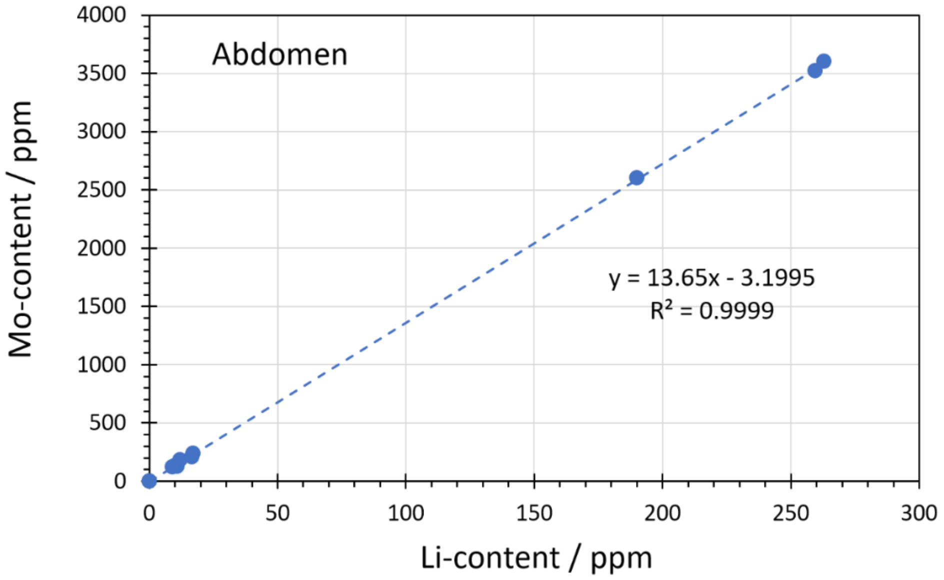


***Figure SVI.3**. Correlation between Mo- and Li-contents in the abdomen of bees fed with **Li-Mo$_2$O$_4$-EDTA**.*

Interestingly, the Mo-content linearly correlates with Li-content in the abdomen. In the complex **Li-Mo$_2$O$_4$-EDTA**, two $Li^+$ cations are present for two Mo(+V) atoms. The molecular masses are of course different (95.94 g/mol for Mo, 6.94 g/mol for Li) with a ratio $MM_{(Mo)}/MM_{(Li)}$ = 13.82. This value is highly similar to the slope of the line drawn in Figure SV.3, which means that in the abdomen, the metals Li and Mo accumulate in agreement with the formula of the complex **Li-Mo$_2$O$_4$-EDTA**.

The figures SVI.4-SVI.5 represent the correlation of Mo- and Li-contents in the thorax and head, respectively.

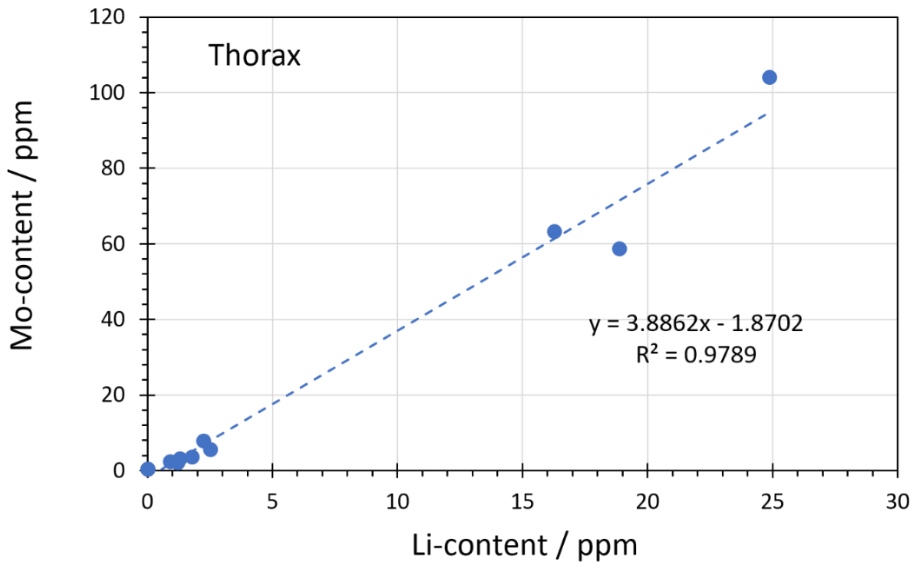


***Figure SVI.4**. Correlation of Mo- and Li-contents in the thorax of bees fed with **Li-Mo$_2$O$_4$-EDTA**.*

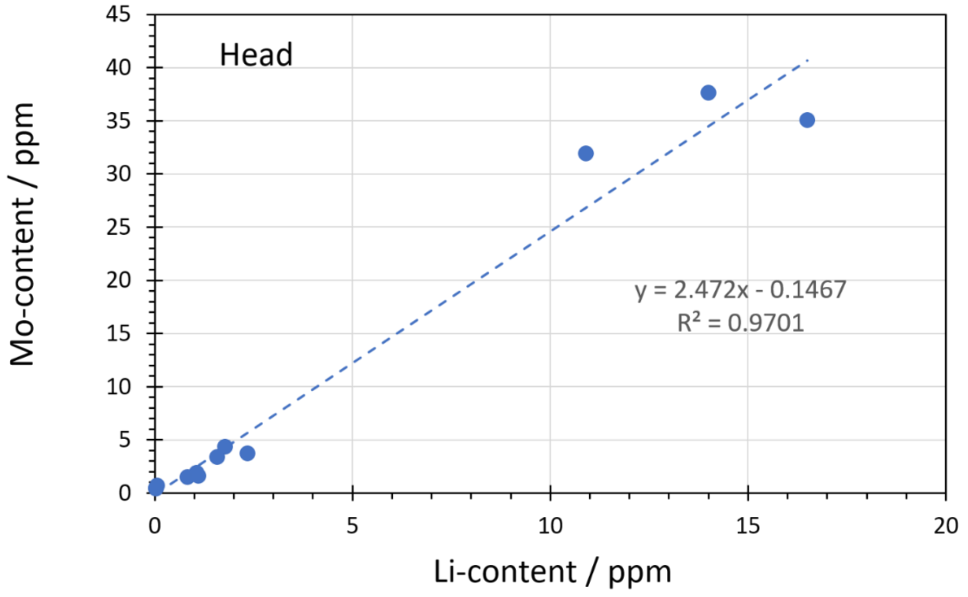


***Figure SVI.5**. Correlation of Mo- and Li-contents in the head of bees fed with **Li-$Mo_2O_4$-EDTA**.*

For these two tagma, the slope significantly differs from the expected value 13.82. It means that the assimilation of Li is stronger than Mo in these tagma. For thorax, a slope of coefficient 3.88 means that 3.56 $Li^+$ are assimilated by bees for 1 Mo atom (13.82/3.88). In the head, this value increases to 5.59 $Li^+$ assimilated per Mo atom.

#### VI.3. Conclusions

These experiments evidence that the complexes **Na-$Mo_2O_4$-EDTA** and **Li-$Mo_2O_4$-EDTA** are assimilated by bees. The analyses show it unambiguously for the head, the thorax and the abdomen. In the abdomen, Mo and Li/Na probably accumulate strongly in faeces, making it difficult to draw any conclusions on the uptake of the two complexes in abdomen tissues. The levels of Mo increase in head and thorax, even with a low concentrated solution of complexes.

Interestingly, the levels of Mo:

- seems dependent on the initial content of Mo in bees;
- depends on the concentration of the complex solution
- does not depend on the duration of feeding (periods 1,2, 3)
- can reach very high values in the head, > 30 ppm, i.e. 100 times higher than the initial level.

Concerning the $Na^+$ cations, the analysis of bees evidence that the Na-content is very different for head, thorax and abdomen. The head contains ca 700 ppm of Na, vs 350 ppm in thorax and 150-200 « only » in the abdomen. By feeding with the 400 mg/L solution, this level in the head increases again up to 800-1000 ppm.

Finally, by feeding with **Li-$Mo_2O_4$-EDTA**, the level of $Li^+$ increases up to 5.6 times higher than the level of Mo, notably in the head, which could have deleterious effects on the health of bees if the level increases up to several mM concentration, which is far from the level reached in our study. [1]

[1] S. Sevin, V. Bommuraj, Y.R. Chen, O. Afik, S. Zarchin, S. Barel, O.C. Arslan, B. Erdem, H. Tutun, J.A. Shimshoni, *Pest Management Science*, **2022**, 78(11), 4507-4516.
