## Supplementary material for "Food supplementation with molybdenum complexes improves honey bee health": X-ray fluorescence studies at synchrotron SOLEIL

### *Supporting Information*

#### Part VII. Synchrotron radiation X-ray fluorescence microscopy

VII.1 Experimental procedures
- 1°) Animal preparation
- 2°) Cross sections
- 3°) X-Ray Fluorescence spectra.

VII.2 Bee Cross-section results

VII.3 Results of synchrotron measurements
- VII.3.1 X-Ray Fluorescence spectrum of resin on $Si_3N_4$ membrane
- VII.3.2 X-Ray fluorescence spectra of a honey bee abdomen
- VII.3.3 X-Ray fluorescence spectra of a honey bee head
  - 1°) Localization of the Mo in a large area of the head
  - 2°) Focus on the neurolemma
  - 3°) Focus on hypopharyngeal glands
- VII.3.4 X-Ray fluorescence spectra of cuticle / thorax

VII.4 Conclusions of X-Ray fluorescence studies

### Part VII. Synchrotron radiation X-ray fluorescence microscopy

In part VI, we evidenced the presence of Molybdenum in the different tagmata of the bee body thanks to ICP-MS. This element is naturally present in bees (see part I) and a feeding with our Mo-complexes provoked an increase in the level of Mo in all bee tagmata. In this part, we aim to locate more precisely the Mo, in natural bees and in bees fed with **Na-$Mo_2O_4$-EDTA**. X-ray fluorescence spectra were recorded at the NANOSCOPIUM beamline of the Synchrotron SOLEIL (Gif-sur-Yvette, France) on cross-sections of head, thorax and abdomen and on hypopharyngeal glands extracted from the head of worker bees.

#### VII.1 Experimental procedures

##### 1°) Animal preparation

The study was conducted on age-controlled bees. The bees were sampled before their emergence by placing a capped brood comb in an incubator at 35°C and 50% humidity. The day of their emergence, bees were collected and placed into plexiglass cages of Pain-type (see Part III), in which they were fed with a sugar syrup containing **Na-$Mo_2O_4$-EDTA** at 400 mg/L during 14 days, while a control group was constituted in the same manner except that bees only received syrup.

After 14 days of feeding in cages, the bees were collected, narcosed with $CO_2$ and cryogenized to preserve the organic tissues and organs. To do so, individuals were soaked into isopentane (Sigma-Aldrich) cooled to -50°C with dry ice for approximately 1 minute. After being cryopreserved, bees were cut on a dry ice bed in order to separate the three tagmata. Samples were stored at -80°C.

##### 2°) Cross sections

This part of the work was performed using a Leica CM1950 cryostat with diamond-coated blades. Prior to sectioning, the specimens were embedded in tissue freezing medium (Leica) and then rapidly frozen in Carboglass before being mounted on a sample holder adapted to the cryostat.

Croos-sections of head, thorax and abdomen with a thickness in the range 20-50 µm were prepared. The cryostat chamber temperature was set at -20°C while the sample was cooled at -18°C. Figure SVII.1 indicates the positions of the sections we selected in head, thorax and abdomen aiming to focus our measurements on brain, muscles and rectum respectively.

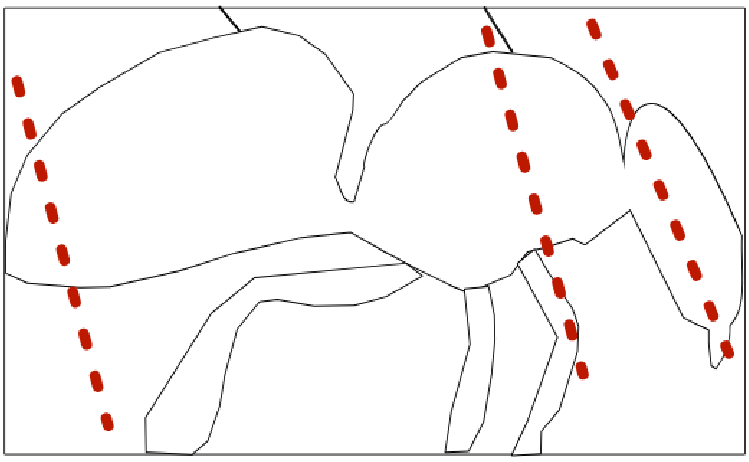

***Figure SVII.1.*** *Positions of the cross-sections of bees used in this study. Draw of the bee adapted from http://marmie.bernard.free.fr/apiculture/RUCHES_morphologie_et_anatomie_de-l_abeille.pdf*

The selected tissue sections were either placed on a silicon nitride, $Si_3N_4$, membrane for X-ray fluorescence imaging (5 mm x 5 mm or 7 mm x 7 mm, 500 nm membrane thickness provided by Silson company, see Figure SVII.2) or on classical glass slides (Superfrost) for histological control of the tissues under the microscope.. Specimens cryofixed on glass slides or silicone membranes were fixed in a vacuum desiccator for 10 min.and conserved at -80°C.

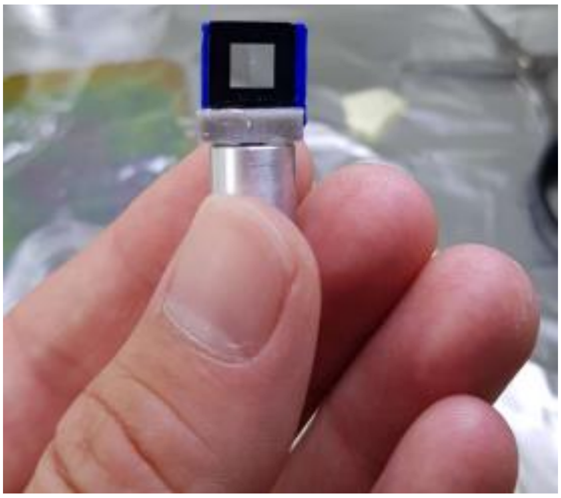

***Figure SVII.2*** *: View of a $Si_3N_4$ membrane used for fluorescence X-ray analysis. The membrane of 5mm x 5 mm, 500 nm thickness is in a frame of silicon 10 mm x 10 mm, 200 µm thick.*

**3°) X-Ray Fluorescence spectra.**

The hard X-ray nano-probe beamline, NANOSCOPIUM of the French national synchrotron facility, SOLEIL, Gif-sur-Yvette, France was used for imaging (see Figure SVII.3). [1]

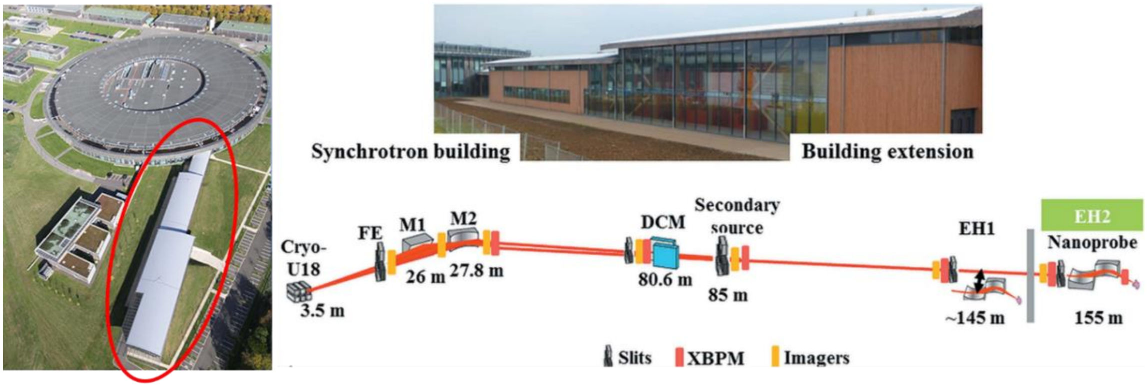


***Figure SVII.3** : View and scheme of the synchrotron building and of the NANOSCOPIUM 155 m-long scanning hard X-ray nanoprobe beamline. See ref [1] for more details.*

A multiscale characterisation of the selected samples has been performed by fast scanning X-ray Fluorescence imaging. The sub-micron scale variation of the Mo distribution has been studied at 20.1 keV excitation energy just above the Mo K-edge. The X-ray beam was focused by a Kirkpatrick-Baez nano-focusing mirror to few hundred of nanometers size at the sample position. For the maps the FLYSCAN continuous scanning mode (https://doi.org/10.1107/S1600577515009364) was used with 300 nm image pixel-size. The full XRF spectra were collected in each pixel by two Si-drift detectors (VITUS H50, KETEK GmbH) in order to increase the detection solid angle. The XRF spectra of the two detectors were added and the sum was used for calculating the elemental maps. The initial large scans in head were obtained at a low resolution of 1 µm at a dwell time of 100 ms to assess the presence of elements in the samples (15 hours of acquisition for the larger scans). Once the Mo element had been located, much smaller scans were performed at a higher resolution scan with 500 nm pixel size and 200 ms dwell time per pixel was performed. Due to the trace quantities of Mo and several other metals in the sample, a duration in the range 2-20 hours was necessary for the scan of each sample. Samples of natural bees (control) and bees fed with **Na-$Mo_2O_4$-EDTA** at 400 mg/L (group "Mo400") were measured. For each type of sample, the spectra were acquired for at least two different bees. In particular, X-Ray fluorescence spectra on neurolemma and on hypopharyngeal glands were measured from two bees form the control groups and two bees from the "Mo400" group. In each case, spectra were acquired in different zone to obtain at least 6-7 spectra of neurolemma and hypopharyngeal glands from both groups.

The XRF sum-spectrum of each scan was fitted by an open source software PyMCA [2] of the European Synchrotron Radiation Facility including background correction. An in-house algorithm developed in MATLAB was used to obtain the elemental distribution maps and the mean XRF spectrum of the cell within a mask region.

#### VII.2 Bee Cross-section results

Bee cross sections were prepared at the IJCLab (Laboratory of the Physics of the two Infinities Irène Joliot-Curie) at Orsay, France. The quality of cross sections was controlled by optical microscopy. Figures SVII.4-A, B, and C, as well as SVII.6, SVII.7, and SVII.8, show sections from the head, thorax, and abdomen, respectively.

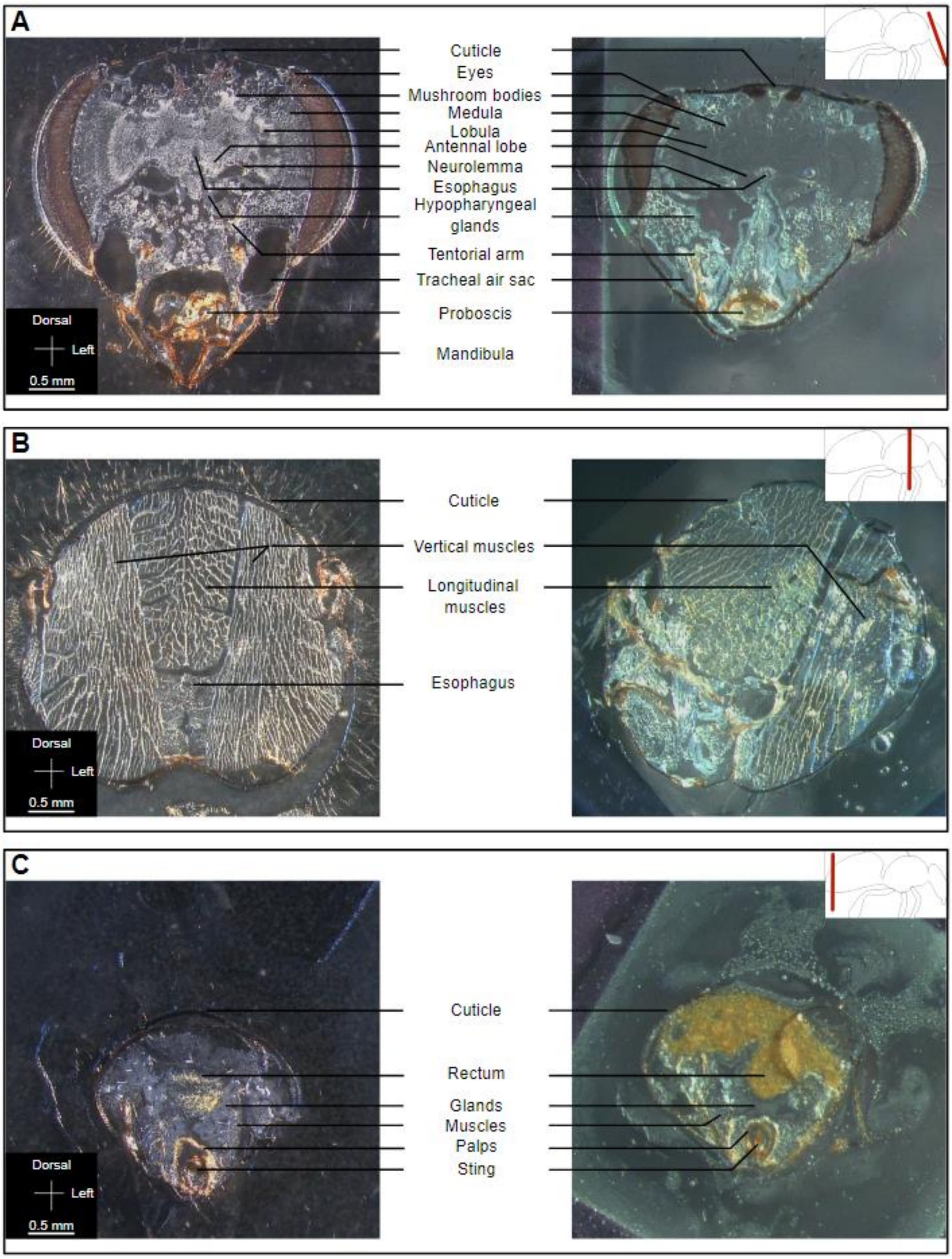


***Figure SVII.4 :*** *Cross-sections of a worker bee. Location of the sample inside the body is shown in the upper right corner of the picture, while orientation and scale are shown in the lower-left corner. A) Head : 40 µm thick sample deposited on a Superfrost glass (left) on a* $Si_3N_4$ *membrane (Right). B) Thorax : 40 µm thick sample deposited on a Superfrost glass (left) or on a* $Si_3N_4$ *membrane (right). C) Abdomen : 50 µm thick sample deposited on a Superfrost glass (left), 40 µm thick sample deposited on a* $Si_3N_4$ *membrane (right).*

Significant effort was dedicated to optimizing the protocol, particularly to ensure the integrity of the cuticle during cutting. After determining the most effective experimental conditions for producing these 20-50 µm thick cross-sections, we found that some structures are better preserved by the cut than others. The muscles and glands inside the head and abdomen were particularly sensitive to temperature variations during handling. On the Superfrost slides, the tissues showed better integrity, whereas on the Si3N4 membranes, shrinkage of certain tissues was sometimes visible, especially after the vacuum phase. This step was therefore limited or excluded for samples prepared for synchrotron measurements.

#### VII.3 Results of synchrotron measurements

##### VII.3.1 X-Ray Fluorescence spectrum of resin on $Si_3N_4$ membrane

As previously seen, the cross-sections were made from bee body parts immobilized in a resin (Tissue Freezing Medium, Leica) and the sections with this resin were deposited on silicon nitride membranes.

The NANOSCOPIUM beamline can be used to detect metals in extremely small quantities. To begin with, we recorded an X-ray fluorescence spectrum on the resin deposited on $Si_3N_4$. The X-ray fluorescence spectrum obtained is shown in Figure SVII.5.

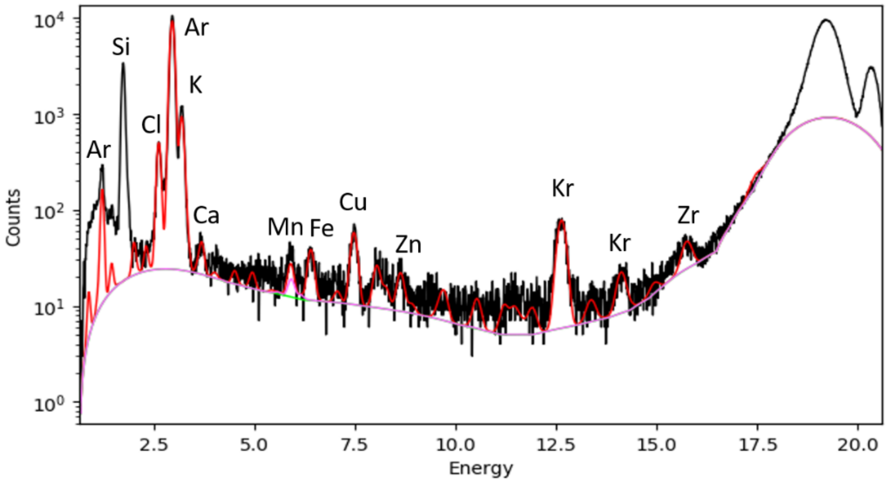


***Figure SVII.5**. X-ray Fluorescence spectrum of a resin sample (Tissue Freezing Medium) deposited on a silicon nitride membrane. The experimental spectrum is traced in black. The simulated spectrum is shown as a red line.*

The X-Ray fluorescence spectrum of the resin of $Si_3N_4$ exhibits a few signals assigned to various elements. Rare gases Kr and Ar are coming from the atmosphere and are detected thank to the high sensitivity of the technique. The presence of Si is due to the membrane. The elements Zr, Zn, Cu, Fe, Mn, Ca, K, Cl are identified as trace in the resin we used but no interfering lines are observed in the energy range expected for Mo (17.37 and 17.48 eV respectively for MoKL1 and MoKL2 transitions).

##### VII.3.2 X-Ray fluorescence spectra of a honey bee abdomen

The bee sections studied were 20 µm thick and the pixel resolution was 1x1 µm² with an integration time of 150 ms/pixel. Under these conditions, we expected to measure quantities of Mo in the nanogram range or less. Our initial studies therefore focused on analysing the rectum of a bee fed with the complex **Na-$Mo_2O_4$-EDTA** at 400 mg/L during 14 days. As we saw in Part IV, in captivity the abdomen becomes loaded with Mo, probably in the faeces. The objective of the analysis of this sample was to verify the feasibility of this analysis by X-ray fluorescence on the Synchrotron. The results of this analysis are presented in Figure SVII.6 in comparison with a control sample registered in the same conditions (Figure SVII.7). The experimental details are given in the legends of the figures.

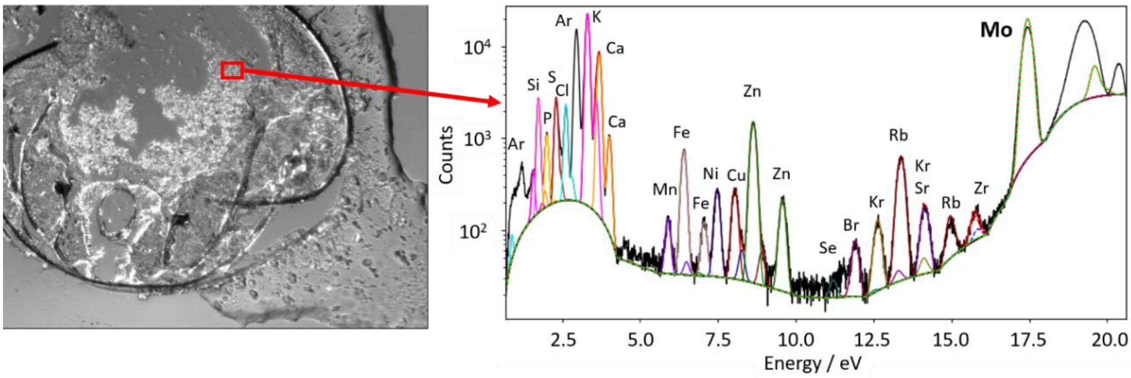


***Figure SVII.6.*** *Picture of a 20 µm thick section from of the abdomen of a bee fed with* **Na-$Mo_2O_4$-EDTA** *(left). X-ray fluorescence spectrum corresponding to the area indicated by the red rectangle in the photograph (right). Size 139 x 144 µm², pixel size 1x1 µm², acquisition time/pixel 150 ms.*

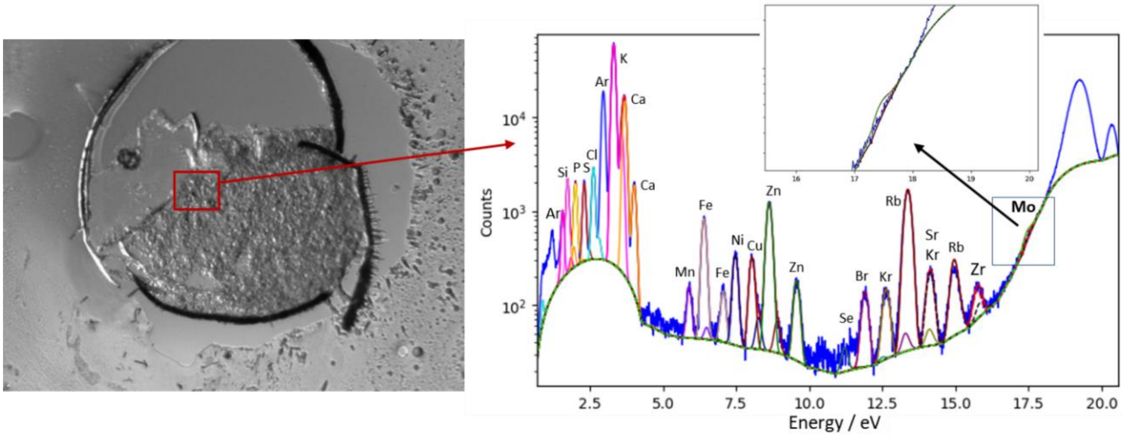


***Figure SVII.7.*** *Picture of a 20 µm thick cross-section of the abdomen of a control bee (left). X-ray fluorescence spectrum corresponding to the area indicated by the red rectangle in the photograph (right). The blue curve corresponds to the experimental data. Size 160 µm x 162 µm, pixel size 1x1 µm², acquisition time/pixel 150 ms.*

As can be seen from these two figures, the X-ray fluorescence spectrum of the control abdomen does not show the presence of Mo in the rectum. On the contrary, the rectum of a bee fed with a sugar solution containing Mo shows a very large and intense peak at the energy expected for Molybdenum, which validates the use of this technique for our study.

#### VII.3.3 X-Ray fluorescence spectra of a honey bee head

##### 1°) Localization of the Mo in a large area of the head

As shown in part IV of the Supporting Information, the level of Mo significantly increases in head by feeding bees with syrup containing the complex **Na-$Mo_2O_4$-EDTA**. In this study, the head was thus our main objective. First, we decided to investigate a large area of the head, including the brain, the neurolemma, the cuticle and the oesophagus as a muscular part. Figure SVII.8A shows a 20 µm thick slice of the head of a bee fed with the complex **Na-$Mo_2O_4$-EDTA** with the zone we studied represented by a red rectangle. Figure SVII.8B shows the total X-ray fluorescence spectrum obtained for this whole region, while Figures SVII.8C-K show the distribution of selected elements based on the X-ray fluorescence of each of element.

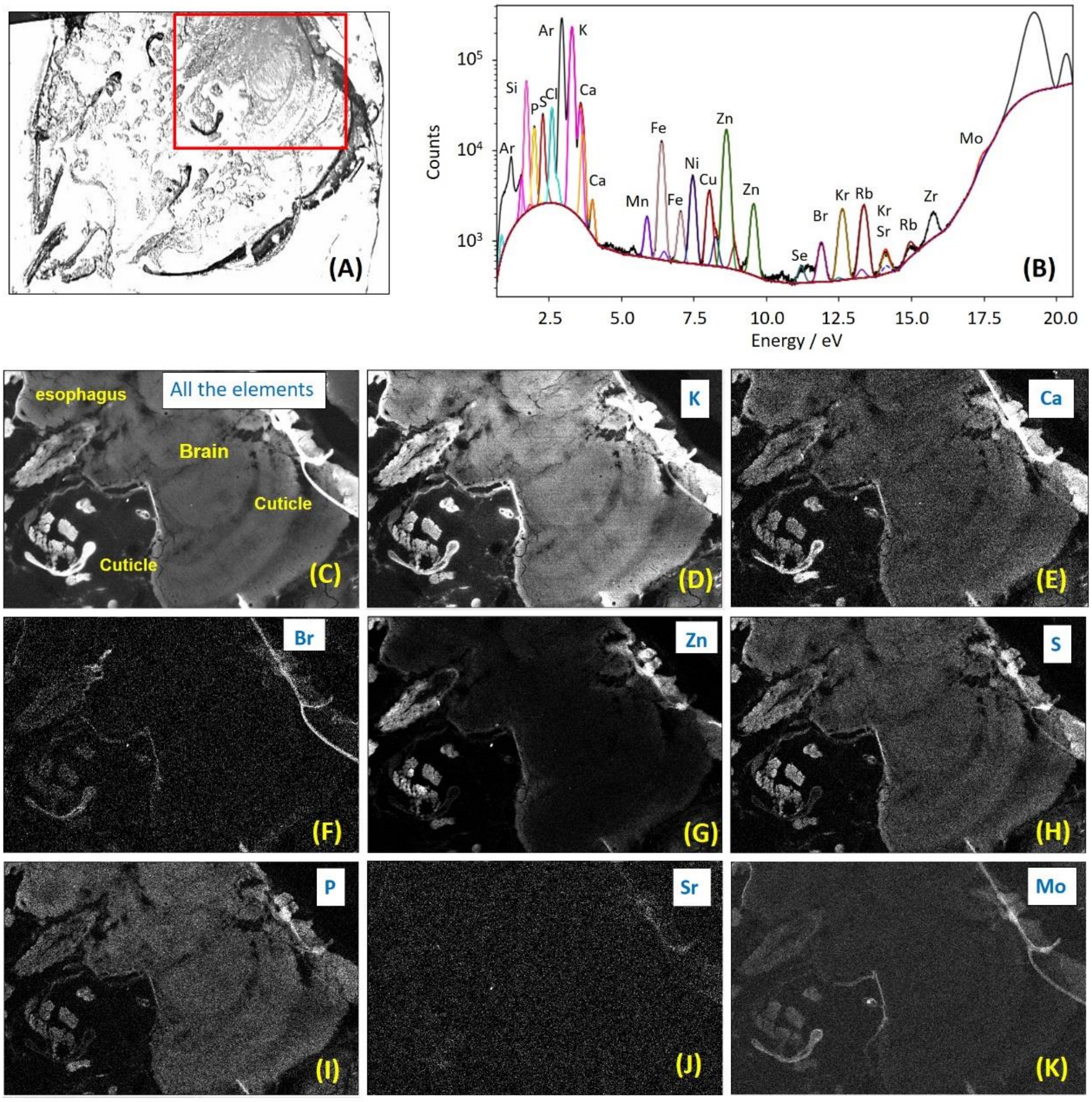


***Figure SVII.8.*** *Picture of a 20 µm thickness cross-section of the head of a worker bee fed with Mo (A). The red rectangle corresponds to the zone we analysed; total X-Ray fluorescence spectrum of the zone analysed with assignments of the peaks (B); Mapping of the elements in the zone analysed thanks to the X-ray fluorescence of all elements (C), K (D), Ca (E), Br (F), Zn (G), S (H), P (I), Sr (J), and Mo (K). Size 2000 µm x 1000 µm, pixel size 2x2 µm², acquisition time/pixel 100 ms.*

If we focus on the X-ray fluorescence of Mo (Figure SVII.8K), the white zones indicate the presence of this element. In comparison with the X-Ray fluorescence of Sr, which is almost negligible at all locations, Figure SVII.8K shows the presence of Mo in the cuticle (external and tentorial arms), in all the brain and especially at the level of the neurolemma around the brain. The presence of Mo in the cuticle is not linked to feeding with our complex, as Mo is also found in the cuticle of control bees as shown on Figure SVII.9 (and *vide infra*). The tentorial arm (pilar) appears indeed clearer on the picture obtained by the X-ray fluorescence of Mo for 20 µm thick slice of a head of a control worker bee (Figure SVII.C), which indicates the presence of Mo in this part, while the other parts of the brain do not exhibit significant fluorescence of Mo.

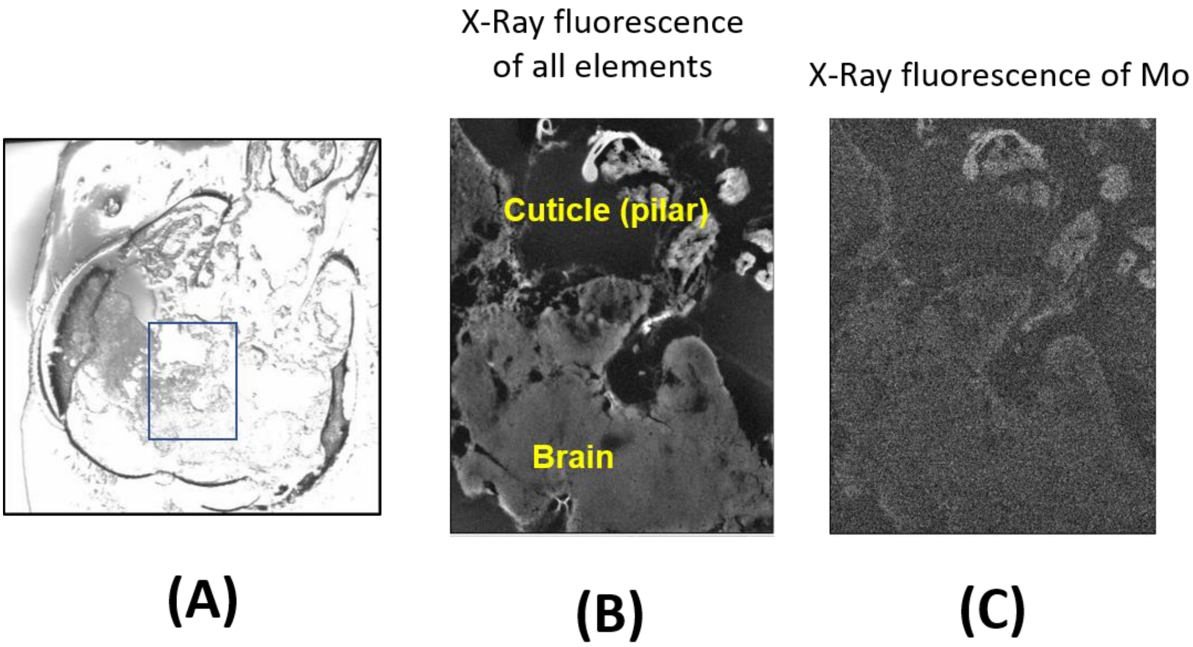


***Figure SVII.9.*** *Microscope picture of a 20 µm thickness cross-section of the head of a worker bee control (A). The blue rectangle corresponds to the zone we analysed; (B) picture of the analysed thanks to the X-ray fluorescence of all elements (B) and Mo only (C). Size 1500 µm x 1000 µm, pixel size 2x2 µm², acquisition time/pixel 100 ms.*

In Figure SVII.10, to evidence that the Mo detected here in neurolemma of a bee fed with Mo is directly linked to feeding with **Na-$Mo_2O_4$-EDTA**, we thus focused our attention on the neurolemma (green rectangle on Figure SVII.10B) to get a more precise analysis of this membrane. Figure SVII.10C presents the fluorescence of Mo in this small zone, clearly showing the neurolemma, while Figure SVII.10D represents the intensity of Mo X-ray fluorescence pixel by pixel in this zone (blue for low level; green, yellow or red for higher levels of Mo). The neurolemma membrane can be clearly distinguished from the Mo levels of fluorescence. The presence of this element is confirmed by the X-Ray fluorescence spectrum corresponding to the sum of pixels delimiting the membrane with a peak characteristic of Mo at *ca* 17.4 eV (Figure SVII.10F), while the X-ray fluorescence of the entire region shows only a shoulder for Mo due to its low average content in the overall sample (Figure SVII.10E).

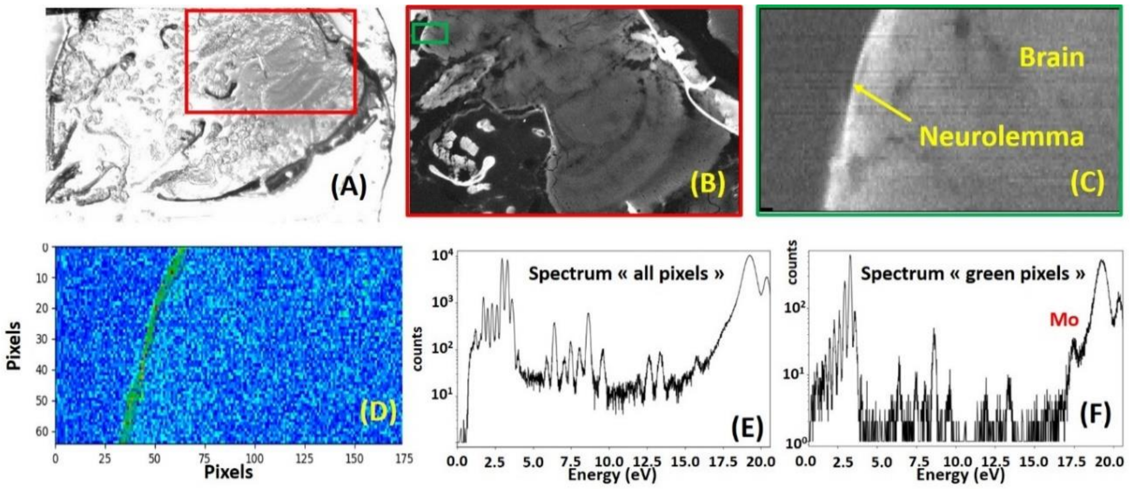


***Figure SVII.10.*** *Picture of a 20 µm thick section from the head of a bee fed with Mo (A). X-ray Fluorescence of all elements in the region corresponding to the red rectangle in A (B); X-ray fluorescence of Mo for a smaller region (green rectangle in B, 181 x 64 µm² ; pixel size 1x1 µm², acquisition time/pixel 100 ms) focusing on the neurolemma (C) ; Intensity of X-ray fluorescence of Mo pixel by pixel (D) ; X-ray Fluorescence spectrum of the entire zone (E) ; X-ray Fluorescence spectrum focused only on the neurolemma membrane (F).*

Figure SVII.11 presents the X-Ray fluorescence obtained on the whole worker head when focusing only on the energy of the Mo element (range 16.9 to 17.9 eV). To emphasize Mo variations in the different parts of the head, we plotted the profile shown in Figure SVII.11 which represents Mo contribution in the hatched zone along X for a bee fed with Mo.

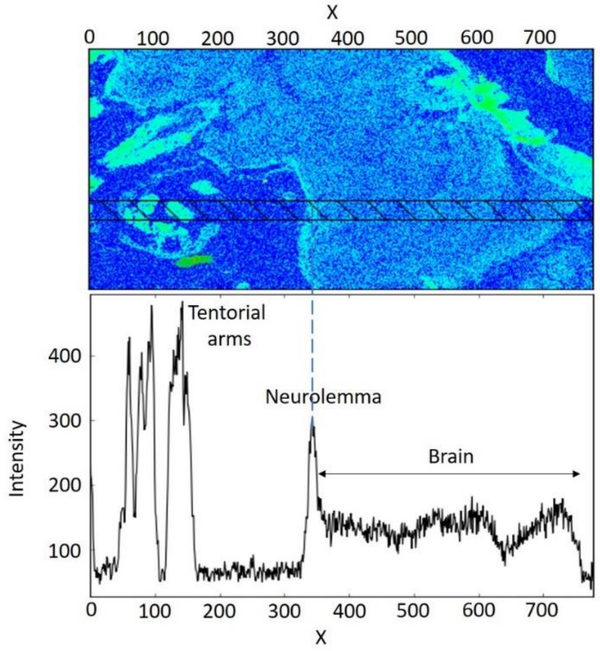


***Figure SVII.11.*** *Variation of Mo contribution (from background between energy* 16.9 to 17.9 eV) *along X in a 20 µm thickness slice of the head of a worker bee fed with Mo evidencing the loading in Mo of tentorial arms (cuticle), neurolemma and brain.*

When Mo is absent, the intensity is close to zero (intensity around 50-60 counts for the baseline). On the other hand, the intensity of the Mo fluorescence radiation is high at the level of the tentorial arms in the head (made of cuticle), then presents a sharp peak at the level of the neurolemma. In the brain, the level of Mo does not fall back to the level of the baseline but stay higher at an average value of 130-140 counts along X, while at similar experimental conditions the intensity of Mo signal in the brain of a control bee is found very close to that of the baseline (around 70 counts) as shown in Figure SVII.12. It suggests a diffuse and continuous presence of this element Mo in this structure after feeding with the complex **Na-$Mo_2O_4$-EDTA**.

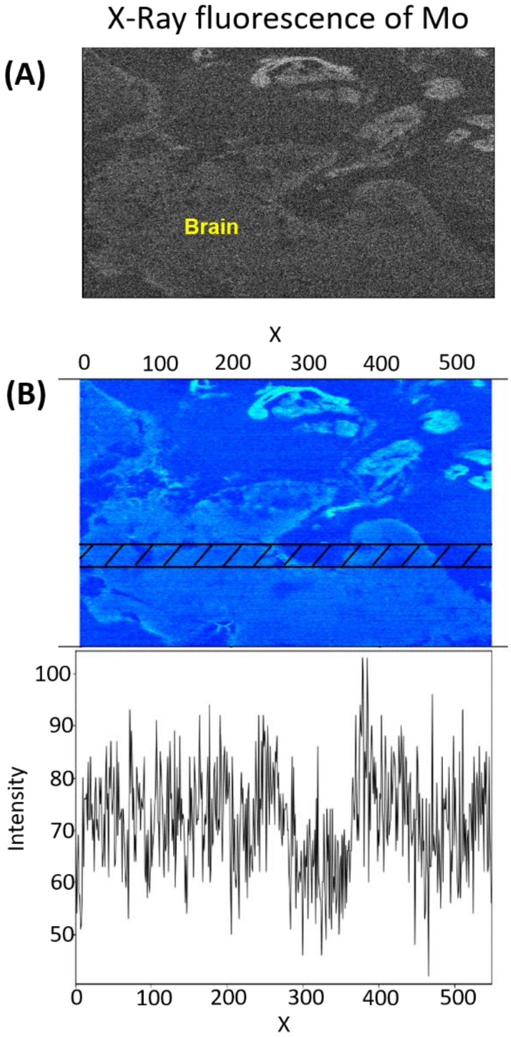


***Figure SVII.12.*** *Variation of Mo contribution (from background between energy* 16.9 to 17.9 eV) *along X in a 20 µm thickness slice of the head of a worker bee control showing a low intensity level of Mo within the brain along X in the hatched zones.*

**2°) Focus on the neurolemma**

To detail the presence of Mo in the neurolemma and to confirm that its presence is linked to or enhanced by feeding with the **Na-$Mo_2O_4$-EDTA** complex, we carried out two series of spectra focusing on the neurolemma of several individuals from the control group or from the group fed with the complex. To improve the quality of the measure, the thickness of the sections was increased to 40 µm (x 2), the size of the areas analysed was reduced, and pixel size was reduced to 0.5 µm x 0.5 µm (/4). Acquisition time was increased to 200-300 ms per pixel (x2 or x3). Figures SVII.13 and SVII.14 present the spectra of neurolemma recorded for control bees and from bees fed with the complex **Na-$Mo_2O_4$-EDTA**.

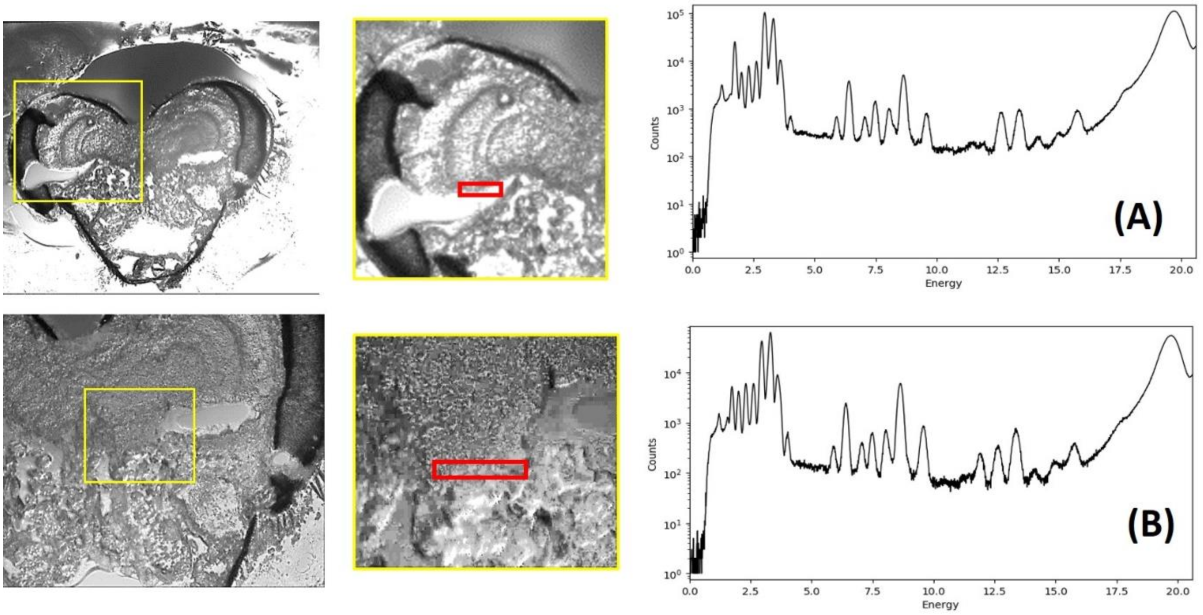


***Figure SVII.13.*** *Pictures of 40 µm thick cross-sections of the head of a worker bee from the control group. Fluorescence spectra of all elements in the red rectangle zones (A and B). Experimental conditions: Pixels 0.5x0.5 $\mu m^2$, size 100x60 $\mu m^2$, acquisition time/pixel 300 ms (spectrum A); Pixels 0.5x0.5 $\mu m^2$, size 218x70 $\mu m^2$, acquisition time/pixel 300 ms (spectrum B).*

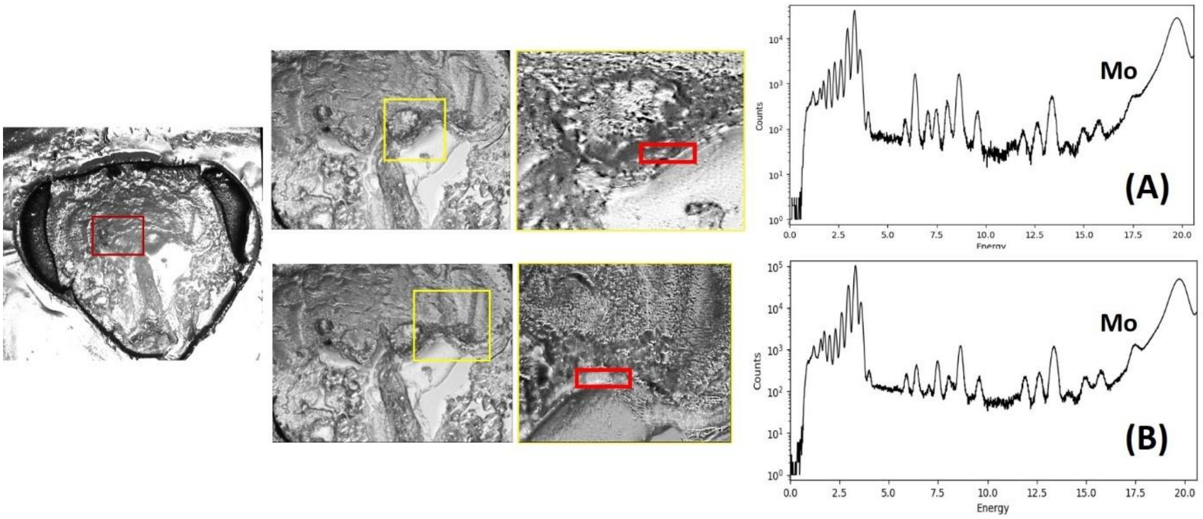


***Figure SVII.14.*** *Pictures of 40 µm thick slices of the head of a worker bee fed with Mo. Fluorescence spectra of all elements in the red rectangle zones (A and B). Experimental conditions: Pixels 0.5x0.5 $\mu m^2$, size 134x25 $\mu m^2$, acquisition time/pixel 200 ms (spectrum A); Pixels 0.5x0.5 $\mu m^2$, size 136x33 $\mu m^2$, acquisition time/pixel 300 ms (spectrum B).*

In a control bee (Figure SVII.13), we note the presence of a shoulder at the energy expected for the element Mo. Thus, Mo is naturally present, although at a low level, in the neurolemma of control bees.

In a bee fed with the complex **Na-$Mo_2O_4$-EDTA** (Figure SVII.14), the fluorescence spectra clearly show the characteristic peak of element Mo in the neurolemma. The observation of a more intense peak in fed bees compared to control bees demonstrates that Mo level increases in the neurolemma with feeding with the **Na-$Mo_2O_4$-EDTA** complex.

From the variation of intensities of this peak from control to fed bees, we estimate that the Mo level is increased by a factor around 10 on average by feeding bees with **Na-$Mo_2O_4$-EDTA** at 400 mg/L.

**3°) Focus on hypopharyngeal glands**

The hypopharyngeal glands are found in the head of honey bee workers (Figure SVII.15). These glands are crucial for bees' nutrition and health, notably through the production of vitellogenin, proteins, lipids, vitamins, enzymes. They are therefore of interest for understanding the many effects of feeding with Mo complexes observed in our study. Glands were taken from control bees or bees fed with the complex **Na-$Mo_2O_4$-EDTA** at 400 mg/L during 14 days. Two series of glands for each bee group were deposited on silicon nitride membranes and analysed by X-Ray fluorescence. Figure SVII.16 presents the results obtained for glands of 2 bees from the control group (6 acini measured in total), while Figure VII.17 shows the results obtained for glands of 2 bees from the "Mo 400" group (7 acini measured in total).

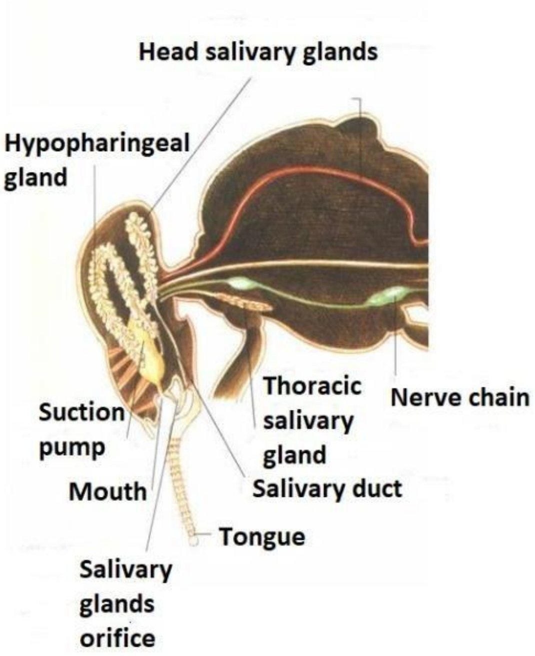


***Figure SVII.15.*** *Anatomy of the head of a worker honey bee. Picture reproduced with permission from* https://abejas.org/en/bees-internal-anatomy

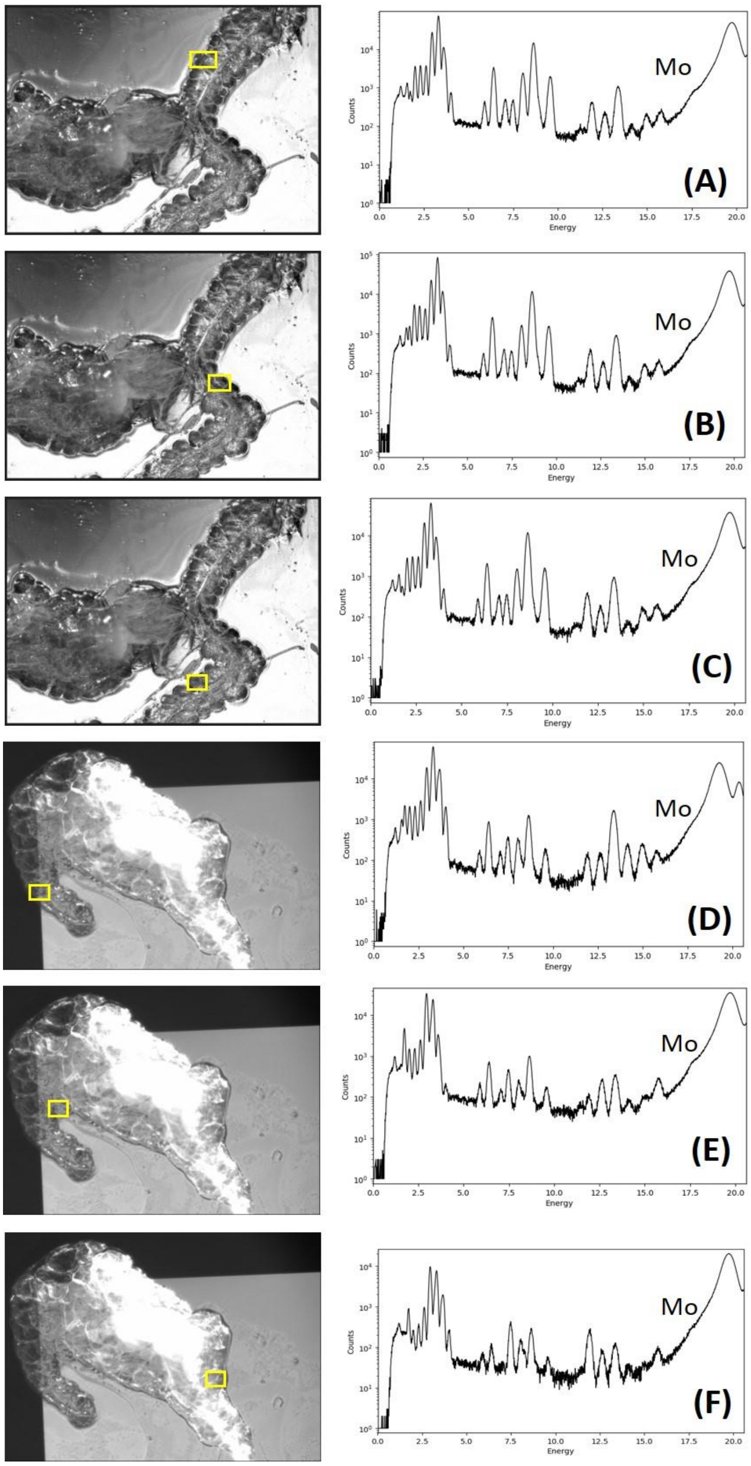


***Figure SVII.16.*** *Pictures of hypopharyngeal glands from workers of the control group. The yellow rectangles indicate the part of the gland on which the fluorescence spectrum was measured. Experimental conditions : Pixels 0.5x0.5 µm², size 108x49 µm², acquisition time/pixel 200 ms (spectrum A); Pixels 0.5x0.5 µm², size 72x59 µm², acquisition time/pixel 200 ms (spectrum B); Pixels 0.5x0.5 µm², size 76x53 µm², acquisition time/pixel 200 ms (spectrum C); Pixels 0.5x0.5 µm², 60x90 µm², acquisition time/pixel 200 ms (spectrum D); Pixels 0.5x0.5 µm², size 70x72 µm², acquisition time/pixel 200 ms (spectrum E); Pixels 0.5x0.5 µm², size 70x70 µm², acquisition time/pixel 200 ms (spectrum F);*

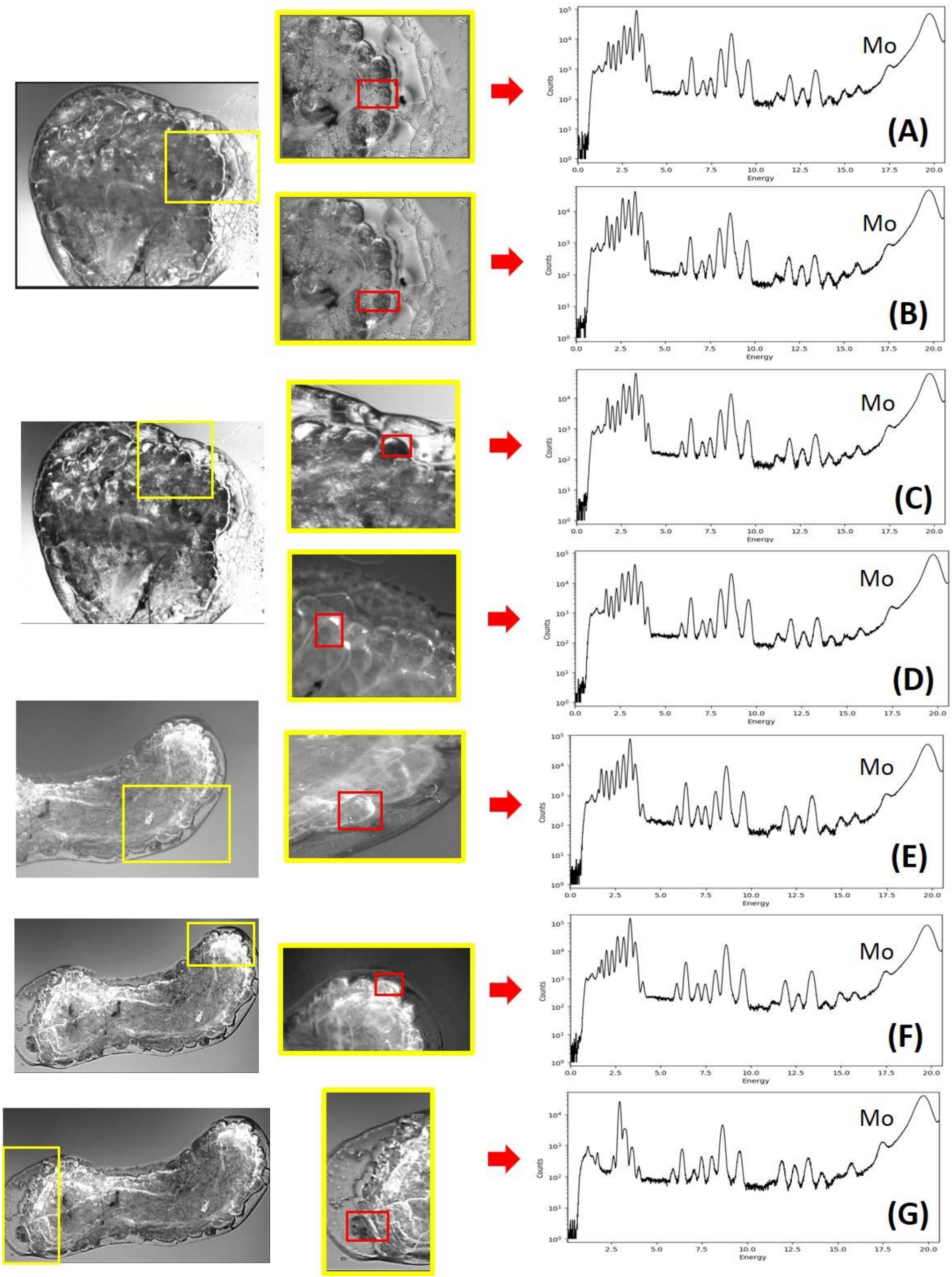


***Figure SVII.17.*** *Pictures of hypopharyngeal glands from workers of the "Mo400" group. The yellow rectangles indicate the parts of the gland on which the X-ray fluorescence spectrum was measured. Experimental conditions: Pixels 0.5x0.5 µm², size 104x51 µm², acquisition time/pixel 200 ms (spectrum A); Pixels 0.5x0.5 µm², size 85x52 µm², acquisition time/pixel 200 ms (spectrum B); Pixels 0.5x0.5 µm², size 70x70 µm², acquisition time/pixel 200 ms (spectrum C); Pixels 0.5x0.5 µm², size 70x70 µm², acquisition time/pixel 200 ms (spectrum D); Pixels 0.5x0.5 µm², size 90x50 µm², acquisition time/pixel 200 ms (spectrum E); Pixels 0.5x0.5 µm², size 100x60 µm², acquisition time/pixel 200 ms (spectrum F); Pixels 0.5x0.5 µm², size 70x80 µm², acquisition time/pixel 200 ms (spectrum G);*

In control bees (Figure SVII.16), the fluorescence spectra were similar to those observed for the neurolemma, with a shoulder at 17.4 keV corresponding to a low, natural, presence of molybdenum in these glands. Feeding with a sugar syrup containing the **Na-$Mo_2O_4$-EDTA** complex induced a remarkable fluorescence peak as shown in Figure SVII.17. Here again, the results obtained for 7 *acini* of hypopharyngeal glands from 2 different individuals are similar. The intensity of the peaks attributed to Mo in relation to the baseline allows to estimate an 14x increase for Mo content in the hypopharyngeal glands between the control batch and the batch of bees fed with the Mo complex.

These results evidence that **the hypopharyngeal glands are a prime target for the Mo complex, which reinforces the Mo already present naturally**.

##### VII.3.4 X-Ray fluorescence spectra of cuticle / thorax

As demonstrated in section VII.3.2-1°), the cuticle contains molybdenum (see Figure SVII.8 for example). To confirm this result, we imaged the thoraces of different samples from the control and "Mo400" groups. Figure SVII.18A-B shows a 20 µm thorax section from a control bee. This section shows a poor-quality thorax section. Nevertheless, it contains fragments of cuticle with fragments of muscles which proved useful for our study. Figure VII.18C shows the fluorescence of all the elements present in the section and highlights the pieces of cuticle, here in white. A fluorescence spectrum corresponding to the entire zone is given in Figure SVII.18D. Figures SVII.18E-R show the images associated with the fluorescence of the element Mo (SVII.18D) as well as the elements Ca, P, Mn, Fe, Cl, K, S, Zn, Ni, Sr, Cu, Br and Rb.

This experiment shows that the cuticle naturally contains halides such as Cl and Br but also alkali metals such as K and Rb, alkaline earth metals such as Ca, non-metals such as P, S and transition metals such as Cu and Mo. The metals Mn, Fe, Ni or Zn are almost not detected. Interestingly **Mo is clearly naturally present in the composition of the cuticle**.

Similar experiments were performed on a fragment of thorax containing a piece of cuticle from bees fed with the complex **Na-$Mo_2O_4$-EDTA** (Figure SVII.18). We selected a zone containing both muscles and cuticle (Figure SVII.18B). The X-ray fluorescence spectrum shown in Figure SVII.18C displays only a shoulder for the energy expected for Mo, which indicates only traces of molybdenum. An intensity profile along the horizontal direction of the Mo distribution covering cuticle and muscles (Figures VII.18C) indicates a peak when passing through the cuticle (Figure VII.18D) but the intensity remains low.

This observation suggests that:

- in this part, Mo is mainly contained within the cuticle, and the Mo level in muscles appears very low,
- feeding bees with the **Na-$Mo_2O_4$-EDTA** complex after their emergence probably does not enhance the quantity of Mo in the cuticle, as it is formed before emergence.

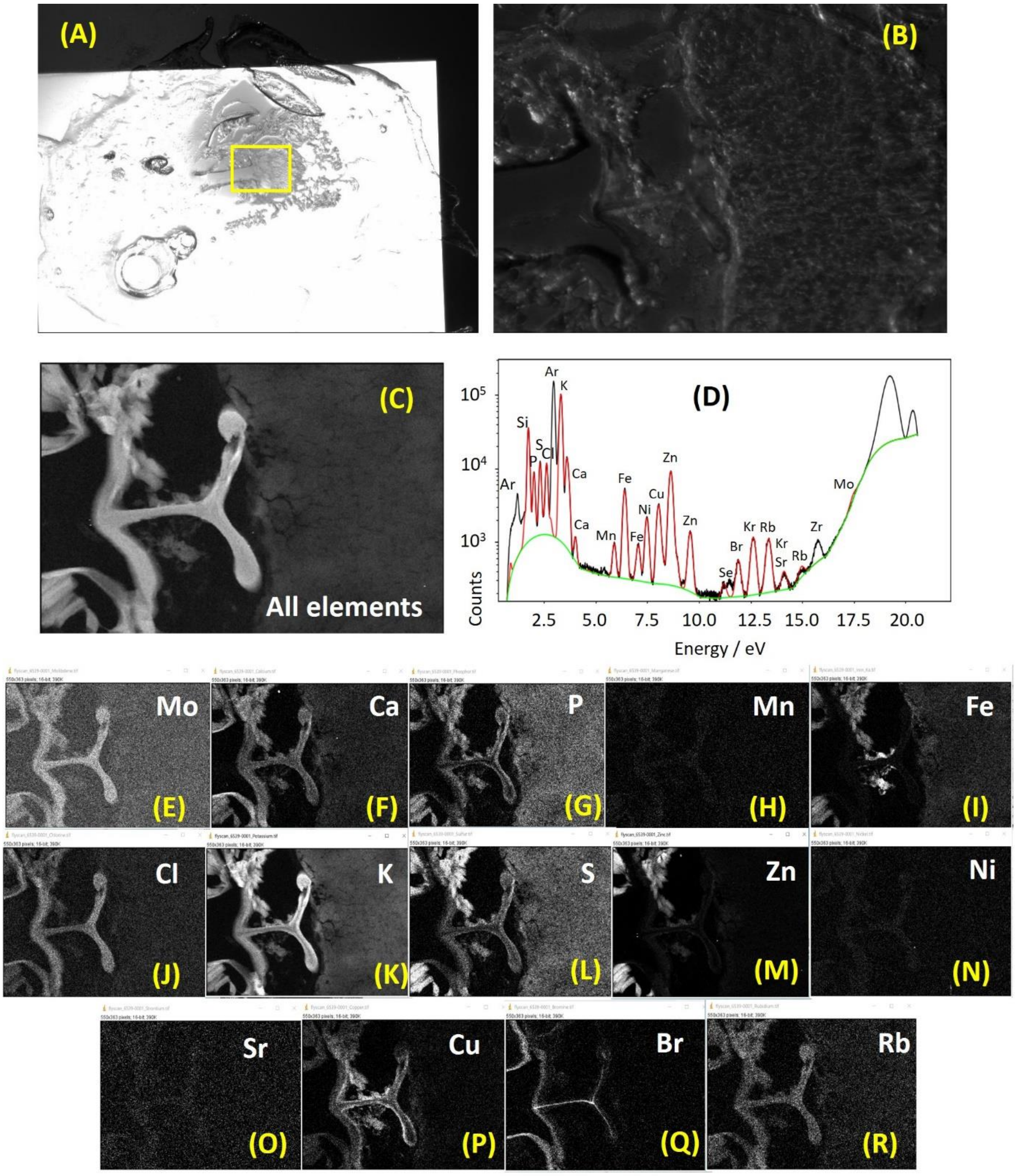


***Figure SVII.18.*** *Picture of a 20 µm thick cross-section from the thorax of a control worker bee (A and B); Picture obtained with the fluorescence of all elements (C); Fluorescence spectrum of the entire zone (D); pictures corresponding to the fluorescence of only the element Mo (E), Ca (F), P (G), Mn (H), Fe (I), Cl (J), K (K), S (L), Zn (M), Ni (N), Sr (O), Cu (P), Br (Q), and Rb (R). Experimental conditions: Size 550 µm x 353 µm, pixel size 1x1 µm², acquisition time/pixel 100 ms.*

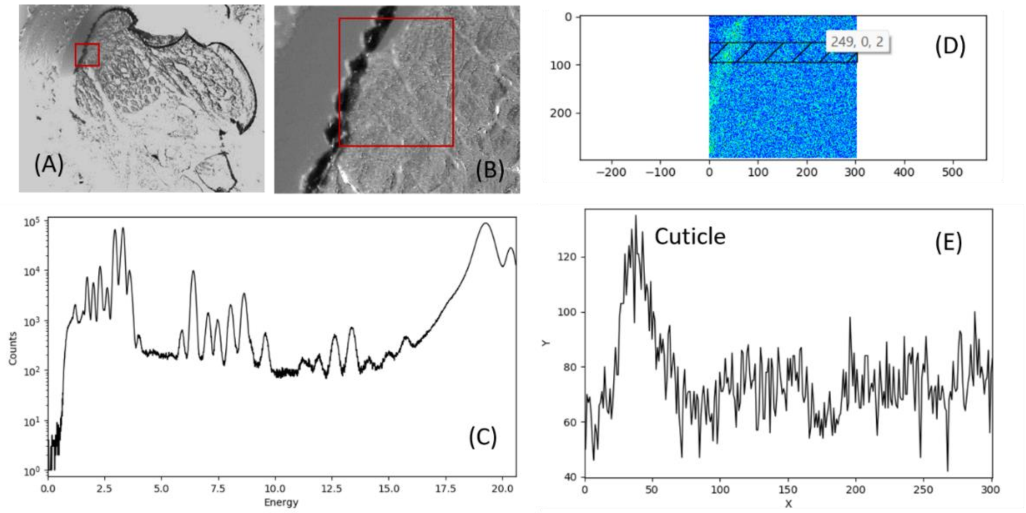


***Figure SVII.19.*** *Picture of a 20 µm thick section from the thorax of a worker bee fed with* **Na-$Mo_2O_4$-EDTA** *(A and B); fluorescence spectrum of the selected zone (C). Intensity of fluorescence of Mo pixel by pixel for the selected zone (D) ; excitation spectrum focused on the energy of Mo along the x axis of the hachured zone in D (E). Experimental conditions: Size 296 µm x 293 µm, pixel size 1x1 µm², acquisition time/pixel 150 ms.*

In addition, to these experiments, Figure SVII.18 shows that the muscles are rich in P, S, K, and contain other metals such as Ca, Rb, Fe. The presence of other metals such as Mn, Ni, Zn or Cu is not clear on the X-ray fluorescence picture but these elements are clearly identified on the fluorescence spectrum given in Figure SVII.18D. Concerning Mo, this fluorescence spectrum displays only a shoulder at the energy expected for Mo. Its presence in muscles is almost negligible. To confirm this point, experiments were performed on 40 µm thick fragments of thorax obtained from bees fed with the **Na-$Mo_2O_4$-EDTA** complex in optimized conditions (small region of analysis, small pixels, high acquisition time, Figure SVII.20).

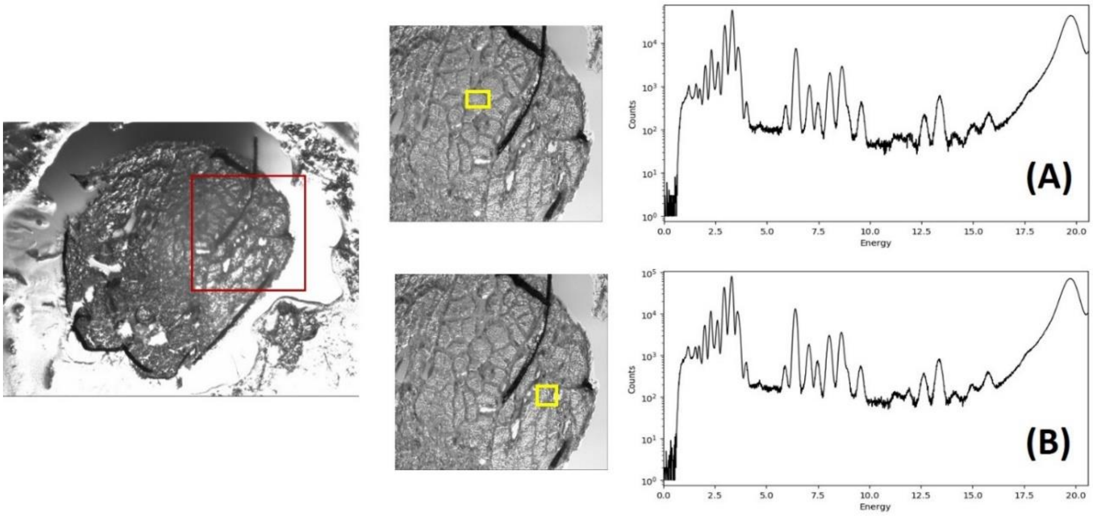


***Figure SVII.20.*** *Picture of a 40 µm thick cross-section of the thorax from a worker bee fed with* **Na-$Mo_2O_4$-EDTA.** *The X-ray fluorescence spectra were recorded for the two zones shown in yellow (spectra A and B). Experimental conditions: Size 116 µm x 43 µm, pixel size 0.5x0.5 µm², acquisition time/pixel 200 ms (A); Size 90 µm x 64 µm, pixel size 0.5x0.5 µm², acquisition time/pixel 300 ms (B).*

#### VII.4 Conclusions of X-Ray fluorescence studies

The X-ray fluorescence studies performed at the NANOSCOPIUM beamline of Synchrotron SOLEIL allowed drawing the following conclusions about Mo in bees:

- Molybdenum is naturally present at low levels in bees in the cuticle, neurolemma, and hypopharyngeal glands. It can also be detected in other parts of the body (muscles, brain) but as trace levels only.
- Feeding bees with the **Na-$Mo_2O_4$-EDTA** complex at a 400 mg/L concentration during 14 days unambiguously provoked a significant increase in Mo levels in the brain, neurolemma and hypopharyngeal glands.
