## Supplementary material for "Food supplementation with molybdenum complexes improves honey bee health": XPS stuides of bee faeces and Anti-Oxydant properties

### Supporting Information

### Part VIII. Antioxidant properties of Na-$Mo_2O_4$-EDTA and Li-$Mo_2O_4$-EDTA complexes

VIII-1 : XPS studies on bees' faeces
VIII-1-1 Experimental section
VIII-1-2 XPS analysis
VIII-1-3 Conclusion
VIII-2 : Antioxidant properties measured in honey bees and products of the hive.
VIII.2.1 Experimental protocols.
1°) Apiary and conditions of feeding.
2°) Sampling
3°) Anti-oxidant activity (AOA) determinations
VIII.2.2 Results.
VIII.2.3 Discussion / conclusions

### Part VIII. Antioxidant properties of Na- $Mo_2O_4$-EDTA and Li- $Mo_2O_4$-EDTA complexes

We have seen that complexes **Na-$Mo_2O_4$-EDTA** and **Li-$Mo_2O_4$-EDTA** have little or no toxicity (Part III), that they are assimilated by the bees, particularly in the head (Parts V, VI, and VII), and that this assimilation leads to beneficial effects measured at the colony level (Part IV). These complexes probably have a range of different effects on bees' physiology. One of these could be as antioxidant species, explaining the beneficial effects observed. One possible role is that of antioxidant species.

In fact, the Mo atoms in the complexes are in the +V oxidation state, leaving the possibility of their oxidation into Mo(+VI), thereby acting as antioxidant. In a recent study, we demonstrated that these two complexes are not directly antioxidant, but that the molybdenum core of these complexes associated with cysteine or histidine leads to compounds with remarkable antioxidant properties [1].

In this section, we investigated this property in two different ways in bees following feeding with **Na-$Mo_2O_4$-EDTA** and **Li-$Mo_2O_4$-EDTA** :

- Using XPS on the faeces of bees fed in the laboratory with **Na-$Mo_2O_4$-EDTA**

- By measuring the antioxidant properties in bees and products of bees in colonies fed with **Na-$Mo_2O_4$-EDTA** and **Li-$Mo_2O_4$-EDTA** as well as sodium molybdate as reference (Mo(+VI))

The results obtained with these two approaches are presented in sections VIII.1 and VIII.2 respectively.

### VIII-1: XPS studies on faeces of bees

#### VIII-1-1 Experimental section

The X-Ray Photoelectron Spectroscopy (XPS) technique is used to characterize elements on surfaces. In particular, the energies detected are characteristic of the elements present and of their degree of oxidation. This technique is therefore ideally suited for studying the fate of the complex **Na-$Mo_2O_4$-EDTA** in bees.

For this purpose, bees used in mortality tests (part III) were used. As mentioned in part IV, the bees do not defecate when they are in captivity in a cage. However, when they die, they sometimes release large quantities of faeces, which were collected on a piece of paper for XPS analysis. For this study we focused our attention on bees fed with the solution of **Na-$Mo_2O_4$-EDTA** at 400 mg/L to increase our chances of detection using XPS.

For surface analysis purposes, **Na-$Mo_2O_4$-EDTA** powder and bee faeces were deposited on carbon tape prior to measurements (Figure SVIII.1). X-ray Photoelectron Spectroscopy (XPS) acquisitions were carried out using a Thermofisher Scientific Nexsa spectrometer with a monochromated Al-Kα X-ray source (hν = 1486.6 eV).

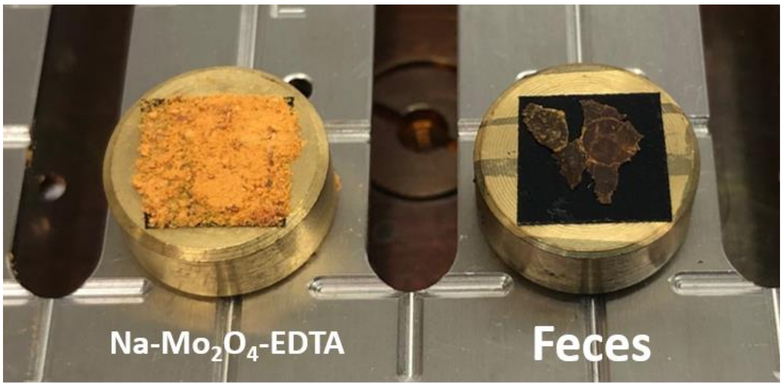


***Figure SVIII.1*** *Samples of* ***Na-$Mo_2O_4$-EDTA*** *powder and bee faeces deposited on carbon tape.*

Both survey spectrum and high energy resolution spectral windows of interest were recorded with a 400 µm spot size. The photoelectron detection was performed perpendicular to the surface using a constant analyzer energy (CAE) mode. Survey spectra were recorded with a 200 eV pass energy and a 1 eV energy step, whereas core level spectra were acquired with a 20 eV CAE and 0.1 eV energy step. The use of low-energy electron and ion flood gun was necessary to perform the analysis. Nevertheless, spectra, especially on the most insulating samples (bee faeces), were then charge corrected by shifting all peaks to the adventitious C1s spectral component (C-C, C-H) binding energy set to 284.8 eV. Quantification was performed using the Thermofisher Scientific Avantage© software. Chemical compositions were obtained from the peak areas after a Shirley type background subtraction and considering "AlThermo1" sensitivity factor library. For the Mo3d peak fitting, a 70% Gaussian/30% Lorentzian peak shape was used. Mo(3d) spin-orbit pair intervals were set at 3.15 eV (Mo(3d3/2) - Mo(3d5/2)), while an area ratio of 1.50 (Mo3d5/2/Mo3d3/2) was used.

#### VIII-1-2 XPS analysis

Among the methods used to characterize **Na-$Mo_2O_4$-EDTA** compounds, X-ray photoelectron spectroscopy (XPS) appears particularly relevant as it gives access to both the elemental composition and the chemical environments of the elements detected at the sample surface. Here, XPS analyses were performed to characterize the synthesized compounds and highlight the presence and the nature of molybdenum in bee faeces.

The XPS survey spectrum analysis of **Na-$Mo_2O_4$-EDTA** powder (Figure SVIII.2) reveals, as expected, the major presence of Mo, C, O and Na by the detection of Mo 3d, C 1s, O 1s and Na 1s core level peaks at 232, 285, 532, and 1071 eV. Traces of Cl 2p are also evidenced at 198 eV, in agreement with traces of NaCl in the complex (<1%).

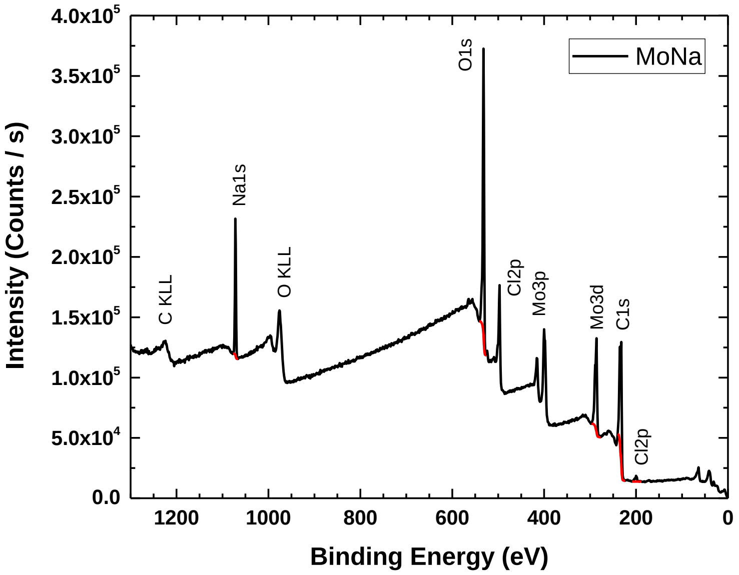


***Figure SVIII.2:*** *XPS survey spectrum of* ***Na-$Mo_2O_4$-EDTA*** *complex (powder)*

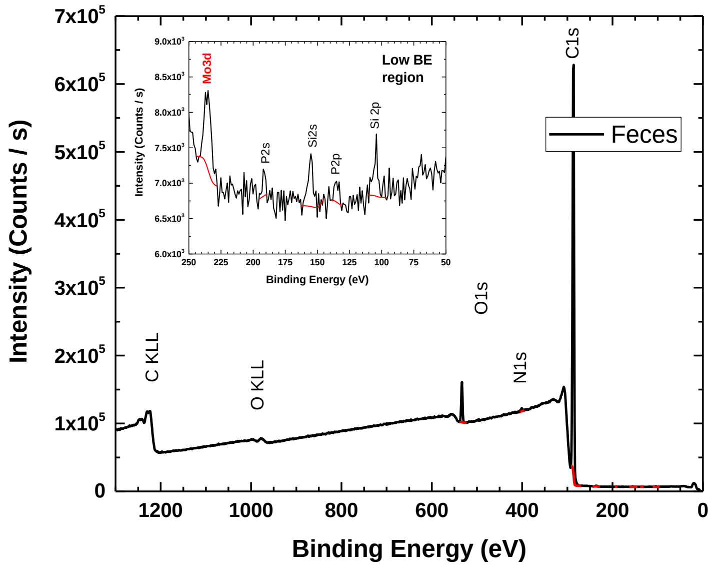


***Figure SVIII.3:*** *XPS survey of bee spectrum faeces after feeding with* ***Na-$Mo_2O_4$-EDTA*** *complex*

The XPS survey of bee faeces (Figure SVIII.3) is dominated by the chemical elements C and O with their main spectral signatures C 1s and O 1s, at the respective binding energies (BE) of 285 and 532 eV. Traces of Si 2p, P 2p, Mo3d and N1s are visible at 102, 133, 233 and 399 eV. Table SVIII.1 summarizes the XPS chemical composition in the two samples obtained from the survey spectra analysis.

*Table SVIII.1: XPS chemical composition of both* ***Na-$Mo_2O_4$-EDTA*** *compound and bee faeces estimated from respective survey spectra analysis*

| **Core level** | **Atomic %** | |
|---|---|---|
| | ***Na-$Mo_2O_4$-EDTA*** | Bee faeces |
| **Si2p** | - | 0.14 |
| **P2p** | - | 0.01 |
| **Cl2p** | 1.1 | - |
| **Mo3d** | 6.1 | 0.03 |
| **C1s** | 46.4 | 95.93 |
| **N1s** | - | 0.56 |
| **O1s** | 39.1 | 3.34 |
| **Na1s** | 7.4 | - |

The Mo 3d spectral region of **Na-$Mo_2O_4$-EDTA** and the bee faeces, as well as their associated reconstruction, are presented in Figure SVIII.4.

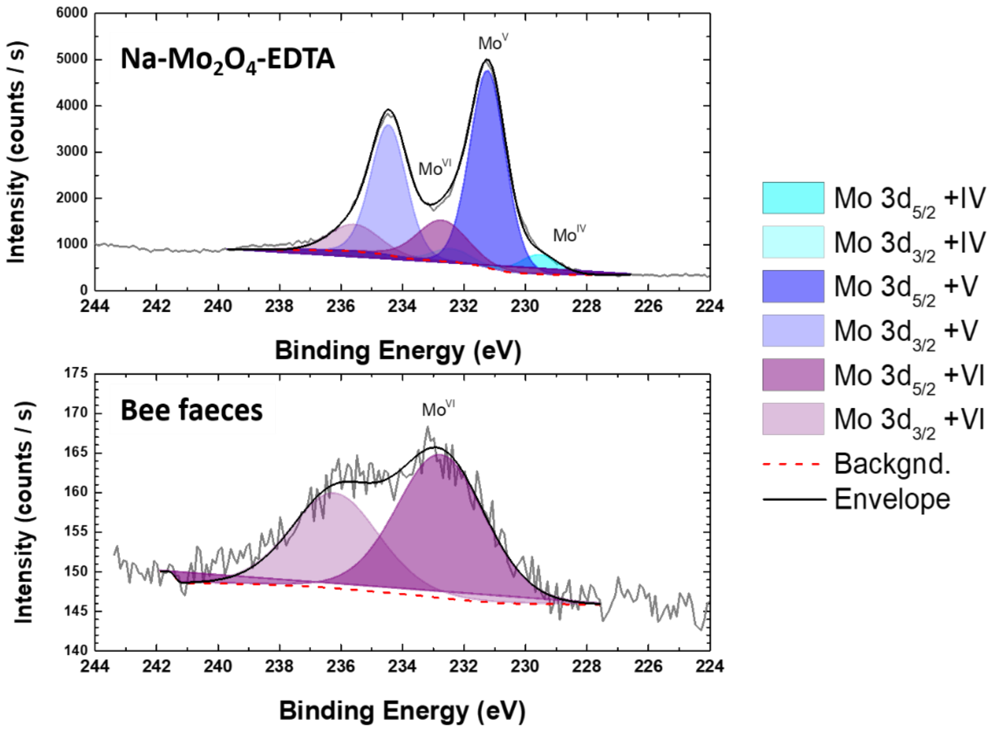


***Figure SVIII.4:*** *XPS high-resolution spectra (Mo3d region) of* ***Na-$Mo_2O_4$-EDTA*** *and the bee faeces. The different contributions and the background used for the reconstruction are plotted as well as the final envelope*

The **Na-$Mo_2O_4$-EDTA** spectrum can be resolved in three doublets, the most intense of which is associated, as expected, to $Mo^{+V}$ (Mo $3d_{5/2}$ = 231.2 eV, FWHM = 1.4 eV). The two other doublets, much less intense, correspond to molybdenum in additional oxidation states. The broader contribution at higher BE (Mo $3d_{5/2}$ = 232.7, Full width at half maximum FWHM = 2.0 eV) can be attributed to $Mo^{+VI}$ phase [2], whereas the sharp component at lower BE (Mo $3d_{5/2}$ = 229.5 eV, FWHM = 1.5 eV) is associated to $Mo^{+IV}$ species [3]. The presence of $Mo^{+VI}$ in the XPS spectrum is typical of crystal surface oxidation since the **Na-$Mo_2O_4$-EDTA** powder was not stored under inert atmosphere prior measurements. The detection of a $Mo^{+IV}$ contribution, meanwhile, can be explained by a partial reduction of the molybdenum under X-ray exposure [4].

For the bee faeces, the Mo 3d core level spectrum was acquired for 4 hours because Mo is a trace element on this surface. In this case, only one broad doublet (Mo $3d_{5/2}$ at BE 232.7 eV, FWHM = 3.5 eV) was detected. The energy position is consistent with a $Mo^{+VI}$ phase [3]. The quite large FWHM observed can be explained by a charging effect not compensated by the flood gun. Surprisingly, no reduction effect under X-ray beam is observed despite the long X-ray exposure. This suggests that even if the oxidation state is the same ($Mo^{+VI}$), the chemical environment of molybdenum in bee faeces is different from that previously observed on the surface of the **Na-$Mo_2O_4$-EDTA** compound.

#### VIII-1-3 Conclusion

The XPS study of bees' faeces sampled after feeding with a 400 mg/L solution of **Na-$Mo_2O_4$-EDTA** unambiguously evidences that the complex **Na-$Mo_2O_4$-EDTA** has been modified inside the bees. From initial Mo(+V) oxidation state in the complex, the XPS study reveals the presence of Mo(+VI) atoms only in the faeces. It suggests that the complex **Na-$Mo_2O_4$-EDTA** has been oxidized, probably into molybdate anion $MoO_4^{2-}$. It constitutes a first element suggesting that the **Na-$Mo_2O_4$-EDTA** complex may act as an antioxidant.

### VIII-2: Antioxidant properties measured in honey bees and products of the beehives

The assessment of Antioxidant Activity (AOA) functions as a pivotal gauge for evaluating the operational state of the antioxidant system in both bees and their larvae. This antioxidative mechanism holds particular significance for insects with high metabolic rates, inherently generating substantial volumes of free radicals[5] Besides, the dietary intake profoundly influences the survival and vitality of *Apis mellifera* colonies, particularly noticeable during the spring season. Implementing proactive measures to mitigate oxidative stress effects can substantially bolster the resilience of *Apis mellifera* organisms[6]. This study aims to study the effect of molybdenum compounds on AOA in bees and in bee products.

#### VIII.2.1 Experimental protocols.

##### 1°) Apiary and conditions of feeding.

The antioxidant properties of the complexes **Na-$Mo_2O_4$-EDTA** and **Li-$Mo_2O_4$-EDTA** were evaluated at the apiary of the Institute of Zoology of the State University of Moldova. The apiary is located in a forested area not far from the city of Chisinau, Republic of Moldova (see Figure SVIII.5). The bees belonged to the *carpatica ecotype of Apis mellifera carnica*.

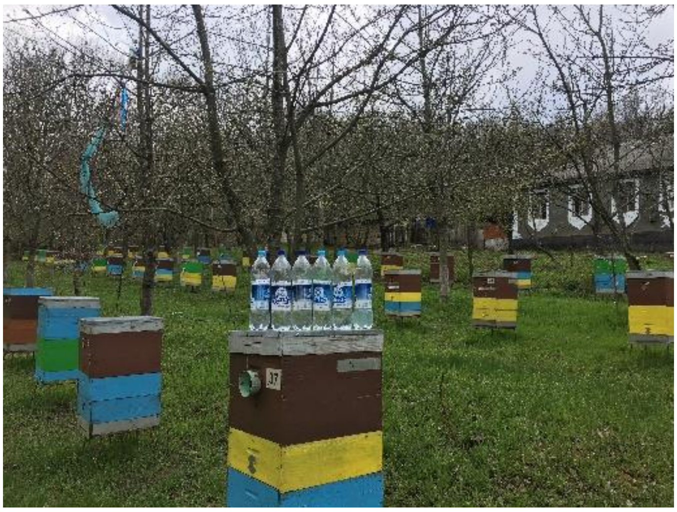

***Figure SVIII.5.*** *Experimental apiary of the institute of Zoology, Moldova.*

Experimental hives were fed for 14 days with 50% sugar syrup enriched or not with bio-active compounds, **Na-$Mo_2O_4$-EDTA** and **Li-$Mo_2O_4$-EDTA**. Aqueous solutions with a concentration of 1mg% (i.e. 1mg/100mL) were prepared and mixed with sugar syrup in a ratio of 60 mL to 1 L of syrup, i.e. 0.6 mg/L, and then administered directly to the beehives.

Colonies were fed in April, at a rate of 120 mL of the mixture for each frame interval populated with bees, every 2 days, for two weeks.

Test hives generally contain 10 frames. On average, 1 liter of syrup is given every 2 days for 14 days, which corresponds to around 7 liters of syrup supplemented with 0.6 mg/L, or around 4.2 mg of **Na-$Mo_2O_4$-EDTA** and **Li-$Mo_2O_4$-EDTA** per colony.

Sodium molybdate $Na_2MoO_4.2H_2O$ was also tested, for comparison, with a Mo content equivalent to that of the **Na-$Mo_2O_4$-EDTA** and **Li-$Mo_2O_4$-EDTA** compounds.

#### 2°) Sampling

The biological samples were registered at the apiary of the Republic of Moldova. Honey, beeswax, propolis, royal jelly, bee bread, larvae and workers bees were collected two weeks after the conclusion of the treatment, at the end of May. The assessment of antioxidant activity (AOA) of the tested compounds and beekeeping products was assessed at the Biological Invasions Research Center, Laboratory of Systematics and Molecular Phylogeny, Institute of Zoology, Moldova State University.

Samples destined for analysis were stored in sterile containers (food-grade plastic or glass), then carefully preserved and transported in portable freezers to the laboratory.

The honey was dissolved in a water-alcohol solution (1:1). Beeswax, propolis, royal jelly, bee bread, hemolymph of bees, and their larvae were dissolved in 96% ethanol and incubated in a thermostat at 25°C for a duration of 2 days. The biological material underwent examination at varying concentrations spanning from 0.01 to 100.00 mg/mL.

In this study, the two most frequently employed methods for evaluating the ability to scavenge free radicals, namely the $DPPH^{\bullet}$ and $ABTS^{\bullet+}$ assays, were conducted.

#### 3°) Anti-oxidant activity (AOA) determinations

The anti-oxidant activity can be evaluated by two methods, namely ABTS or DPPH as follows:

**$ABTS^{\bullet+}$ radical cation scavenging assay**

The antioxidant activity by the ABTS method was assessed according to the protocol described by Re et al. [7] with modifications. The $ABTS^{\bullet+}$ radical cations were generated by mixing a 7 mM solution of ABTS (2,20-azino-bis(3-ethylbenzothiazoline-6-sulphonic acid)) (Sigma) with a 2.45 mM solution of potassium persulfate ($K_2S_2O_8$) (Sigma) at 25°C in the dark for 12–20 hours. The resulting solution was then further diluted with 96% ethanol to achieve an absorbance of 0.7 ± 0.1 at 734 nm. Fresh working solution $ABTS^{\bullet+}$ was prepared for each assay.

Subsequently, 20 µL of different biological samples concentrations diluted in ethanol were allowed to react with 180 µL of the working solution of $ABTS^{\bullet+}$ for 30 min in the dark at 25°C. The absorbance was taken at 734 nm using a hybrid reader (Synergy H1, BioTek). Ethanol with glucose solution was used as control. Blank samples were run by solvent without $ABTS^{\bullet+}$.
The percent of inhibition (I %) of free radical cation production of $ABTS^{\bullet+}$ was calculated according to the following equation:

$$\boldsymbol{I}\,(\%) = \frac{\boldsymbol{Abs_{734\,nm_0}} - \boldsymbol{Abs_{734\,nm_1}}}{\boldsymbol{Abs_{734\,nm_0}}} \times \mathbf{100} \quad \mathbf{,where} \qquad (\mathbf{1})$$

**Abs** $_{734\,nm\,0}$ is the absorbance of the control solution;
**Abs** $_{734\,nm\,1}$ is the absorbance in the presence of sample solutions or standards.

**$DPPH^{\bullet}$ radical scavenging assay**

The 2,2-diphenyl-1-picrylhydrazyl (DPPH) assay was conducted following the method outlined by Brand-Williams et al. with some modifications [8]. The DPPH reagent was diluted with a methanolic solution to achieve an absorbance of 0.7 ± 0.1 at 517 nm.

Subsequently, 20 µL of various concentrations of biological samples diluted in ethanol were allowed to react with 180 µL of the working solution ($DPPH^{\bullet}$ reagent at 0.002% w/v in methanol) for 30 minutes in darkness at 25°C. The measurement was made by a hybrid reader (Synergy H1, BioTek). As a control, ethanol with glucose solution was utilized. Blank samples were run using solvent without $DPPH^{\bullet}$.

The percentage of inhibition (I %) of the $DPPH^{\bullet}$ free radical was calculated using the following equation:

$$\boldsymbol{I}\,(\%) = \frac{\boldsymbol{Abs_{517\,nm_0}} - \boldsymbol{Abs_{517\,nm_1}}}{\boldsymbol{Abs_{517\,nm_0}}} \times \mathbf{100} \quad \mathbf{,where} \qquad (\mathbf{2})$$

**Abs** $_{517\,nm\,0}$ is the absorbance of the control solution;
**Abs** $_{517\,nm\,1}$ is the absorbance in the presence of sample solutions or standards.

**Statistical Analysis**

The antioxidant assay results are presented as the percentage of inhibition (I %) of $DPPH^{\bullet}$ and $ABTS^{\bullet+}$ radicals. To gauge the efficacy of the experimental compounds on antioxidant activity (AOA), the half-maximal inhibitory concentration values ($IC_{50}$) were computed using the dose-response equation derived from the least squares fit method. All data are expressed as means ± standard deviation (SD). Statistical analysis was performed using BIOSTAT and GraphPad software to process and analyze the data.

##### VIII.2.2 Results.

The results of $IC_{50}$ values are depicted in Figures SVIII.6 to SVIII.12. The results are gathered in Table SVIII.2, while all the experimental data obtained with the ABTS and DPPH methods are given for each group and for Trolox as reference in Tables SVIII.3-SVIII.7

The antioxidant properties of hemolymph of bees and larvae are depicted in Figures SVIII.6 and SVIII.7, respectively.

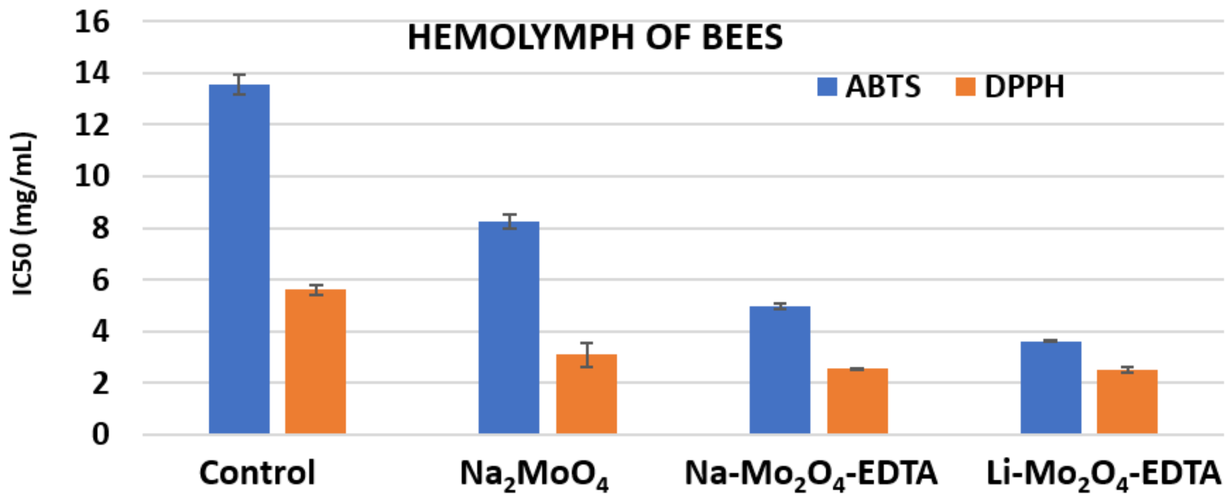


***Figure SVIII.6.*** *Antioxidant properties of hemolymph obtained from worker bees from the control group or fed with sodium molybdate* $Na_2MoO_4.2H_2O$ *or complexes* ***Na-*** $\boldsymbol{Mo_2O_4}$***-EDTA*** *and* ***Li-*** $\boldsymbol{Mo_2O_4}$***-EDTA.*** *Mean values ± SD for both methods ABTS and DDPH.*

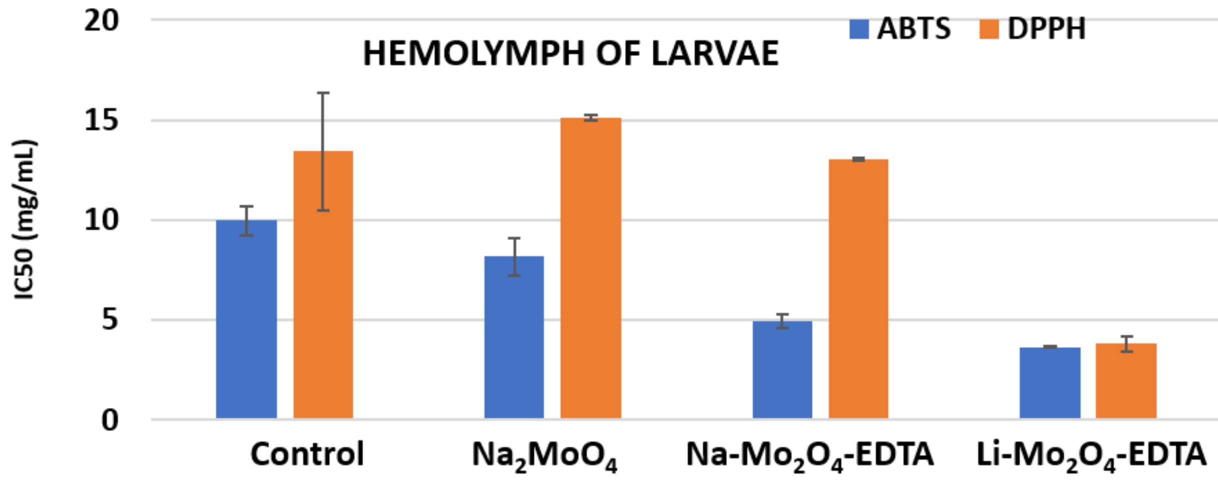


***Figure SVII.7.*** *Antioxidant properties of hemolymph obtained from larvae from the control group or fed with sodium molybdate* $Na_2MoO_4.2H_2O$ *or complexes* ***Na-*** $\boldsymbol{Mo_2O_4}$***-EDTA*** *and* ***Li-*** $\boldsymbol{Mo_2O_4}$***-EDTA.*** *Mean values ± SD for both methods ABTS and DDPH.*

Concerning the hemolymph of worker bees, $IC_{50}$ values obtained with control bees show low values in both methods, indicating a naturally good antioxidant activity. Interestingly, despite the fact that sodium molybdate $Na_2MoO_4$ cannot be chemically an antioxidant, the antioxidant activity (AOA) of the hemolymph of bees treated with this compound significantly increases, thus suggesting another mechanism. Finally, the AOA of hemolymph in bees increases a lot for both complexes tested **Na-$Mo_2O_4$-EDTA** and **Li-$Mo_2O_4$-EDTA**. $IC_{50}$ values are much lower than in control bees, both by the ABTS and DPPH methods. The AOA of both complexes appears of the same order even if it appears slightly better with **Na-$Mo_2O_4$-EDTA**.

In the case of larval hemolymph (Figure SVIII.7), the findings are similar for the control group and sodium molybdate, which shows no increase in AOA. **Li-$Mo_2O_4$-EDTA** complex shows a very significant increase in AOA in larvae for both methods. This increase is less for **Na-$Mo_2O_4$-EDTA** in the case of the DPPH method than for the controls, but is clearly improved by the ABTS method, which scavenges free radicals more effectively than the DPPH method.

The AOA of Honey samples are shown in Figure SVIII.8. The AOA appears similar for control, sodium molybdate and **Na-$Mo_2O_4$-EDTA** groups. On the contrary, the AOA appears strongly enhanced with **Li-$Mo_2O_4$-EDTA**. This result is surprising and could be due either to traces of feeding syrup in the honey for this group of hives or due to an action of this complex on the enzymes produced in their crop to produce honey.

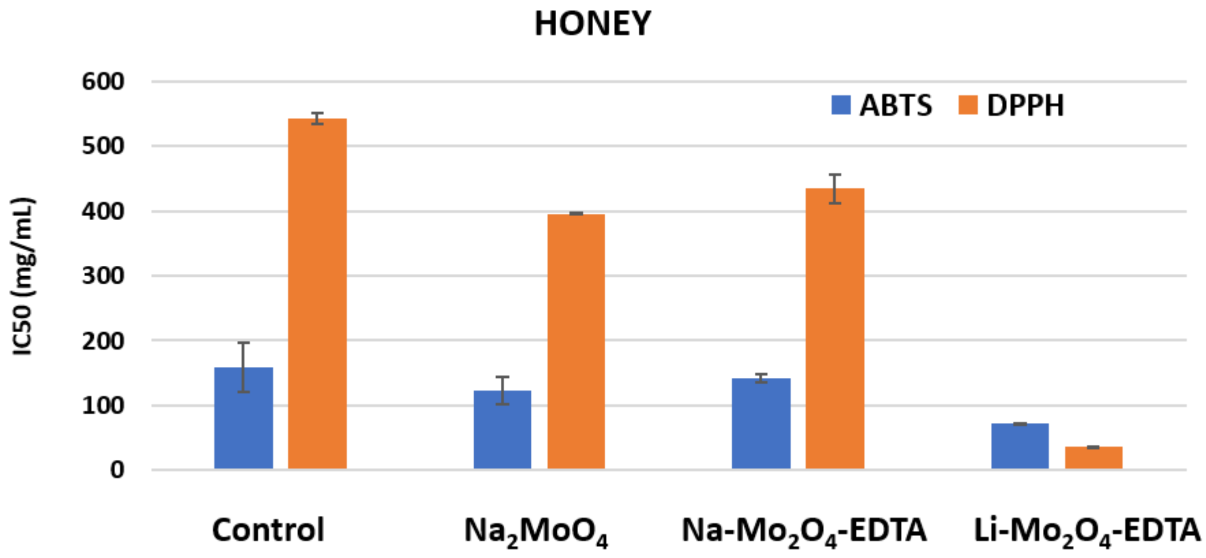


***Figure SVIII.8.*** *Antioxidant properties of honey obtained from hives from the control group or fed with sodium molybdate $Na_2MoO_4.2H_2O$ or complexes **Na-$Mo_2O_4$-EDTA** and **Li-$Mo_2O_4$-EDTA.** Mean values ± SD for both methods ABTS and DDPH.*

The AOA of Royal Jelly and from Bee wax taken from the 4 groups are shown in Figures SVIII.9 and SVIII.10, respectively. The AOA is similar in all cases.

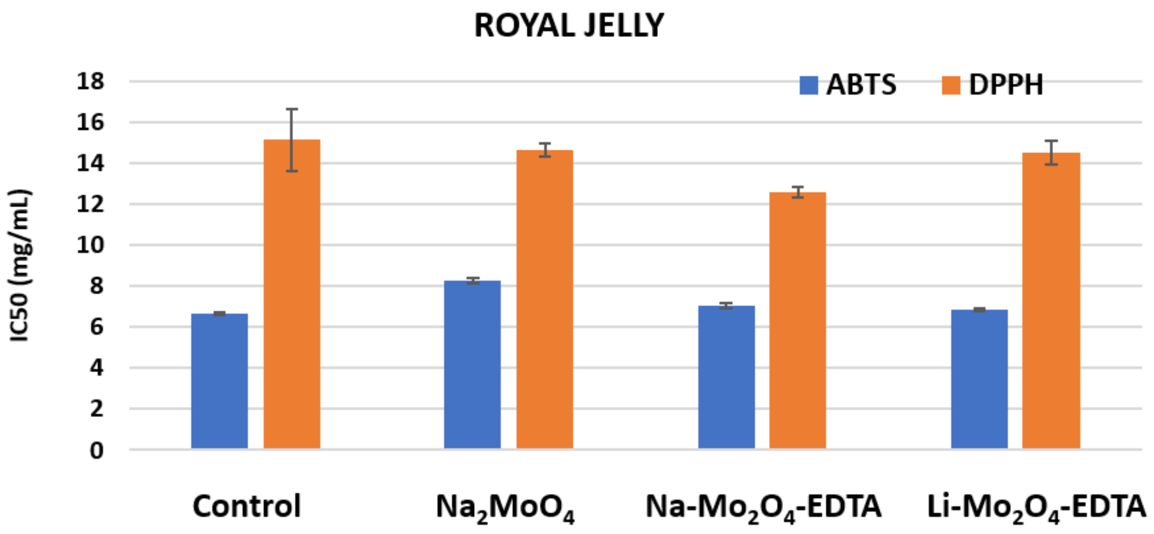


***Figure SVIII.9.*** *Antioxidant properties of royal jelly obtained from hives from the control group or fed with sodium molybdate* $Na_2MoO_4.2H_2O$ *or complexes* ***Na-Mo₂O₄-EDTA*** *and* ***Li-Mo₂O₄-EDTA.*** *Mean values ± SD for both methods ABTS and DDPH.*

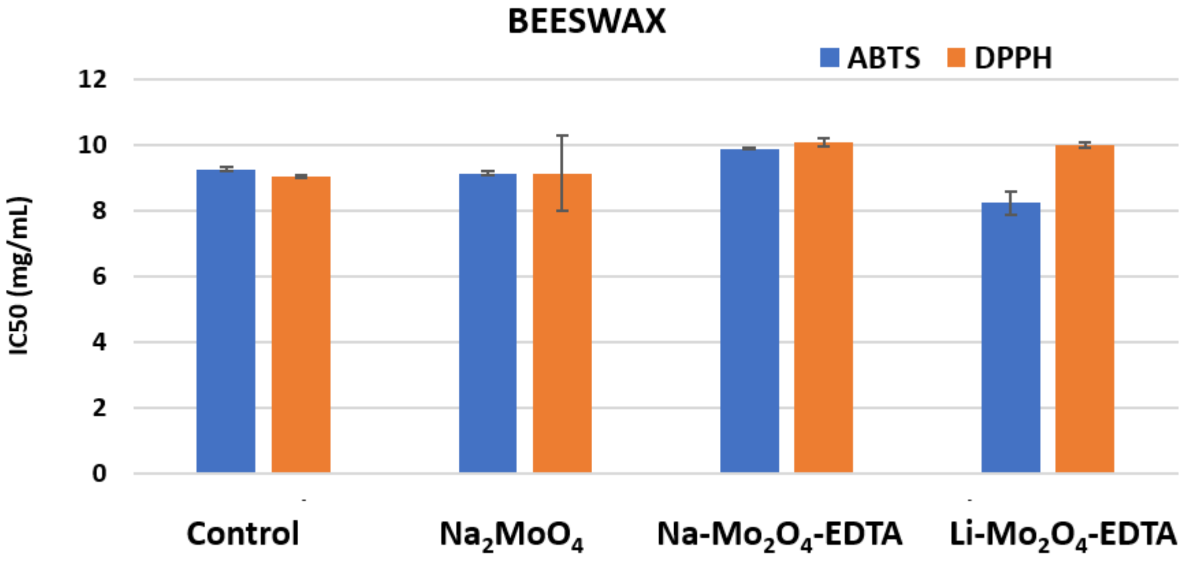


***Figure SVIII.10.*** *Antioxidant properties of beeswax obtained from hives from the control group or fed with sodium molybdate* $Na_2MoO_4.2H_2O$ *or complexes* ***Na-Mo₂O₄-EDTA*** *and* ***Li-Mo₂O₄-EDTA.*** *Mean values ± SD for both methods ABTS and DDPH.*

Similarly, the AOA of propolis (Figure SVIII.11) appears to be very high in all 4 groups. However, AOA activity was lower in the group fed with sodium molybdate, while it appears increased with complex **$Na\text{-}Mo_2O_4\text{-}EDTA$**, especially with DPPH method.

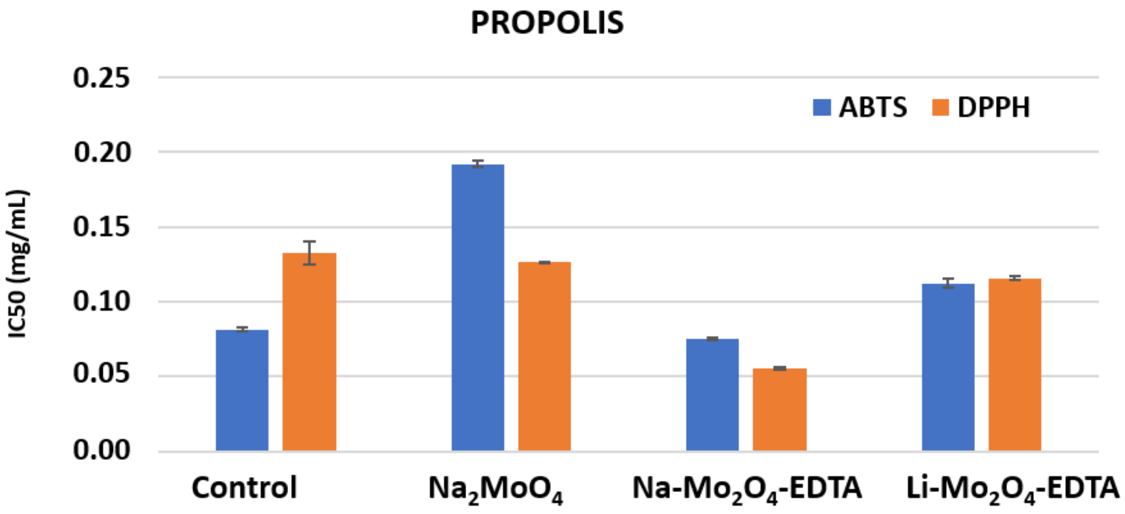


***Figure SVIII.11.*** *Antioxidant properties of propolis obtained from hives from the control group or fed with sodium molybdate $Na_2MoO_4.2H_2O$ or complexes* ***$Na-Mo_2O_4$-EDTA*** *and* ***$Li-Mo_2O_4$-EDTA.*** *Mean values ± SD for both methods ABTS and DDPH.*

Finally, as seen in Figure SVIII.12, both complexes significantly amplified the antioxidant activity of bee bread, showcasing remarkable efficacy in enhancing its overall antioxidative potential.

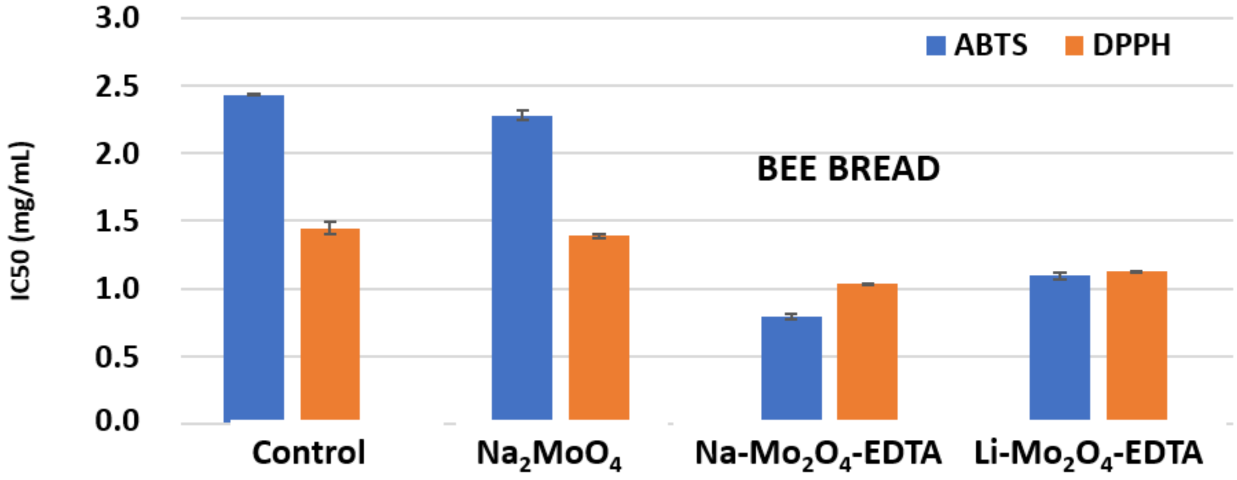


***Figure SVIII.12.*** *Antioxidant properties of bee bread obtained from hives from the control group or fed with sodium molybdate $Na_2MoO_4.2H_2O$ or complexes* ***$Na-Mo_2O_4$-EDTA*** *and* ***$Li-Mo_2O_4$-EDTA.*** *Mean values ± SD for both methods ABTS and DDPH.*

**Table SVIII.2.** Antioxidant activity of the hemolymph of honey bee workers, larvae and products

| Group | Type of sample | ABTS | | DPPH | |
|---|---|---|---|---|---|
| | | $IC_{50}$ (mg/mL) | SD (mg/mL) | $IC_{50}$ (mg/mL) | SD (mg/mL) |
| **Control** | HEMOLYMPH OF BEES | 13.56 | 0.37 | 5.61 | 0.19 |
| | HEMOLYMPH OF LARVAE | 9.95 | 0.70 | 13.44 | 2.95 |
| | HONEY | 159.40 | 8.91 | 543.00 | 38.18 |
| | BEE BREAD | 2.433 | 0.004 | 1.45 | 0.04 |
| | BEESWAX | 9.26 | 0.07 | 9.04 | 0.04 |
| | PROPOLIS | 0.082 | 0.001 | 0.13 | 0.01 |
| | ROYAL JELLY | 6.65 | 0.05 | 15.14 | 1.51 |
| **$Na_2MoO_4.2H_2O$** | HEMOLYMPH OF BEES | 8.26 | 0.26 | 3.4 | 0.5 |
| | HEMOLYMPH OF LARVAE | 8.15 | 0.92 | 15.12 | 0.16 |
| | HONEY | 123.20 | 1.13 | 395.50 | 20.65 |
| | BEE BREAD | 2.28 | 0.04 | 1.39 | 0.01 |
| | BEESWAX | 9.12 | 0.06 | 9.13 | 1.14 |
| | PROPOLIS | 0.19 | 0.01 | 0.13 | 0.01 |
| | ROYAL JELLY | 8.26 | 0.14 | 14.63 | 0.33 |
| **$Na$-$Mo_2O_4$-EDTA** | HEMOLYMPH OF BEES | 4.95 | 0.11 | 2.54 | 0.01 |
| | HEMOLYMPH OF LARVAE | 4.93 | 0.36 | 13.01 | 0.06 |
| | HONEY | 141.95 | 22.27 | 435.00 | 7.07 |
| | BEE BREAD | 0.79 | 0.02 | 1.03 | 0.01 |
| | BEESWAX | 9.89 | 0.01 | 10.09 | 0.12 |
| | PROPOLIS | 0.075 | 0.001 | 0.056 | 0.001 |

| | | | | | |
|---|---|---|---|---|---|
| | ROYAL JELLY | 7.06 | 0.12 | 12.60 | 0.26 |
| **Li-$Mo_2O_4$-EDTA** | HEMOLYMPH OF BEES | 3.63 | 0.05 | 2.51 | 0.12 |
| | HEMOLYMPH OF LARVAE | 3.65 | 0.01 | 3.80 | 0.39 |
| | HONEY | 71.64 | 0.58 | 36.00 | 1.10 |
| | BEE BREAD | 1.09 | 0.02 | 1.127 | 0.001 |
| | BEESWAX | 8.24 | 0.35 | 10.00 | 0.07 |
| | PROPOLIS | 0.112 | 0.003 | 0.116 | 0.001 |
| | ROYAL JELLY | 6.87 | 0.06 | 14.52 | 0.59 |

**Table SVIII.3**. Antioxidant activity of the hemolymph of honey bee workers, larvae and products in control group. $IC_{50}$ are given in mg/mL

| **Group** | **Type of sample** | **Concentration of reactant C (mg/mL)** | **ABTS** | | | | | **DPPH** | | | | |
|---|---|---|---|---|---|---|---|---|---|---|---|---|
| | | | **Abs (734 nm)** | **% inh** | **SD (%)** | **$IC_{50}$ (mg/mL)** | **SD (mg/mL)** | **Abs (517 nm)** | **% inh** | **SD (%)** | **$IC_{50}$ (mg/mL)** | **SD (mg/mL)** |
| Control | HEMOLYMPH OF BEES | 100 | 0.156 | **81.43** | **2.02** | 13.56 | 0.37 | 0.083 | **90.18** | **1.09** | 5.61 | 0.19 |
| | | 10 | 0.483 | **42.56** | **2.27** | | | 0.241 | **71.37** | **1.26** | | |
| | | 1 | 0.702 | **16.49** | **0.42** | | | 0.795 | **5.42** | **0.25** | | |
| | | 0.1 | 0.805 | **4.17** | **0.51** | | | 0.834 | **0.71** | **0.51** | | |
| Control | HEMOLYMPH OF LARVAE | 50 | 0.061 | **92.74** | **4.55** | 9.95 | 0.70 | 0.035 | **95.89** | **1.77** | 13.44 | 2.95 |
| | | 5 | 0.627 | **25.36** | **1.68** | | | 0.762 | **9.29** | **4.88** | | |
| | | 0.5 | 0.756 | **10.06** | **0.08** | | | 0.784 | **6.73** | **1.09** | | |
| | | 0.05 | 0.836 | **0.54** | **0.08** | | | 0.805 | **4.23** | **1.77** | | |

| | | | | | | | | | | | | |
|---|---|---|---|---|---|---|---|---|---|---|---|---|
| Control | HONEY | 100 | 0.506 | **39.76** | **0.51** | 159.40 | 8.91 | 0.562 | **33.15** | **0.08** | 543.00 | 38.18 |
| | | 10 | 0.783 | **6.85** | **0.59** | | | 0.708 | **15.71** | **0.17** | | |
| | | 1 | 0.814 | **3.15** | **0.42** | | | 0.723 | **13.99** | **0.08** | | |
| | | 0.1 | 0.825 | **1.79** | **0.67** | | | 0.759 | **9.64** | **1.18** | | |
| Control | BEE BREAD | 50 | 0.054 | **93.57** | **0.34** | 2.433 | 0.004 | 0.027 | **96.85** | **0.08** | 1.45 | 0.04 |
| | | 5 | 0.369 | **56.07** | **0.34** | | | 0.126 | **85.00** | **0.17** | | |
| | | 0.5 | 0.592 | **29.58** | **0.42** | | | 0.686 | **18.39** | **0.93** | | |
| | | 0.05 | 0.776 | **7.68** | **0.42** | | | 0.802 | **4.52** | **1.85** | | |
| Control | BEESWAX | 50 | 0.064 | **92.44** | **0.42** | 9.26 | 0.07 | 0.019 | **97.740** | **0.001** | 9.04 | 0.04 |
| | | 5 | 0.597 | **28.93** | **0.67** | | | 0.644 | **23.33** | **0.51** | | |
| | | 0.5 | 0.770 | **8.33** | **0.17** | | | 0.754 | **10.30** | **1.26** | | |
| | | 0.05 | 0.838 | **0.24** | **0.17** | | | 0.801 | **4.70** | **0.25** | | |
| Control | PROPOLIS | 5 | 0.060 | **92.86** | **1.52** | 0.082 | 0.001 | 0.093 | **88.99** | **0.42** | 0.13 | 0.01 |
| | | 0.5 | 0.292 | **65.30** | **0.08** | | | 0.145 | **82.74** | **1.52** | | |
| | | 0.05 | 0.404 | **51.90** | **0.17** | | | 0.633 | **24.64** | **0.51** | | |
| | | 0.005 | 0.736 | **12.44** | **0.25** | | | 0.773 | **8.04** | **1.43** | | |
| Control | ROYAL JELLY | 50 | 0.046 | **94.58** | **0.93** | 6.65 | 0.05 | 0.051 | **93.99** | **0.08** | 15.14 | 1.51 |
| | | 5 | 0.512 | **39.05** | **0.34** | | | 0.755 | **10.12** | **0.84** | | |
| | | 0.5 | 0.732 | **12.86** | **0.34** | | | 0.762 | **9.35** | **0.59** | | |
| | | 0.05 | 0.827 | **1.55** | **0.17** | | | 0.796 | **5.24** | **1.01** | | |

**Table SVIII.4.** Antioxidant activity of the hemolymph of honey bee workers, larvae and products after feeding with $Na_2MoO_4 \cdot 2H_2O$ used as reference. $IC_{50}$ are given in mg/mL

| Group | Type of sample | Concentration of reactant C (mg/mL) | ABTS | | | | | DPPH | | | | |
|---|---|---|---|---|---|---|---|---|---|---|---|---|
| | | | Abs (734 nm) | % inh | SD (%) | $IC_{50}$ (mg/mL) | SD (mg/mL) | Abs (517 nm) | % inh | SD (%) | $IC_{50}$ (mg/mL) | SD (mg/mL) |
| **$Na_2MoO_4 \cdot 2H_2O$** | HEMOLYMPH OF BEES | 100 | 0.094 | **88.8** | **0.2** | 8.3 | 0.2 | 0.049 | **94.17** | **3.20** | 3.10 | 0.47 |
| | | 10 | 0.476 | **43.3** | **0.3** | | | 0.155 | **81.61** | **6.5** | | |
| | | 0.1 | 0.592 | **29.52** | **0.67** | | | 0.675 | **19.64** | **0.84** | | |
| | | 0.01 | 0.785 | **6.55** | **0.17** | | | 0.798 | **5.06** | **0.76** | | |
| **$Na_2MoO_4 \cdot 2H_2O$** | HEMOLYMPH OF LARVAE | 50 | 0.025 | **97.08** | **3.96** | 8.15 | 0.92 | 0.069 | **91.79** | **0.00** | 15.12 | 0.16 |
| | | 5 | 0.597 | **28.93** | **0.67** | | | 0.759 | **9.64** | **0.34** | | |
| | | 0.5 | 0.770 | **8.33** | **0.67** | | | 0.806 | **4.11** | **0.08** | | |
| | | 0.05 | 0.831 | **1.07** | **0.51** | | | 0.808 | **3.81** | **0.17** | | |
| **$Na_2MoO_4 \cdot 2H_2O$** | HONEY | 100 | 0.463 | **44.88** | **0.17** | 123.20 | 1.13 | 0.527 | **37.32** | **0.42** | 395.50 | 20.65 |
| | | 10 | 0.784 | **6.73** | **0.08** | | | 0.706 | **16.01** | **0.42** | | |
| | | 1 | 0.810 | **3.63** | **0.08** | | | 0.773 | **7.98** | **1.01** | | |
| | | 0.1 | 0.827 | **1.55** | **0.34** | | | 0.783 | **6.85** | **0.93** | | |
| **$Na_2MoO_4 \cdot 2H_2O$** | BEE BREAD | 50 | 0.039 | **95.42** | **0.08** | 2.28 | 0.04 | 0.039 | **95.42** | **0.08** | 1.39 | 0.01 |
| | | 5 | 0.359 | **57.26** | **0.17** | | | 0.114 | **86.43** | **0.01** | | |
| | | 0.5 | 0.593 | **29.40** | **0.51** | | | 0.684 | **18.63** | **0.42** | | |
| | | 0.05 | 0.769 | **8.51** | **0.59** | | | 0.789 | **6.13** | **0.08** | | |
| **$Na_2MoO_4 \cdot 2H_2O$** | BEESWAX | 50 | 0.051 | **93.99** | **0.25** | 9.12 | 0.06 | 0.021 | **97.56** | **2.95** | 9.13 | 1.14 |
| | | 5 | 0.604 | **28.15** | **0.59** | | | 0.653 | **22.32** | **0.08** | | |
| | | 0.5 | 0.765 | **8.99** | **0.59** | | | 0.755 | **10.12** | **0.34** | | |

| | | | | | | | | | | | | |
|---|---|---|---|---|---|---|---|---|---|---|---|---|
| | | 0.05 | 0.811 | **3.45** | **0.51** | | | 0.795 | **5.42** | **0.08** | | |
| **$Na_2MoO_4 \cdot 2H_2O$** | PROPOLIS | 5 | 0.050 | **94.05** | **0.17** | 0.19 | 0.00 | 0.015 | **98.27** | **0.25** | 0.13 | 0.00 |
| | | 0.5 | 0.358 | **57.44** | **0.25** | | | 0.139 | **83.51** | **0.08** | | |
| | | 0.05 | 0.543 | **35.42** | **0.59** | | | 0.637 | **24.23** | **0.08** | | |
| | | 0.005 | 0.771 | **8.21** | **0.34** | | | 0.756 | **10.00** | **0.17** | | |
| **$Na_2MoO_4 \cdot 2H_2O$** | ROYAL JELLY | 50 | 0.057 | **93.27** | **0.25** | 8.26 | 0.14 | 0.090 | **89.35** | **0.59** | 14.63 | 0.33 |
| | | 5 | 0.566 | **32.68** | **0.42** | | | 0.728 | **13.39** | **0.25** | | |
| | | 0.5 | 0.747 | **11.07** | **0.67** | | | 0.776 | **7.62** | **0.17** | | |
| | | 0.05 | 0.821 | **2.26** | **0.34** | | | 0.818 | **2.68** | **0.59** | | |

**Table SVIII.5.** Antioxidant activity of the hemolymph of honey bee workers, larvae and products after feeding with complex Na-$Mo_2O_4$-EDTA. $IC_{50}$ are given in mg/mL

| **Group** | **Type of sample** | **Concentration of reactant C (mg/mL)** | **ABTS** | | | | | **DPPH** | | | | |
|---|---|---|---|---|---|---|---|---|---|---|---|---|
| | | | **Abs (734 nm)** | **% inh** | **SD (%)** | **$IC_{50}$ (mg/mL)** | **SD (mg/mL)** | **Abs (517 nm)** | **% inh** | **SD (%)** | **$IC_{50}$ (mg/mL)** | **SD (mg/mL)** |
| **Na-$Mo_2O_4$-EDTA** | HEMOLYMPH OF BEES | 100 | 0.197 | **76.55** | **0.01** | 4.95 | 0.11 | 0.049 | **94.23** | **2.44** | 2.54 | 0.01 |
| | | 10 | 0.389 | **53.75** | **0.08** | | | 0.085 | **89.88** | **0.17** | | |
| | | 1 | 0.491 | **41.55** | **0.67** | | | 0.684 | **18.63** | **0.08** | | |
| | | 0.1 | 0.736 | **12.38** | **0.17** | | | 0.765 | **8.93** | **0.34** | | |
| **Na-$Mo_2O_4$-EDTA** | HEMOLYMPH OF LARVAE | 50 | 0.037 | **95.60** | **5.89** | 4.93 | 0.36 | 0.073 | **91.37** | **0.25** | 13.01 | 0.06 |
| | | 5 | 0.462 | **45.00** | **1.01** | | | 0.708 | **15.77** | **0.08** | | |
| | | 0.5 | 0.692 | **17.62** | **0.17** | | | 0.753 | **10.36** | **0.51** | | |
| | | 0.05 | 0.795 | **5.42** | **0.42** | | | 0.776 | **7.62** | **0.17** | | |

| | | | | | | | | | | | | |
|---|---|---|---|---|---|---|---|---|---|---|---|---|
| **Na-$Mo_2O_4$-EDTA** | HONEY | 100 | 0.473 | **43.69** | **2.69** | 141.95 | 22.27 | 0.552 | **34.29** | **1.85** | 435.00 | 7.07 |
| | | 10 | 0.753 | **10.36** | **0.17** | | | 0.715 | **14.94** | **0.59** | | |
| | | 1 | 0.814 | **3.15** | **0.25** | | | 0.722 | **14.05** | **0.17** | | |
| | | 0.1 | 0.789 | **6.07** | **0.34** | | | 0.740 | **11.90** | **0.67** | | |
| **Na-$Mo_2O_4$-EDTA** | BEE BREAD | 50 | 0.045 | **94.70** | **0.08** | 0.79 | 0.02 | 0.073 | **91.31** | **0.17** | 1.030 | 0.001 |
| | | 5 | 0.357 | **57.56** | **0.08** | | | 0.128 | **84.82** | **0.08** | | |
| | | 0.5 | 0.409 | **51.37** | **0.42** | | | 0.580 | **30.95** | **0.00** | | |
| | | 0.05 | 0.656 | **21.96** | **0.42** | | | 0.754 | **10.24** | **0.17** | | |
| **Na-$Mo_2O_4$-EDTA** | BEESWAX | 50 | 0.053 | **93.69** | **0.00** | 9.89 | 0.01 | 0.045 | **94.64** | **0.51** | 10.09 | 0.12 |
| | | 5 | 0.632 | **24.82** | **0.08** | | | 0.647 | **23.04** | **0.08** | | |
| | | 0.5 | 0.775 | **7.80** | **0.59** | | | 0.738 | **12.14** | **0.00** | | |
| | | 0.05 | 0.807 | **3.99** | **0.25** | | | 0.763 | **9.23** | **0.25** | | |
| **Na-$Mo_2O_4$-EDTA** | PROPOLIS | 5 | 0.050 | **94.11** | **0.08** | 0.075 | 0.001 | 0.057 | **93.21** | **2.36** | 0.056 | 0.001 |
| | | 0.5 | 0.355 | **57.80** | **0.25** | | | 0.139 | **83.45** | **0.34** | | |
| | | 0.05 | 0.370 | **56.01** | **0.25** | | | 0.421 | **49.88** | **0.00** | | |
| | | 0.005 | 0.691 | **17.74** | **0.34** | | | 0.743 | **11.55** | **0.17** | | |
| **Na-$Mo_2O_4$-EDTA** | ROYAL JELLY | 50 | 0.058 | **93.15** | **0.08** | 7.06 | 0.12 | 0.051 | **93.99** | **0.25** | 12.60 | 0.26 |
| | | 5 | 0.526 | **37.44** | **0.42** | | | 0.724 | **13.81** | **0.51** | | |
| | | 0.5 | 0.729 | **13.27** | **0.25** | | | 0.746 | **11.19** | **0.34** | | |
| | | 0.05 | 0.815 | **2.98** | **0.17** | | | 0.788 | **6.25** | **0.08** | | |

**Table SVIII.6.** Antioxidant activity of the hemolymph of honey bee workers, larvae and products after feeding with complex Li-$Mo_2O_4$-EDTA. $IC_{50}$ are given in mg/mL

| Group | Type of sample | Concentration of reactant C (mg/mL) | ABTS | | | | | DPPH | | | | |
|---|---|---|---|---|---|---|---|---|---|---|---|---|
| | | | Abs (734 nm) | % inh | SD (%) | $IC_{50}$ (mg/mL) | SD (mg/mL) | Abs (517 nm) | % inh | SD (%) | $IC_{50}$ (mg/mL) | SD (mg/mL) |
| **Li-$Mo_2O_4$-EDTA** | HEMOLYMPH OF BEES | 100 | 0.188 | **77.68** | **0.76** | 3.63 | 0.05 | 0.045 | **94.38** | **0.00** | 2.51 | 0.12 |
| | | 10 | 0.392 | **53.39** | **0.08** | | | 0.085 | **89.44** | **1.68** | | |
| | | 1 | 0.429 | **48.99** | **0.25** | | | 0.643 | **19.69** | **0.09** | | |
| | | 0.1 | 0.723 | **13.99** | **0.08** | | | 0.758 | **5.25** | **0.35** | | |
| **Li-$Mo_2O_4$-EDTA** | HEMOLYMPH OF LARVAE | 50 | 0.062 | **92.62** | **0.67** | 3.65 | 0.01 | 0.060 | **92.56** | **5.04** | 3.80 | 0.39 |
| | | 5 | 0.395 | **53.04** | **0.08** | | | 0.340 | **57.50** | **2.12** | | |
| | | 0.5 | 0.680 | **19.11** | **0.25** | | | 0.713 | **10.88** | **0.18** | | |
| | | 0.05 | 0.784 | **6.67** | **0.34** | | | 0.775 | **3.19** | **0.09** | | |
| **Li-$Mo_2O_4$-EDTA** | HONEY | 100 | 0.345 | **58.99** | **0.25** | 71.64 | 0.58 | 0.224 | **72.00** | **1.41** | 36.00 | 1.10 |
| | | 10 | 0.758 | **9.82** | **0.08** | | | 0.625 | **21.94** | **0.44** | | |
| | | 1 | 0.788 | **6.19** | **0.51** | | | 0.717 | **10.38** | **0.01** | | |
| | | 0.1 | 0.788 | **6.19** | **0.51** | | | 0.746 | **6.81** | **0.27** | | |
| **Li-$Mo_2O_4$-EDTA** | BEE BREAD | 50 | 0.040 | **95.30** | **0.08** | 1.09 | 0.02 | 0.070 | **91.25** | **5.13** | 1.127 | 0.001 |
| | | 5 | 0.346 | **58.87** | **0.08** | | | 0.104 | **87.00** | **0.53** | | |
| | | 0.5 | 0.464 | **44.82** | **0.25** | | | 0.587 | **26.69** | **0.09** | | |
| | | 0.05 | 0.693 | **17.56** | **0.59** | | | 0.745 | **6.94** | **0.27** | | |
| **Li-$Mo_2O_4$-EDTA** | BEESWAX | 50 | 0.065 | **92.26** | **3.03** | 8.24 | 0.35 | 0.036 | **95.50** | **0.18** | 10.00 | 0.07 |
| | | 5 | 0.560 | **33.39** | **0.08** | | | 0.623 | **22.19** | **0.09** | | |
| | | 0.5 | 0.751 | **10.60** | **0.34** | | | 0.708 | **11.50** | **0.18** | | |
| | | 0.05 | 0.778 | **7.44** | **0.42** | | | 0.749 | **6.44** | **0.09** | | |

| | | | | | | | | | | | | |
|---|---|---|---|---|---|---|---|---|---|---|---|---|
| **Li-$Mo_2O_4$-EDTA** | PROPOLIS | 5 | 0,044 | **94.76** | **0.17** | 0.112 | 0.003 | 0.033 | **95.94** | **0.09** | 0.116 | 0.001 |
| | | 0.5 | 0.346 | **58.81** | **0.00** | | | 0.137 | **82.94** | **0.27** | | |
| | | 0.05 | 0.455 | **45.83** | **0.34** | | | 0.572 | **28.50** | **0.18** | | |
| | | 0.005 | 0.712 | **15.30** | **0.42** | | | 0.741 | **7.44** | **0.09** | | |
| **Li-$Mo_2O_4$-EDTA** | ROYAL JELLY | 50 | 0.049 | **94.23** | **0.08** | 6.87 | 0.06 | 0.067 | **91.63** | **1.41** | 14.52 | 0.59 |
| | | 5 | 0.519 | **38.21** | **0.17** | | | 0.710 | **11.25** | **3.01** | | |
| | | 0.5 | 0.736 | **12.38** | **0.34** | | | 0.727 | **9.19** | **0.62** | | |
| | | 0.05 | 0.806 | **4.05** | **0.34** | | | 0.764 | **4.56** | **0.80** | | |

**Table SVIII.7.** Antioxidant activity of Trolox, used as reference. $IC_{50}$ are given in µM.

| **TROLOX** | **C (uM)** | **ABTS** | | | | | **DPPH** | | | | |
|---|---|---|---|---|---|---|---|---|---|---|---|
| | | **Abs (734 nm)** | **% inh** | **SD (%)** | **$IC_{50}$ (uM)** | **SD (uM)** | **Abs (517 nm)** | **% inh** | **SD (%)** | **$IC_{50}$ (uM)** | **SD (uM)** |
| | 100.00 | 0.153 | 81.73 | 0.25 | **23.91** | **0.30** | 0.218 | 73.99 | 1.26 | **37.72** | **0.91** |
| | 10.00 | 0.600 | 28.57 | 0.17 | | | 0.679 | 19.11 | 0.42 | | |
| | 1.00 | 0.805 | 4.11 | 0.42 | | | 0.803 | 4.35 | 0.25 | | |

#### VIII.2.3 Discussion / conclusions

The results obtained from this experiments evidence that:

- The two complexes **Na-$Mo_2O_4$-EDTA** and **Li-$Mo_2O_4$-EDTA** provoke an important increase of the AOA of the hemolymph of honey bee workers and larvae, which means that these complexes act on the organisms of both larvae and adults as antioxidants, inhibiting the activity of $ABTS^{\bullet+}$ and $DPPH^{\bullet}$ free radicals.

- Bee products have a very high AOA. The use of **Na-$Mo_2O_4$-EDTA** and **Li-$Mo_2O_4$-EDTA** has a moderate or no effect on the AOA of beeswax and royal jelly. However, both complexes **Na-$Mo_2O_4$-EDTA** and **Li-$Mo_2O_4$-EDTA** exhibits strong effects on the AOA of beebread, **Na-$Mo_2O_4$-EDTA** has a significant effect on the increase of AOA of propolis and the **Li-$Mo_2O_4$-EDTA** complex appears to significantly increase the AOA of honey.

Bee products usually exhibit a high antioxidant activity (AOA), which is correlated with the concentration of phenolic compounds, amino acids, peptides, proteins, organic acids, enzymes, vitamins, minerals, superoxide dismutase, catalase, glutathione [9]. Among bee products, propolis emerges as an exceptionally potent antioxidant, boasting the highest concentrations of phenols and flavonoids, closely followed by pollen/bee bread and royal jelly Our assessment of AOA in bee products within the control group consistently aligns with the reported data from literature data [10].

In response to the escalating interest in utilizing bioactive natural compounds with pronounced antioxidant properties to enhance health and mitigate certain diseases, our studies revealed that our Mo complexes allows augmenting the antioxidant capacity of *Apis mellifera* bee products, notably beebread, honey, and propolis. Remarkably, the AOA exhibited by these bee products often surpasses that of a water-soluble analog of vitamin E - Trolox, commonly employed as a standard antioxidant in biochemical techniques targeting oxidative stress reduction.
